# Single cell RNA-seq identifies developing corneal cell fates in the human cornea organoid

**DOI:** 10.1101/2021.12.23.471999

**Authors:** George Maiti, Maithe Rocha Monteiro de Barros, Nan Hu, Mona Roshan, Karl J Wahlin, Shukti Chakravarti

## Abstract

The cornea is a protective and refractive barrier in the eye crucial for vision. Understanding the human cornea in health, disease and cell-based treatments can be greatly advanced with cornea organoids developed in culture from induced pluripotent stem cells. While a limited number of studies have investigated the single-cell transcriptomic composition of the human cornea, its organoids have not been examined similarly. Here we elucidated the transcriptomic cell fate map of 4 month-old human cornea organoids and the central cornea from three donors. The organoids harbor cell clusters representing corneal epithelium, stroma and endothelium with sub populations that capture signatures of early developmental states. Unlike the adult cornea where the largest cell population is stromal, the organoids develop almost equal proportion of the three major cell types. These corneal organoids offer a three-dimensional platform to model corneal diseases and integrated responses of the different cell types to treatments.

**Teaser:** Transcriptomic Map of Cornea Organoid and Human Cornea

## Introduction

The cornea is the outermost transparent layer of the eye that serves as a protective barrier, and together with the lens refracts light onto the retina for visual processing (*1, 2*). The 3^rd^ leading cause of blindness globally is due to disorders of the cornea, with surgical transplantation of donor cornea often the only treatment option (*3*). An estimated 12.7 million people need cornea transplantation, and only ~1 in 70 of the needs are addressed worldwide. Consequently, there is a great interest in cell-based regenerative approaches to develop treatments for blinding corneal diseases. *A priori*, the cornea appears to be a simple tissue of three major cell types - a stratified epithelium above a basement membrane and an acellular Bowman’s layer, keratocytes, a specialized fibroblast-like cell type embedded in the central stroma of highly organized extracellular matrix that it produces, and an innermost single-cell layered endothelium that lies under the Descemet’s membrane, a basement membrane produced by the endothelium (*4, 5*). However, it is increasingly appreciated that within each cell type there is considerable heterogeneity, as well as subgroups of cells with stem, progenitor and transit amplifying qualities. Recognizing this heterogeneity, treatments for epithelial disorders, early on, have used marker-specific assessment of stemness in limbal epithelial cells (*6*). Stromal and endothelial cell-based treatments are also under development (*7, 8*). Comprehensive phenotyping of corneal cells by single cell RNA sequencing can lead to better characterization of cell populations for such cell-based therapies. Yet, there are just a few reported examples of single cell RNA sequence analyses of the cornea, with a majority of these focused on the limbal epithelial region, and very few on the central cornea (*9–13*). Here we performed single cell RNA sequencing of the central human cornea to determine its epithelial, stromal and endothelial cellular landscape. In addition, we performed scRNA-seq on human cornea organoids, and compared the organoid single cell transcriptome with that of the adult human cornea.

Organoid technology, which represents the study of 3D tissues developed from stem cells, has revolutionized the study of developmental stages and disease pathogenesis in tissue culture dishes. A majority of these, including our cornea organoid used induced pluripotent stem cells (iPSC). Thus, gut (*14*), liver (*15*), retina (*16–18*), cornea organoids (*19*) and mini-cornea organoids (*20*) have been developed from iPSC. Another study reported development of multiple cell linages of the eye from iPSC grown in 2D cultures. Multipotent somatic cells have also been used to generate corneal limbus (*21*) and lacrimal gland organoids (*22*). ScRNA-seq was used in the latter to address cellular heterogeneity of lacrimal glands (*22*). We performed scRNA-seq of iPSC-derived cornea organoids to elucidate its cellular complexity and relatedness to the human cornea. In our previous study we reported the presence of major epithelial, stromal and endothelial markers by immunohistology, and the presence of collagen fibrils by TEM in the cornea organoid without further characterization of its cellular constituents (*19, 23*). Thus far, modelling of corneal diseases to investigate underlying mechanisms *in vitro* have largely relied on individual cell types, such as the epithelium and the stroma for keratoconus (*24, 25*), and cultured endothelial cells for Fuchs endothelial dystrophy (*26*). These 2D cell culture models, though invaluable, lack information about tissue development and the consequences of interactions between cell types. Clearly, there is an unmet need for 3D-corneal organoids that harbor some of the cellular complexity and functionality of the human cornea.

We compared the single cell transcriptome of the central human cornea and iPSC-derived cornea organoid keeping key features of the cornea in mind. The cornea develops from surface ectodermal cells that express the paired box transcription factor *PAX6*, and receives inductive signals from the lens vesicle (*27*). The epithelium becomes stratified into 6-8 layers postnatally (mouse) or by birth (human) with the most proliferative capacity residing in the basal layer and these proliferative basal cells are replenished from pools of limbal epithelial stem cells (*9*), a realization that has had vast clinical impact. Multipotent neural crest cells migrate into the space between the epithelium and the lens vesicle to produce the stromal keratocytes and the single cell layered endothelium (*27–30*). The stroma is produced by keratocytes and is composed of glycoproteins, proteoglycans and collagen types I and V primarily, that form uniformly thin collagen fibrils organized into orthogonally stacked lamellae to make up a unique transparent and refractive extracellular matrix (*31–33*). The stromal keratocytes are quiescent under normal corneal homeostasis. Perturbed by infection or injury keratocytes acquire gene expression programs consistent with apoptosis, cell migration, activation and extracellular matrix (ECM) synthesis (*34, 35*). The adult corneal endothelium expresses tight junction marker ZO-1 and the Na+/K+-ATPase pump which maintains corneal hydration for optimal vision (*36*). The corneal endothelium produces an acellular, collagen type VIII-rich basement membrane called the Descemet’s membrane that helps to anchor the endothelium to the cornea (*37*). Interactions between all the three major layers are imperative for proper functioning of the cornea, underscoring the need for a 3D organoid model system. Our data indicates a significant transcriptomic match between the cornea organoid and the human cornea.

## Results

### scRNA-seq of human iPSC derived cornea organoids and human corneal tissue shows shared and unique genes

Central corneas from three human donors (**Fig. S1A**) and three cultured organoids (**Fig. S1B**) were digested separately with collagenase type 1 to prepare single cell suspension with about 80-90 % viability and processed for scRNA-seq using 10X Genomics platform. The scRNA-seq was performed individually on the three organoids and the three human corneas. Each scRNA-seq datasets from the three organoid and three human cornea replicates were pooled and integrated using Cell Ranger Single-Cell software suite v3.01. After doublet cell exclusion, quality control filtering and data integration, a total 25,885 and 29,356 cells were pooled for further analysis from the three organoids and three human cornea, respectively. Henceforth, upon integration the three organoids and the three human corneas are referred to as organoid and human cornea, respectively. The average reads per cell was 40,288 for the organoid which was comparable to that of the human cornea (37,015), indicating acquisition of sufficient reads for accurate cell type classification and biomarker identification. However, the median number of expressed genes per cell was higher in the organoid (3672) compared to that of the cornea (2684) (**Fig S1C**).

Hierarchical clustering of the organoid and the human cornea samples based on gene expression revealed that the biological replicates within each sample type were, as expected, highly similar (**Fig 1A**). Differential gene expression analysis showed 559 genes to be uniquely expressed in the organoid, 389 in the cornea and 971 genes expressed in common (**Fig 1B**). The organoid-expressed genes relate to cell-cell adhesion, cell-extracellular matrix interaction, tight junction, gap junction, focal adhesion and cell cycle (**Fig 1C**). Whereas the genes expressed in the corneas are primarily from signaling pathways that involve TNF-α, NF-κB, chemokines, toll-like and NOD-like receptors suggesting innate immune response activities in the cornea (**Fig 1D**, **Table S1, S2 & S3**).

**Fig. 1.**
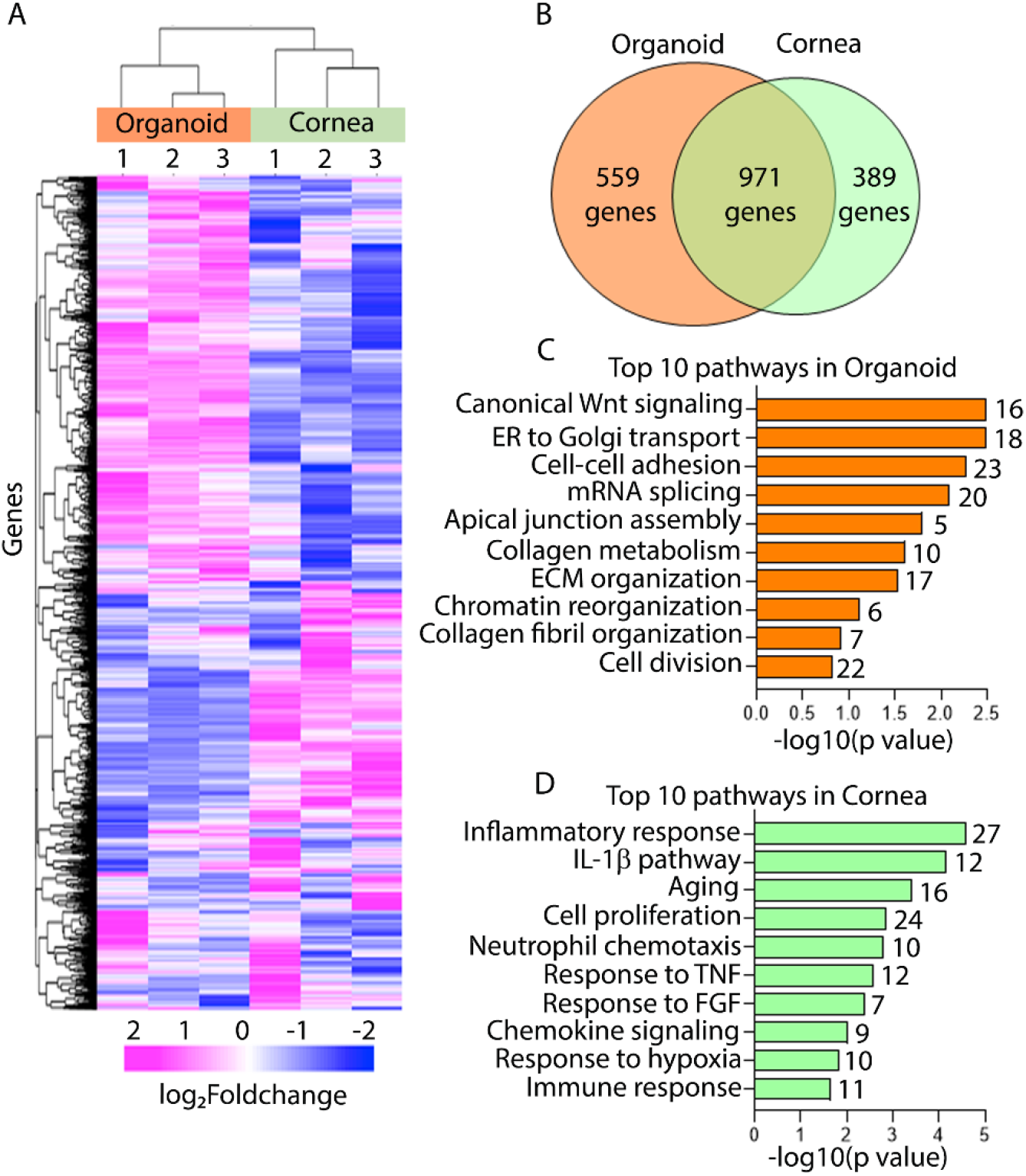
scRNA-seq analysis of the cornea organoid and human cornea. **(A)** Unbiased hierarchical clustering using 2494 differentially expressed genes from the three cornea organoids and the human corneal scRNA-seq data. **(B)** Distribution of unique and commonly expressed genes in the organoid and human cornea samples. **(C-D)** Gene ontology (GO) analysis of the significantly (p < 0.05) expressed unique genes in the organoid (C) and the human cornea (D) showing top 10 significant -log_10_ (p < 0.05) biological processes with the number of genes indicated next to each bar.

### Epithelial, stromal and endothelial cell clusters (CL) in the cornea and organoid

An integrated analysis of the cornea and organoid yielded 30 distinct cell clusters (CL). Of these, 20 CL were detected in the cornea and 14 in the organoid samples (**Fig 2A**). The organoid and the cornea samples share 4 CL (CL17, 22, 27 and 29), while 10 are unique to the organoid (CL 1, 2, 3, 10, 14, 18, 19, 21, 25 and 26), and 16 CL (CL 4, 5, 6, 7, 8, 9, 11, 12, 13, 15, 16, 20, 23, 24, 28 and 30) to the cornea (**Fig. 2B & 2C**). These CL were further analyzed using previously published and broadly validated markers for each of the major cell types **(Fig. S2A)**.

**Fig.2.**
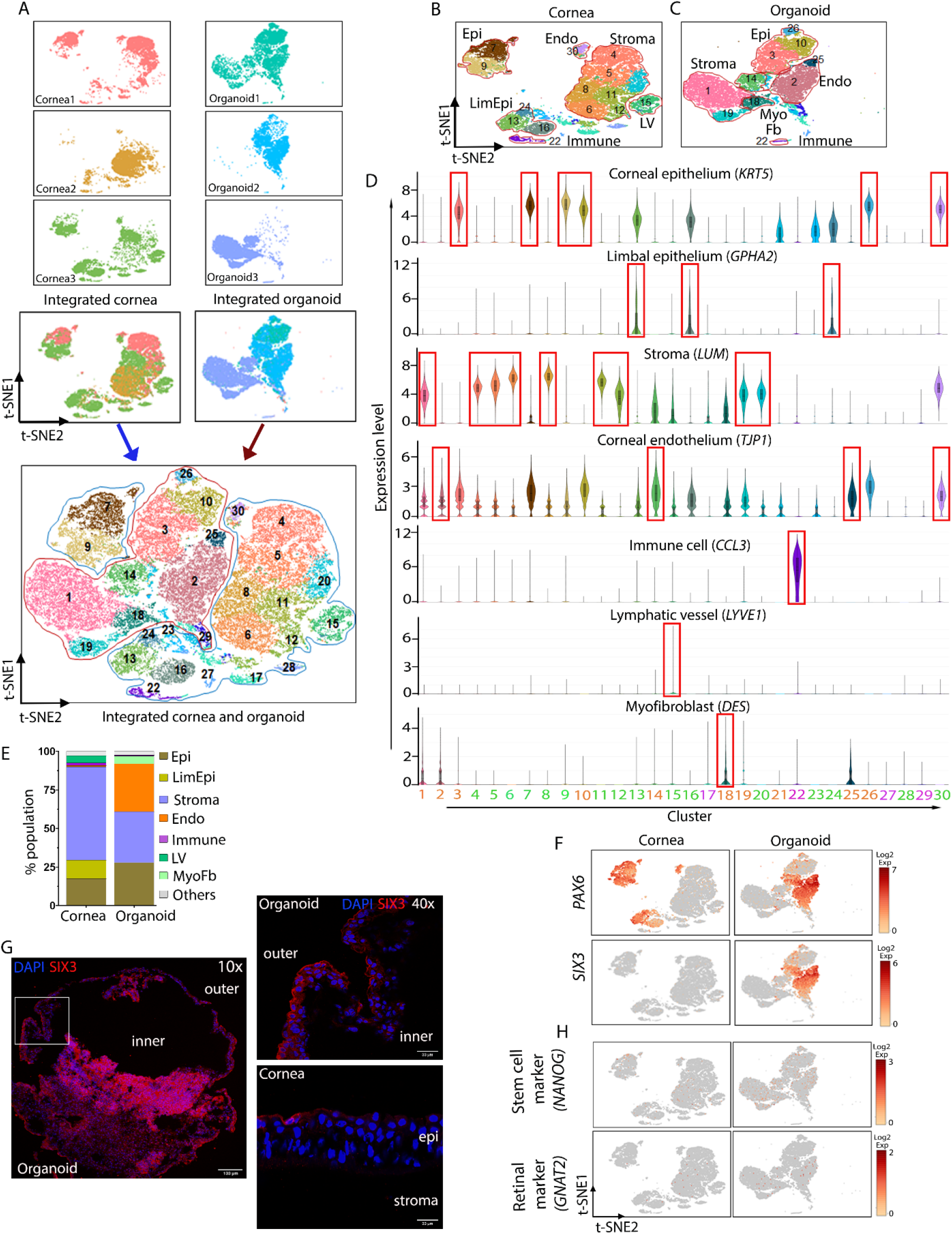
Major cell type clusters in the cornea and the organoid samples. **(A)** Data integration and unbiased clustering of cells identify 30 clusters (CL) in a t-SNE1/2 plot. **(B-C)** t-SNE representation of 20 CL in the human cornea (B) and 14 CL in the organoid indicating seven major cell types (C). Abbreviations used for B &C include, Epi: Corneal epithelial clusters; Stroma: Corneal stromal clusters; LimEpi: Limbal epithelial clusters; Endo: Corneal endothelial clusters; Immune: Immune cell cluster; LV: Lymphatic vessel cell cluster and MyoFb: Myofibroblast. **(D)** Violin plots showing the log_2_expression level of markers for corneal epithelium (*KRT5*), limbal epithelium (*GPHA2*), stroma (*LUM*), endothelium (*TJP1*), immune cells (*CCL3*), lymphatic vessel (*LYVE1*) and myofibroblast (*DES*) in the human cornea and organoid CL. Color code for cluster number in this figure and others: Orange-organoid clusters, Green-human cornea clusters, Pink-organoid and human cornea shared clusters. **(E)** Relative proportions of cell clusters in the organoid and human cornea. **(F)** t-SNE plot showing the expression of the major transcription factors *PAX6* and *SIX3*. **(G)** Immunostaining of SIX3 in the organoid and human cornea sections. Low magnification scale bar = 50 μm. High magnification scale bar=33 μm. **(H)** Lack of expression of the stem cell marker *NANOG* and retinal marker *GNAT2*.

Thus, we initially used keratin 5 (*KRT5*) to delineate epithelial cells, lumican (*LUM*) for stromal keratocytes, tight junction protein 1 (*TJP1*) for endothelial cells, glycoprotein hormone alpha 2 (*GPHA2*) for limbal epithelium, CC-motif ligand 3 (*CCL3*) for immune cells, desmin (*DES*) for myofibroblasts and lymphatic vessel endothelial hyaluronan receptor 1 (*LYVE1*) for lymphatic vessel cell (LV) CL (**Fig. 2D**). Subsequently, we sought expression of other known markers for each cell type (Fig.S2A). Unlike the cornea, where the largest group consists of stromal cells, the organoid shows almost equal proportion of epithelial (28%), stromal (33%) and endothelial (31%) cell contribution. Furthermore, we detected a minor population myofibroblast-like cells (5%) in the organoid (**Fig. 2E**). Importantly, the paired box 6 transcription factor (*PAX6*) which is a master regulator of epidermal cell fate in various organs, including the ocular surface ectoderm, lens vesicle, inner and outer optic cup, optic stalk, the developing and adult corneal epithelia and conjunctiva (*38*), is highly expressed in the epithelial/endothelial clusters of the cornea and organoid samples (**Fig. 2F**). SIX3, a homeobox transcription factor that controls early eye development and is itself regulated by PAX6, is also expressed in the same organoid CL, but its expression in the adult human cornea is limited (**Fig. 2F**). Similarly, the SIX3 protein is present widely in the organoid, and limited to a few epithelial cells in the cornea (**Fig. 2G**).

As organoids differentiate, they are expected to lose the expression of undifferentiated stem cell markers like *NANOG*. Indeed, *NANOG* is expressed at very low levels in the organoid, confirming its differentiating status (**Fig. 2H**). Further analysis shows near-absence of the retinal cone photoreceptor marker, G protein subunit alpha transducing 2 (*GNAT2*) and very low levels of *VSX2* in the organoid (**Fig. 2H, S2B & S2C**). Taken together our data indicate that the organoid harbor cellular states that represent corneal tissue layers.

### Epithelial sub-populations in the organoid

We called *KRT5* expressing clusters 3, 10 and 26 (organoid) and clusters 7 and 9 (corneal) as epithelial (**Fig. 3A & 3B**). Heterodimers of basic and acidic cytokeratin polypeptide chains form functional intermediate filaments (IF) that provide structural support to the epithelial cells by spanning through desmosomes that forms cell-cell junctions and help in epithelial cell adhesion (*39*). Keratin filaments also insert between the hemidesmosomes that help the epithelial layer to attach with the basement membrane (*40*). In the mammalian cornea, KRT5/KRT14 IFs are present widely in corneal and limbal basal epithelial cells, while KRT3/KRT12 is predominant in the basal/suprabasal differentiating corneal epithelial cells (*41*). We detected *KRT3, KRT12, KRT14* and *KRT24* in the corneal CL 7 and 9, and *KRT14* in the organoid CL 3 (**Fig. 3C & 3D**). *KRT19* is broadly present in many simple epithelia, and we detected its presence in the corneal, limbal and conjunctival epithelial CL (not shown) (*42*). Both CL 7 and 9 from the cornea conform to differentiating epithelial layers expressing *KRT12*, but CL 9 expressed many more epithelial keratins, *KRT5/14, KRT3, KRT13, KRT15* and *KRT24* (Fig. 3D). The *KRT5+* CL from the organoid include CL 3, 10 and 26. Of these, CL 3 expressed the widest array of epithelial cytokeratins, *KRT5/14, KRT13, KRT15, KRT18*, but little of *KRT3* or *KRT12* transcripts (**Fig. S3A**). Immunohistology showed the presence of Keratin 5 in the basal epithelium of the cornea and cells in the outer rim of the organoid (**Fig. 2E**). *CENPF*, a primary cilia marker that we identified in the cornea before (*43*) is present in the epithelial and stromal cells of the cornea, and in patches of cells in the organoid (**Fig. 2E**). *TP63*, a transcription factor that regulates epithelial proliferation and stratification and considered an epithelial stem cell marker, is expressed by the corneal CL 7 and 9 and the organoid CL 26 (**Fig. 2B**). Also, the TP63 protein has been shown before to have a cytoplasmic location in basal epithelial cells, while a more nuclear location in epithelial stem cells (*6*). Here, we detected cytoplasmic immunostaining of TP63 in the basal epithelial layer of the cornea, but more nuclear staining of cells with some cytoplasmic staining and colocalization with Keratin3 in the organoid (**Fig. 2E**). Amphiregulin (*AREG*), an epithelial growth factor known to be associated with the mature corneal epithelia is also expressed in the corneal epithelial CL 7 and 9, but undetectable in the organoid epithelial CL (**Fig. S3B**).

**Fig.3.**
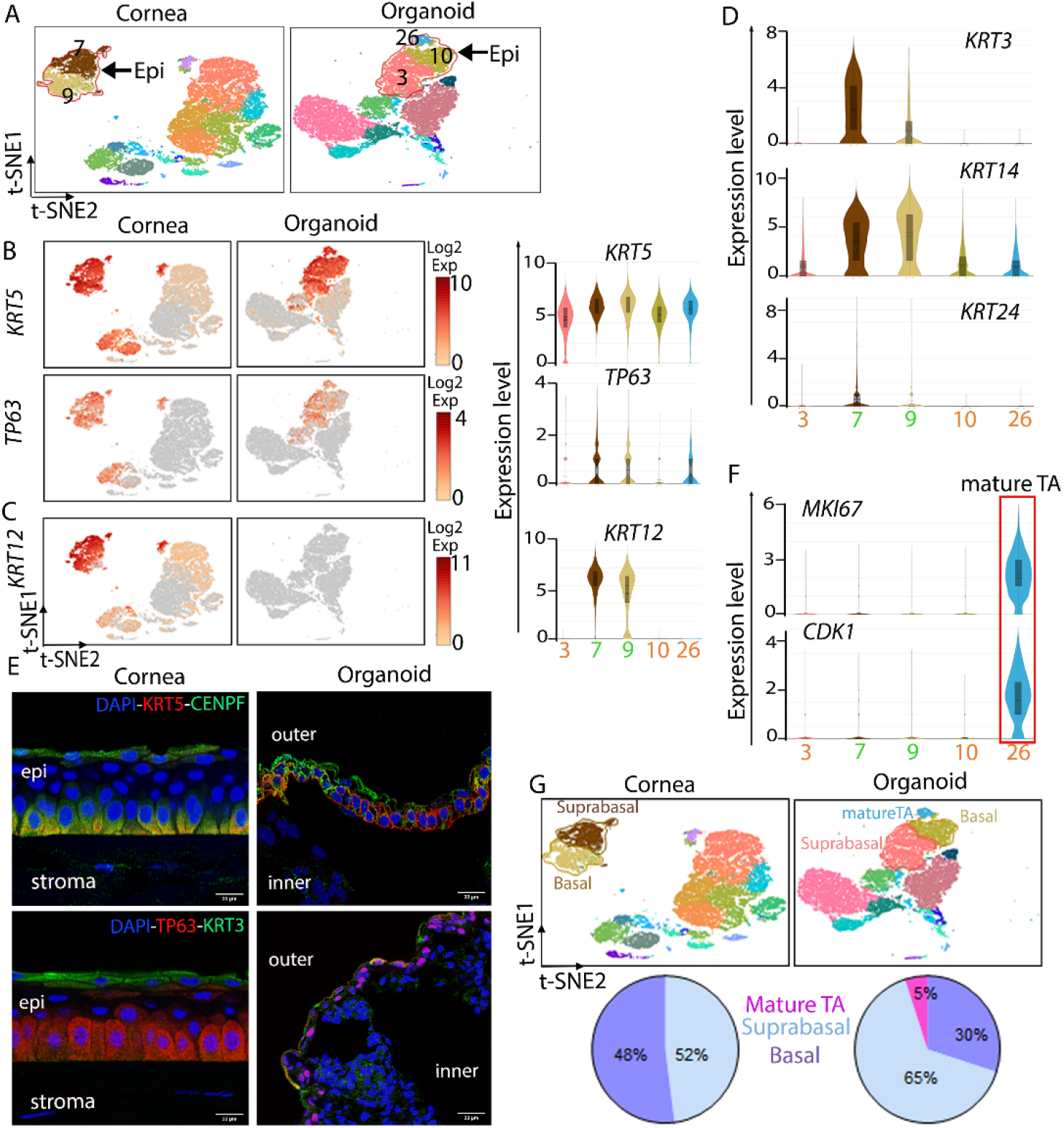
Corneal epithelial-like clusters in the organoid. **(A)** t-SNE representation showing 2 major epithelial CL 7 and 9 in the cornea and CL 3, 10 and 26 in the organoid. **(B-C)** Expression of corneal epithelial markers *KRT5, TP63* (B) and *KRT12* (C) in the human cornea and organoid. Violin plots showing log_2_expression levels CL 7 and 9 (human cornea) and CL 3, 10 and 26 (organoid). **(D)** Expression levels of corneal suprabasal and basal epithelial cell markers (*KRT3, KRT14, KRT24*) in CL 3, 7, 9, 10 and 26. **(E)** Immunostaining of corneal epithelial proteins TP63, KRT3, KRT5 and CENPF. Scale bars =33 μm. **(F)** Gene expression of the mature transiently amplifying (mTA) corneal epithelial cell markers (*MKI67* and *CDK1*) in CL 26. **(G)** t-SNE representation showing the basal, suprabasal and mTA subclusters within the cornea and organoid epithelial CL. Relative proportions of basal, suprabasal and mTA cells in the corneas and the organoids.

Taken together these indicate a lack of fully differentiated epithelial cells in the organoid. CL 26, from the organoid, also expressed marker of proliferation Ki-67 (*MKI67*), cyclin dependent kinase 1 (*CDK1*) and the abnormal spindle microtubule assembly (*ASPM*), that promote cell cycle progression and division (**Fig 3F, S3C**). These findings suggest the presence of a small cluster of mature transiently amplifying (TA) basal epithelial cell states that are in advanced stages of cell cycle (S, G2 and M) in the organoid. However, the organoid epithelial CL 3, 10 and 26 show little to no expression of Leucine rich repeats and immunoglobulin like domains 1 (*LRIG1*), considered to be a marker for early TA cells (*44*) (**Fig. S3D**). Overall, about 30% of epithelial cells (2074 cells) in the organoid are similar to basal epithelial layer cells, 65% to suprabasal (4596 cells) and 5% to mature TA population (371 cells). While in the human cornea 52% of the cells (2444 cells) are basal corneal epithelial and 48% (2682 cells) are suprabasal epithelial cells. Not surprisingly, we could not detect a defined mature TA population in the central cornea (**Fig. 3G**).

Interestingly, a subset of cells within organoid CL 3 and cornea CL 9 also express mucins (*MUC1, MUC4* and *MUC20*) (*45*), known to be expressed by conjunctival epithelial cells (**Fig. S3E**). About 378 cells in the human cornea and 207 cells in the organoid express conjunctival epithelial cell markers. There is no significant expression of *MUC5AC*, a goblet cell mucin in the organoid or the cornea (**Fig. S3F**) (*46*). Moreover, the epithelial clusters also express several other human ocular surface mucins such as *MUC16, MUC2, MUC4, MUC15, MUC21* and *MUC22* **(Fig. S3G)**. *GPHA2* has been identified as a marker for limbal progenitor cells (*10, 47*). High expression of *GPHA2*, along with *KRT15* in the corneal CL 13, 16 and 24 (**Fig. S3H**), both associated with the limbal epithelium defines these clusters in the human cornea as limbal epithelium. The organoids do not express *GPHA2*, indicating an absence of limbal epithelial progenitor cell fate (**Fig. S3I**).

Of note, we detected the expression of SARS-CoV-2 receptor *ACE2* and the plasma membrane bound protease *TMPRSS2*, in the epithelial clusters in both the organoid and human cornea (**Fig. S3J**). Previously, ACE2 and TMPRSS2 were detected in the human corneal epithelial cells (*48, 49*).

Altogether, our data demonstrate that both the organoid and the human cornea express markers for basal corneal epithelium, suprabasal and superficial corneal epithelium. The organoids also show mature TA corneal epithelial-like cells population, but for the most part lack limbal epithelial-like cells.

### Subpopulations of stromal cells

Generally, the corneal stroma is considered to be a single tissue type and its major cellular content defined as keratocytes. The scRNA-seq however, identified seven closely related stromal subpopulations or cellular states (CL 4, 5, 6, 8, 11,12 and 20) in the cornea, and two, CL 1 and 19 in the organoid (**Fig. 4A**). We first focused on the distribution of stromal cell types expressing the major cornea-typical collagens and the proteoglycans. The adult healthy corneal stroma contains fibrillar collagen types I and V (**Fig. 4B**), and lesser amounts of others, such as collagen types VI, XII, XIII, XIV, and XXIV (**Fig. S4A**). The presence of collagen type III in the adult healthy cornea is debatable, but it is present in the developing and in the post-injury remodeling cornea (*50, 51*). The stroma-like CL 1 and 19 cells from the organoids express genes encoding collagen types I, V, as well as III, indicating their similarity with a developing or remodeling stroma (**Fig. 4B**). Immunohistology of the organoid also shows extensive co-localization of collagen types III and V (**Fig. 4C**). Major collagen fibril-associated proteoglycans of the cornea include lumican (*LUM*), decorin (*DCN*), low levels of biglycan (*BGN*) and keratocan (*KERA*), where *KERA* expression is more restricted to the cornea than the either *LUM* or *DCN* (*52, 53*). We found expression of *LUM*, *KERA* and *DCN* in all of the cornea and the organoid stromal CL (**Fig. 4D**). To our surprise, *DCN* is transcriptionally very active in all of the corneal stromal CL, with the expression of *LUM* and *KERA* 10-fold lower than that of *DCN* (**Fig. S4B**). As the genes for these corneal proteoglycans are closely linked, *LUM* and *KERA* may be under some *DCN* mediated cis-regulation (*54*). Fibromodulin (*FMOD*), another collagen-associated proteoglycan, known to localize very specifically to the peripheral and limbal corneal region (*55*), is expressed at very low levels in the stromal CL of the cornea, and is also detectable in the organoid CL 1 and 19 (**Fig. S4C**). Fibrillin1 (*FBN1*), known to make up microfibrils that associate with elastin and provide structural support to the cornea (*56*), is expressed in multiple CL from the cornea and the organoid (**Fig. S4D**). Of all the corneal stromal clusters, CL 6 and 8 show a classical stroma-typical gene expression profile, expressing fibrillar collagen, *COL1A2* and *COL5A2* and the proteoglycan genes, *LUM*, *KERA* and *DCN*, although *COL1A1* or *COL5A1* transcripts are not detectable (**Fig. 4E & S4E**). Immunohistology shows typical staining of decorin along collagen fibrils in the corneal stroma, while the organoid shows staining between cells (**Fig. 4F**). The corneal stromal cell clusters CL 4, 5, 12 and 20, which make up 48% of all cells, express numerous inflammatory genes (**Fig. 4G**). These include pentraxin 3 (*PTX3*), interleukins (*IL6*, *IL32*), C-X-C motif chemokine ligands (*CXCL1, CXCL2, CXCL3* and *CXCL8*), C-C motif chemokine ligand 2 (*CCL2*), serum amyloids (*SSA1, SSA2*) and matrix metalloproteinases (*MMP1, MMP3*) consistent with an activated, somewhat macrophage/monocyte-like gene signature. Contrastingly, keratocytes in CL 6, 8 and 11 have low expression of these genes and are potentially in a resting state (**Fig. 4H)**.

**Fig.4.**
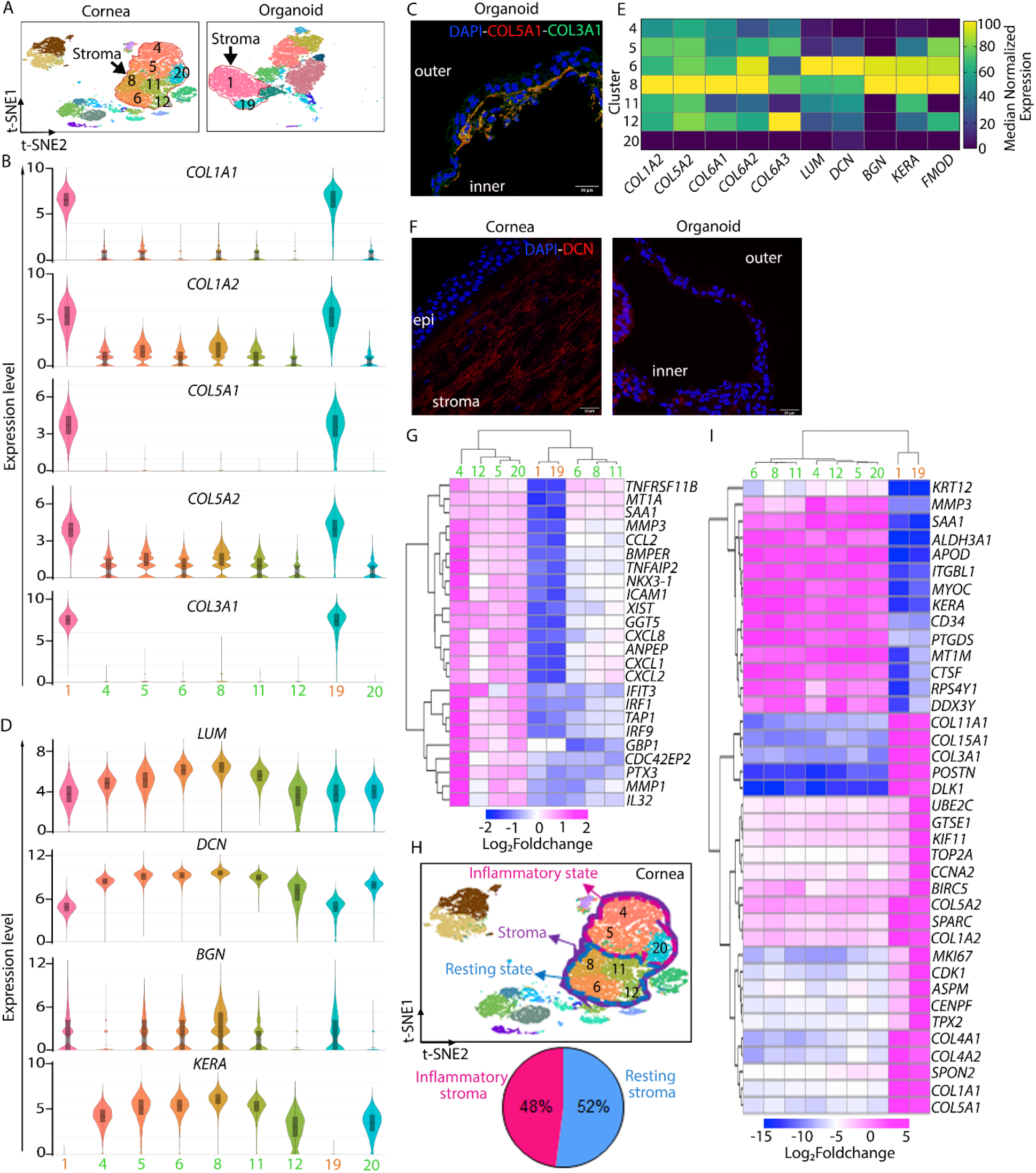
Stromal cell-like clusters in the organoids. **(A)** Stromal cell CL in the cornea include 4, 5, 6, 8, 11, 12, 20, and in the organoids CL 1 and 19. **(B)** Log_2_expression levels of *COLIA1, COL1A2*, *COL5A1, COL5A2* and *COL3A1* in the corneal and the organoid stromal CL. **(C)** Immunostaining of COL5A1 and COL3A1 in organoid sections. Scale bars =33 μm. **(D)** Violin plots showing the log_2_expression level of major stromal proteoglycans *LUM, DCN, BGN* and *KERA* in the cornea and organoid stromal CL. **(E)** Heatmap summarizing collagen and proteoglycan gene expression in the corneal and organoid stromal CL. **(F)** Immunolocalization of decorin in the cornea along collagen fibrils in the stroma. The organoid shows some diffused staining of decorin around cells. **(G)** Heatmap showing the expression of top 24 DEG between the organoid and corneal stromal CL. **(H)** t-SNE plot summarizing CL related to inflammatory (CL 4, 5 and 20) and resting states (CL 8, 6, 11 and 12) in the human corneas. **(I)** Expression of top 38 DEGs in all of the stromal CLs from cornea and organoid samples. Organoids show over expression of fibrillar collagen genes *COL1A1, COL3A1* and *COL5A1*, as well as basement membrane and hemidesmosomal collagens (*COL15A1, COL4A1* and *COL4A2*) associated with an epithelial phenotype.

The organoid CL 19 shows high expression of DNA topoisomerase II alpha (*TOP2A*), Centromere protein F (*CENPF*), *ASPM, MKI67*, Cyclin A2 (*CCNA2*), *EID1, EMP2, PTEN, FSTL1, GPC3, IGF2, CDKN1C* microtubule nucleation factor (*TPX2*), and an apoptosis inhibitor gene *BIRC5*, all consistent with a profile of remodeling and proliferative stromal cells (**Fig. 4I & S4F**). Several other wound healing related stromal matrix genes (*POSTN* and *SPARC*) are elevated in the organoid CL as well (**Fig. S4G**). The transketolase (*TKT*) and aldehyde dehydrogenases (*ALDH*), often described as corneal crystallins (*57*), *TKT, ALDH1A1* and *ALDH3A2* are widely expressed in the corneal and the organoid CL, but *ALDH3A1* is more specifically expressed in the cornea (**Fig. S4H**). From mouse studies, these corneal crystallins are known to increase with eyelid opening and with the acquisition of corneal transparency (*58*). Other transcription factors *KLF4, IRF1* and *MEF2A*, known to be expressed in the adult cornea, are expressed at low levels in the organoid (**Fig. S4I**).

### Clusters with corneal endothelial cell characteristics in the organoid

The corneal endothelium is composed of hexagonally arranged polarized cells that form tight junctions and are involved in fluid and nutrient transport between the cornea and the anterior chamber (*59*). Our data indicates that genes encoding membrane-associated transporters (*ATPB1, SLC20A2*), water channel proteins (*AQP1*) (*60*), cell-cell and cell-ECM adhesion (*NCAM1, CDH2*), tight junction associated proteins (*TJP1, CLDN3*), typically seen in the corneal endothelium, are primarily co-expressed in 3 clusters in the organoids, CL 2, 14 and 25. The human cornea, by contrast shows one, relatively small cluster, CL 30 that co-expresses these endothelial-typical genes (**Fig. 5A-B & S5A**). However, multiple CL that we consider as having stromal properties, individually also express these markers, but lack expression of cell proliferation restriction genes.

**Fig.5.**
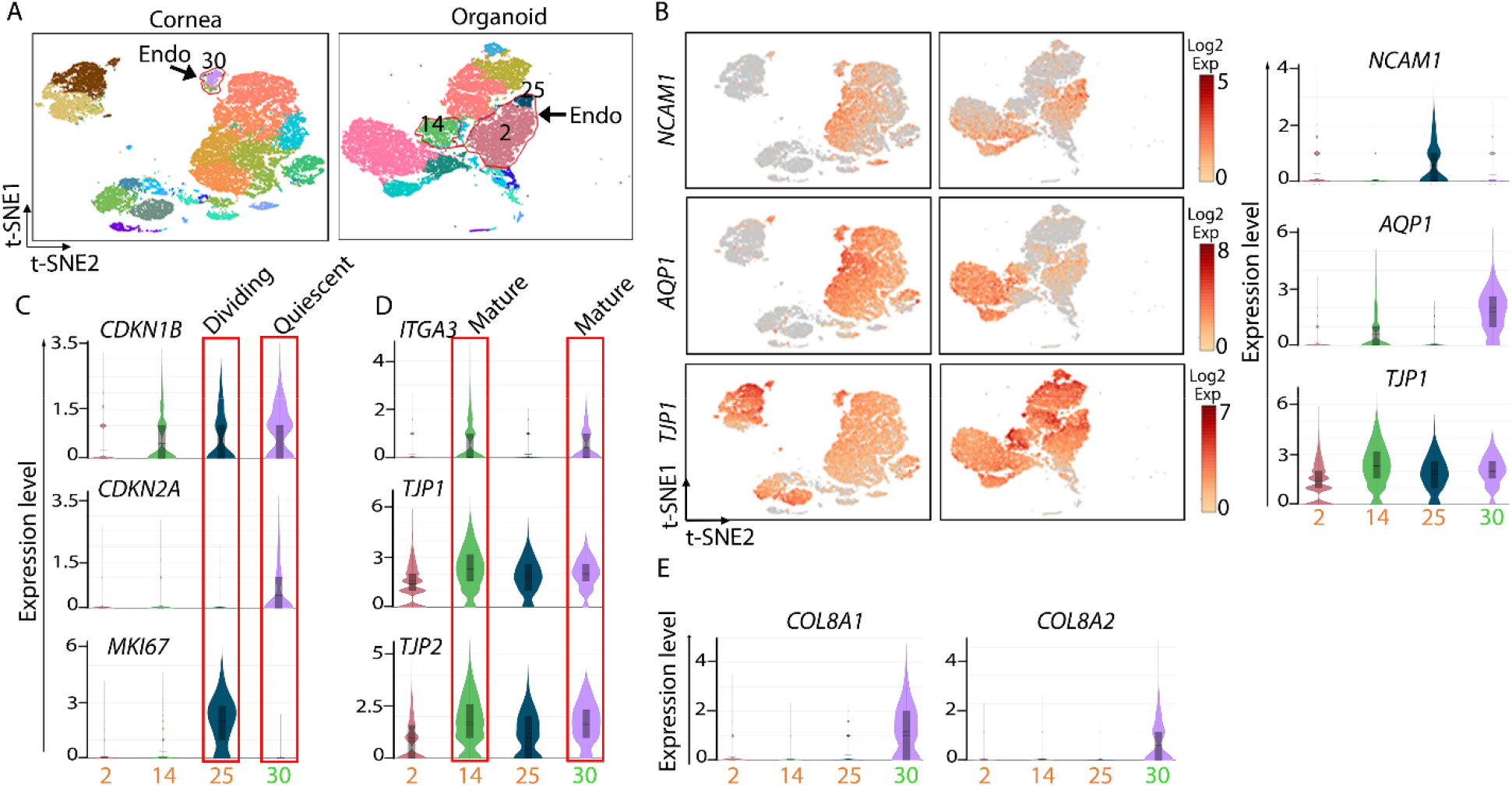
Corneal endothelial cell lineages in organoids. **(A)** t-SNE representation of endothelial cell clusters, CL 30 in the human cornea and CL 2, 14 and 25 in the organoid. **(B)** Expression of major corneal endothelial markers *NCAM1, AQP1* and *TJP1*. Violin plots show the log_2_expression of the genes in CL 30 (human cornea) and CL 2, 14 and 25 (organoid). **(C)** Log_2_expression of cell cycle (*CDKN1B* and *CDKN2A*) and proliferation marker *MKI67* in the endothelial CL 2, 14, 25 and 30. CL 25 coincides with a dividing cell-state and CL 30 with a quiescent cell-state in the organoid and human cornea, respectively. **(D)** Log_2_expression level of the mature corneal endothelial markers (*ITGA3, TJP2* and *TJP1*) in CL 14 and 30. **(E)** Descemet’s membrane marker *COL8A1* and *COL8A2* in CL 2 (organoid) and 30 (human cornea).

In humans, the corneal endothelium has a finite number of mitotically quiescent, G1-arrested cells (*61, 62*). Quiescence is maintained by cyclin-dependent kinase inhibitors of the Cip/Kip family, p27kip1 (*CDKN1B*) and p21 (*CDKN1A*), and INK family p16INK4a (*CDKN2A*) which bind cyclin-CDK complexes to prevent their kinase activity in the mature endothelium (*63*). Consistently, our data shows that the corneal CL 30 has high expressions of *CDKN1B* and *CDKN2A* compared to clusters 2, 14 and 25 from the organoid (Fig. 5C). Moreover, expression of *MKI67, TOP2A, CENPK* and *CCNA2*, associated with cell proliferation, are low in CL 30 (**Fig. 5C & S5B**). Interestingly, cluster 25 expresses high level of Leucine-rich repeat-containing G protein-coupled receptor 5 (*LGR5*), a target of Wnt signaling that is critical for maintaining corneal endothelial cell phenotypes (*64*), CL 2 also expresses *LGR5* but to a lesser extent (**Fig. S5C**). Of the organoid endothelial clusters CL 14 is more consistent with a mature, non-dividing cell state. Unlike CL 2 and 25, we found that CL 14, like corneal CL 30, expresses high levels of *TJP1, TJP2* and *ITGA3*, encoding the alpha 3 subunit of α3β1 integrin (**Fig. 5D**). The TJP proteins mark the basolateral junction while α3β1 integrin mediates adhesion to the underlying Descemet’s membrane in mature endothelial cells. Similarly, *TGFB1*, known to arrest endothelial cells at the G1 phase (*65*), is expressed at high levels in CL 14 and 30 (**Fig. S5D**). A small subset of CL 14 cells also expresses *LUM*, *COL3A1*, *COL1A1* and *COL1A2*, typically seen in stromal keratocytes (**Fig. S5E**). We also detected corneal epithelial markers *KRT3* and *KRT12* in CL 30 indicating inclusion of some cells with epithelial fate (**Fig. S5F**).

Collagen type VIII is normally enriched in the endothelial cell derived Descemet’s membrane where it creates a porous structure that allows nutrients to pass into the stroma and anchors the endothelium to the cornea (*66, 67*). We detected robust expression of both *COL8A1* and *COL8A2* in CL 30 from the cornea (**Fig. 5E**). *COL8A1* is detectable in CL 2 and 14 from the organoid, but *COL8A2* expression is low, suggesting very little production of the functional collagen type VIII heterotrimer by the organoid CL (**Fig. 5E**). Other basement membrane genes, perlecan/heparan sulfate proteoglycan2 (*HSPG2*), fibronectin (*FN1*), nidogen-1 and 2 (*NID1* and *NID2*) and laminins (*LAMA3, LAMA5* and *LAMB1*) are detectable in the organoid endothelial CL (**Fig. S5G**). Overall, the organoid CL 2 captures some of the expression pattern seen in the adult corneal endothelium, while CL 25 may represent a group of dividing endothelial-like cells.

### Organoid harbors immune cell and myofibroblast-like clusters but lack lymphatic vessel-related cells

The organoid and human cornea share CL 22 (159 cells from the organoid and 394 from the cornea sample) which shows high expression of CC-motif chemokine ligands (*CCL3* and *CCL4*) that prompted us to define it as an immune cell cluster (Fig. 6A). Additional inflammatory and immune response related genes expressed in CL 22 are shown in Table S4. We observed that a majority of the cornea-derived cells of CL 22 cells express MHCII-related genes and are likely to be antigen presenting cells (APC) monocytes, macrophages and dendritic cells. By contrast, the CL 22 cells originating from the organoid express high levels of S100 genes (*S100A8* and *S100A9*), cathepsin K (*CTSK*), phospholipase A2 group VII (*PLA2G27*) and acyl-CoA synthetase long chain family member 4 (*ACSL4*) which are typically secreted by granulocytes (**Fig. 6B & S6A**).

**Fig.6.**
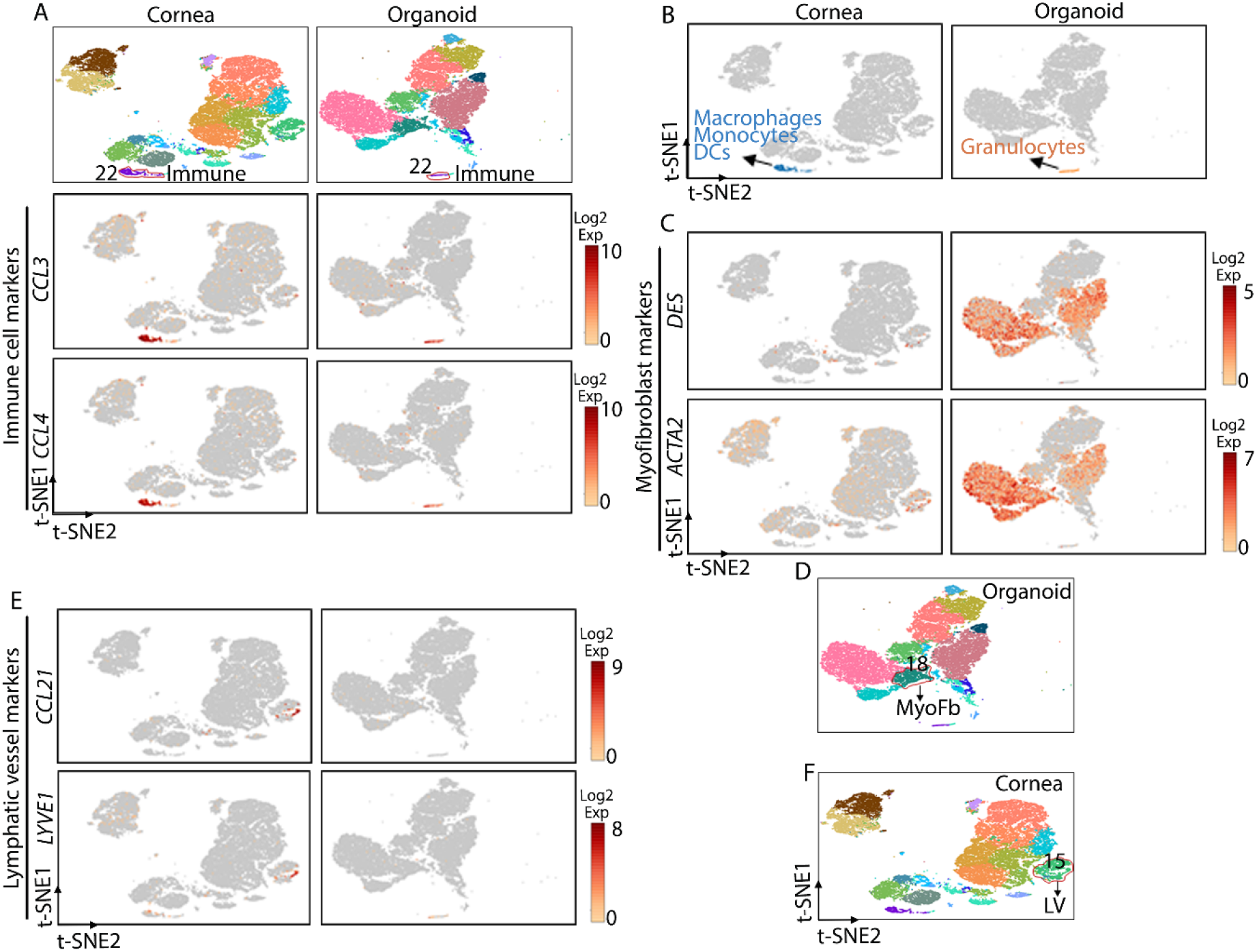
Organoids display immune and myofibroblast-like cell clusters. **(A)** Expression of immune cell markers *CCL3* and *CCL4* in CL 22 shared by the human cornea and organoid samples. **(B)** Based on DEG (Fig. S5A) CL 22 is enriched in macrophages, monocytes and DCs in the cornea and granulocytes in the organoid. **(C-D)** Cells expressing myofibroblast markers *DES* and *ACTA2* in the organoid CL 18 which also expresses high levels of *VIM* (Fig.S5B). **(E-F)** Expression of lymphatic vessel markers *CCL21* and *LYVE1* presence of the lymphatic vessel CL 15 in the human cornea only.

We delineate CL 18 from the organoid as mature myofibroblast-like as it shows high expression of desmin (*DES*), actin (*ACTA2*) and vimentin (*VIM*) (**Fig. 6C-D & S6B**). Further analysis of CL18 shows high expression of genes encoding titin (*TTN*), myosin heavy chain and light chain (*MYH6, MYL3* and *MYL4*), myosin light chain kinase (*MYLK*) and tropomyosin-binding subunit 2 (*TNNT2*) (**Fig. S6C**). These cytoskeletal genes regulate cell adhesion, cell migration, cargo transport, cilia formation and cell division in repair-related myofibroblasts (*68*). In the cornea myofibroblasts are both beneficial and harmful; they regulate corneal healing after trauma, infections and surgeries, but can also cause vision-debilitating haze (Wilson, 2020). *CCL21* and *LYVE1*, two lymphatic vessel markers, are predominantly expressed in CL 15 leading us to define this cluster as lymphatic vessel-derived cells, present in the cornea, but absent in the organoid (**Fig. 6E-F**).

## Discussion

In this study we performed concurrent scRNA-seq analyses of three human iPSC-derived cornea organoids and full-thickness central corneas from three different adult human donors. Our human corneal dataset corroborates recent transcriptomic studies of the full thickness human cornea. Human cornea organoids have not been analyzed at the transcriptomic level before. Here we show the repertoire of cornea-like cells present in the organoid. As a whole, the organoid display features of a developing or a healing cornea. In two earlier studies, Ligocki et al., reported 16 distinct clusters from the cornea, and Collin et al., found 21 clusters from the cornea and adjacent conjunctival tissue. In our study we removed most of the peripheral limbal region from the cornea, and identified two major epithelial and 3 limbal clusters, while Ligocki reported 8 epithelial and 3 conjunctival clusters. We detected expression of hallmark genes of the epithelium (*PAX6, KRT3*, and *KRT12*), stroma (*DCN, KERA*, and *LUM*) and the endothelium (*AQP1, TJP1, COL8A1*, and *COL8A2*) in the adult donor human corneas. While we identified seven closely related subpopulations within the corneal stromal cluster, the Ligocki study identified one stromal cluster. All major ECM genes are represented in both studies. The stromal subpopulations in our study distinguished between quiescent and proliferating cells, and those with a distinct immune cell-related profile. In fact, somewhat unexpectedly, ~48% of the corneal stromal CL express genes related to inflammation and immune response. In an earlier array-based transcriptomic study of the mouse cornea we detected expression of many immune response related genes typically seen in macrophages and considered this to have some species-specific underpinnings (*69*). Our current study however shows that the human cornea also contains a substantial amount of inflammation and immune response related cells in the stromal cell clusters.

In the scRNA-seq of the organoid we detected all major corneal cell types. Additionally, markers of the undifferentiation iPS cells (*NANOG, OCT4*) are no longer detectable in the organoid, indicating its progression into differentiated cell fates. The gene expression pattern of the organoid resembles the immature developing cornea more closely than the adult stage. *PAX6*, that helps to define the eye field and early embryonic development (*70*), is highly expressed in the epithelial and endothelial clusters of the organoid. *SIX3*, a member of the homeobox transcription factor, known to suppress WNT signaling and regulate *PAX6* to promote eye development, shows high *PAX6*-overlapping expression in the organoid, and expectedly little to no expression in our adult corneal clusters. The human cornea continues to grow and mature for at least 6 months after birth, at which time it is influenced by various developmental signaling pathways as well as external cues. The major events in the developing cornea include stratification of the squamous epithelial cells, increase in stromal matrix deposition and a decline in endothelial cell density (*27–29, 71*). The gene expression pattern of the organoid sample type conforms to this general developmental scheme. Expression of *KRT5* and *KRT14* in the organoid indicates the presence of less differentiated basal and suprabasal-like cells. We also detected mature TA population only in the organoid indicative of their high proliferative potential (*44*). *KRT12*, a hallmark of differentiating stratified corneal epithelium (*72*), was not detectable in the organoid. High expression of several cell growth factors (*GPC3, IGF2, FSTL1, EGF* and *EID1*) and cell proliferation proteins (*MKI67, CENPF* and *ITM2A*) also support the active cell growth and proliferative state of the organoid analogous to immature developing corneas.

Type IV collagen, which is a hallmark of basement membranes is abundant in the cornea, but transcripts for *COL4A1* and *COL4A2* were present at very low levels in our corneal epithelial CLs, because its turnover in the adult healthy cornea is low. The organoid CLs on the other hand showed robust expression of *COL4A1* and *COL4A2* as reported for infant human corneas (*73*). The transcription factor *KLF4*, widely expressed in the cornea and known to regulate epithelial homeostasis (*74*), was expressed at low levels in the organoid epithelial and endothelial clusters. The ECM collagen types I and V are the most abundant proteins of the corneal stroma, yet the adult corneas show very low levels of the respective transcripts. This is to be expected as these proteins are highly stable with little turn over in a homeostatic cornea. Corroboratively, in our earlier transcriptomic study of the postnatal day 10 and the adult mouse cornea the transcripts for collagen types I and V were markedly elevated in the postnatal cornea compared to the adult (*75*). In the same vein, here we noted the organoid stromal CL to have high levels of transcripts for *COL1A1/COL1A2* and *COL5A1/COL5A2*, as well as *COL3A1* that encodes for type III collagen which is typical of the immature and injured cornea. Elevated *POSTN* in the organoid is also consistent with its role in enhanced collagen fibrillogenesis and stromal maturation (*76*). Moreover, expression of proliferation-related genes separates the two stromal CL of the organoid; CL 19 shows a more proliferative profile compared to CL 1.

Our findings show that overall human iPSC-derived cornea organoids harbor cells and expression patterns that resemble the cornea. Although the organoids were matured in culture for 4 months, they resemble immature developing corneas rather than the adult cornea. The organoids characterized here were biological replicates from the same batch and have similar gene expression patterns. Additional standardizations are thus required to determine whether organoids generated using the same protocol but at different times would be as consistent. Genetically manipulated iPSC-derived cornea organoids can be used to explore the role of specific genes in the differentiation of specific cell types. This type of genetic lineage analysis, using human iPSCs was previously demonstrated in mouse kidney nephrogenesis (*77*). While SARS-CoV-2 was not the focus of the current study, co-expression of ACE2 and TMPRSS2 in the organoid supports its suitability to model SARS-CoV-2 infection and study the accompanying cellular and molecular changes. The cornea organoids may be used in functional studies of genes potentially involved in cell-cell and cell-ECM adhesion and tissue-level organization during early corneal development. The cellular complexity and organizational similarity of the organoid to the human cornea also makes it suitable for screening the effects of drugs and their toxicities on the different cell types that mimic corneal cells.

## Materials and Methods

### Induced pluripotent stem cell (iPSC) maintenance

The IMR90.4 iPSCs were obtained from WiCell (Madison, Wisconsin) and grown at UC San Diego with IRB approval. Stem cells were maintained antibiotic free on 1% (vol/vol) Matrigel-GFR (#354230; BD Biosciences) coated dishes at 37°C under hypoxia conditions (10% CO_2_/5%O_2_) in mTeSR1 (StemCell Technologies). These were passaged every 4-6 days with Accutase (#A6964; Sigma) for 8-10 minutes, dissociated to single, quenched in mTeSR1 plus 5μM (−) blebbistatin (B; #B0560; Sigma), pelleted at 80 g for 5 minutes, and resuspended in mTeSR1+B and plated at 5,000 cells per 35mm dish. After 48 hours, cells were fed without B.

### Cell culture medium

As described in Wahlin et al, 2017, E6 supplement for making BE6.2 medium for cell maintenance consists of DMEM/F12 (1:1) (#11330-032; Invitrogen) with 19.4 mg/L insulin (#11376497001; Roche), 64 mg/L L-ascorbic acid (#A8960; Sigma), 14 μg/L sodium selenium (#S5261; Sigma), 10.7 mg/L transferrin (#T0665; Sigma), 19.4 mg/L NaHCO3. Osmolarity was raised +30 mOsm to ~330-340 mOsm by adding 0.88 g/L NaCl. BE6.2-NIM (B27 + E6 at 2X concentration) (neural induction medium) for cell differentiation consists of DMEM (#11965; Invitrogen) with 1% B27 vitamin A (−) (#12587010; Invitrogen), 10% heat inactivated FBS (#16140071; Invitrogen), 1 mM pyruvate (#11360; Invitrogen), 1x NEAA (#11140; Invitrogen), 1x Glutamax (#35050061; Invitrogen) and 1mM taurine (#T-8691; Sigma).

### Human corneal organoid differentiation

Human induced pluripotent stem cells (hiPSCs) and hiPSC-corneal organoids were cultured as previously described (*18, 19, 23*). Briefly, stem cells were passaged with Accutase for 12 min and 1000-3000 cells in 50 μl of mTeSR1+B were seeded per well into polystyrene 96-well U-bottom plate (#650180; Greiner). Over the first 4 days, aggregate were transitioned to neural induction medium (BE6.2-NIM) by adding 50 μl of BE6.2 + 2% MG on day 1 and 50 μl of BE6.2 + 2% MG each day thereafter. On D4-8 a 50% medium exchange (100 μl) was performed daily and every other day thereafter. NIM also contained 3μM of the WNT antagonist (IWR-1-endo; #681669; EMD Millipore) from D1-6. Organoids were grown in BE6.2 + 300nM Smoothened agonist (SAG; #566660; EMD Millipore) from D10-D12 and then LTR+SAG from D12-D18. To separate vesicles, we used sharpened tungsten needles to excise optic vesicles from D10-12 as previously described (*18, 19*). Unlike retinas that require low density to thrive, corneas develop at higher concentration. Organoids were maintained in suspension in LTR and fed every 2-3 days. Around day 31, the corneal organoids have a translucent cystic-like appearance and allowed to mature in culture for ~ 4 months.

### Single-cell isolation of cornea organoid and human cornea

Three cadaverous donor human corneas (aged 51, 41 and 33 years old) were received from Lions Eye Institute for Transplantation and Research, Tampa, Florida (**Fig. S1A**). The central region of the human corneas were dissected away from the peripheral limbal area (**Fig S1B**). The samples were cut into small pieces and digested in 2 ml of digestion mix: 2mg/ml collagenase type 1 (#SCR103; EMD Millipore) in DMEM-F12 supplemented with 5% FBS (#10438026; Gibco), 1% antibiotic-antimycotic (#15240062; Gibco), 1mM L-ascorbic acid using rotatory nutator at 37°C for 6h at 5% CO_2._ Every 2h the supernatant was collected by allowing the undigested tissue to settle under gravity and this was repeated 2-3 times. The collected supernatant samples were stored on ice at all stages and pooled after a final 6h of digestion, and centrifuged at 500g for 5 min. The supernatant was discarded and the pellet resuspended in 5 ml of 1x Accutase and incubated at 37°C for 20 min in a 5% CO_2_ incubator. Fresh 5 ml of DMEM-F12 with 5% FBS was added and passed through a 40 μm cell strainer and centrifuged at 500g for 5 min/4°C. Supernatant was discarded and the pellet resuspend in PBS with 0.04% BSA or complete DMEM-F12 to make a single cell suspension. Live cells were counted by trypan-blue exclusion in a Countess® II automated cell counter.

### Single-cell library preparation and sequencing

For scRNA-seq libraries, we used single cell suspension from 3 different cornea organoids and the central cornea dissected from 3 different human donor corneas, separately. A total 6 individual (3 organoids and 3 human corneas) libraries were prepared using the Chromium Single Cell 3’ Library & Gel Bead Kit v2 (10x Genomics). The six libraries were run on an Illumina HiSeq 4000 as 150-bp paired-end reads.

### Analysis of scRNA-seq and pre-processing of scRNA-seq data

#### Quality control and integration

Sequencing results were demultiplexed and converted to FASTQ format using the Illumina bcl2fastq software. The Cell Ranger Single-Cell Software Suite v3.01 (http://support.10xgenomics.com/single-cell-gene-expression/software/pipeline/latest/what-is-cell-ranger) was used to perform sample demultiplexing, barcode processing and single-cell 3’ gene counting. The cDNA insert was aligned to the hg19/GRCh38-3.0.0 reference genome. Meaningful reads were calculated as the sum of four metrics: (1) valid barcodes; (2) valid unique molecular identifier (UMI); (3) associated with a cell barcode; and (4) confidently mapped to exons (*78*). Only confidently mapped, non-PCR duplicates with valid barcode and unique molecular identifiers were used to generate the gene-cell-barcode matrix that contained 7809 to 11728 cells. Further analysis including quality filtering, identification of highly variable genes, dimensionality reduction, standard unsupervised clustering algorithms and discovery of the differentially expressed genes (DEG) was performed using Cell Ranger Single-Cell Software Suite v3.01 and Loupe Browser 5.1.0. To exclude low quality cells, we removed all cells with less than 1000 detected genes. To exclude cells that were extreme outliers and that may include multiple cells or doublets, we removed all cells in the top 2% quantile. We also removed cells with more than 5% of transcripts derived from mitochondrial genes. After quality filtering, the mean and median numbers of detected genes per cell were 26,434 to 46,569 and 2,592 to 3,878, respectively. Integration of the three human cornea and three organoid samples was performed using the Cell Ranger Single-Cell Software Suite v3.01. Gene-cell-barcode matrix from each of the six samples was linked and merged. The integrated dataset has 55,241 cells in the gene-cell-barcode matrix and the mean and median number of genes per cell were 2,932 and 9,634, respectively.

#### Dimensionality reduction, clustering, differential expression and data visualization: Dimensionality reduction

To reduce the gene expression matrix, Cell ranger uses Principal Component Analysis (PCA) to change the dimensionality of the dataset. For 2-D visualization, Cell Ranger passes the PCA-reduced dataset into t-SNE (t-Stochastic Neighbor Embedding), a nonlinear dimensionality reduction method.

#### Clustering

The datasets were then clustered using a graph-based clustering algorithm where cells are clustered based on expression similarity, followed by Louvian Modularity Optimization (LMO), an algorithm that finds highly connected “modules” in the graph. Additional hierarchical cluster-merging was done if there were no differentially expressed genes between two sibling clusters. This hierarchical clustering and merging was repeated until there were no more cluster-pairs to merge. To remove batch effects Cell Ranger uses an algorithm based on mutual nearest neighbors (MNN) to identify similar cell subpopulations between batches. The matched cell subpopulation between batches were then used to merge multiple batches together.

#### Differential expression

Differentially expressed genes (log2-foldchange > 0.5, FDR < 0.05) for a specific cluster were identified as the difference between the average expression by cells in the cluster and the average expression by cells not in the cluster. In order to identify specific genes in each cluster, Cell Ranger checks for each gene and each cluster *a*, whether the mean expression in cluster *a*, differs from the mean expression across all other cells (global) or within cells in a selected cluster (local). The mean expression of a gene in cluster *a*, is calculated as the total number of UMI count from that gene in the cluster *a*, divided by the sum of size factors for cells in cluster *a*. The log_2_ fold-change of a gene expression in cluster *a* relative to other clusters is the log_2_ ratio of mean expression within cluster *a*, and outside of cluster *a*.

#### Data visualization

Loupe Browser 5.1.0 was used to analyze and visualize the data. Cell Ranger cloupe files were used to generate gene expression information for cells in the samples. Various gene expression-based clustering information for the cells, t-SNE projections, differential gene expression, location of different cell types, subtypes and functional groups were done using Loupe. Violin plots were generated using graph-based log_2_ expression data and heatmaps were generated using graph-based log_2_ fold-changes.

### Immunofluorescence staining

The organoids were fixed with 4% paraformaldehyde (PFA) for 4h at 4°C on a rocker. Tissues were rinsed with 1X PBS, followed by overnight incubation in 10%, 20% and 30% sucrose at 4°C on a rocker. After removing the extra sucrose, the tissues were embedded into OCT containing molds on dry ice and stored in −80°C. The unfixed human donor corneas were directly placed into the OCT containing mold on liquid nitrogen and stored in −80C freezer. The frozen sections of human donor cornea were air dried for 1h and fixed in 4% PFA for 15 min. The 10 μm organoid and human cornea sections were permeabilized with 0.1% Triton X-100 in PBS for 5 min at room temperature (RT). Followed by 2 wash with PBS and blocked using 3% BSA, 5% normal goat serum in PBS (blocking buffer) for 1h. Primary antibodies were diluted in 1X PBS with 0.05% Tween-20 (PBST) at appropriate concentrations and slides were incubated for overnight at 4C in a humidified chamber. Slides were then washed 3 x 5 min with PBST and incubated with secondary antibodies diluted in blocking buffer for 1h at RT. Followed by washing 3 x 5 min with PBST and counter-stained with 1μg/ml DAPI (#564907; BD Biosciences) for 2 min. Slides were washed twice with ice-cold 1X PBS to remove extra DAPI and mounted in Vectashield antifade mounting media (#H-1000; Vector Laboratories) with coverslips. The slides were visualized using Zeiss LSM-700 laser scanning confocal microscope and images were analyzed using Fiji. Primary antibodies and dilution: Anti-mouse CENPF (1:100; #MA1-23185) from Invitrogen; anti-rabbit COL5A1 (1:500; #PA5-83158) and anti-rabbit SIX3 (1:200; #PA5-103901) from Thermo Fisher; anti-mouse COL3A1 (1:300; #ab6310), anti-rabbit ZO1 (1:100; #ab96587), anti-rabbit DCN (1:1000; #ab277636), anti-rabbit KRT5 (1:50; #ab64081) and anti-mouse KRT3 (1:100; #ab77869) from abcam; anti-rabbit TP63 (1:100; #CS-13109T) from Cell Signaling. Secondary antibodies and dilution: Goat Anti-Mouse IgG-AF488 (1:500; #ab150117) and Goat Anti-Rabbit IgG-AF555 (1:500; #ab150086) from abcam.

### Statistical analysis

Statistical analysis of two groups was performed with Student’s two-tailed *t* test. P < 0.05 was considered as statistically significant and the significance is marked by * p<0.05, ** p<0.01, *** p<0.001.

## Acknowledgements

We thank the NYU Langone Health core facilities: Genome Technology Center for single cell RNA sequencing, Experimental Pathology Research Laboratory for tissue sectioning and Microscopy Laboratory for confocal imaging.

## Author contributions

Conceptualization: SC, GM. Investigation: GM, SC. Supervision: SC. Data Analysis: GM, SC. Methodology: GM, KJW, MR, MRMB, NH. Writing-original draft: GM, SC. Writing-review & editing: SC, GM, MRMB, KJW, NH.

## Data and material availability

Accession numbers for the scRNA-seq from Gene Expression Omnibus will be provided later upon deposition.

## Supplementary Materials PDF file includes

Figs. S1 to S6

Legends for tables S1, S2, S3 and S4

