## Supplementary Figures for "Single cell RNA-seq identifies developing corneal cell fates in the human cornea organoid"

**This PDF file includes:**

Figs. S1 to S6
Legends for Table S1, S2, S3 and S4

A

| Samples | Age (years) | Gender | Death to Preservation time (hrs) |
|---|---|---|---|
| Cornea1 | 41 | F | 10:23 |
| Cornea2 | 33 | M | 9:25 |
| Cornea3 | 51 | M | 9:00 |

B

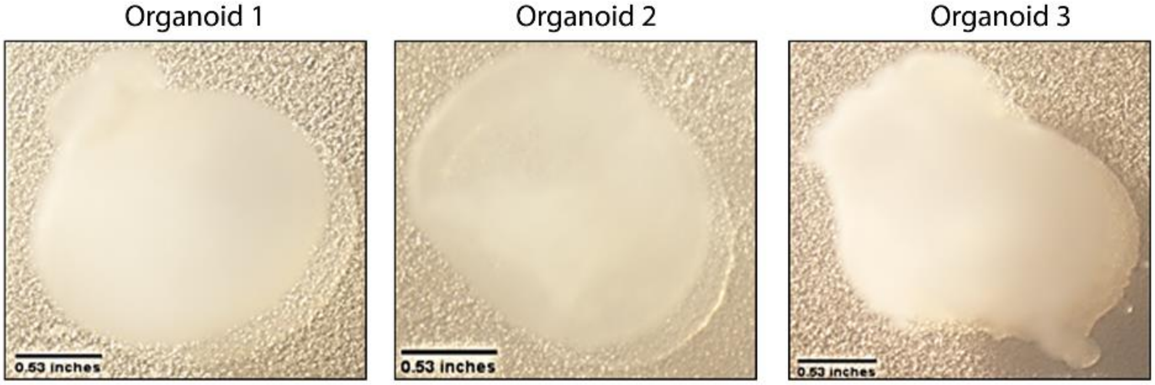


C

| Samples | Cell count (cells/ml) | Viability (%) | No. of cells analyzed | Mean genes /cell | Median no. of genes/cell | Total genes detected |
|---|---|---|---|---|---|---|
| Organoid1 | 400000 | 89 | 9,096 | 39,898 | 3,754 | 29,531 |
| Organoid2 | 320000 | 81 | 7,090 | 46,569 | 3,878 | 29,228 |
| Organoid3 | 480000 | 83 | 9,547 | 34,399 | 3,383 | 29,288 |
| Cornea1 | 1960000 | 88 | 10,455 | 38,064 | 2,769 | 26,102 |
| Cornea2 | 820000 | 86 | 7,809 | 46,549 | 2,592 | 25,623 |
| Cornea3 | 1200000 | 83 | 11,728 | 26,434 | 2,692 | 28,801 |

**Fig.S1. Organoid and human cornea quality. (A)** Demography and death to preservation time for the human donor corneas. **(B)** Light microscopic images of the organoids used in this study. Scale bar= 0.53 inches. **(C)** Cell counts, viability and expressed gene counts in the corneas and organoids.

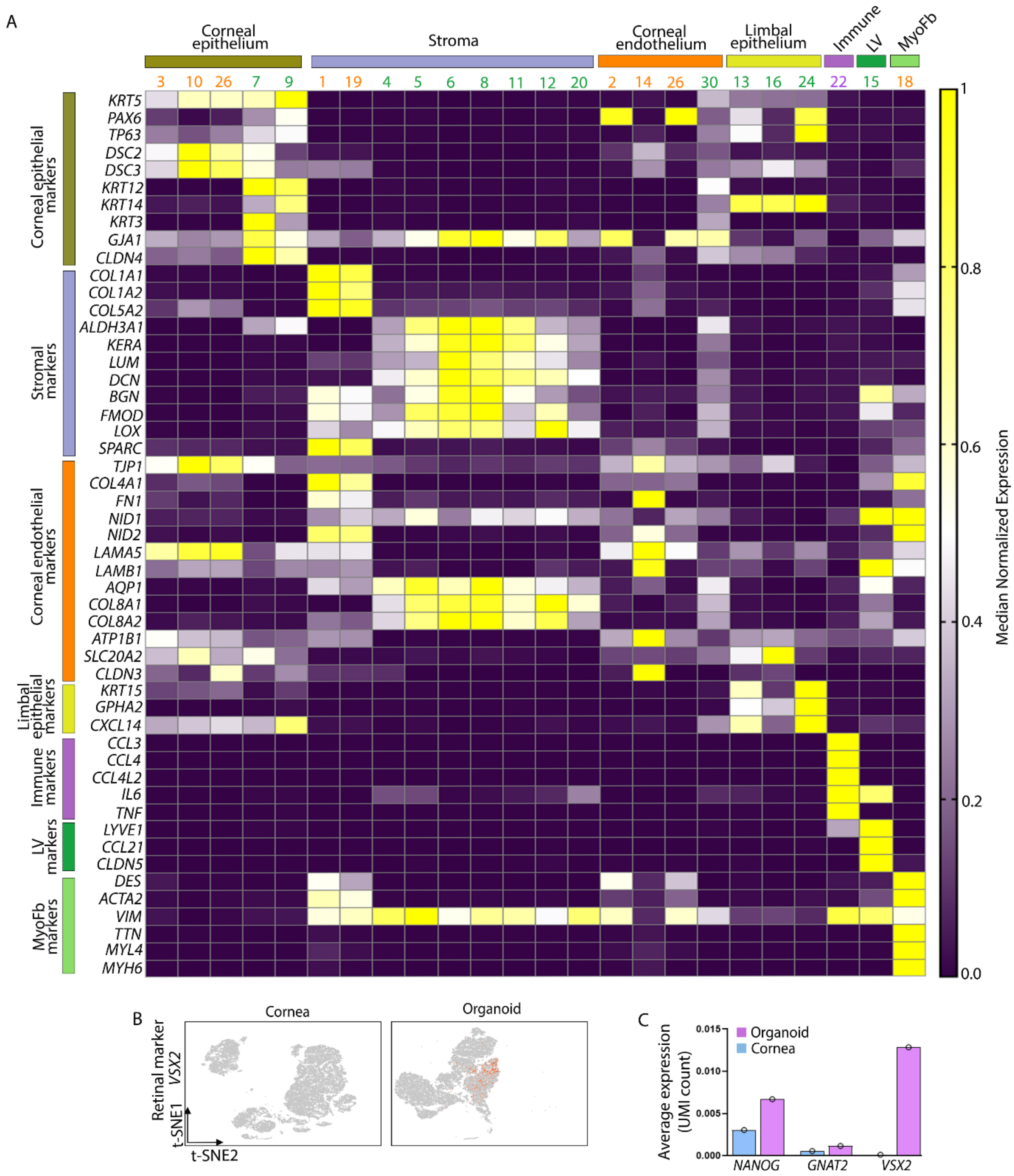


**Fig.S2. Key markers for cluster identification. (A)** Heatmap showing the normalized median expression level of the key marker genes used to identify corneal epithelium, stroma, corneal endothelium, limbal epithelium, immune cells, lymphatic vessels (LV) and myofibroblast (MyoFb) clusters. **(B)** t-SNE plot showing very low expression of retinal marker *VSX2* in the organoid. **(C)** Average expression level (UMI count) of the stem cell marker, *NANOG* and the retinal markers *GNAT2* and *VSX2*. Color code for cluster number in all figures: Orange-organoid clusters, Green-human cornea clusters, Pink-organoid and human cornea shared clusters.

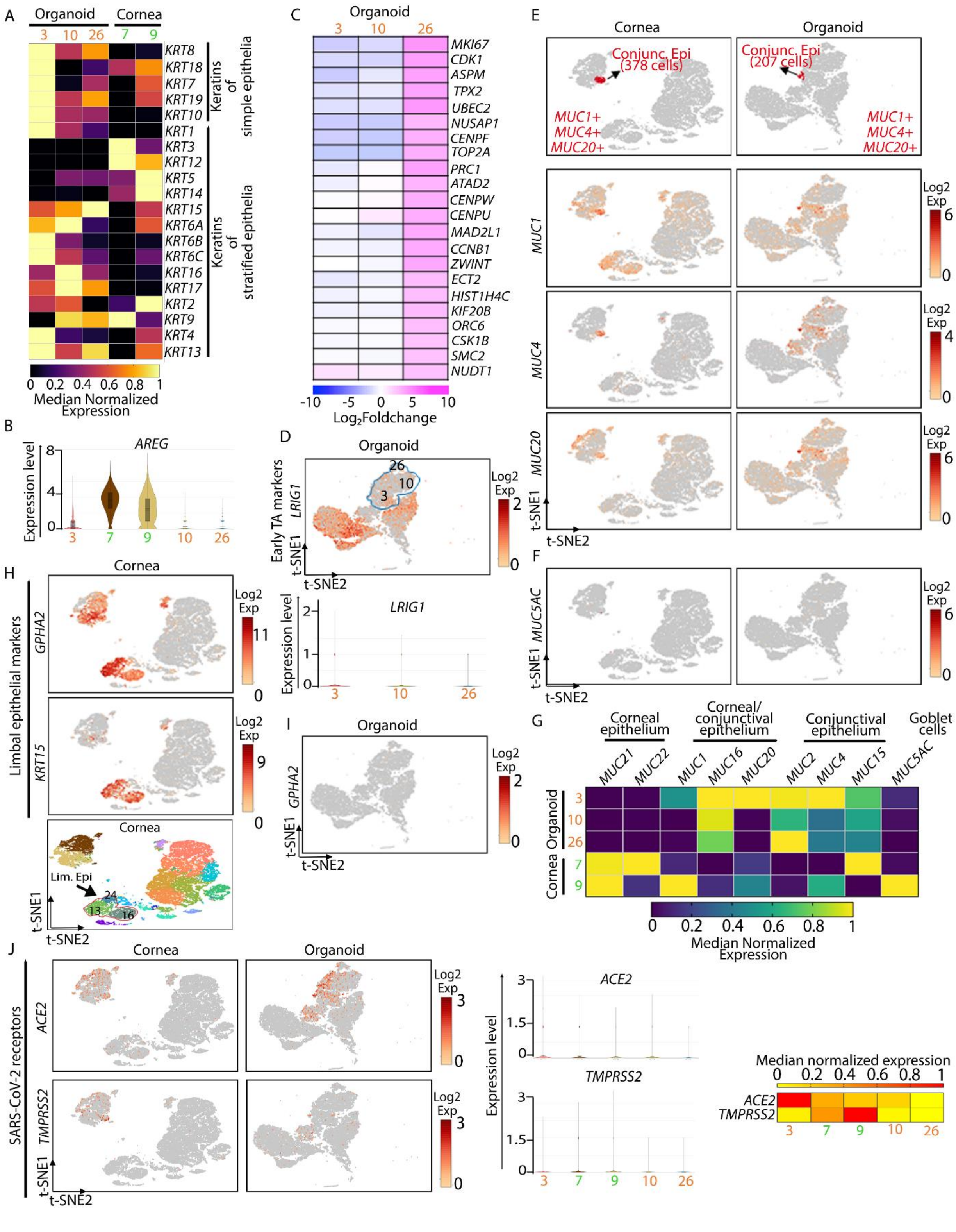


**Fig.S3. Organoids express corneal epithelial markers and SARS-CoV-2 receptor. (A)** Heatmap showing the normalized median expression level of simple and stratified epithelial keratins in the epithelial CL 7 and 9 (cornea) and 3, 10 and 26 (organoid). **(B)** Expression of *AREG* in CL 3, 7, 9, 10 and 26. **(C)** Top 22 DEG within the epithelial CL 3, 10 and 26 in the organoid. **(D)** Expression of stem/early TA marker, *LRIG1* in the organoid. Violin plot showing the expression level of *LRIG1* in the organoid CL 3, 10 and 26. **(E)** t-SNE representation of a small subset of conjunctival epithelial cells (*$MUC1^{+}MUC4^{+}MUC20^{+}$)* within the epithelial cluster in human cornea (378 cells) and organoid (207 cells). Heatmap showing expression of mucins that are expressed in corneal and conjunctival epithelium in the CL 3, 7, 9, 10 and 26. **(F)** Very little to no expression of goblet cell marker *MUC5AC* in the cornea or organoid. **(G)** Heatmap showing expression level of ocular surface mucins across the epithelial clusters. **(H-I)** Expression of limbal epithelial markers *GPHA2* and *KRT15* in CL 13, 16 and 24 in the human cornea (H), and absence of *GPHA2* expression in the organoid (I). **(J)** t-SNE plot showing the expression of SARS-CoV-2 receptor, *ACE2* and the cell membrane associated protease, *TMPRSS2* in the human cornea and organoid and violin plots showing their expression level in the corneal epithelial CL 3, 7, 9, 10 and 26. Heatmap summarizing *ACE2* and *TMPRSS2* expression in the corneal and organoid epithelial CL.

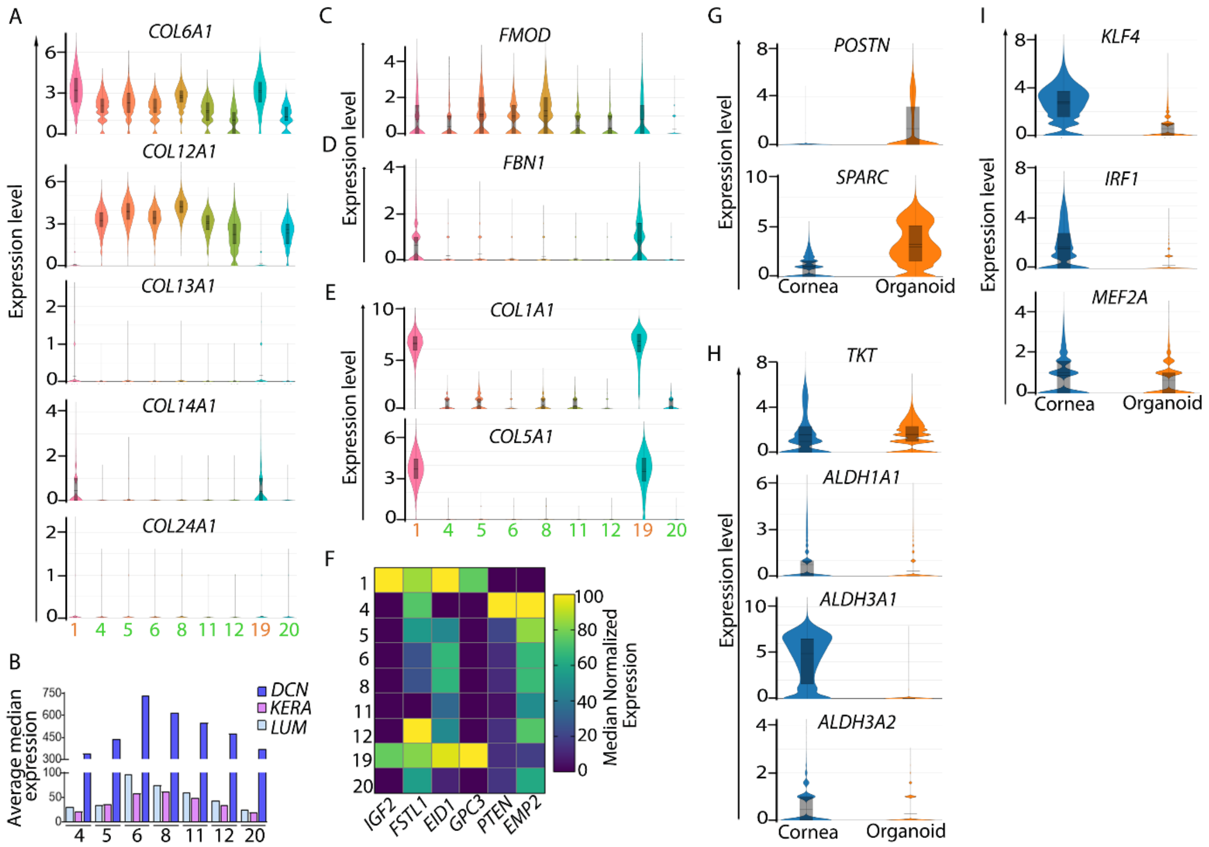


**Fig.S4. Organoids express corneal typical and atypical stromal cell markers. (A)** Violin plots showing expression of collagen genes *COL6A1*, *COL12A1*, *COL13A1*, *COL14A1* and *COL24A1* in the stromal CL 1 and 19 (organoid), and CL 4, 5, 6, 8, 11, 12, and 20 (cornea). **(B)** Average median expression of the major stromal proteoglycan genes *DCN*, *KERA* and *LUM* in the corneal stromal CL. **(C-D)** expression of *FMOD* and *FBN1* in all stromal clusters. **(E)** Expression of *COL1A1* and *COL5A1*. **(F)** High expression of *IGF2*, *FSTL1*, *EID1* and *GPC3* in CL19 of organoids. **(G)** Cumulative expression of stromal matrix genes (*POSTN* and *SPARC*) in the human cornea and organoid. **(H)** Cumulative expressions of corneal crystalline genes *ALDH1A1*, *ALDH3A1*, *ALDH3A2* and *TKT* in the human cornea and organoid. **(I)** Cumulative expressions of transcription factors *KLF4*, *IRF1* and *MEF2A* in the human cornea.

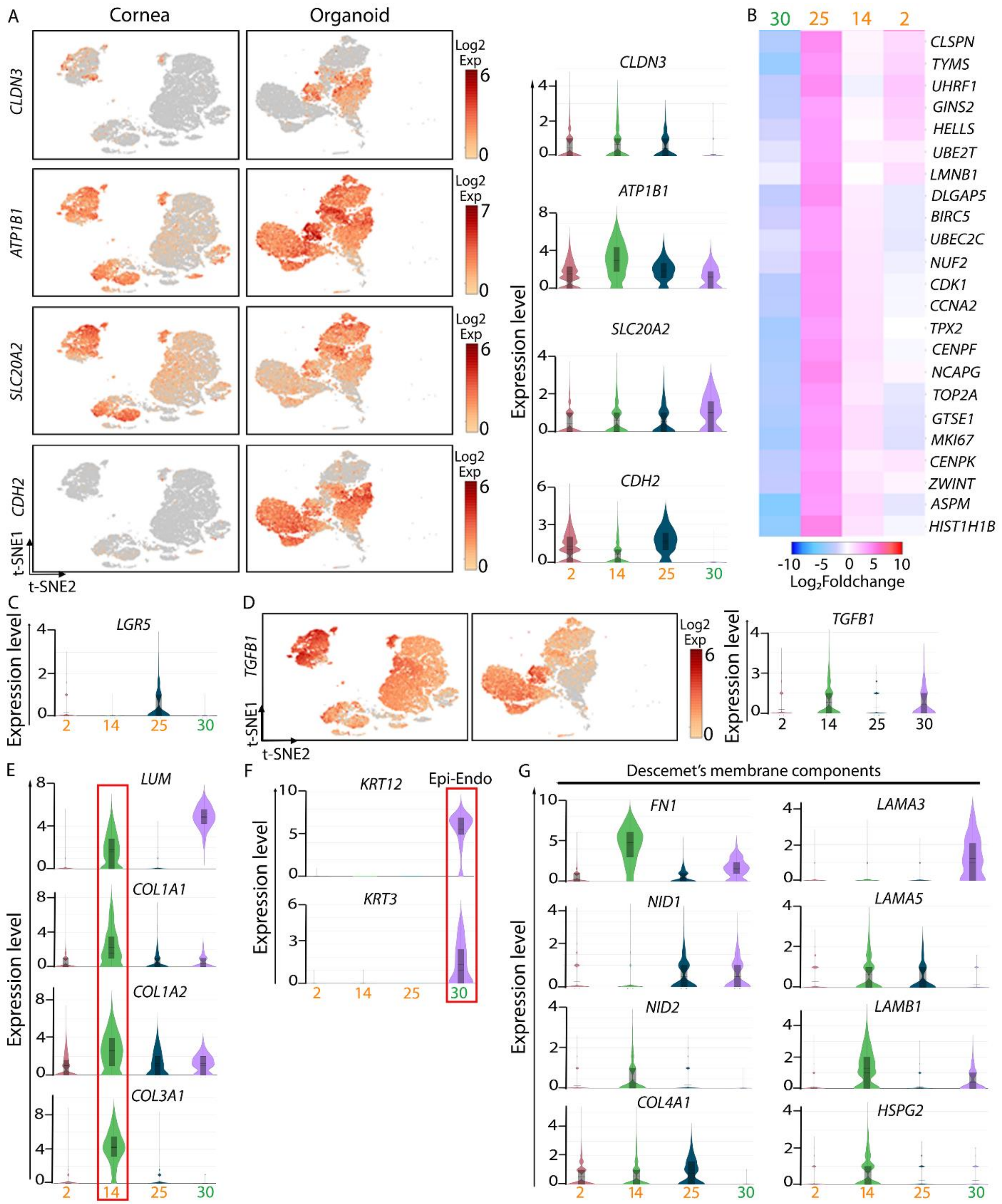


**Fig. S5. Organoids express corneal endothelial markers. (A)** t-SNE plot showing the expression of corneal endothelial markers *CLDN3*, *ATP1B1*, *SLC20A2* and *CDH2* in the human cornea and organoid and violin plots showing their expression level in the endothelial CL 2, 14, and 25 (organoid) and 30 (human cornea). **(B)** Top 23 DEG in the organoid CL 2, 14, 25 and corneal CL 30. **(C)** Expression levels of *LGR5*, a Wnt signaling target in CL25. **(D)** *TGFB1*

expression in CL 2, 14, 25 and 30. **(E)** Expression level of stromal matrix genes *COL1A1*, *COL1A2*, *COL3A1* and *LUM* in CL 2, 14, 25 and 30. **(F)** Expression of corneal epithelial keratin genes *KRT3* and *KRT12* in CL30 of the human cornea. **(G)** Expression of basement membrane genes *FN1*, *NID1*, *NID2*, *COL4A1*, *LAMA3*, *LAMA5*, *LAMB1* and *HSPG2*.

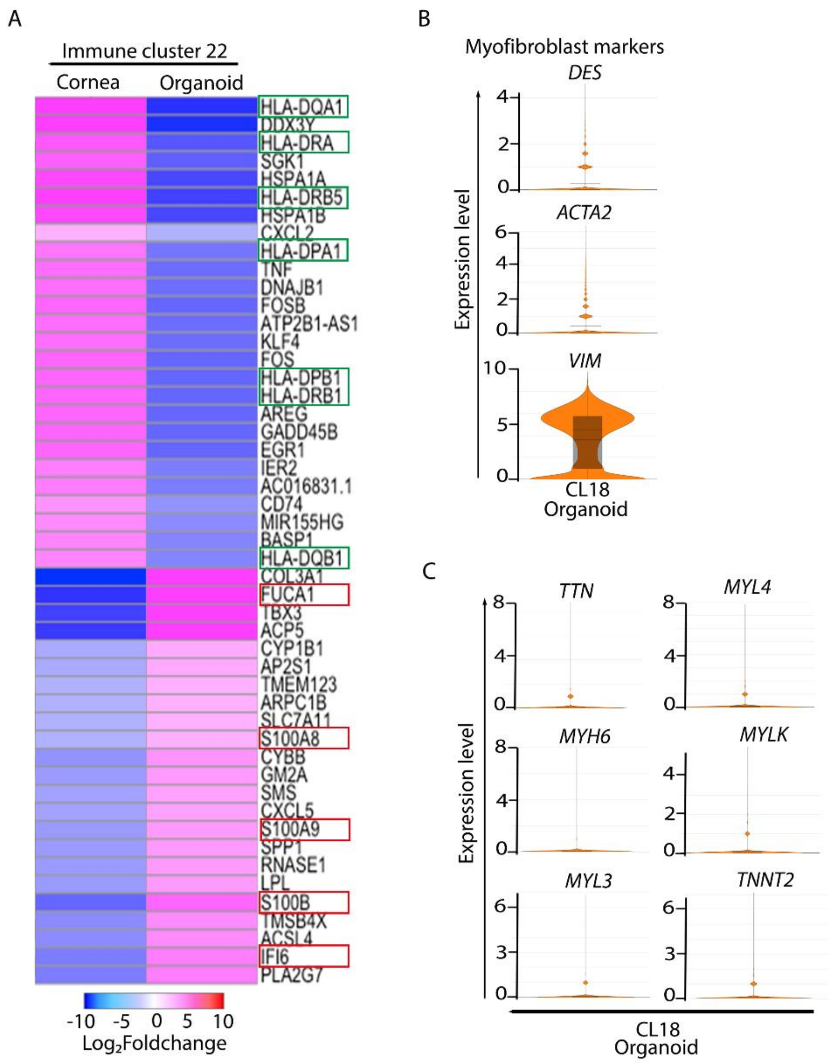


**Fig. S6. Expression of selected immune cell and myofibroblast markers. (A)** Top 50 DEG in the immune CL 22 detected in the human corneas and organoids. Genes (*HLA-DQA1*, *HLA-DRB5*, *HLA-DRA*, *HLA-DRB1* and *HLA-DPB1*) in the green boxes encode Major Histocompatibility Complex (MHC) proteins. Genes (*S100B*, *FUCA1*, *CTSK*, *IFI6* and *S100A9*) in the red boxes encode bactericidal secretory proteins. **(B)** Violin plots showing the expression of myofibroblast markers *DES*, *ACTA2* and *VIM* in CL 18 of the organoid. **(C)** Expression of muscle cell markers *TTN*, *MYH6*, *MYL3*, *MYL4*, *MYLK* and *TNNT2* in the organoid CL18.

**Table S1. List of all the genes that are expressed in the cornea organoid only.** Average expression of 559 genes detected in the integrated cornea organoid only.

**Table S2. List of all the genes that are expressed in the human cornea only.** Average expression of 389 genes detected in the integrated human cornea only.

**Table S3. List of all the genes that are expressed in both the human cornea and the cornea organoid.** Expression of 971 genes common to the integrated human cornea and the integrated organoid data.

**Table S4. List of all the significant genes expressed in the immune cluster.** Average expression level, log2foldchange and p-values for genes that are expressed significantly in cluster 22 in the human cornea and the organoid.
