## Supplementary material for "Single cell RNA-seq identifies developing corneal cell fates in the human cornea organoid": Table S1

| Serial No. | FeatureID | FeatureName | Organoid.Average | Organoid.Log2.Fold.Change | Organoid.P.Value |
|---|---|---|---|---|---|
| 1 | ENSG00000185559 | DLK1 | 17.65683002 | 11.55252888 | 0 |
| 2 | ENSG00000168542 | COL3A1 | 53.59996417 | 9.167753028 | 2.83E-261 |
| 3 | ENSG00000133110 | POSTN | 5.458473257 | 8.976630907 | 3.74E-245 |
| 4 | ENSG00000113196 | HAND1 | 1.400629283 | 9.035541987 | 1.95E-228 |
| 5 | ENSG00000145423 | SFRP2 | 1.750151028 | 8.771977603 | 3.91E-223 |
| 6 | ENSG00000159217 | IGF2BP1 | 1.000650308 | 8.533343525 | 3.71E-214 |
| 7 | ENSG00000060718 | COL11A1 | 1.176577649 | 8.222190346 | 3.36E-208 |
| 8 | ENSG00000130635 | COL5A1 | 4.304716512 | 7.011508091 | 1.43E-183 |
| 9 | ENSG00000167244 | IGF2 | 2.069015595 | 7.051139607 | 1.13E-179 |
| 10 | ENSG00000164107 | HAND2 | 1.069113022 | 7.107470847 | 3.40E-176 |
| 11 | ENSG00000170558 | CDH2 | 1.118104689 | 6.953100195 | 5.35E-173 |
| 12 | ENSG00000204291 | COL15A1 | 2.622489828 | 6.760954435 | 3.31E-170 |
| 13 | ENSG00000147257 | GPC3 | 2.19912167 | 6.551223488 | 5.00E-164 |
| 14 | ENSG00000108821 | COL1A1 | 29.09300445 | 6.307626346 | 1.35E-158 |
| 15 | ENSG00000130303 | BST2 | 1.239626754 | 6.381329333 | 9.24E-156 |
| 16 | ENSG00000242265 | PEG10 | 1.913994954 | 5.77245451 | 4.94E-140 |
| 17 | ENSG00000166426 | CRABP1 | 1.549099756 | 5.378454498 | 1.20E-124 |
| 18 | ENSG00000103485 | QPRT | 1.112272348 | 5.374203138 | 6.57E-124 |
| 19 | ENSG00000134871 | COL4A2 | 1.11804487 | 5.067971167 | 3.54E-115 |
| 20 | ENSG00000069011 | PITX1 | 1.598689611 | 5.062164247 | 1.09E-114 |
| 21 | ENSG00000102109 | PCSK1N | 1.103329425 | 5.02026108 | 9.05E-114 |
| 22 | ENSG00000187498 | COL4A1 | 1.386093294 | 5.017274807 | 7.37E-113 |
| 23 | ENSG00000038427 | VCAN | 2.057620097 | 5.000470452 | 8.81E-113 |
| 24 | ENSG00000130600 | H19 | 12.19569482 | 4.979015487 | 1.85E-112 |
| 25 | ENSG00000163209 | SPRR3 | 1.630064616 | 7.069280245 | 1.90E-108 |
| 26 | ENSG00000169469 | SPRR1B | 1.615079986 | 4.958310859 | 1.04E-97 |
| 27 | ENSG00000140416 | TPM1 | 7.308791008 | 4.457831749 | 4.66E-95 |
| 28 | ENSG00000105048 | TNNT1 | 1.219318243 | 4.445684361 | 1.16E-94 |
| 29 | ENSG00000135916 | ITM2C | 1.239357569 | 4.417699997 | 4.81E-94 |
| 30 | ENSG00000114115 | RBP1 | 2.322169116 | 4.334674241 | 4.37E-91 |
| 31 | ENSG00000110492 | MDK | 5.69063035 | 4.217329072 | 8.46E-88 |
| 32 | ENSG00000119888 | EPCAM | 1.741806294 | 4.205144011 | 3.11E-86 |
| 33 | ENSG00000170421 | KRT8 | 7.290665886 | 4.032817045 | 8.76E-82 |
| 34 | ENSG00000196923 | PDLIM7 | 1.036780915 | 3.923785188 | 5.65E-78 |
| 35 | ENSG00000134138 | MEIS2 | 2.022326955 | 3.892711555 | 4.01E-77 |
| 36 | ENSG00000106484 | MEST | 1.335007966 | 3.872301867 | 2.38E-76 |
| 37 | ENSG00000118971 | CCND2 | 1.434965323 | 3.842579566 | 2.36E-75 |
| 38 | ENSG00000179222 | MAGED1 | 3.023784818 | 3.758609015 | 3.54E-73 |
| 39 | ENSG00000171604 | CXXC5 | 1.662725728 | 3.476884568 | 4.37E-64 |
| 40 | ENSG00000154380 | ENAH | 2.525224321 | 3.409552418 | 2.40E-62 |
| 41 | ENSG00000154277 | UCHL1 | 1.635867048 | 3.407471491 | 5.49E-62 |
| 42 | ENSG00000161013 | MGAT4B | 1.035644356 | 3.343067596 | 4.31E-60 |
| 43 | ENSG00000130203 | APOE | 3.239820721 | 3.221411004 | 3.73E-56 |
| 44 | ENSG00000146386 | ABRACL | 1.986854356 | 3.075252261 | 2.46E-52 |
| 45 | ENSG00000035403 | VCL | 4.433267296 | 3.031042612 | 2.95E-51 |

| | | | | | |
|---|---|---|---|---|---|
| 46 | ENSG00000101335 | MYL9 | 1.083230279 | 3.070935666 | 4.94E-51 |
| 47 | ENSG00000039560 | RAI14 | 1.034627435 | 2.953359552 | 8.83E-49 |
| 48 | ENSG00000136542 | GALNT5 | 1.802133639 | 2.871875152 | 5.22E-46 |
| 49 | ENSG00000182670 | TTC3 | 4.869346968 | 2.82647978 | 1.50E-45 |
| 50 | ENSG00000198695 | MT-ND6 | 1.65082177 | 2.849580658 | 3.23E-45 |
| 51 | ENSG00000182871 | COL18A1 | 1.38630266 | 2.763546098 | 1.43E-43 |
| 52 | ENSG00000102226 | USP11 | 1.206756277 | 2.734659503 | 7.60E-43 |
| 53 | ENSG00000102172 | SMS | 3.335889851 | 2.625657702 | 4.68E-40 |
| 54 | ENSG00000109472 | CPE | 1.597911966 | 2.622024611 | 1.10E-39 |
| 55 | ENSG00000077942 | FBLN1 | 2.799553728 | 2.60449448 | 2.19E-39 |
| 56 | ENSG00000272398 | CD24 | 4.289313149 | 2.582822398 | 9.85E-39 |
| 57 | ENSG00000073792 | IGF2BP2 | 1.823369343 | 2.574314986 | 1.13E-38 |
| 58 | ENSG00000223573 | TINCR | 2.153420041 | 2.584240422 | 1.20E-38 |
| 59 | ENSG00000204262 | COL5A2 | 4.879396541 | 2.561575316 | 2.45E-38 |
| 60 | ENSG00000102038 | SMARCA1 | 2.418118608 | 2.539919746 | 7.77E-38 |
| 61 | ENSG00000122378 | PRXL2A | 1.116519489 | 2.502748027 | 1.03E-36 |
| 62 | ENSG00000180964 | TCEAL8 | 1.508452823 | 2.474155698 | 4.12E-36 |
| 63 | ENSG00000111371 | SLC38A1 | 2.208961876 | 2.457607617 | 1.15E-35 |
| 64 | ENSG00000100345 | MYH9 | 2.91691838 | 2.454770503 | 1.23E-35 |
| 65 | ENSG00000103196 | CRISPLD2 | 1.002026143 | 2.463169468 | 1.51E-35 |
| 66 | ENSG00000185630 | PBX1 | 1.18728523 | 2.445042945 | 2.92E-35 |
| 67 | ENSG00000046604 | DSG2 | 2.032466256 | 2.435819573 | 4.88E-35 |
| 68 | ENSG00000140937 | CDH11 | 2.234384902 | 2.436203532 | 4.94E-35 |
| 69 | ENSG00000185624 | P4HB | 3.510052534 | 2.419823449 | 8.57E-35 |
| 70 | ENSG00000180447 | GAS1 | 1.338447552 | 2.414709355 | 2.34E-34 |
| 71 | ENSG00000184226 | PCDH9 | 1.273902975 | 2.420678001 | 3.24E-34 |
| 72 | ENSG00000187840 | EIF4EBP1 | 1.549727854 | 2.373510573 | 1.52E-33 |
| 73 | ENSG00000257923 | CUX1 | 1.888900932 | 2.35848145 | 3.12E-33 |
| 74 | ENSG00000118495 | PLAGL1 | 1.06707918 | 2.35642093 | 4.79E-33 |
| 75 | ENSG00000111885 | MAN1A1 | 1.456021571 | 2.346023136 | 7.84E-33 |
| 76 | ENSG00000188290 | HES4 | 1.064536878 | 2.357165601 | 8.74E-33 |
| 77 | ENSG00000149212 | SESN3 | 1.217583495 | 2.329287971 | 2.37E-32 |
| 78 | ENSG00000102316 | MAGED2 | 1.622736803 | 2.318421862 | 2.95E-32 |
| 79 | ENSG00000108312 | UBTF | 1.137067276 | 2.321969746 | 2.97E-32 |
| 80 | ENSG00000169851 | PCDH7 | 1.043301173 | 2.317325652 | 5.89E-32 |
| 81 | ENSG00000070961 | ATP2B1 | 3.177010892 | 2.29336771 | 1.23E-31 |
| 82 | ENSG00000198467 | TPM2 | 2.050172646 | 2.231110551 | 5.84E-30 |
| 83 | ENSG00000123131 | PRDX4 | 4.058412253 | 2.218739217 | 6.71E-30 |
| 84 | ENSG00000091986 | CCDC80 | 1.127885077 | 2.22229989 | 1.53E-29 |
| 85 | ENSG00000127616 | SMARCA4 | 1.064088236 | 2.206453044 | 1.88E-29 |
| 86 | ENSG00000172201 | ID4 | 2.824019652 | 2.195972938 | 3.63E-29 |
| 87 | ENSG00000071282 | LMCD1 | 1.404098778 | 2.171097093 | 1.96E-28 |
| 88 | ENSG00000130741 | EIF2S3 | 2.897028601 | 2.149702124 | 2.81E-28 |
| 89 | ENSG00000057019 | DCBLD2 | 2.974404329 | 2.131744703 | 1.29E-27 |
| 90 | ENSG00000064666 | CNN2 | 1.020121355 | 2.106156908 | 3.95E-27 |
| 91 | ENSG00000058668 | ATP2B4 | 1.120826448 | 2.098276439 | 5.81E-27 |
| 92 | ENSG00000131711 | MAP1B | 3.381770936 | 2.084720299 | 1.07E-26 |

| | | | | | |
|---|---|---|---|---|---|
| 93 | ENSG00000103187 | COTL1 | 1.13790474 | 2.054364308 | 5.70E-26 |
| 94 | ENSG00000145817 | YIPF5 | 1.910106727 | 2.032097052 | 1.48E-25 |
| 95 | ENSG00000095739 | BAMBI | 1.100458118 | 2.039607176 | 1.71E-25 |
| 96 | ENSG00000149591 | TAGLN | 3.855117773 | 2.036010066 | 2.12E-25 |
| 97 | ENSG00000224078 | SNHG14 | 1.087327873 | 2.016991128 | 4.52E-25 |
| 98 | ENSG00000131016 | AKAP12 | 3.458189561 | 2.00641662 | 8.55E-25 |
| 99 | ENSG00000103202 | NME4 | 1.295796687 | 1.983729484 | 1.91E-24 |
| 100 | ENSG00000143153 | ATP1B1 | 2.634902247 | 1.9616987 | 5.74E-24 |
| 101 | ENSG00000104341 | LAPTM4B | 1.376402635 | 1.957284463 | 7.35E-24 |
| 102 | ENSG00000153250 | RBMS1 | 3.461210415 | 1.942490425 | 1.23E-23 |
| 103 | ENSG00000102898 | NUTF2 | 1.981949207 | 1.941875445 | 1.33E-23 |
| 104 | ENSG00000263001 | GTF2I | 2.399485025 | 1.933252084 | 2.05E-23 |
| 105 | ENSG00000101966 | XIAP | 1.511144673 | 1.925592083 | 3.20E-23 |
| 106 | ENSG00000138448 | ITGAV | 1.723651262 | 1.905273156 | 9.12E-23 |
| 107 | ENSG00000114744 | COMMD2 | 1.029453102 | 1.887782006 | 2.19E-22 |
| 108 | ENSG00000058262 | SEC61A1 | 1.10626055 | 1.865745517 | 6.49E-22 |
| 109 | ENSG00000163939 | PBRM1 | 1.164554053 | 1.864508553 | 6.93E-22 |
| 110 | ENSG00000182512 | GLRX5 | 2.116242604 | 1.855155224 | 9.41E-22 |
| 111 | ENSG00000145247 | OCIAD2 | 1.50381686 | 1.852251064 | 1.30E-21 |
| 112 | ENSG00000090615 | GOLGA3 | 1.053769478 | 1.846333311 | 1.74E-21 |
| 113 | ENSG00000133134 | BEX2 | 1.200176199 | 1.839123326 | 2.49E-21 |
| 114 | ENSG00000100697 | DICER1 | 1.474685063 | 1.834523613 | 2.65E-21 |
| 115 | ENSG00000173473 | SMARCC1 | 1.448454482 | 1.831877953 | 3.02E-21 |
| 116 | ENSG00000137962 | ARHGAP29 | 2.128116653 | 1.830299144 | 3.77E-21 |
| 117 | ENSG00000164294 | GPX8 | 1.055982777 | 1.81580947 | 7.35E-21 |
| 118 | ENSG00000113328 | CCNG1 | 1.217792861 | 1.813620109 | 7.56E-21 |
| 119 | ENSG00000196924 | FLNA | 1.989187293 | 1.805234905 | 1.07E-20 |
| 120 | ENSG00000105568 | PPP2R1A | 1.113827639 | 1.793109486 | 1.91E-20 |
| 121 | ENSG00000072682 | P4HA2 | 1.290143802 | 1.795467834 | 1.94E-20 |
| 122 | ENSG00000069329 | VPS35 | 1.590135511 | 1.788757838 | 2.23E-20 |
| 123 | ENSG00000110092 | CCND1 | 1.789063213 | 1.775532815 | 5.55E-20 |
| 124 | ENSG00000141429 | GALNT1 | 1.103897704 | 1.769391019 | 6.22E-20 |
| 125 | ENSG00000135446 | CDK4 | 1.265318965 | 1.751858104 | 1.27E-19 |
| 126 | ENSG00000102401 | ARMCX3 | 1.231760571 | 1.751158257 | 1.36E-19 |
| 127 | ENSG00000105223 | PLD3 | 1.862640441 | 1.746864196 | 1.50E-19 |
| 128 | ENSG00000196141 | SPATS2L | 1.709205001 | 1.747787802 | 1.52E-19 |
| 129 | ENSG00000075618 | FSCN1 | 1.448843304 | 1.744835466 | 1.83E-19 |
| 130 | ENSG00000182287 | AP1S2 | 1.195420598 | 1.739492042 | 2.53E-19 |
| 131 | ENSG00000087088 | BAX | 1.015006841 | 1.732664477 | 3.30E-19 |
| 132 | ENSG00000074696 | HACD3 | 1.445074715 | 1.730091676 | 3.47E-19 |
| 133 | ENSG00000162909 | CAPN2 | 2.887517398 | 1.700469692 | 1.19E-18 |
| 134 | ENSG00000171401 | KRT13 | 2.373344172 | 1.724657805 | 1.99E-18 |
| 135 | ENSG00000241685 | ARPC1A | 1.284221733 | 1.672869422 | 4.44E-18 |
| 136 | ENSG00000170035 | UBE2E3 | 1.223146651 | 1.66954746 | 5.23E-18 |
| 137 | ENSG00000161203 | AP2M1 | 2.562222302 | 1.659373046 | 7.24E-18 |
| 138 | ENSG00000175582 | RAB6A | 1.178372216 | 1.661777344 | 7.38E-18 |
| 139 | ENSG00000111530 | CAND1 | 1.341737591 | 1.654328718 | 9.84E-18 |

| | | | | | |
|---|---|---|---|---|---|
| 140 | ENSG00000101752 | MIB1 | 1.034447978 | 1.653731341 | 1.10E-17 |
| 141 | ENSG00000135862 | LAMC1 | 1.24949687 | 1.653148589 | 1.11E-17 |
| 142 | ENSG00000196914 | ARHGEF12 | 1.017848238 | 1.647928435 | 1.41E-17 |
| 143 | ENSG00000102024 | PLS3 | 2.769345191 | 1.645095615 | 1.52E-17 |
| 144 | ENSG00000176101 | SSNA1 | 1.173167973 | 1.633283417 | 2.50E-17 |
| 145 | ENSG00000049449 | RCN1 | 2.552322276 | 1.630411945 | 2.64E-17 |
| 146 | ENSG00000205531 | NAP1L4 | 1.147296305 | 1.627412069 | 3.29E-17 |
| 147 | ENSG00000196591 | HDAC2 | 1.372663954 | 1.610860801 | 6.47E-17 |
| 148 | ENSG00000126934 | MAP2K2 | 1.525501205 | 1.607634493 | 7.28E-17 |
| 149 | ENSG00000163479 | SSR2 | 2.996597135 | 1.605112392 | 7.69E-17 |
| 150 | ENSG00000103275 | UBE2I | 2.382705828 | 1.595912747 | 1.15E-16 |
| 151 | ENSG00000178209 | PLEC | 1.528312693 | 1.58892934 | 1.99E-16 |
| 152 | ENSG00000105810 | CDK6 | 1.095493151 | 1.588424565 | 2.02E-16 |
| 153 | ENSG00000117713 | ARID1A | 1.050060707 | 1.583907104 | 2.16E-16 |
| 154 | ENSG00000102931 | ARL2BP | 1.082003992 | 1.574582755 | 3.12E-16 |
| 155 | ENSG00000197102 | DYNC1H1 | 1.645557708 | 1.572893449 | 3.32E-16 |
| 156 | ENSG00000123595 | RAB9A | 1.98718336 | 1.56727131 | 4.14E-16 |
| 157 | ENSG00000129116 | PALLD | 2.513230635 | 1.566761566 | 4.24E-16 |
| 158 | ENSG00000167522 | ANKRD11 | 2.65242918 | 1.565676405 | 4.29E-16 |
| 159 | ENSG00000197006 | METTL9 | 1.603295665 | 1.553790736 | 6.97E-16 |
| 160 | ENSG00000143545 | RAB13 | 2.828715434 | 1.552126624 | 7.26E-16 |
| 161 | ENSG00000204217 | BMPR2 | 1.013212274 | 1.551474164 | 8.48E-16 |
| 162 | ENSG00000253719 | ATXN7L3B | 1.40027037 | 1.542365311 | 1.16E-15 |
| 163 | ENSG00000113161 | HMGCR | 1.102551779 | 1.541421352 | 1.31E-15 |
| 164 | ENSG00000156113 | KCNMA1 | 1.418933862 | 1.546667888 | 1.43E-15 |
| 165 | ENSG00000160075 | SSU72 | 1.926138188 | 1.535995651 | 1.44E-15 |
| 166 | ENSG00000136026 | CKAP4 | 2.617764136 | 1.534826338 | 1.51E-15 |
| 167 | ENSG00000131435 | PDLIM4 | 1.847655811 | 1.53479172 | 1.58E-15 |
| 168 | ENSG00000112473 | SLC39A7 | 1.214293456 | 1.526677808 | 2.26E-15 |
| 169 | ENSG00000163902 | RPN1 | 1.02203556 | 1.526157833 | 2.37E-15 |
| 170 | ENSG00000155506 | LARP1 | 1.116130666 | 1.523459442 | 2.59E-15 |
| 171 | ENSG00000244038 | DDOST | 1.375295985 | 1.517929057 | 3.10E-15 |
| 172 | ENSG00000257727 | CNPY2 | 2.084568505 | 1.513682897 | 3.59E-15 |
| 173 | ENSG00000013375 | PGM3 | 1.143019255 | 1.516255402 | 3.63E-15 |
| 174 | ENSG00000101972 | STAG2 | 1.876608151 | 1.511306758 | 3.97E-15 |
| 175 | ENSG00000141367 | CLTC | 2.371699153 | 1.509147535 | 4.24E-15 |
| 176 | ENSG00000077097 | TOP2B | 1.054337758 | 1.510336842 | 4.58E-15 |
| 177 | ENSG00000172301 | COPRS | 1.284431099 | 1.49154185 | 9.22E-15 |
| 178 | ENSG00000065243 | PKN2 | 1.259366986 | 1.477230246 | 1.73E-14 |
| 179 | ENSG00000188042 | ARL4C | 1.405743798 | 1.472029934 | 2.43E-14 |
| 180 | ENSG00000116649 | SRM | 1.609726196 | 1.458279264 | 3.44E-14 |
| 181 | ENSG00000239672 | NME1 | 1.62076278 | 1.457378429 | 3.57E-14 |
| 182 | ENSG00000166347 | CYB5A | 1.176607559 | 1.459282448 | 3.66E-14 |
| 183 | ENSG00000148248 | SURF4 | 1.240015577 | 1.454484963 | 4.06E-14 |
| 184 | ENSG00000225470 | JPX | 1.54566017 | 1.450577015 | 5.04E-14 |
| 185 | ENSG00000104332 | SFRP1 | 1.185849576 | 1.451837555 | 6.14E-14 |
| 186 | ENSG00000086232 | EIF2AK1 | 1.083918196 | 1.424713293 | 1.33E-13 |

| | | | | | |
|---|---|---|---|---|---|
| 187 | ENSG00000140391 | TSPAN3 | 1.29672388 | 1.423261691 | 1.39E-13 |
| 188 | ENSG00000122218 | COPA | 1.715814988 | 1.420177031 | 1.50E-13 |
| 189 | ENSG00000054148 | PHPT1 | 2.362427226 | 1.415409173 | 1.79E-13 |
| 190 | ENSG00000198743 | SLC5A3 | 1.122531287 | 1.409300448 | 2.61E-13 |
| 191 | ENSG00000143753 | DEGS1 | 1.62506974 | 1.403279875 | 3.06E-13 |
| 192 | ENSG00000110917 | MLEC | 1.548202472 | 1.400338698 | 3.28E-13 |
| 193 | ENSG00000125304 | TM9SF2 | 1.000650308 | 1.393551442 | 4.51E-13 |
| 194 | ENSG00000035687 | ADSS | 1.15399602 | 1.386150873 | 5.87E-13 |
| 195 | ENSG00000181061 | HIGD1A | 1.6518686 | 1.386340461 | 5.88E-13 |
| 196 | ENSG00000133703 | KRAS | 1.651569506 | 1.38303556 | 6.40E-13 |
| 197 | ENSG00000166266 | CUL5 | 1.023710489 | 1.3836136 | 6.47E-13 |
| 198 | ENSG00000115306 | SPTBN1 | 2.379236332 | 1.381698532 | 6.65E-13 |
| 199 | ENSG00000176014 | TUBB6 | 1.484256084 | 1.376534788 | 8.44E-13 |
| 200 | ENSG00000071054 | MAP4K4 | 2.460111465 | 1.374036756 | 8.88E-13 |
| 201 | ENSG00000177425 | PAWR | 1.629137424 | 1.362872254 | 1.44E-12 |
| 202 | ENSG00000113648 | H2AFY | 1.928381396 | 1.35194784 | 1.99E-12 |
| 203 | ENSG00000095139 | ARCN1 | 1.630333801 | 1.351680577 | 2.02E-12 |
| 204 | ENSG00000137693 | YAP1 | 1.196317881 | 1.348836446 | 2.34E-12 |
| 205 | ENSG00000169967 | MAP3K2 | 1.273753428 | 1.348450485 | 2.37E-12 |
| 206 | ENSG00000102054 | RBBP7 | 1.304679791 | 1.344093393 | 2.75E-12 |
| 207 | ENSG00000138814 | PPP3CA | 1.016053671 | 1.342034309 | 3.07E-12 |
| 208 | ENSG00000170955 | CAVIN3 | 1.081764716 | 1.340183357 | 3.49E-12 |
| 209 | ENSG00000159210 | SNF8 | 1.336892261 | 1.32046484 | 6.39E-12 |
| 210 | ENSG00000108010 | GLRX3 | 1.543716056 | 1.319519974 | 6.65E-12 |
| 211 | ENSG00000124783 | SSR1 | 1.190395812 | 1.311543492 | 9.00E-12 |
| 212 | ENSG00000105379 | ETFB | 2.182342472 | 1.305476357 | 1.07E-11 |
| 213 | ENSG00000215021 | PHB2 | 1.193237209 | 1.303748257 | 1.19E-11 |
| 214 | ENSG00000141279 | NPEPPS | 1.712136127 | 1.298932811 | 1.40E-11 |
| 215 | ENSG00000213977 | TAX1BP3 | 1.175171905 | 1.298624729 | 1.43E-11 |
| 216 | ENSG00000136628 | EPRS | 1.10733729 | 1.298946013 | 1.44E-11 |
| 217 | ENSG00000170017 | ALCAM | 1.571980479 | 1.29608679 | 1.70E-11 |
| 218 | ENSG00000146701 | MDH2 | 1.242976612 | 1.291156189 | 1.88E-11 |
| 219 | ENSG00000101294 | HM13 | 1.506089977 | 1.285478446 | 2.28E-11 |
| 220 | ENSG00000157954 | WIPI2 | 1.036691187 | 1.285775144 | 2.29E-11 |
| 221 | ENSG00000101557 | USP14 | 1.875351954 | 1.284073843 | 2.32E-11 |
| 222 | ENSG00000145012 | LPP | 1.015455482 | 1.283233764 | 2.61E-11 |
| 223 | ENSG00000172531 | PPP1CA | 1.198620908 | 1.276868897 | 3.11E-11 |
| 224 | ENSG00000166165 | CKB | 1.721138869 | 1.281486165 | 3.32E-11 |
| 225 | ENSG00000198380 | GFPT1 | 1.067976463 | 1.275018594 | 3.39E-11 |
| 226 | ENSG00000167085 | PHB | 1.250663338 | 1.27331393 | 3.50E-11 |
| 227 | ENSG00000130313 | PGLS | 1.380829232 | 1.272893288 | 3.51E-11 |
| 228 | ENSG00000106443 | PHF14 | 1.213097079 | 1.273669109 | 3.52E-11 |
| 229 | ENSG00000112419 | PHACTR2 | 1.553227259 | 1.270561195 | 4.02E-11 |
| 230 | ENSG00000078369 | GNB1 | 1.8884822 | 1.26777335 | 4.09E-11 |
| 231 | ENSG00000173905 | GOLIM4 | 1.39383984 | 1.267138849 | 4.38E-11 |
| 232 | ENSG00000114867 | EIF4G1 | 1.905560491 | 1.259065423 | 5.61E-11 |
| 233 | ENSG00000167397 | VKORC1 | 1.493228917 | 1.259518641 | 5.62E-11 |

| | | | | | |
|---|---|---|---|---|---|
| 234 | ENSG00000064651 | SLC12A2 | 1.185909395 | 1.261672982 | 5.88E-11 |
| 235 | ENSG00000119185 | ITGB1BP1 | 1.56175145 | 1.251930398 | 7.22E-11 |
| 236 | ENSG00000253729 | PRKDC | 1.216147842 | 1.251127763 | 7.71E-11 |
| 237 | ENSG00000055609 | KMT2C | 1.289784889 | 1.249657423 | 8.30E-11 |
| 238 | ENSG00000004059 | ARF5 | 1.731487536 | 1.241373831 | 1.03E-10 |
| 239 | ENSG00000114857 | NKTR | 1.521642887 | 1.239143701 | 1.16E-10 |
| 240 | ENSG00000130520 | LSM4 | 1.312665613 | 1.237323601 | 1.23E-10 |
| 241 | ENSG00000073584 | SMARCE1 | 1.358187784 | 1.228966485 | 1.62E-10 |
| 242 | ENSG00000154639 | CXADR | 1.355226749 | 1.228985823 | 1.79E-10 |
| 243 | ENSG00000258947 | TUBB3 | 1.579697115 | 1.227883828 | 1.84E-10 |
| 244 | ENSG00000038219 | BOD1L1 | 2.154347234 | 1.224612416 | 1.86E-10 |
| 245 | ENSG00000061676 | NCKAP1 | 1.453000717 | 1.220266507 | 2.17E-10 |
| 246 | ENSG00000107854 | TNKS2 | 1.267352807 | 1.204454289 | 3.77E-10 |
| 247 | ENSG00000057608 | GDI2 | 1.761247431 | 1.202707958 | 3.80E-10 |
| 248 | ENSG00000184432 | COPB2 | 1.802971104 | 1.196604773 | 4.66E-10 |
| 249 | ENSG00000129083 | COPB1 | 1.800398891 | 1.191111025 | 5.59E-10 |
| 250 | ENSG00000094916 | CBX5 | 1.084965027 | 1.191101467 | 5.99E-10 |
| 251 | ENSG00000151135 | TMEM263 | 1.224163572 | 1.181186807 | 8.01E-10 |
| 252 | ENSG00000104549 | SQLE | 1.559867155 | 1.174410168 | 1.02E-09 |
| 253 | ENSG00000149547 | EI24 | 1.057508159 | 1.166621248 | 1.30E-09 |
| 254 | ENSG00000123908 | AGO2 | 1.289695161 | 1.16591222 | 1.34E-09 |
| 255 | ENSG00000141522 | ARHGDIA | 1.171403316 | 1.16475451 | 1.37E-09 |
| 256 | ENSG00000204272 | NBDY | 1.343352701 | 1.162517763 | 1.46E-09 |
| 257 | ENSG00000006451 | RALA | 1.47109593 | 1.160912847 | 1.56E-09 |
| 258 | ENSG00000169895 | SYAP1 | 1.476090806 | 1.158505486 | 1.66E-09 |
| 259 | ENSG00000188994 | ZNF292 | 1.327590424 | 1.143276652 | 2.87E-09 |
| 260 | ENSG00000114062 | UBE3A | 1.617921383 | 1.141503513 | 2.87E-09 |
| 261 | ENSG00000106541 | AGR2 | 1.341887138 | 1.157509733 | 3.45E-09 |
| 262 | ENSG00000212907 | MT-ND4L | 2.162243327 | 1.150891127 | 3.63E-09 |
| 263 | ENSG00000120805 | ARL1 | 1.471514662 | 1.13160168 | 3.96E-09 |
| 264 | ENSG00000077147 | TM9SF3 | 1.774108492 | 1.110548933 | 7.57E-09 |
| 265 | ENSG00000198887 | SMC5 | 1.186866497 | 1.111046919 | 7.74E-09 |
| 266 | ENSG00000160551 | TAOK1 | 1.202449317 | 1.109980168 | 7.97E-09 |
| 267 | ENSG00000113658 | SMAD5 | 1.034537707 | 1.109858675 | 8.03E-09 |
| 268 | ENSG00000197930 | ERO1A | 1.257482691 | 1.10844081 | 8.67E-09 |
| 269 | ENSG00000099901 | RANBP1 | 1.950813478 | 1.102123885 | 9.84E-09 |
| 270 | ENSG00000101266 | CSNK2A1 | 1.025445236 | 1.10076184 | 1.07E-08 |
| 271 | ENSG00000092148 | HECTD1 | 1.424586746 | 1.099318263 | 1.10E-08 |
| 272 | ENSG00000166326 | TRIM44 | 1.60736335 | 1.098693962 | 1.12E-08 |
| 273 | ENSG00000073614 | KDM5A | 1.412951973 | 1.096686111 | 1.18E-08 |
| 274 | ENSG00000137710 | RDX | 2.057769644 | 1.08879606 | 1.52E-08 |
| 275 | ENSG00000198961 | PJA2 | 1.708247899 | 1.081300862 | 1.88E-08 |
| 276 | ENSG00000175324 | LSM1 | 1.326842688 | 1.076424772 | 2.19E-08 |
| 277 | ENSG00000044115 | CTNNA1 | 1.999924782 | 1.069045884 | 2.71E-08 |
| 278 | ENSG00000136770 | DNAJC1 | 1.135691441 | 1.065004178 | 3.16E-08 |
| 279 | ENSG00000213246 | SUPT4H1 | 1.231940027 | 1.062733679 | 3.32E-08 |
| 280 | ENSG00000054523 | KIF1B | 1.325855677 | 1.066345226 | 3.63E-08 |

| | | | | | |
|---|---|---|---|---|---|
| 281 | ENSG00000149257 | SERPINH1 | 1.468254532 | 1.057158039 | 4.07E-08 |
| 282 | ENSG00000099800 | TIMM13 | 1.83913162 | 1.05287585 | 4.39E-08 |
| 283 | ENSG00000205420 | KRT6A | 1.973933477 | 1.081038661 | 4.40E-08 |
| 284 | ENSG00000168385 | SEPTIN2 | 1.897993402 | 1.050267533 | 4.71E-08 |
| 285 | ENSG00000149357 | LAMTOR1 | 1.413819347 | 1.049534038 | 4.88E-08 |
| 286 | ENSG00000173692 | PSMD1 | 1.070758042 | 1.044924073 | 5.74E-08 |
| 287 | ENSG00000134318 | ROCK2 | 1.333422766 | 1.029086774 | 9.08E-08 |
| 288 | ENSG00000105176 | URI1 | 1.34233578 | 1.019878959 | 1.17E-07 |
| 289 | ENSG00000120265 | PCMT1 | 1.714379335 | 1.018862709 | 1.20E-07 |
| 290 | ENSG00000145050 | MANF | 1.763101817 | 1.014587041 | 1.36E-07 |
| 291 | ENSG00000128699 | ORMDL1 | 1.098663551 | 1.011200868 | 1.53E-07 |
| 292 | ENSG00000102007 | PLP2 | 2.008867706 | 1.010783665 | 1.55E-07 |
| 293 | ENSG00000088832 | FKBP1A | 1.037797836 | 1.009478521 | 1.61E-07 |
| 294 | ENSG00000175166 | PSMD2 | 1.318797048 | 1.008832283 | 1.62E-07 |
| 295 | ENSG00000182827 | ACBD3 | 1.378944937 | 1.004336773 | 1.84E-07 |
| 296 | ENSG00000131626 | PPFIA1 | 1.014408652 | 0.998816322 | 2.21E-07 |
| 297 | ENSG00000130332 | LSM7 | 1.430688273 | 0.992990335 | 2.52E-07 |
| 298 | ENSG00000141002 | TCF25 | 1.612298408 | 0.991275655 | 2.66E-07 |
| 299 | ENSG00000129351 | ILF3 | 1.252368177 | 0.98843286 | 2.95E-07 |
| 300 | ENSG00000116747 | RO60 | 1.275936817 | 0.986855191 | 3.02E-07 |
| 301 | ENSG00000145919 | BOD1 | 1.065703346 | 0.986561151 | 3.06E-07 |
| 302 | ENSG00000110801 | PSMD9 | 1.602248835 | 0.984687705 | 3.15E-07 |
| 303 | ENSG00000135387 | CAPRIN1 | 1.157944067 | 0.985154272 | 3.19E-07 |
| 304 | ENSG00000119318 | RAD23B | 1.533337479 | 0.98220872 | 3.42E-07 |
| 305 | ENSG00000130706 | ADRM1 | 1.302795497 | 0.980211491 | 3.63E-07 |
| 306 | ENSG00000165280 | VCP | 1.929727321 | 0.979389565 | 3.65E-07 |
| 307 | ENSG00000102409 | BEX4 | 1.173945618 | 0.975401558 | 4.18E-07 |
| 308 | ENSG00000047849 | MAP4 | 1.732683914 | 0.972820149 | 4.42E-07 |
| 309 | ENSG00000091317 | CMTM6 | 1.121574184 | 0.971206989 | 4.70E-07 |
| 310 | ENSG00000118482 | PHF3 | 1.584213441 | 0.964455272 | 5.60E-07 |
| 311 | ENSG00000116350 | SRSF4 | 1.48231197 | 0.961893634 | 6.00E-07 |
| 312 | ENSG00000129292 | PHF20L1 | 1.15988818 | 0.96017348 | 6.39E-07 |
| 313 | ENSG00000174851 | YIF1A | 1.158602074 | 0.954656854 | 7.43E-07 |
| 314 | ENSG00000135750 | KCNK1 | 1.000710127 | 0.956089802 | 7.63E-07 |
| 315 | ENSG00000156675 | RAB11FIP1 | 1.120617082 | 0.954038222 | 8.33E-07 |
| 316 | ENSG00000152234 | ATP5F1A | 1.543327233 | 0.948809083 | 8.54E-07 |
| 317 | ENSG00000172239 | PAIP1 | 1.186836588 | 0.944398615 | 9.73E-07 |
| 318 | ENSG00000168701 | TMEM208 | 1.504833781 | 0.943725928 | 9.81E-07 |
| 319 | ENSG00000075391 | RASAL2 | 1.04240389 | 0.944055875 | 1.05E-06 |
| 320 | ENSG00000165283 | STOML2 | 1.212887712 | 0.940261777 | 1.09E-06 |
| 321 | ENSG00000100836 | PABPN1 | 1.301569209 | 0.938334334 | 1.14E-06 |
| 322 | ENSG00000143727 | ACP1 | 1.828753043 | 0.932766926 | 1.30E-06 |
| 323 | ENSG00000009954 | BAZ1B | 1.052184278 | 0.932450663 | 1.37E-06 |
| 324 | ENSG00000106263 | EIF3B | 1.087058688 | 0.930848276 | 1.42E-06 |
| 325 | ENSG00000205937 | RNPS1 | 1.689375041 | 0.926882333 | 1.54E-06 |
| 326 | ENSG00000177200 | CHD9 | 1.352026439 | 0.92601968 | 1.60E-06 |
| 327 | ENSG00000180992 | MRPL14 | 1.448783485 | 0.924676707 | 1.64E-06 |

| | | | | | |
|---|---|---|---|---|---|
| 328 | ENSG00000137100 | DCTN3 | 1.488892048 | 0.922980672 | 1.70E-06 |
| 329 | ENSG00000112701 | SENP6 | 1.502889667 | 0.921208674 | 1.80E-06 |
| 330 | ENSG00000129128 | SPCS3 | 1.554363817 | 0.919670776 | 1.87E-06 |
| 331 | ENSG00000166562 | SEC11C | 1.375924083 | 0.91950375 | 1.89E-06 |
| 332 | ENSG00000182004 | SNRPE | 1.593096546 | 0.917570414 | 1.96E-06 |
| 333 | ENSG00000075292 | ZNF638 | 1.187943237 | 0.917771681 | 2.01E-06 |
| 334 | ENSG00000149923 | PPP4C | 1.301838394 | 0.916929539 | 2.02E-06 |
| 335 | ENSG00000069956 | MAPK6 | 1.226496509 | 0.91137774 | 2.35E-06 |
| 336 | ENSG00000143106 | PSMA5 | 1.626924125 | 0.90858247 | 2.48E-06 |
| 337 | ENSG00000152082 | MZT2B | 1.860636509 | 0.908406953 | 2.49E-06 |
| 338 | ENSG00000170275 | CRTAP | 1.0649257 | 0.90891029 | 2.53E-06 |
| 339 | ENSG00000147140 | NONO | 1.65886741 | 0.904569453 | 2.76E-06 |
| 340 | ENSG00000100528 | CNIH1 | 1.019313801 | 0.90069972 | 3.11E-06 |
| 341 | ENSG00000085733 | CTTN | 1.449441493 | 0.896903964 | 3.39E-06 |
| 342 | ENSG00000169139 | UBE2V2 | 1.028107177 | 0.895898737 | 3.52E-06 |
| 343 | ENSG00000147548 | NSD3 | 1.244292627 | 0.895033136 | 3.60E-06 |
| 344 | ENSG00000011304 | PTBP1 | 1.020749454 | 0.893607394 | 3.75E-06 |
| 345 | ENSG00000131236 | CAP1 | 1.71949385 | 0.890477062 | 3.96E-06 |
| 346 | ENSG00000141759 | TXNL4A | 1.287182768 | 0.887031293 | 4.39E-06 |
| 347 | ENSG00000108829 | LRRC59 | 1.426351404 | 0.886632354 | 4.41E-06 |
| 348 | ENSG00000138760 | SCARB2 | 1.151214442 | 0.886749573 | 4.48E-06 |
| 349 | ENSG00000174437 | ATP2A2 | 1.759662231 | 0.885700627 | 4.56E-06 |
| 350 | ENSG00000243927 | MRPS6 | 1.687401018 | 0.884153866 | 4.70E-06 |
| 351 | ENSG00000198231 | DDX42 | 1.082183449 | 0.883936113 | 4.81E-06 |
| 352 | ENSG00000198563 | DDX39B | 1.624052818 | 0.87935995 | 5.28E-06 |
| 353 | ENSG00000134851 | TMEM165 | 1.78759765 | 0.875588406 | 5.79E-06 |
| 354 | ENSG00000175334 | BANF1 | 1.400360098 | 0.874792963 | 5.96E-06 |
| 355 | ENSG00000173801 | JUP | 1.697181405 | 0.873612405 | 6.44E-06 |
| 356 | ENSG00000164733 | CTSB | 1.658927228 | 0.872900245 | 6.52E-06 |
| 357 | ENSG00000117139 | KDM5B | 1.710042466 | 0.869341023 | 6.86E-06 |
| 358 | ENSG00000058272 | PPP1R12A | 1.45069769 | 0.867851259 | 7.15E-06 |
| 359 | ENSG00000124486 | USP9X | 1.043360992 | 0.862842503 | 8.22E-06 |
| 360 | ENSG00000137714 | FDX1 | 1.347928845 | 0.86114775 | 8.47E-06 |
| 361 | ENSG00000134970 | TMED7 | 1.150107793 | 0.857796622 | 9.21E-06 |
| 362 | ENSG00000168036 | CTNNB1 | 1.641579752 | 0.851019959 | 1.08E-05 |
| 363 | ENSG00000109113 | RAB34 | 1.313622715 | 0.845928309 | 1.22E-05 |
| 364 | ENSG00000037241 | RPL26L1 | 1.049761613 | 0.845614744 | 1.24E-05 |
| 365 | ENSG00000243317 | STMP1 | 1.350800152 | 0.845437185 | 1.24E-05 |
| 366 | ENSG00000106261 | ZKSCAN1 | 1.001667229 | 0.843701737 | 1.32E-05 |
| 367 | ENSG00000146066 | HIGD2A | 1.226586237 | 0.828903963 | 1.85E-05 |
| 368 | ENSG00000177697 | CD151 | 1.204692525 | 0.828708524 | 1.87E-05 |
| 369 | ENSG00000100554 | ATP6V1D | 1.740669735 | 0.8261949 | 1.96E-05 |
| 370 | ENSG00000253352 | TUG1 | 1.07084777 | 0.823346728 | 2.14E-05 |
| 371 | ENSG00000187051 | RPS19BP1 | 1.677171988 | 0.819131923 | 2.31E-05 |
| 372 | ENSG00000102390 | PBDC1 | 1.225120674 | 0.819432899 | 2.34E-05 |
| 373 | ENSG00000100592 | DAAM1 | 1.276265821 | 0.818679365 | 2.41E-05 |
| 374 | ENSG00000101596 | SMCHD1 | 1.034477888 | 0.818192324 | 2.44E-05 |

| | | | | | |
|---|---|---|---|---|---|
| 375 | ENSG00000114354 | TFG | 1.732953099 | 0.816660131 | 2.44E-05 |
| 376 | ENSG00000156256 | USP16 | 1.171971595 | 0.816271852 | 2.51E-05 |
| 377 | ENSG00000163527 | STT3B | 1.013182365 | 0.812451675 | 2.77E-05 |
| 378 | ENSG00000100129 | EIF3L | 1.157555244 | 0.811144481 | 2.84E-05 |
| 379 | ENSG00000078674 | PCM1 | 1.392284549 | 0.809771191 | 2.93E-05 |
| 380 | ENSG00000164754 | RAD21 | 1.667511238 | 0.806223966 | 3.15E-05 |
| 381 | ENSG00000134049 | IER3IP1 | 1.424467109 | 0.805197411 | 3.23E-05 |
| 382 | ENSG00000129515 | SNX6 | 1.168292734 | 0.800621575 | 3.62E-05 |
| 383 | ENSG00000108055 | SMC3 | 1.259785718 | 0.799802627 | 3.71E-05 |
| 384 | ENSG00000152700 | SAR1B | 1.094924871 | 0.799557662 | 3.73E-05 |
| 385 | ENSG00000166275 | BORCS7 | 1.096001611 | 0.797845617 | 3.88E-05 |
| 386 | ENSG00000145734 | BDP1 | 1.504833781 | 0.79498433 | 4.14E-05 |
| 387 | ENSG00000126067 | PSMB2 | 1.545181619 | 0.79173743 | 4.41E-05 |
| 388 | ENSG00000135720 | DYNC1LI2 | 1.203855061 | 0.789161776 | 4.74E-05 |
| 389 | ENSG00000197982 | C1orf122 | 1.656773749 | 0.786033281 | 5.02E-05 |
| 390 | ENSG00000123159 | GIPC1 | 1.07994024 | 0.783983602 | 5.46E-05 |
| 391 | ENSG00000110717 | NDUFS8 | 1.612089041 | 0.782431033 | 5.47E-05 |
| 392 | ENSG00000141027 | NCOR1 | 1.044706917 | 0.778405558 | 6.11E-05 |
| 393 | ENSG00000077549 | CAPZB | 1.411905143 | 0.77481323 | 6.51E-05 |
| 394 | ENSG00000118579 | MED28 | 1.014677837 | 0.771141567 | 7.15E-05 |
| 395 | ENSG00000105677 | TMEM147 | 1.023919855 | 0.769036642 | 7.50E-05 |
| 396 | ENSG00000006128 | TAC1 | 1.157585153 | 0.780452046 | 7.91E-05 |
| 397 | ENSG00000135829 | DHX9 | 1.118972063 | 0.75856078 | 9.48E-05 |
| 398 | ENSG00000130726 | TRIM28 | 1.010819519 | 0.752292008 | 0.000109348 |
| 399 | ENSG00000007372 | PAX6 | 1.461225814 | 0.753458984 | 0.000114245 |
| 400 | ENSG00000092201 | SUPT16H | 1.084157472 | 0.746974011 | 0.000121913 |
| 401 | ENSG00000147274 | RBMX | 1.609606558 | 0.744890436 | 0.000126567 |
| 402 | ENSG00000136235 | GPNMB | 1.405205428 | 0.744424199 | 0.000136402 |
| 403 | ENSG00000170242 | USP47 | 1.149629242 | 0.739148712 | 0.000144383 |
| 404 | ENSG00000108671 | PSMD11 | 1.464665399 | 0.737414347 | 0.000148338 |
| 405 | ENSG00000116489 | CAPZA1 | 1.022215016 | 0.733001125 | 0.000165759 |
| 406 | ENSG00000114446 | IFT57 | 1.185849576 | 0.730137592 | 0.000177659 |
| 407 | ENSG00000173141 | MRPL57 | 1.304560154 | 0.725843895 | 0.000191521 |
| 408 | ENSG00000071127 | WDR1 | 1.624621098 | 0.723346114 | 0.00020138 |
| 409 | ENSG00000153317 | ASAP1 | 1.053619931 | 0.716796383 | 0.000235514 |
| 410 | ENSG00000143612 | C1orf43 | 1.059123268 | 0.714975196 | 0.000242416 |
| 411 | ENSG00000054118 | THRAP3 | 1.608200814 | 0.710695218 | 0.000262383 |
| 412 | ENSG00000257103 | LSM14A | 1.104944534 | 0.706862019 | 0.000286561 |
| 413 | ENSG00000173418 | NAA20 | 1.206875914 | 0.701622728 | 0.000318366 |
| 414 | ENSG00000186832 | KRT16 | 1.088673798 | 0.709495623 | 0.00033245 |
| 415 | ENSG00000139433 | GLTP | 1.041028056 | 0.701240032 | 0.000334338 |
| 416 | ENSG00000177189 | RPS6KA3 | 1.383311716 | 0.697975144 | 0.000346864 |
| 417 | ENSG00000196072 | BLOC1S2 | 1.113678092 | 0.697446006 | 0.000347948 |
| 418 | ENSG00000144747 | TMF1 | 1.528940791 | 0.696867716 | 0.000349526 |
| 419 | ENSG00000017797 | RALBP1 | 1.193805488 | 0.691793907 | 0.00039125 |
| 420 | ENSG00000100216 | TOMM22 | 1.179149861 | 0.688748912 | 0.000414628 |
| 421 | ENSG00000138594 | TMOD3 | 1.5977026 | 0.683259154 | 0.000463593 |

| | | | | | |
|---|---|---|---|---|---|
| 422 | ENSG00000164258 | NDUFS4 | 1.143737081 | 0.680926023 | 0.000485808 |
| 423 | ENSG00000164611 | PTTG1 | 1.206636639 | 0.679937592 | 0.000506742 |
| 424 | ENSG00000168066 | SF1 | 1.594891112 | 0.678278499 | 0.000510061 |
| 425 | ENSG00000135837 | CEP350 | 1.090767459 | 0.676228719 | 0.000539202 |
| 426 | ENSG00000001630 | CYP51A1 | 1.232747582 | 0.673928913 | 0.000563827 |
| 427 | ENSG00000182944 | EWSR1 | 1.324270476 | 0.669594531 | 0.000609573 |
| 428 | ENSG00000060339 | CCAR1 | 1.02415913 | 0.665267741 | 0.000667469 |
| 429 | ENSG00000127774 | EMC6 | 1.274650711 | 0.662703427 | 0.000697499 |
| 430 | ENSG00000099795 | NDUFB7 | 1.298817541 | 0.661164595 | 0.000717971 |
| 431 | ENSG00000078140 | UBE2K | 1.301060749 | 0.66008651 | 0.000731847 |
| 432 | ENSG00000067182 | TNFRSF1A | 1.115263292 | 0.660319666 | 0.000732505 |
| 433 | ENSG00000118705 | RPN2 | 1.475582346 | 0.656842191 | 0.000779047 |
| 434 | ENSG00000168724 | DNAJC21 | 1.422612723 | 0.655707816 | 0.00079743 |
| 435 | ENSG00000115464 | USP34 | 1.057328702 | 0.654666948 | 0.000822454 |
| 436 | ENSG00000119314 | PTBP3 | 1.163746498 | 0.65264886 | 0.000859471 |
| 437 | ENSG00000103266 | STUB1 | 1.184384014 | 0.648829764 | 0.000913952 |
| 438 | ENSG00000242616 | GNG10 | 1.482640974 | 0.647023956 | 0.000942887 |
| 439 | ENSG00000136450 | SRSF1 | 1.083918196 | 0.646405749 | 0.000964661 |
| 440 | ENSG00000112697 | TMEM30A | 1.171283678 | 0.642342723 | 0.001037295 |
| 441 | ENSG00000127483 | HP1BP3 | 1.552359885 | 0.641447709 | 0.001049299 |
| 442 | ENSG00000105254 | TBCB | 1.017130411 | 0.640151975 | 0.001083245 |
| 443 | ENSG00000143222 | UFC1 | 1.183127817 | 0.637923372 | 0.001126002 |
| 444 | ENSG00000179933 | C14orf119 | 1.051436542 | 0.637691683 | 0.001135072 |
| 445 | ENSG00000089248 | ERP29 | 1.513058877 | 0.634897717 | 0.001189364 |
| 446 | ENSG00000163430 | FSTL1 | 1.165899978 | 0.630330711 | 0.00131506 |
| 447 | ENSG00000085719 | CPNE3 | 1.015096569 | 0.629279346 | 0.001334358 |
| 448 | ENSG00000133318 | RTN3 | 1.095642698 | 0.629202357 | 0.001335046 |
| 449 | ENSG00000097033 | SH3GLB1 | 1.131115297 | 0.622059943 | 0.001521335 |
| 450 | ENSG00000114978 | MOB1A | 1.510097843 | 0.616769261 | 0.001670042 |
| 451 | ENSG00000106682 | EIF4H | 1.283503906 | 0.615757273 | 0.001702224 |
| 452 | ENSG00000145833 | DDX46 | 1.413609981 | 0.611891325 | 0.001831714 |
| 453 | ENSG00000100883 | SRP54 | 1.149120781 | 0.611122766 | 0.001860645 |
| 454 | ENSG00000134453 | RBM17 | 1.15345765 | 0.602248378 | 0.002188558 |
| 455 | ENSG00000158604 | TMED4 | 1.059930823 | 0.599827197 | 0.00228786 |
| 456 | ENSG00000115540 | MOB4 | 1.135182981 | 0.59778923 | 0.002366009 |
| 457 | ENSG00000115241 | PPM1G | 1.271330763 | 0.5915939 | 0.002643404 |
| 458 | ENSG00000187079 | TEAD1 | 1.220335164 | 0.589669003 | 0.002755226 |
| 459 | ENSG00000171314 | PGAM1 | 1.357290501 | 0.58865749 | 0.002779297 |
| 460 | ENSG00000075420 | FNDC3B | 1.275159172 | 0.588669175 | 0.002788127 |
| 461 | ENSG00000108774 | RAB5C | 1.074406994 | 0.579385245 | 0.003284827 |
| 462 | ENSG00000174775 | HRAS | 1.189677985 | 0.577996073 | 0.003378477 |
| 463 | ENSG00000166557 | TMED3 | 1.18564021 | 0.575566798 | 0.003504184 |
| 464 | ENSG00000126261 | UBA2 | 1.18297827 | 0.575554388 | 0.003506317 |
| 465 | ENSG00000087502 | ERGIC2 | 1.102701326 | 0.573245198 | 0.003647003 |
| 466 | ENSG00000128534 | LSM8 | 1.261939198 | 0.566677409 | 0.004078846 |
| 467 | ENSG00000067900 | ROCK1 | 1.144993278 | 0.56674086 | 0.004096463 |
| 468 | ENSG00000155115 | GTF3C6 | 1.051286995 | 0.564866916 | 0.004219506 |

| | | | | | |
|---|---|---|---|---|---|
| 469 | ENSG00000116954 | RRAGC | 1.084007925 | 0.563250487 | 0.004347953 |
| 470 | ENSG00000113407 | TARS | 1.026103244 | 0.560921159 | 0.004543365 |
| 471 | ENSG00000146963 | LUC7L2 | 1.273573971 | 0.547203083 | 0.005675811 |
| 472 | ENSG00000065526 | SPEN | 1.01826697 | 0.542826938 | 0.006189385 |
| 473 | ENSG00000117335 | CD46 | 1.147415943 | 0.540092396 | 0.006422633 |
| 474 | ENSG00000169223 | LMAN2 | 1.112212529 | 0.53973688 | 0.00643398 |
| 475 | ENSG00000114209 | PDCD10 | 1.04886433 | 0.536473116 | 0.006815243 |
| 476 | ENSG00000143420 | ENSA | 1.249137957 | 0.534428246 | 0.007007889 |
| 477 | ENSG00000164163 | ABCE1 | 1.129021636 | 0.53401717 | 0.007068716 |
| 478 | ENSG00000145555 | MYO10 | 1.024248859 | 0.533612706 | 0.007186466 |
| 479 | ENSG00000114353 | GNAI2 | 1.350650605 | 0.529802854 | 0.007540087 |
| 480 | ENSG00000116459 | ATP5PB | 1.408764651 | 0.52646221 | 0.007951939 |
| 481 | ENSG00000130811 | EIF3G | 1.337400722 | 0.526302108 | 0.007989573 |
| 482 | ENSG00000139684 | ESD | 1.090468364 | 0.525099779 | 0.008150884 |
| 483 | ENSG00000172428 | COPS9 | 1.253474826 | 0.520524435 | 0.008743149 |
| 484 | ENSG00000189067 | LITAF | 1.272706597 | 0.519694454 | 0.008893525 |
| 485 | ENSG00000130725 | UBE2M | 1.096629709 | 0.511792183 | 0.010060594 |
| 486 | ENSG00000090621 | PABPC4 | 1.316852935 | 0.511736259 | 0.010061981 |
| 487 | ENSG00000127884 | ECHS1 | 1.131683576 | 0.505322359 | 0.011132258 |
| 488 | ENSG00000155366 | RHOC | 1.195809421 | 0.491986518 | 0.013665747 |
| 489 | ENSG00000125870 | SNRPB2 | 1.19853118 | 0.481370231 | 0.015963103 |
| 490 | ENSG00000150316 | CWC15 | 1.076052013 | 0.474190033 | 0.017773399 |
| 491 | ENSG00000119139 | TJP2 | 1.230683831 | 0.472495747 | 0.018453587 |
| 492 | ENSG00000134153 | EMC7 | 1.314519998 | 0.470589092 | 0.018687353 |
| 493 | ENSG00000143771 | CNIH4 | 1.12393703 | 0.464356946 | 0.020476753 |
| 494 | ENSG00000198818 | SFT2D1 | 1.067527822 | 0.462775387 | 0.020978995 |
| 495 | ENSG00000144566 | RAB5A | 1.205171076 | 0.45467938 | 0.023508853 |
| 496 | ENSG00000166337 | TAF10 | 1.025684512 | 0.454146948 | 0.02375132 |
| 497 | ENSG00000123066 | MED13L | 1.039592403 | 0.453693716 | 0.023957643 |
| 498 | ENSG00000153827 | TRIP12 | 1.263015938 | 0.449730628 | 0.025227679 |
| 499 | ENSG00000107341 | UBE2R2 | 1.176816925 | 0.448121716 | 0.025738254 |
| 500 | ENSG00000131507 | NDFIP1 | 1.18564021 | 0.447253734 | 0.026025305 |
| 501 | ENSG00000054654 | SYNE2 | 1.184952293 | 0.444204131 | 0.027793713 |
| 502 | ENSG00000176845 | METRNL | 1.070339309 | 0.435019223 | 0.031030971 |
| 503 | ENSG00000124214 | STAU1 | 1.273843156 | 0.433431286 | 0.031428117 |
| 504 | ENSG00000176915 | ANKLE2 | 1.161353743 | 0.431965548 | 0.032333355 |
| 505 | ENSG00000080345 | RIF1 | 1.090887096 | 0.423438607 | 0.036105206 |
| 506 | ENSG00000036257 | CUL3 | 1.275906908 | 0.422656646 | 0.036362083 |
| 507 | ENSG00000174780 | SRP72 | 1.328936349 | 0.412478709 | 0.041585369 |
| 508 | ENSG00000124614 | RPS10 | 1.062144122 | 0.409413334 | 0.043360851 |
| 509 | ENSG00000090266 | NDUFB2 | 1.288827787 | 0.403305885 | 0.046846583 |
| 510 | ENSG00000125652 | ALKBH7 | 1.279675498 | 0.394954537 | 0.052054003 |
| 511 | ENSG00000112739 | PRPF4B | 1.126808337 | 0.388827414 | 0.056482484 |
| 512 | ENSG00000100644 | HIF1A | 1.208790119 | 0.388407266 | 0.056652499 |
| 513 | ENSG00000075142 | SRI | 1.145142825 | 0.385565423 | 0.058560353 |
| 514 | ENSG00000168259 | DNAJC7 | 1.171522953 | 0.383599557 | 0.059947004 |
| 515 | ENSG00000137076 | TLN1 | 1.055534135 | 0.378280238 | 0.064093033 |

| | | | | | |
|---|---|---|---|---|---|
| 516 | ENSG00000170860 | LSM3 | 1.235917983 | 0.374359898 | 0.067102907 |
| 517 | ENSG00000117395 | EBNA1BP2 | 1.016173309 | 0.368927385 | 0.07174139 |
| 518 | ENSG00000177156 | TALDO1 | 1.012733723 | 0.368061185 | 0.072609749 |
| 519 | ENSG00000152558 | TMEM123 | 1.159469448 | 0.362109423 | 0.077871842 |
| 520 | ENSG00000102760 | RGCC | 1.037409013 | 0.359515194 | 0.081781367 |
| 521 | ENSG00000152952 | PLOD2 | 1.02846609 | 0.354014981 | 0.085870125 |
| 522 | ENSG00000106355 | LSM5 | 1.060887926 | 0.346157484 | 0.09305004 |
| 523 | ENSG00000106299 | WASL | 1.091694651 | 0.345608185 | 0.093636449 |
| 524 | ENSG00000128609 | NDUFA5 | 1.261490556 | 0.336878812 | 0.103219936 |
| 525 | ENSG00000241468 | ATP5MF | 1.168023549 | 0.330796259 | 0.110055546 |
| 526 | ENSG00000101654 | RNMT | 1.078953229 | 0.327444923 | 0.114418353 |
| 527 | ENSG00000164104 | HMGB2 | 1.034866711 | 0.326737266 | 0.116658906 |
| 528 | ENSG00000163629 | PTPN13 | 1.167335631 | 0.313114424 | 0.134311368 |
| 529 | ENSG00000099246 | RAB18 | 1.112631261 | 0.294008902 | 0.162205562 |
| 530 | ENSG00000173207 | CKS1B | 1.162280936 | 0.291374121 | 0.166946359 |
| 531 | ENSG00000124767 | GLO1 | 1.189468619 | 0.286343885 | 0.174218894 |
| 532 | ENSG00000055211 | GINM1 | 1.035913541 | 0.265522752 | 0.210926953 |
| 533 | ENSG00000186298 | PPP1CC | 1.164374597 | 0.264871455 | 0.21201355 |
| 534 | ENSG00000106244 | PDAP1 | 1.186148671 | 0.25844639 | 0.225281621 |
| 535 | ENSG00000196419 | XRCC6 | 1.136229811 | 0.254044619 | 0.23446423 |
| 536 | ENSG00000242247 | ARFGAP3 | 1.031427125 | 0.250035242 | 0.243497554 |
| 537 | ENSG00000126756 | UXT | 1.094865052 | 0.240051435 | 0.26579347 |
| 538 | ENSG00000198355 | PIM3 | 1.111225517 | 0.234651795 | 0.278761766 |
| 539 | ENSG00000060237 | WNK1 | 1.172210871 | 0.233004971 | 0.282425991 |
| 540 | ENSG00000122644 | ARL4A | 1.096719438 | 0.227538391 | 0.296856054 |
| 541 | ENSG00000132485 | ZRANB2 | 1.040370048 | 0.213950184 | 0.327372502 |
| 542 | ENSG00000157106 | SMG1 | 1.047608133 | 0.198020967 | 0.369057147 |
| 543 | ENSG00000177576 | C18orf32 | 1.08101698 | 0.197659754 | 0.369575457 |
| 544 | ENSG00000180398 | MCFD2 | 1.043331083 | 0.195234446 | 0.375615581 |
| 545 | ENSG00000104964 | TLE5 | 1.033371238 | 0.188860581 | 0.394658042 |
| 546 | ENSG00000136522 | MRPL47 | 1.021766375 | 0.182207094 | 0.414644717 |
| 547 | ENSG00000128951 | DUT | 1.0563716 | 0.180477987 | 0.420109459 |
| 548 | ENSG00000145494 | NDUFS6 | 1.043121717 | 0.160155546 | 0.483561758 |
| 549 | ENSG00000109920 | FNBP4 | 1.109730045 | 0.15343227 | 0.506035812 |
| 550 | ENSG00000153048 | CARHSP1 | 1.000889584 | 0.150797459 | 0.514617744 |
| 551 | ENSG00000177556 | ATOX1 | 1.054008754 | 0.150740425 | 0.514617744 |
| 552 | ENSG00000245694 | CRNDE | 1.044467642 | 0.150132424 | 0.516866356 |
| 553 | ENSG00000110367 | DDX6 | 1.086011857 | 0.149253793 | 0.519885489 |
| 554 | ENSG00000113580 | NR3C1 | 1.000769946 | 0.140845017 | 0.549794681 |
| 555 | ENSG00000107863 | ARHGAP21 | 1.02738935 | 0.133476388 | 0.575318732 |
| 556 | ENSG00000129235 | TXNDC17 | 1.056401509 | 0.093822772 | 0.703161893 |
| 557 | ENSG00000169738 | DCXR | 1.004508626 | 0.083713275 | 0.735444693 |
| 558 | ENSG00000145337 | PYURF | 1.000082029 | 0.073188817 | 0.77385328 |
| 559 | ENSG00000101367 | MAPRE1 | 1.015993852 | 0.034550885 | 0.922344404 |
