## Supplementary material for "Single cell RNA-seq identifies developing corneal cell fates in the human cornea organoid": Table S2

| Serial No. | FeatureID | FeatureName | Cornea.Average | Cornea.Log2.Fold.Change | Cornea.P.Value |
|---|---|---|---|---|---|
| 1 | ENSG00000189058 | APOD | 147.8889919 | 11.79861608 | 0 |
| 2 | ENSG00000108602 | ALDH3A1 | 52.70891756 | 11.62489148 | 0 |
| 3 | ENSG00000171819 | ANGPTL7 | 24.5128464 | 11.53598029 | 0 |
| 4 | ENSG00000173432 | SAA1 | 52.70593072 | 11.43604808 | 0 |
| 5 | ENSG00000139330 | KERA | 22.36419271 | 11.39841542 | 0 |
| 6 | ENSG00000173110 | HSPA6 | 14.77039346 | 10.68485753 | 0 |
| 7 | ENSG00000067048 | DDX3Y | 1.654902789 | 13.43388284 | 1.05E-307 |
| 8 | ENSG00000198542 | ITGBL1 | 9.854618509 | 9.967897664 | 2.91E-298 |
| 9 | ENSG00000034971 | MYOC | 1.825566029 | 11.99051513 | 5.67E-294 |
| 10 | ENSG00000187242 | KRT12 | 21.35711754 | 9.783905854 | 8.88E-291 |
| 11 | ENSG00000205364 | MT1M | 5.603924976 | 9.991920629 | 9.08E-291 |
| 12 | ENSG00000149968 | MMP3 | 5.70299926 | 9.861285548 | 2.67E-275 |
| 13 | ENSG00000127954 | STEAP4 | 7.364288814 | 9.309676354 | 4.63E-274 |
| 14 | ENSG00000140403 | DNAJA4 | 2.130192446 | 9.930217497 | 2.83E-272 |
| 15 | ENSG00000174080 | CTSF | 1.064810249 | 11.11967618 | 3.25E-271 |
| 16 | ENSG00000148671 | ADIRF | 9.705117384 | 9.324793575 | 2.21E-270 |
| 17 | ENSG00000205362 | MT1A | 1.841008654 | 9.932278148 | 8.89E-269 |
| 18 | ENSG00000109610 | SOD3 | 13.3082375 | 8.913100983 | 1.64E-267 |
| 19 | ENSG00000134443 | GRP | 2.706621759 | 9.161766242 | 2.59E-257 |
| 20 | ENSG00000177675 | CD163L1 | 1.053752567 | 10.24663542 | 3.78E-255 |
| 21 | ENSG00000124491 | F13A1 | 1.226671828 | 9.516468548 | 8.74E-251 |
| 22 | ENSG00000113296 | THBS4 | 4.58223333 | 8.632640654 | 3.32E-248 |
| 23 | ENSG00000149735 | GPHA2 | 2.475999194 | 10.45442307 | 3.99E-243 |
| 24 | ENSG00000000971 | CFH | 4.136780348 | 8.503909014 | 1.42E-242 |
| 25 | ENSG00000204389 | HSPA1A | 38.57195767 | 7.884097435 | 2.27E-227 |
| 26 | ENSG00000081041 | CXCL2 | 22.69115692 | 7.64625734 | 1.57E-216 |
| 27 | ENSG00000118849 | RARRES1 | 9.054302919 | 7.562887808 | 1.24E-214 |
| 28 | ENSG00000186442 | KRT3 | 1.19874165 | 8.432614775 | 1.10E-213 |
| 29 | ENSG00000147689 | FAM83A | 1.229626899 | 7.876083562 | 2.66E-208 |
| 30 | ENSG00000150782 | IL18 | 2.366661602 | 7.365018974 | 7.48E-198 |
| 31 | ENSG00000128383 | APOBEC3A | 7.600821519 | 6.660577404 | 2.96E-174 |
| 32 | ENSG00000163739 | CXCL1 | 12.13729893 | 6.362166459 | 3.03E-167 |
| 33 | ENSG00000169429 | CXCL8 | 34.9295955 | 6.374466308 | 9.95E-166 |
| 34 | ENSG00000143369 | ECM1 | 3.192365374 | 6.245270291 | 3.42E-165 |
| 35 | ENSG00000125148 | MT2A | 58.24106293 | 6.219748473 | 2.07E-164 |
| 36 | ENSG00000165949 | IFI27 | 2.884974537 | 6.172827489 | 6.79E-162 |
| 37 | ENSG00000166741 | NNMT | 17.38753697 | 6.100534203 | 5.83E-160 |
| 38 | ENSG00000136688 | IL36G | 1.945135155 | 6.352306112 | 8.78E-158 |
| 39 | ENSG00000109846 | CRYAB | 31.36104653 | 5.939497396 | 4.75E-153 |
| 40 | ENSG00000126368 | NR1D1 | 1.461107387 | 6.002483166 | 2.23E-152 |
| 41 | ENSG00000166033 | HTRA1 | 12.01833352 | 5.684913992 | 1.97E-144 |
| 42 | ENSG00000095713 | CRTAC1 | 1.43241461 | 5.753102039 | 5.35E-141 |
| 43 | ENSG00000137463 | MGARP | 3.764568619 | 5.479031224 | 1.39E-136 |
| 44 | ENSG00000196136 | SERPINA3 | 2.718823339 | 5.420147381 | 1.08E-133 |
| 45 | ENSG00000115008 | IL1A | 1.226703603 | 5.63561053 | 1.65E-132 |

| | | | | | |
|---|---|---|---|---|---|
| 46 | ENSG00000204388 | HSPA1B | 20.58991145 | 5.314003389 | 1.23E-130 |
| 47 | ENSG00000170345 | FOS | 32.95973944 | 5.296986963 | 3.06E-130 |
| 48 | ENSG00000170891 | CYTL1 | 4.491134557 | 5.296392375 | 1.54E-127 |
| 49 | ENSG00000119147 | ECRG4 | 1.944023032 | 5.244954261 | 1.97E-127 |
| 50 | ENSG00000248323 | LUCAT1 | 2.640180345 | 5.156477797 | 8.91E-124 |
| 51 | ENSG00000152583 | SPARCL1 | 1.9180629 | 5.077686505 | 3.93E-121 |
| 52 | ENSG00000124216 | SNAI1 | 1.240716355 | 5.011572117 | 8.58E-119 |
| 53 | ENSG00000169715 | MT1E | 4.296290582 | 4.983980235 | 6.87E-118 |
| 54 | ENSG00000154175 | ABI3BP | 2.121231911 | 4.965484149 | 1.37E-117 |
| 55 | ENSG00000132002 | DNAJB1 | 26.9866849 | 4.939636716 | 4.56E-117 |
| 56 | ENSG00000111799 | COL12A1 | 6.653387925 | 4.912945968 | 5.22E-116 |
| 57 | ENSG00000125740 | FOSB | 8.038426089 | 4.895125747 | 2.44E-115 |
| 58 | ENSG00000188257 | PLA2G2A | 1.850096288 | 5.296442074 | 5.99E-115 |
| 59 | ENSG00000163734 | CXCL3 | 4.695352142 | 4.981249751 | 3.84E-112 |
| 60 | ENSG00000164761 | TNFRSF11B | 2.127682225 | 4.843962852 | 4.76E-112 |
| 61 | ENSG00000073756 | PTGS2 | 14.8814469 | 4.67559158 | 1.95E-107 |
| 62 | ENSG00000120738 | EGR1 | 10.07008442 | 4.520758998 | 4.10E-102 |
| 63 | ENSG00000113070 | HBEGF | 3.613669396 | 4.527162877 | 7.56E-102 |
| 64 | ENSG00000189129 | PLAC9 | 4.788134987 | 4.43851872 | 4.93E-99 |
| 65 | ENSG00000123610 | TNFAIP6 | 1.385451239 | 4.444143351 | 6.10E-96 |
| 66 | ENSG00000186352 | ANKRD37 | 1.227466202 | 4.277633498 | 1.45E-92 |
| 67 | ENSG00000074527 | NTN4 | 1.118891208 | 4.237678539 | 8.26E-92 |
| 68 | ENSG00000169908 | TM4SF1 | 12.45063168 | 4.236081681 | 1.32E-91 |
| 69 | ENSG00000125347 | IRF1 | 5.284459659 | 4.228983784 | 2.27E-91 |
| 70 | ENSG00000206075 | SERPINB5 | 3.303958988 | 4.176289752 | 3.02E-89 |
| 71 | ENSG00000123689 | G0S2 | 4.500063317 | 4.191964984 | 7.01E-89 |
| 72 | ENSG00000187094 | CCK | 1.761189987 | 4.12922255 | 6.67E-88 |
| 73 | ENSG00000181019 | NQO1 | 8.599889402 | 4.120299288 | 7.76E-88 |
| 74 | ENSG00000186847 | KRT14 | 12.35384519 | 4.136108217 | 1.66E-87 |
| 75 | ENSG00000120129 | DUSP1 | 7.726936284 | 3.948875609 | 3.12E-82 |
| 76 | ENSG00000149131 | SERPING1 | 3.391880266 | 3.935488356 | 5.79E-82 |
| 77 | ENSG00000282885 | AL627171.2 | 1.734499031 | 3.987531872 | 2.79E-81 |
| 78 | ENSG00000144802 | NFKBIZ | 3.829929685 | 3.821389814 | 6.28E-78 |
| 79 | ENSG00000070190 | DAPP1 | 1.224860657 | 3.846971876 | 2.40E-77 |
| 80 | ENSG00000178381 | ZFAND2A | 2.765754936 | 3.787212007 | 1.17E-76 |
| 81 | ENSG00000090339 | ICAM1 | 1.179962656 | 3.762005261 | 1.65E-75 |
| 82 | ENSG00000162616 | DNAJB4 | 4.788770486 | 3.725982201 | 1.10E-74 |
| 83 | ENSG00000116132 | PRRX1 | 2.120215113 | 3.714258197 | 2.11E-74 |
| 84 | ENSG00000143333 | RGS16 | 2.09199896 | 3.730353564 | 2.78E-74 |
| 85 | ENSG00000136826 | KLF4 | 9.057448639 | 3.660107352 | 6.22E-73 |
| 86 | ENSG00000121931 | LRIF1 | 3.236055926 | 3.654005546 | 1.44E-72 |
| 87 | ENSG00000185201 | IFITM2 | 4.514425593 | 3.602863364 | 7.04E-71 |
| 88 | ENSG00000170962 | PDGFD | 1.009553616 | 3.611302098 | 7.54E-71 |
| 89 | ENSG00000108551 | RASD1 | 1.291524495 | 3.61596807 | 1.39E-70 |
| 90 | ENSG00000128016 | ZFP36 | 6.77038328 | 3.56126685 | 1.40E-69 |
| 91 | ENSG00000176046 | NUPR1 | 2.303651882 | 3.527995057 | 2.38E-68 |
| 92 | ENSG00000197989 | SNHG12 | 3.914260395 | 3.471411237 | 1.29E-66 |

| | | | | | |
|---|---|---|---|---|---|
| 93 | ENSG00000159403 | C1R | 1.191687612 | 3.460307347 | 7.03E-66 |
| 94 | ENSG00000112149 | CD83 | 1.106340104 | 3.438471225 | 6.72E-65 |
| 95 | ENSG00000175197 | DDIT3 | 4.683150562 | 3.385390789 | 7.56E-64 |
| 96 | ENSG00000175592 | FOSL1 | 2.177791317 | 3.386223042 | 1.38E-63 |
| 97 | ENSG00000148926 | ADM | 2.307401325 | 3.369785613 | 4.34E-63 |
| 98 | ENSG00000130222 | GADD45G | 1.557417252 | 3.365582786 | 2.52E-62 |
| 99 | ENSG00000155011 | DKK2 | 1.164392932 | 3.297733768 | 7.01E-61 |
| 100 | ENSG00000163661 | PTX3 | 4.208845928 | 3.351870036 | 1.50E-60 |
| 101 | ENSG00000119720 | NRDE2 | 1.208496559 | 3.492921853 | 2.85E-60 |
| 102 | ENSG00000008517 | IL32 | 1.535270114 | 3.290832069 | 3.56E-60 |
| 103 | ENSG00000108691 | CCL2 | 9.32381802 | 3.27447277 | 6.47E-60 |
| 104 | ENSG00000071967 | CYBRD1 | 1.433336084 | 3.219490311 | 1.27E-58 |
| 105 | ENSG00000123358 | NR4A1 | 1.86712766 | 3.203450831 | 6.60E-58 |
| 106 | ENSG00000187479 | C11orf96 | 5.382580695 | 3.20334815 | 5.74E-57 |
| 107 | ENSG00000118503 | TNFAIP3 | 4.479091852 | 3.153505694 | 2.02E-56 |
| 108 | ENSG00000114541 | FRMD4B | 3.231480334 | 3.161630075 | 3.96E-56 |
| 109 | ENSG00000104635 | SLC39A14 | 1.763191808 | 3.122525187 | 1.34E-55 |
| 110 | ENSG00000184205 | TSPYL2 | 2.213887656 | 3.108709415 | 3.11E-55 |
| 111 | ENSG00000132386 | SERPINF1 | 7.579881829 | 3.061809272 | 5.48E-54 |
| 112 | ENSG00000163565 | IFI16 | 2.64316719 | 3.01531601 | 1.52E-52 |
| 113 | ENSG00000151929 | BAG3 | 3.181117043 | 3.014932031 | 1.71E-52 |
| 114 | ENSG00000139318 | DUSP6 | 1.410235697 | 3.026654967 | 4.63E-52 |
| 115 | ENSG00000109321 | AREG | 3.582561723 | 3.022102918 | 2.59E-51 |
| 116 | ENSG00000185885 | IFITM1 | 2.738587356 | 2.955986164 | 1.16E-50 |
| 117 | ENSG00000173559 | NABP1 | 1.570794504 | 2.9505149 | 1.74E-50 |
| 118 | ENSG00000111912 | NCOA7 | 2.904357255 | 2.951986689 | 4.25E-50 |
| 119 | ENSG00000182326 | C1S | 1.007424694 | 2.93122981 | 9.21E-50 |
| 120 | ENSG00000277632 | CCL3 | 2.21185406 | 3.633459155 | 1.67E-49 |
| 121 | ENSG00000143322 | ABL2 | 2.799213955 | 2.859327006 | 7.16E-48 |
| 122 | ENSG00000275302 | CCL4 | 1.050320873 | 3.862103073 | 1.34E-47 |
| 123 | ENSG00000119862 | LGALSL | 2.091712985 | 2.815294127 | 2.13E-46 |
| 124 | ENSG00000114315 | HES1 | 3.493337672 | 2.802730267 | 3.29E-46 |
| 125 | ENSG00000188313 | PLSCR1 | 1.622174594 | 2.77061797 | 2.85E-45 |
| 126 | ENSG00000141753 | IGFBP4 | 4.156512591 | 2.769554943 | 3.08E-45 |
| 127 | ENSG00000185022 | MAFF | 2.538246315 | 2.74797724 | 1.06E-44 |
| 128 | ENSG00000141232 | TOB1 | 2.525345687 | 2.74474323 | 1.17E-44 |
| 129 | ENSG00000013441 | CLK1 | 2.922564299 | 2.742211261 | 1.37E-44 |
| 130 | ENSG00000198805 | PNP | 2.282235567 | 2.734429997 | 2.41E-44 |
| 131 | ENSG00000163660 | CCNL1 | 5.78097498 | 2.72391261 | 4.22E-44 |
| 132 | ENSG00000204569 | PPP1R10 | 4.613976502 | 2.723675305 | 4.46E-44 |
| 133 | ENSG00000187134 | AKR1C1 | 1.669995889 | 2.685097057 | 1.50E-42 |
| 134 | ENSG00000122691 | TWIST1 | 2.734234188 | 2.654436512 | 4.93E-42 |
| 135 | ENSG00000162772 | ATF3 | 5.703158135 | 2.642676861 | 9.39E-42 |
| 136 | ENSG00000118515 | SGK1 | 3.591363383 | 2.633013407 | 1.67E-41 |
| 137 | ENSG00000173334 | TRIB1 | 2.339239822 | 2.630871842 | 2.91E-41 |
| 138 | ENSG00000003402 | CFLAR | 1.016035705 | 2.642642046 | 2.92E-41 |
| 139 | ENSG00000276085 | CCL3L1 | 1.089880682 | 3.447616636 | 5.46E-41 |

| | | | | | |
|---|---|---|---|---|---|
| 140 | ENSG00000109099 | PMP22 | 2.376956684 | 2.618510179 | 5.51E-41 |
| 141 | ENSG00000086544 | ITPKC | 1.04606303 | 2.593706399 | 6.94E-40 |
| 142 | ENSG00000069667 | RORA | 1.333054351 | 2.562884044 | 2.07E-39 |
| 143 | ENSG00000196352 | CD55 | 4.696400715 | 2.544050478 | 4.69E-39 |
| 144 | ENSG00000166592 | RRAD | 1.766528178 | 2.542209443 | 3.63E-38 |
| 145 | ENSG00000101160 | CTSZ | 1.063539251 | 2.495672488 | 1.45E-37 |
| 146 | ENSG00000124107 | SLPI | 1.636854619 | 2.555580478 | 2.13E-37 |
| 147 | ENSG00000165494 | PCF11 | 2.416611818 | 2.480865427 | 2.18E-37 |
| 148 | ENSG00000083799 | CYLD | 1.846950569 | 2.44736563 | 1.80E-36 |
| 149 | ENSG00000113369 | ARRDC3 | 2.228440582 | 2.447105157 | 2.05E-36 |
| 150 | ENSG00000125538 | IL1B | 1.863664191 | 2.974462041 | 2.89E-36 |
| 151 | ENSG00000121742 | GJB6 | 1.123021951 | 2.463872275 | 4.07E-36 |
| 152 | ENSG00000144136 | SLC20A1 | 3.111847658 | 2.398134737 | 3.41E-35 |
| 153 | ENSG00000136997 | MYC | 1.345192381 | 2.388944877 | 8.28E-35 |
| 154 | ENSG00000021355 | SERPINB1 | 2.944552562 | 2.377723551 | 1.37E-34 |
| 155 | ENSG00000185633 | NDUFA4L2 | 2.852500541 | 2.370416044 | 2.26E-34 |
| 156 | ENSG00000138166 | DUSP5 | 1.281896686 | 2.353413377 | 7.14E-34 |
| 157 | ENSG00000106546 | AHR | 2.49973508 | 2.332891008 | 1.57E-33 |
| 158 | ENSG00000135047 | CTSL | 3.971042225 | 2.330042488 | 2.55E-33 |
| 159 | ENSG00000110172 | CHORDC1 | 2.650284778 | 2.325778854 | 2.63E-33 |
| 160 | ENSG00000162783 | IER5 | 1.960990853 | 2.327223442 | 2.72E-33 |
| 161 | ENSG00000111859 | NEDD9 | 1.510994054 | 2.303200653 | 2.42E-32 |
| 162 | ENSG00000134107 | BHLHE40 | 1.782574526 | 2.287721999 | 2.50E-32 |
| 163 | ENSG00000184557 | SOCS3 | 1.623096067 | 2.273385574 | 6.98E-32 |
| 164 | ENSG00000112110 | MRPL18 | 4.162422731 | 2.266392052 | 9.11E-32 |
| 165 | ENSG00000232956 | SNHG15 | 1.671393987 | 2.257992879 | 1.36E-31 |
| 166 | ENSG00000115009 | CCL20 | 1.058042185 | 2.978298859 | 2.82E-31 |
| 167 | ENSG00000122861 | PLAU | 2.306479852 | 2.264823247 | 3.51E-31 |
| 168 | ENSG00000116679 | IVNS1ABP | 1.963088 | 2.196844133 | 4.71E-30 |
| 169 | ENSG00000197943 | PLCG2 | 2.511777785 | 2.263768663 | 1.58E-29 |
| 170 | ENSG00000099968 | BCL2L13 | 1.724617023 | 2.150892085 | 5.98E-29 |
| 171 | ENSG00000204103 | MAFB | 3.127194958 | 2.149807434 | 6.11E-29 |
| 172 | ENSG00000105939 | ZC3HAV1 | 1.111424096 | 2.153754182 | 6.21E-29 |
| 173 | ENSG00000169242 | EFNA1 | 1.256000105 | 2.107463375 | 9.12E-28 |
| 174 | ENSG00000159388 | BTG2 | 2.271400311 | 2.085934461 | 2.48E-27 |
| 175 | ENSG00000153071 | DAB2 | 1.719151732 | 2.061414605 | 7.56E-27 |
| 176 | ENSG00000163584 | RPL22L1 | 3.047852915 | 2.046608929 | 1.46E-26 |
| 177 | ENSG00000127528 | KLF2 | 1.466286704 | 2.047175592 | 2.17E-26 |
| 178 | ENSG00000162407 | PLPP3 | 2.287605533 | 2.023606178 | 5.79E-26 |
| 179 | ENSG00000006652 | IFRD1 | 3.185120686 | 2.005388454 | 1.29E-25 |
| 180 | ENSG00000246705 | H2AFJ | 1.523290959 | 2.004399262 | 1.54E-25 |
| 181 | ENSG00000197329 | PELI1 | 1.691253328 | 1.951106287 | 2.52E-24 |
| 182 | ENSG00000151632 | AKR1C2 | 1.382432619 | 1.951676822 | 3.45E-24 |
| 183 | ENSG00000117616 | RSRP1 | 3.253087298 | 1.91033237 | 1.86E-23 |
| 184 | ENSG00000117143 | UAP1 | 2.497097759 | 1.902030439 | 2.80E-23 |
| 185 | ENSG00000117228 | GBP1 | 1.036244571 | 1.907861773 | 5.50E-23 |
| 186 | ENSG00000100591 | AHSA1 | 2.521342044 | 1.864771518 | 1.85E-22 |

| | | | | | |
|---|---|---|---|---|---|
| 187 | ENSG00000165092 | ALDH1A1 | 1.788452891 | 1.866278352 | 2.64E-22 |
| 188 | ENSG00000103642 | LACTB | 1.388056784 | 1.855352686 | 3.48E-22 |
| 189 | ENSG00000171867 | PRNP | 2.764992337 | 1.831437156 | 1.01E-21 |
| 190 | ENSG00000196628 | TCF4 | 2.248141049 | 1.81125356 | 2.68E-21 |
| 191 | ENSG00000206573 | THUMPD3-AS1 | 2.569512863 | 1.80640906 | 3.38E-21 |
| 192 | ENSG00000157557 | ETS2 | 3.405956568 | 1.78356949 | 9.76E-21 |
| 193 | ENSG00000169155 | ZBTB43 | 1.784766997 | 1.7737761 | 1.82E-20 |
| 194 | ENSG00000186834 | HEXIM1 | 2.380261279 | 1.768979254 | 2.09E-20 |
| 195 | ENSG00000132823 | OSER1 | 1.677526551 | 1.754506539 | 3.91E-20 |
| 196 | ENSG00000078401 | EDN1 | 1.201124771 | 1.775299597 | 4.30E-20 |
| 197 | ENSG00000106624 | AEBP1 | 1.320852771 | 1.751061035 | 5.26E-20 |
| 198 | ENSG00000175906 | ARL4D | 1.79537983 | 1.750119108 | 5.74E-20 |
| 199 | ENSG00000124145 | SDC4 | 2.340065971 | 1.740028904 | 8.07E-20 |
| 200 | ENSG00000122257 | RBBP6 | 3.048520189 | 1.720954862 | 1.86E-19 |
| 201 | ENSG00000158270 | COLEC12 | 1.30388495 | 1.715899033 | 2.70E-19 |
| 202 | ENSG00000163738 | MTHFD2L | 1.077393128 | 1.712064095 | 3.66E-19 |
| 203 | ENSG00000157514 | TSC22D3 | 2.710307653 | 1.694268049 | 7.08E-19 |
| 204 | ENSG00000105835 | NAMPT | 3.165960393 | 1.683878753 | 1.08E-18 |
| 205 | ENSG00000196428 | TSC22D2 | 2.166066361 | 1.680802585 | 1.45E-18 |
| 206 | ENSG00000176597 | B3GNT5 | 1.108310151 | 1.684209776 | 1.72E-18 |
| 207 | ENSG00000137267 | TUBB2A | 1.406232054 | 1.674532183 | 1.73E-18 |
| 208 | ENSG00000186594 | MIR22HG | 3.086427701 | 1.661962792 | 2.85E-18 |
| 209 | ENSG00000135919 | SERPINE2 | 2.739159305 | 1.665953004 | 3.19E-18 |
| 210 | ENSG00000107968 | MAP3K8 | 1.407344177 | 1.655924737 | 4.32E-18 |
| 211 | ENSG00000104419 | NDRG1 | 1.349609098 | 1.656170734 | 4.60E-18 |
| 212 | ENSG00000123975 | CKS2 | 2.538977139 | 1.647722943 | 6.01E-18 |
| 213 | ENSG00000158716 | DUSP23 | 1.343317659 | 1.628918855 | 1.34E-17 |
| 214 | ENSG00000132510 | KDM6B | 1.918761949 | 1.622030284 | 1.91E-17 |
| 215 | ENSG00000171940 | ZNF217 | 1.249645115 | 1.609967847 | 3.51E-17 |
| 216 | ENSG00000050165 | DKK3 | 1.229150274 | 1.603714435 | 4.22E-17 |
| 217 | ENSG00000065618 | COL17A1 | 2.686889517 | 1.607346182 | 4.34E-17 |
| 218 | ENSG00000119138 | KLF9 | 1.146884937 | 1.603373101 | 4.68E-17 |
| 219 | ENSG00000189266 | PNRC2 | 2.5774566 | 1.574424802 | 1.35E-16 |
| 220 | ENSG00000213145 | CRIP1 | 1.184411149 | 1.574540776 | 1.62E-16 |
| 221 | ENSG00000250722 | SELENOP | 2.392971258 | 1.564091403 | 2.31E-16 |
| 222 | ENSG00000196878 | LAMB3 | 1.019658049 | 1.573225428 | 2.66E-16 |
| 223 | ENSG00000035862 | TIMP2 | 2.517815024 | 1.545293659 | 4.96E-16 |
| 224 | ENSG00000102786 | INTS6 | 1.87310135 | 1.535880097 | 8.29E-16 |
| 225 | ENSG00000118523 | CCN2 | 2.22977513 | 1.524400103 | 1.57E-15 |
| 226 | ENSG00000197019 | SERTAD1 | 2.267714417 | 1.499597115 | 3.54E-15 |
| 227 | ENSG00000163435 | ELF3 | 2.075475987 | 1.508735577 | 3.94E-15 |
| 228 | ENSG00000134202 | GSTM3 | 1.359427557 | 1.496759139 | 4.04E-15 |
| 229 | ENSG00000058085 | LAMC2 | 1.098332818 | 1.508304162 | 4.20E-15 |
| 230 | ENSG00000049759 | NEDD4L | 1.003516376 | 1.492565943 | 5.48E-15 |
| 231 | ENSG00000107290 | SETX | 2.76905953 | 1.485432107 | 1.07E-14 |
| 232 | ENSG00000118689 | FOXO3 | 1.079490274 | 1.471985245 | 1.15E-14 |
| 233 | ENSG00000165997 | ARL5B | 2.510125487 | 1.468499345 | 1.39E-14 |

| | | | | | |
|---|---|---|---|---|---|
| 234 | ENSG00000134853 | PDGFRA | 1.094742249 | 1.464659132 | 1.67E-14 |
| 235 | ENSG00000148175 | STOM | 1.217298219 | 1.447990559 | 3.01E-14 |
| 236 | ENSG00000116161 | CACYBP | 2.197364684 | 1.44234458 | 3.71E-14 |
| 237 | ENSG00000140836 | ZFHX3 | 1.398065893 | 1.44112335 | 4.11E-14 |
| 238 | ENSG00000142867 | BCL10 | 1.986347261 | 1.432044811 | 5.69E-14 |
| 239 | ENSG00000007944 | MYLIP | 1.07567728 | 1.414187335 | 1.21E-13 |
| 240 | ENSG00000243147 | MRPL33 | 1.292477744 | 1.383361086 | 3.98E-13 |
| 241 | ENSG00000152377 | SPOCK1 | 1.698593341 | 1.365305839 | 7.61E-13 |
| 242 | ENSG00000137947 | GTF2B | 2.008557949 | 1.353377335 | 1.15E-12 |
| 243 | ENSG00000105856 | HBP1 | 1.365909646 | 1.346753997 | 1.53E-12 |
| 244 | ENSG00000281991 | TMEM265 | 1.69659152 | 1.344167749 | 1.94E-12 |
| 245 | ENSG00000138279 | ANXA7 | 1.205795688 | 1.330483455 | 2.82E-12 |
| 246 | ENSG00000177426 | TGIF1 | 2.131495219 | 1.330066544 | 2.82E-12 |
| 247 | ENSG00000148634 | HERC4 | 1.210021756 | 1.322272816 | 4.26E-12 |
| 248 | ENSG00000186660 | ZFP91 | 2.22977513 | 1.314435283 | 5.05E-12 |
| 249 | ENSG00000115641 | FHL2 | 1.442519043 | 1.314931371 | 5.37E-12 |
| 250 | ENSG00000222041 | CYTOR | 1.194166058 | 1.314253718 | 5.76E-12 |
| 251 | ENSG00000240583 | AQP1 | 1.537653235 | 1.310085918 | 6.54E-12 |
| 252 | ENSG00000109320 | NFKB1 | 1.376236504 | 1.294580095 | 1.15E-11 |
| 253 | ENSG00000178695 | KCTD12 | 1.101955162 | 1.282971013 | 1.80E-11 |
| 254 | ENSG00000136158 | SPRY2 | 1.026108363 | 1.276425266 | 2.38E-11 |
| 255 | ENSG00000069020 | MAST4 | 1.497998101 | 1.270306387 | 3.04E-11 |
| 256 | ENSG00000151247 | EIF4E | 1.758997515 | 1.264323542 | 3.20E-11 |
| 257 | ENSG00000174718 | RESF1 | 1.247643294 | 1.266129985 | 3.59E-11 |
| 258 | ENSG00000175984 | DENND2C | 1.573082301 | 1.248031829 | 6.75E-11 |
| 259 | ENSG00000168209 | DDIT4 | 1.759728339 | 1.245781427 | 6.87E-11 |
| 260 | ENSG00000013588 | GPRC5A | 2.148939665 | 1.239855692 | 8.94E-11 |
| 261 | ENSG00000115884 | SDC1 | 1.548901566 | 1.236933245 | 9.31E-11 |
| 262 | ENSG00000115520 | COQ10B | 1.306268071 | 1.231120082 | 1.03E-10 |
| 263 | ENSG00000183722 | LHFPL6 | 1.272713727 | 1.228186409 | 1.17E-10 |
| 264 | ENSG00000089327 | FXYD5 | 1.13875055 | 1.22713427 | 1.22E-10 |
| 265 | ENSG00000173221 | GLRX | 1.303313001 | 1.220890685 | 1.63E-10 |
| 266 | ENSG00000164463 | CREBRF | 1.089912457 | 1.217297348 | 1.73E-10 |
| 267 | ENSG00000118985 | ELL2 | 1.134969331 | 1.212787516 | 2.16E-10 |
| 268 | ENSG00000117000 | RLF | 1.092994627 | 1.208541535 | 2.29E-10 |
| 269 | ENSG00000155090 | KLF10 | 2.248617674 | 1.177506176 | 6.39E-10 |
| 270 | ENSG00000128590 | DNAJB9 | 1.3185332 | 1.170396602 | 8.37E-10 |
| 271 | ENSG00000154734 | ADAMTS1 | 1.065795272 | 1.170271977 | 1.05E-09 |
| 272 | ENSG00000026508 | CD44 | 2.172453126 | 1.160941241 | 1.18E-09 |
| 273 | ENSG00000164442 | CITED2 | 1.831825694 | 1.160034584 | 1.20E-09 |
| 274 | ENSG00000255717 | SNHG1 | 1.485955396 | 1.131389017 | 2.97E-09 |
| 275 | ENSG00000055044 | NOP58 | 1.329304907 | 1.126931241 | 3.42E-09 |
| 276 | ENSG00000244879 | GABPB1-AS1 | 1.213612325 | 1.124591744 | 4.22E-09 |
| 277 | ENSG00000267368 | UPK3BL1 | 2.07967028 | 1.123692634 | 5.22E-09 |
| 278 | ENSG00000113083 | LOX | 1.578420492 | 1.111240209 | 6.02E-09 |
| 279 | ENSG00000100911 | PSME2 | 1.04593593 | 1.105456655 | 6.94E-09 |
| 280 | ENSG00000109220 | CHIC2 | 1.648516025 | 1.104517766 | 7.03E-09 |

| | | | | | |
|---|---|---|---|---|---|
| 281 | ENSG00000182534 | MXRA7 | 1.83128552 | 1.099604132 | 8.17E-09 |
| 282 | ENSG00000148516 | ZEB1 | 1.043870558 | 1.094285228 | 1.01E-08 |
| 283 | ENSG00000143337 | TOR1AIP1 | 1.138273926 | 1.061027618 | 2.79E-08 |
| 284 | ENSG00000164949 | GEM | 1.578039192 | 1.06103353 | 3.17E-08 |
| 285 | ENSG00000123983 | ACSL3 | 2.052439151 | 1.039270076 | 5.18E-08 |
| 286 | ENSG00000143514 | TP53BP2 | 1.22603633 | 1.022723322 | 9.19E-08 |
| 287 | ENSG00000135334 | AKIRIN2 | 1.847967367 | 1.014417026 | 1.09E-07 |
| 288 | ENSG00000126709 | IFI6 | 1.205636814 | 1.004823847 | 1.62E-07 |
| 289 | ENSG00000068305 | MEF2A | 1.271093204 | 1.001869645 | 1.62E-07 |
| 290 | ENSG00000005483 | KMT2E | 1.886542153 | 0.989745838 | 2.25E-07 |
| 291 | ENSG00000113742 | CPEB4 | 1.419768181 | 0.949938058 | 7.05E-07 |
| 292 | ENSG00000150938 | CRIM1 | 1.165282631 | 0.948688577 | 7.35E-07 |
| 293 | ENSG00000253738 | OTUD6B-AS1 | 1.064905573 | 0.932607919 | 1.14E-06 |
| 294 | ENSG00000130164 | LDLR | 1.357489285 | 0.93157988 | 1.17E-06 |
| 295 | ENSG00000133773 | CCDC59 | 1.166744278 | 0.926258991 | 1.33E-06 |
| 296 | ENSG00000185129 | PURA | 1.259908423 | 0.920679799 | 1.56E-06 |
| 297 | ENSG00000116731 | PRDM2 | 1.071832512 | 0.91879207 | 1.65E-06 |
| 298 | ENSG00000147872 | PLIN2 | 1.58836605 | 0.913969107 | 1.97E-06 |
| 299 | ENSG00000204138 | PHACTR4 | 1.036117471 | 0.911988218 | 2.04E-06 |
| 300 | ENSG00000116774 | OLFML3 | 1.112282019 | 0.902990026 | 2.57E-06 |
| 301 | ENSG00000126653 | NSRP1 | 1.279322916 | 0.891723591 | 3.33E-06 |
| 302 | ENSG00000164548 | TRA2A | 1.618202725 | 0.889134548 | 3.55E-06 |
| 303 | ENSG00000089356 | FXYD3 | 1.480839629 | 0.877913798 | 5.10E-06 |
| 304 | ENSG00000152484 | USP12 | 1.065000898 | 0.8736392 | 5.44E-06 |
| 305 | ENSG00000123384 | LRP1 | 1.238682758 | 0.859679694 | 7.77E-06 |
| 306 | ENSG00000157654 | PALM2-AKAP2 | 1.221873812 | 0.859122253 | 8.02E-06 |
| 307 | ENSG00000179119 | SPTY2D1 | 1.339631765 | 0.848266595 | 1.02E-05 |
| 308 | ENSG00000169641 | LUZP1 | 1.048064851 | 0.832585831 | 1.52E-05 |
| 309 | ENSG00000145390 | USP53 | 1.592337918 | 0.82878808 | 1.69E-05 |
| 310 | ENSG00000198730 | CTR9 | 1.044188308 | 0.823432362 | 1.89E-05 |
| 311 | ENSG00000155957 | TMBIM4 | 1.481157379 | 0.802138546 | 3.09E-05 |
| 312 | ENSG00000145779 | TNFAIP8 | 1.127375119 | 0.799499269 | 3.63E-05 |
| 313 | ENSG00000173848 | NET1 | 1.470163247 | 0.783891714 | 4.87E-05 |
| 314 | ENSG00000027697 | IFNGR1 | 1.162391111 | 0.770375025 | 6.58E-05 |
| 315 | ENSG00000235162 | C12orf75 | 1.089435832 | 0.760141388 | 8.50E-05 |
| 316 | ENSG00000083896 | YTHDC1 | 1.487194619 | 0.75700431 | 8.81E-05 |
| 317 | ENSG00000057704 | TMCC3 | 1.120225756 | 0.750701216 | 0.000111224 |
| 318 | ENSG00000153922 | CHD1 | 1.513122976 | 0.725069417 | 0.000177885 |
| 319 | ENSG00000111817 | DSE | 1.120924805 | 0.725079485 | 0.000181759 |
| 320 | ENSG00000269028 | MTRNR2L12 | 1.323617192 | 0.732900879 | 0.000188496 |
| 321 | ENSG00000059804 | SLC2A3 | 1.268805408 | 0.719530739 | 0.000207957 |
| 322 | ENSG00000102908 | NFAT5 | 1.320217272 | 0.717911077 | 0.00020833 |
| 323 | ENSG00000067334 | DNTTIP2 | 1.105069106 | 0.714103456 | 0.000227169 |
| 324 | ENSG00000163171 | CDC42EP3 | 1.062681327 | 0.712512175 | 0.000242652 |
| 325 | ENSG00000093167 | LRRFIP2 | 1.446872211 | 0.705823619 | 0.000271965 |
| 326 | ENSG00000164023 | SGMS2 | 1.036562321 | 0.707139051 | 0.000272277 |
| 327 | ENSG00000109814 | UGDH | 1.231787595 | 0.681245199 | 0.000451434 |

| | | | | | |
|---|---|---|---|---|---|
| 328 | ENSG00000116752 | BCAS2 | 1.213326351 | 0.675160562 | 0.000507707 |
| 329 | ENSG00000134757 | DSG3 | 1.326540487 | 0.674409901 | 0.000552004 |
| 330 | ENSG00000100603 | SNW1 | 1.131537637 | 0.663484191 | 0.000640688 |
| 331 | ENSG00000115816 | CEBPZ | 1.146821387 | 0.658820775 | 0.000705952 |
| 332 | ENSG00000234545 | FAM133B | 1.368737616 | 0.650114121 | 0.00083563 |
| 333 | ENSG00000108588 | CCDC47 | 1.532315044 | 0.646321413 | 0.000893461 |
| 334 | ENSG00000112715 | VEGFA | 1.188446567 | 0.641131341 | 0.001014804 |
| 335 | ENSG00000168303 | MPLKIP | 1.114379166 | 0.626299567 | 0.001326107 |
| 336 | ENSG00000161813 | LARP4 | 1.043489259 | 0.593642667 | 0.002431934 |
| 337 | ENSG00000162642 | C1orf52 | 1.043425709 | 0.572370203 | 0.003537886 |
| 338 | ENSG00000136003 | ISCU | 1.272872601 | 0.566689587 | 0.003888525 |
| 339 | ENSG00000185950 | IRS2 | 1.253299234 | 0.547851921 | 0.005422386 |
| 340 | ENSG00000072401 | UBE2D1 | 1.209545132 | 0.531688754 | 0.007073769 |
| 341 | ENSG00000183283 | DAZAP2 | 1.268392334 | 0.528054804 | 0.007463966 |
| 342 | ENSG00000188342 | GTF2F2 | 1.434702406 | 0.524075058 | 0.008021058 |
| 343 | ENSG00000121039 | RDH10 | 1.092581552 | 0.521725118 | 0.00841493 |
| 344 | ENSG00000196199 | MPHOSPH8 | 1.066970945 | 0.518858379 | 0.008703343 |
| 345 | ENSG00000075426 | FOSL2 | 1.017402028 | 0.511362979 | 0.009821178 |
| 346 | ENSG00000169439 | SDC2 | 1.110947472 | 0.509769905 | 0.010082555 |
| 347 | ENSG00000117691 | NENF | 1.285995655 | 0.504547145 | 0.010876025 |
| 348 | ENSG00000133104 | SPART | 1.205986338 | 0.491329736 | 0.013331768 |
| 349 | ENSG00000005893 | LAMP2 | 1.09397965 | 0.469376329 | 0.018536894 |
| 350 | ENSG00000108510 | MED13 | 1.169032075 | 0.469040199 | 0.018646142 |
| 351 | ENSG00000145912 | NHP2 | 1.086353663 | 0.45454146 | 0.022975275 |
| 352 | ENSG00000005810 | MYCBP2 | 1.063030852 | 0.452205076 | 0.023826772 |
| 353 | ENSG00000220205 | VAMP2 | 1.079299624 | 0.443457 | 0.026808225 |
| 354 | ENSG00000258289 | CHURC1 | 1.27967244 | 0.441187671 | 0.027598392 |
| 355 | ENSG00000255112 | CHMP1B | 1.347670827 | 0.431330681 | 0.031830903 |
| 356 | ENSG00000131503 | ANKHD1 | 1.114283841 | 0.426450956 | 0.033885026 |
| 357 | ENSG00000076053 | RBM7 | 1.078441701 | 0.425607933 | 0.034233946 |
| 358 | ENSG00000086062 | B4GALT1 | 1.249645115 | 0.425454743 | 0.0343219 |
| 359 | ENSG00000086666 | ZFAND6 | 1.218664542 | 0.423351243 | 0.035411268 |
| 360 | ENSG00000134046 | MBD2 | 1.191655837 | 0.412208076 | 0.041025529 |
| 361 | ENSG00000057757 | PITHD1 | 1.274238924 | 0.411437255 | 0.041278754 |
| 362 | ENSG00000166233 | ARIH1 | 1.155209973 | 0.409988096 | 0.04220919 |
| 363 | ENSG00000168288 | MMADHC | 1.211165654 | 0.401541906 | 0.04695797 |
| 364 | ENSG00000121579 | NAA50 | 1.093947875 | 0.36126511 | 0.07720048 |
| 365 | ENSG00000180957 | PITPNB | 1.110502622 | 0.351311445 | 0.086691342 |
| 366 | ENSG00000105355 | PLIN3 | 1.166140554 | 0.346841796 | 0.090959102 |
| 367 | ENSG00000070081 | NUCB2 | 1.20163317 | 0.313619447 | 0.131271621 |
| 368 | ENSG00000162368 | CMPK1 | 1.062045828 | 0.305705791 | 0.142570206 |
| 369 | ENSG00000134152 | KATNBL1 | 1.071038138 | 0.302434947 | 0.148332982 |
| 370 | ENSG00000213523 | SRA1 | 1.054451616 | 0.297715128 | 0.154768201 |
| 371 | ENSG00000111615 | KRR1 | 1.167602202 | 0.293528519 | 0.161209327 |
| 372 | ENSG00000152601 | MBNL1 | 1.033925 | 0.293344735 | 0.161779324 |
| 373 | ENSG00000130429 | ARPC1B | 1.156036121 | 0.283430455 | 0.177267136 |
| 374 | ENSG00000168610 | STAT3 | 1.021024372 | 0.283385813 | 0.177483278 |

| | | | | | |
|---|---|---|---|---|---|
| 375 | ENSG00000117614 | SYF2 | 1.081555646 | 0.281342544 | 0.180763091 |
| 376 | ENSG00000152492 | CCDC50 | 1.16289951 | 0.280973249 | 0.181451182 |
| 377 | ENSG00000171456 | ASXL1 | 1.170398397 | 0.280988966 | 0.181549346 |
| 378 | ENSG00000165416 | SUGT1 | 1.134238508 | 0.242314416 | 0.258500816 |
| 379 | ENSG00000169905 | TOR1AIP2 | 1.07914075 | 0.230053706 | 0.287888307 |
| 380 | ENSG00000117984 | CTSD | 1.006249021 | 0.229105525 | 0.291062701 |
| 381 | ENSG00000104131 | EIF3J | 1.13913185 | 0.219788154 | 0.31226028 |
| 382 | ENSG00000168264 | IRF2BP2 | 1.032685777 | 0.215219945 | 0.322743077 |
| 383 | ENSG00000159216 | RUNX1 | 1.121496754 | 0.201830233 | 0.356205128 |
| 384 | ENSG00000119541 | VPS4B | 1.019181425 | 0.129270228 | 0.584376939 |
| 385 | ENSG00000022267 | FHL1 | 1.011333013 | 0.120527715 | 0.602230226 |
| 386 | ENSG00000100519 | PSMC6 | 1.057216036 | 0.103331136 | 0.664815018 |
| 387 | ENSG00000142634 | EFHD2 | 1.02493269 | 0.084872876 | 0.729633363 |
| 388 | ENSG00000125356 | NDUFA1 | 1.006534996 | 0.069767854 | 0.784381317 |
| 389 | ENSG00000129824 | RPS4Y1 | 1.833223792 | 14.90344468 | 7.78E-319 |
