## Supplementary material for "Single cell RNA-seq identifies developing corneal cell fates in the human cornea organoid": Table S3

| Serial No. | FeatureID | FeatureName | Cornea.Average | Cornea.Log2.Fold.Change | Cornea.P.Value | Organoid.Average | Organoid.Log2.Fold.Change | Organoid.P.Value |
|---|---|---|---|---|---|---|---|---|
| 1 | ENSG00000107317 | PTGDS | 217.88 | 7.7618 | 5.58E-223 | 1.003850619 | -7.7617876 | 5.58E-223 |
| 2 | ENSG00000011465 | DCN | 278.99 | 5.0107 | 8.78E-120 | 8.653998094 | -5.01070716 | 8.78E-120 |
| 3 | ENSG00000120885 | CLU | 22.889 | 4.3337 | 4.81E-95 | 1.135093252 | -4.3337122 | 4.81E-95 |
| 4 | ENSG00000101439 | CST3 | 48.059 | 4.2971 | 1.88E-94 | 2.444558555 | -4.29714148 | 1.88E-94 |
| 5 | ENSG00000187193 | MT1X | 21.514 | 4.2782 | 1.47E-93 | 1.108772943 | -4.27818702 | 1.47E-93 |
| 6 | ENSG00000099860 | GADD45B | 21.319 | 4.1237 | 1.87E-88 | 1.222907376 | -4.12371176 | 1.87E-88 |
| 7 | ENSG00000196154 | S100A4 | 17.585 | 3.957 | 1.54E-82 | 1.132311674 | -3.95698386 | 1.54E-82 |
| 8 | ENSG00000164692 | COL1A2 | 1.2368 | -3.8097 | 3.43E-74 | 17.3444558 | 3.809749186 | 3.43E-74 |
| 9 | ENSG00000113140 | SPARC | 1.6167 | -3.5774 | 2.13E-67 | 19.30026415 | 3.577436612 | 2.13E-67 |
| 10 | ENSG00000167779 | IGFBP6 | 16.211 | 3.2511 | 9.03E-60 | 1.702744562 | -3.25105125 | 9.03E-60 |
| 11 | ENSG00000160888 | IER2 | 12.553 | 3.1906 | 6.00E-58 | 1.374966981 | -3.19055601 | 6.00E-58 |
| 12 | ENSG00000087074 | PPP1R15A | 16.255 | 3.1816 | 1.05E-57 | 1.791515787 | -3.18161949 | 1.05E-57 |
| 13 | ENSG00000137331 | IER3 | 28.511 | 3.1575 | 1.03E-56 | 3.19534538 | -3.15747598 | 1.03E-56 |
| 14 | ENSG00000177606 | JUN | 23.36 | 3.0782 | 1.83E-54 | 2.765815877 | -3.0782442 | 1.83E-54 |
| 15 | ENSG00000100906 | NFKBIA | 16.377 | 2.9886 | 1.05E-51 | 2.063392619 | -2.98855149 | 1.05E-51 |
| 16 | ENSG00000115414 | FN1 | 1.2455 | -3.0524 | 2.12E-51 | 10.33206737 | 3.052359054 | 2.12E-51 |
| 17 | ENSG00000134531 | EMP1 | 18.056 | 2.9262 | 6.88E-50 | 2.375497652 | -2.92618263 | 6.88E-50 |
| 18 | ENSG00000163347 | CLDN1 | 8.9853 | 2.9265 | 2.17E-49 | 1.181871621 | -2.92645217 | 2.17E-49 |
| 19 | ENSG00000112096 | SOD2 | 15.321 | 2.8064 | 2.95E-46 | 2.190059108 | -2.80643342 | 2.95E-46 |
| 20 | ENSG00000135821 | GLUL | 7.5819 | 2.7988 | 3.06E-46 | 1.089541172 | -2.79881145 | 3.06E-46 |
| 21 | ENSG00000143384 | MCL1 | 7.0876 | 2.7347 | 1.89E-44 | 1.064746244 | -2.73474929 | 1.89E-44 |
| 22 | ENSG00000171223 | JUNB | 8.7379 | 2.7341 | 2.44E-44 | 1.313263801 | -2.73410281 | 2.44E-44 |
| 23 | ENSG00000106211 | HSPB1 | 15.233 | 2.6536 | 4.16E-42 | 2.420810457 | -2.65358491 | 4.16E-42 |
| 24 | ENSG00000133639 | BTG1 | 16.386 | 2.6299 | 1.98E-41 | 2.647195028 | -2.62988233 | 1.98E-41 |
| 25 | ENSG00000184292 | TACSTD2 | 17.565 | 2.6391 | 2.03E-41 | 2.819712692 | -2.63906029 | 2.03E-41 |
| 26 | ENSG00000120694 | HSPH1 | 12.065 | 2.5933 | 2.01E-40 | 1.999177046 | -2.5932812 | 2.01E-40 |
| 27 | ENSG00000150991 | UBC | 37.511 | 2.5866 | 2.30E-40 | 6.2447925 | -2.58656606 | 2.30E-40 |
| 28 | ENSG00000139329 | LUM | 28.801 | 2.5919 | 2.38E-40 | 4.777046428 | -2.59193879 | 2.38E-40 |
| 29 | ENSG00000141682 | PMAIP1 | 7.7543 | 2.5954 | 3.41E-40 | 1.283085174 | -2.59535409 | 3.41E-40 |
| 30 | ENSG00000163659 | TIPARP | 5.9478 | 2.5693 | 8.68E-40 | 1.002115871 | -2.56927488 | 8.68E-40 |
| 31 | ENSG00000181649 | PHLDA2 | 9.8583 | 2.4786 | 3.20E-37 | 1.768754701 | -2.47858311 | 3.20E-37 |
| 32 | ENSG00000204592 | HLA-E | 6.2706 | 2.4555 | 8.21E-37 | 1.143168802 | -2.45552349 | 8.21E-37 |
| 33 | ENSG00000197956 | S100A6 | 39.547 | 2.4345 | 3.00E-36 | 7.315789817 | -2.43446848 | 3.00E-36 |
| 34 | ENSG00000142089 | IFITM3 | 14.984 | 2.3502 | 4.74E-34 | 2.938453179 | -2.3502323 | 4.74E-34 |
| 35 | ENSG00000146278 | PNRC1 | 11.195 | 2.341 | 7.89E-34 | 2.209470337 | -2.34104103 | 7.89E-34 |
| 36 | ENSG00000269893 | SNHG8 | 5.1735 | 2.2493 | 1.80E-31 | 1.088075609 | -2.24933762 | 1.80E-31 |
| 37 | ENSG00000137801 | THBS1 | 8.0735 | 2.2416 | 3.48E-31 | 1.707171159 | -2.24155957 | 3.48E-31 |
| 38 | ENSG00000171345 | KRT19 | 1.6428 | -2.22 | 9.09E-30 | 7.653916065 | 2.219996792 | 9.09E-30 |
| 39 | ENSG00000170315 | UBB | 26.052 | 2.1622 | 2.44E-29 | 5.820257873 | -2.16221986 | 2.44E-29 |
| 40 | ENSG00000136240 | KDELR2 | 1.1447 | -2.1226 | 1.19E-27 | 4.985006782 | 2.122566669 | 1.19E-27 |
| 41 | ENSG00000166681 | BEX3 | 1.0538 | -2.0863 | 7.99E-27 | 4.475080697 | 2.086346383 | 7.99E-27 |
| 42 | ENSG00000135046 | ANXA1 | 64.853 | 2.0486 | 1.28E-26 | 15.67631646 | -2.04859223 | 1.28E-26 |
| 43 | ENSG00000111011 | RSRC2 | 5.7728 | 2.0443 | 1.55E-26 | 1.399522634 | -2.04431305 | 1.55E-26 |

| | | | | | | | | |
|---|---|---|---|---|---|---|---|---|
| 44 | ENSG00000205542 | TMSB4X | 16.908 | -2.0674 | 2.19E-26 | 70.86542914 | 2.067412714 | 2.19E-26 |
| 45 | ENSG00000080824 | HSP90AA1 | 58.2 | 2.0209 | 5.44E-26 | 14.34085985 | -2.02089398 | 5.44E-26 |
| 46 | ENSG00000100234 | TIMP3 | 4.3915 | 2.0277 | 5.70E-26 | 1.077009115 | -2.02765763 | 5.70E-26 |
| 47 | ENSG00000096696 | DSP | 3.6992 | -2.0491 | 7.58E-26 | 15.30944724 | 2.049114766 | 7.58E-26 |
| 48 | ENSG00000197632 | SERPINB2 | 5.3915 | 2.0291 | 1.99E-25 | 1.320980438 | -2.02906179 | 1.99E-25 |
| 49 | ENSG00000115963 | RND3 | 10.367 | 1.9832 | 4.48E-25 | 2.622041187 | -1.98318466 | 4.48E-25 |
| 50 | ENSG00000062582 | MRPS24 | 5.8344 | 1.9792 | 6.03E-25 | 1.479799577 | -1.97916357 | 6.03E-25 |
| 51 | ENSG00000142871 | CCN1 | 7.1055 | 1.9664 | 1.11E-24 | 1.818254829 | -1.96637102 | 1.11E-24 |
| 52 | ENSG00000107372 | ZFAND5 | 7.5724 | 1.9383 | 4.04E-24 | 1.975787862 | -1.93830689 | 4.04E-24 |
| 53 | ENSG00000115541 | HSPE1 | 11.226 | 1.9411 | 4.05E-24 | 2.923289091 | -1.94113676 | 4.05E-24 |
| 54 | ENSG00000100292 | HMOX1 | 13.679 | 1.9222 | 1.09E-23 | 3.609232246 | -1.92217957 | 1.09E-23 |
| 55 | ENSG00000198856 | OSTC | 1.4457 | -1.9389 | 1.46E-23 | 5.542967432 | 1.938872546 | 1.46E-23 |
| 56 | ENSG00000120708 | TGFBI | 5.4757 | 1.9177 | 1.46E-23 | 1.449262037 | -1.91770429 | 1.46E-23 |
| 57 | ENSG00000128422 | KRT17 | 9.3109 | -1.9395 | 2.27E-23 | 35.71316046 | 1.939459561 | 2.27E-23 |
| 58 | ENSG00000138061 | CYP1B1 | 1.5758 | -1.9304 | 3.07E-23 | 6.006414244 | 1.930415728 | 3.07E-23 |
| 59 | ENSG00000206503 | HLA-A | 4.7465 | 1.8962 | 3.60E-23 | 1.275129262 | -1.89619669 | 3.60E-23 |
| 60 | ENSG00000109971 | HSPA8 | 15.34 | 1.8645 | 1.69E-22 | 4.212655248 | -1.86453376 | 1.69E-22 |
| 61 | ENSG00000086061 | DNAJA1 | 17.431 | 1.8513 | 3.27E-22 | 4.831092791 | -1.85125686 | 3.27E-22 |
| 62 | ENSG00000051108 | HERPUD1 | 3.9052 | 1.8312 | 9.29E-22 | 1.097467174 | -1.83119005 | 9.29E-22 |
| 63 | ENSG00000139289 | PHLDA1 | 5.0137 | 1.8257 | 1.40E-21 | 1.414357717 | -1.82571525 | 1.40E-21 |
| 64 | ENSG00000196230 | TUBB | 1.2079 | -1.8177 | 5.50E-21 | 4.258237239 | 1.817700789 | 5.50E-21 |
| 65 | ENSG00000176788 | BASP1 | 1.0135 | -1.8132 | 7.36E-21 | 3.561736051 | 1.81316782 | 7.36E-21 |
| 66 | ENSG00000230937 | MIR205HG | 4.6742 | 1.7924 | 8.33E-21 | 1.349424317 | -1.79235727 | 8.33E-21 |
| 67 | ENSG00000221869 | CEBPD | 8.6689 | 1.7649 | 2.28E-20 | 2.550737076 | -1.76492119 | 2.28E-20 |
| 68 | ENSG00000131981 | LGALS3 | 12.001 | 1.7618 | 2.73E-20 | 3.538765599 | -1.76178538 | 2.73E-20 |
| 69 | ENSG00000167460 | TPM4 | 2.6001 | -1.7723 | 4.35E-20 | 8.882087502 | 1.772332489 | 4.35E-20 |
| 70 | ENSG00000042753 | AP2S1 | 1.0188 | -1.7696 | 4.95E-20 | 3.473563015 | 1.769560341 | 4.95E-20 |
| 71 | ENSG00000105438 | KDELR1 | 1.2996 | -1.7598 | 7.65E-20 | 4.401264193 | 1.75983273 | 7.65E-20 |
| 72 | ENSG00000150347 | ARID5B | 3.9786 | 1.7376 | 8.79E-20 | 1.192997933 | -1.73764367 | 8.79E-20 |
| 73 | ENSG00000125944 | HNRNPR | 1.2634 | -1.7476 | 1.34E-19 | 4.242774057 | 1.747632482 | 1.34E-19 |
| 74 | ENSG00000124762 | CDKN1A | 3.9532 | 1.7198 | 1.96E-19 | 1.20014629 | -1.71977804 | 1.96E-19 |
| 75 | ENSG00000156976 | EIF4A2 | 5.326 | 1.7137 | 2.45E-19 | 1.623753724 | -1.7136961 | 2.45E-19 |
| 76 | ENSG00000167996 | FTH1 | 165.09 | 1.7106 | 2.81E-19 | 50.43964275 | -1.71058266 | 2.81E-19 |
| 77 | ENSG00000175793 | SFN | 16.307 | 1.711 | 3.71E-19 | 4.981028826 | -1.71099794 | 3.71E-19 |
| 78 | ENSG00000175130 | MARCKSL1 | 1.0202 | -1.7204 | 5.13E-19 | 3.361731609 | 1.720370583 | 5.13E-19 |
| 79 | ENSG00000189143 | CLDN4 | 9.4767 | 1.6639 | 3.74E-18 | 2.990675065 | -1.66389938 | 3.74E-18 |
| 80 | ENSG00000116044 | NFE2L2 | 4.4359 | 1.6372 | 8.15E-18 | 1.426052309 | -1.63717015 | 8.15E-18 |
| 81 | ENSG00000165272 | AQP3 | 1.9123 | -1.647 | 1.96E-17 | 5.989096676 | 1.647006698 | 1.96E-17 |
| 82 | ENSG00000111057 | KRT18 | 1.8042 | -1.6313 | 3.74E-17 | 5.589297158 | 1.631307409 | 3.74E-17 |
| 83 | ENSG00000143546 | S100A8 | 2.6235 | -1.6637 | 5.97E-17 | 8.311714434 | 1.663662007 | 5.97E-17 |
| 84 | ENSG00000146457 | WTAP | 4.5002 | 1.5931 | 6.50E-17 | 1.491643717 | -1.59306163 | 6.50E-17 |
| 85 | ENSG00000116717 | GADD45A | 5.824 | 1.5785 | 1.26E-16 | 1.949976013 | -1.5785366 | 1.26E-16 |
| 86 | ENSG00000234745 | HLA-B | 3.21 | 1.5703 | 1.68E-16 | 1.080897343 | -1.57032995 | 1.68E-16 |
| 87 | ENSG00000116285 | ERRFI1 | 2.9716 | 1.5624 | 2.37E-16 | 1.006093827 | -1.56242612 | 2.37E-16 |
| 88 | ENSG00000096384 | HSP90AB1 | 45.259 | 1.558 | 2.60E-16 | 15.37082142 | -1.55801189 | 2.60E-16 |
| 89 | ENSG00000011422 | PLAUR | 5.5717 | 1.5619 | 2.96E-16 | 1.887196094 | -1.56186548 | 2.96E-16 |
| 90 | ENSG00000137440 | FGFBP1 | 3.6743 | 1.5653 | 3.18E-16 | 1.241570868 | -1.56528875 | 3.18E-16 |

| | | | | | | | | |
|---|---|---|---|---|---|---|---|---|
| 91 | ENSG00000213639 | PPP1CB | 1.3655 | -1.568 | 3.70E-16 | 4.048542137 | 1.56799015 | 3.70E-16 |
| 92 | ENSG00000172216 | CEBPB | 4.0803 | 1.5436 | 5.17E-16 | 1.39970209 | -1.54355013 | 5.17E-16 |
| 93 | ENSG00000144381 | HSPD1 | 12.978 | 1.5421 | 5.87E-16 | 4.456417205 | -1.54211367 | 5.87E-16 |
| 94 | ENSG00000105372 | RPS19 | 11.905 | -1.539 | 1.16E-15 | 34.59412857 | 1.538968303 | 1.16E-15 |
| 95 | ENSG00000179218 | CALR | 1.5368 | -1.5365 | 1.36E-15 | 4.45827159 | 1.536512196 | 1.36E-15 |
| 96 | ENSG00000128595 | CALU | 1.4709 | -1.5342 | 1.49E-15 | 4.260001896 | 1.534173541 | 1.49E-15 |
| 97 | ENSG00000204256 | BRD2 | 4.1167 | 1.5179 | 1.50E-15 | 1.437447807 | -1.51794554 | 1.50E-15 |
| 98 | ENSG00000204525 | HLA-C | 3.086 | 1.5182 | 1.51E-15 | 1.077368028 | -1.518191 | 1.51E-15 |
| 99 | ENSG00000102804 | TSC22D1 | 8.0009 | 1.511 | 2.04E-15 | 2.807210545 | -1.51101267 | 2.04E-15 |
| 100 | ENSG00000140988 | RPS2 | 19.317 | -1.5235 | 2.20E-15 | 55.53453683 | 1.523507652 | 2.20E-15 |
| 101 | ENSG00000198034 | RPS4X | 22.096 | -1.488 | 9.25E-15 | 61.9794235 | 1.488006704 | 9.25E-15 |
| 102 | ENSG00000115677 | HDLBP | 1.2986 | -1.4871 | 1.02E-14 | 3.640367975 | 1.487059667 | 1.02E-14 |
| 103 | ENSG00000143870 | PDIA6 | 1.3866 | -1.4523 | 4.11E-14 | 3.794371695 | 1.452328469 | 4.11E-14 |
| 104 | ENSG00000136527 | TRA2B | 3.0941 | 1.4206 | 8.16E-14 | 1.155790587 | -1.42062263 | 8.16E-14 |
| 105 | ENSG00000166794 | PPIB | 3.0388 | -1.4246 | 1.18E-13 | 8.157501348 | 1.424619888 | 1.18E-13 |
| 106 | ENSG00000137309 | HMGA1 | 1.2522 | -1.4255 | 1.21E-13 | 3.363645813 | 1.425517947 | 1.21E-13 |
| 107 | ENSG00000102265 | TIMP1 | 8.2759 | 1.4099 | 1.45E-13 | 3.114410429 | -1.40994269 | 1.45E-13 |
| 108 | ENSG00000108106 | UBE2S | 3.8213 | 1.3941 | 2.34E-13 | 1.453898 | -1.39412414 | 2.34E-13 |
| 109 | ENSG00000167658 | EEF2 | 3.9358 | -1.3976 | 3.30E-13 | 10.36930462 | 1.397595789 | 3.30E-13 |
| 110 | ENSG00000197746 | PSAP | 3.903 | 1.3623 | 7.96E-13 | 1.518143483 | -1.36227215 | 7.96E-13 |
| 111 | ENSG00000198840 | MT-ND3 | 5.2531 | -1.3812 | 9.01E-13 | 13.68321103 | 1.381172326 | 9.01E-13 |
| 112 | ENSG00000188643 | S100A16 | 1.6341 | -1.3516 | 2.19E-12 | 4.170303478 | 1.351621687 | 2.19E-12 |
| 113 | ENSG00000198938 | MT-CO3 | 14.188 | -1.3418 | 3.63E-12 | 35.96203693 | 1.341759901 | 3.63E-12 |
| 114 | ENSG00000163931 | TKT | 6.1761 | 1.3117 | 6.04E-12 | 2.488046884 | -1.31167642 | 6.04E-12 |
| 115 | ENSG00000181467 | RAP2B | 3.6381 | 1.3079 | 7.24E-12 | 1.46948082 | -1.30786093 | 7.24E-12 |
| 116 | ENSG00000142534 | RPS11 | 10.313 | -1.2882 | 1.88E-11 | 25.18686117 | 1.288202245 | 1.88E-11 |
| 117 | ENSG00000175061 | SNHG29 | 6.535 | -1.2844 | 2.16E-11 | 15.91834367 | 1.28443771 | 2.16E-11 |
| 118 | ENSG00000114942 | EEF1B2 | 5.9832 | -1.2789 | 2.62E-11 | 14.5187014 | 1.27892346 | 2.62E-11 |
| 119 | ENSG00000184009 | ACTG1 | 18.254 | -1.2786 | 2.65E-11 | 44.28694142 | 1.27864916 | 2.65E-11 |
| 120 | ENSG00000166441 | RPL27A | 8.6462 | -1.278 | 2.71E-11 | 20.96783521 | 1.27803107 | 2.71E-11 |
| 121 | ENSG00000130066 | SAT1 | 22.276 | 1.2644 | 3.05E-11 | 9.272944097 | -1.26440724 | 3.05E-11 |
| 122 | ENSG00000060138 | YBX3 | 9.6578 | 1.2622 | 3.15E-11 | 4.026439059 | -1.26217735 | 3.15E-11 |
| 123 | ENSG00000143320 | CRABP2 | 1.5331 | -1.2705 | 3.72E-11 | 3.698631569 | 1.27054529 | 3.72E-11 |
| 124 | ENSG00000135486 | HNRNPA1 | 4.9326 | -1.2574 | 5.57E-11 | 11.79188744 | 1.257385514 | 5.57E-11 |
| 125 | ENSG00000198830 | HMGN2 | 1.3729 | -1.2553 | 6.46E-11 | 3.277416891 | 1.255314977 | 6.46E-11 |
| 126 | ENSG00000096746 | HNRNPH3 | 4.1093 | 1.2322 | 9.28E-11 | 1.749193926 | -1.23217331 | 9.28E-11 |
| 127 | ENSG00000168028 | RPSA | 7.5925 | -1.239 | 1.05E-10 | 17.92084066 | 1.238997175 | 1.05E-10 |
| 128 | ENSG00000167553 | TUBA1C | 3.3409 | 1.2204 | 1.46E-10 | 1.433828764 | -1.22036444 | 1.46E-10 |
| 129 | ENSG00000204387 | SNHG32 | 1.4254 | -1.2287 | 1.55E-10 | 3.340705271 | 1.228744913 | 1.55E-10 |
| 130 | ENSG00000117906 | RCN2 | 1.1586 | -1.2249 | 1.82E-10 | 2.70821029 | 1.224923939 | 1.82E-10 |
| 131 | ENSG00000265972 | TXNIP | 4.229 | 1.2148 | 1.85E-10 | 1.822023418 | -1.21476738 | 1.85E-10 |
| 132 | ENSG00000004779 | NDUFAB1 | 1.1225 | -1.2209 | 2.02E-10 | 2.616537849 | 1.22090461 | 2.02E-10 |
| 133 | ENSG00000169100 | SLC25A6 | 2.1343 | -1.2122 | 2.67E-10 | 4.945137495 | 1.212221921 | 2.67E-10 |
| 134 | ENSG00000175183 | CSRP2 | 6.0601 | 1.2064 | 2.72E-10 | 2.626049052 | -1.20642531 | 2.72E-10 |
| 135 | ENSG00000198899 | MT-ATP6 | 12.466 | -1.2104 | 4.10E-10 | 28.84693947 | 1.21036743 | 4.10E-10 |
| 136 | ENSG00000163453 | IGFBP7 | 3.025 | 1.1941 | 4.62E-10 | 1.322057178 | -1.19411845 | 4.62E-10 |
| 137 | ENSG00000152518 | ZFP36L2 | 3.4109 | 1.18 | 5.94E-10 | 1.505342241 | -1.18003527 | 5.94E-10 |

| | | | | | | | | |
|---|---|---|---|---|---|---|---|---|
| 138 | ENSG00000115091 | ACTR3 | 1.454 | -1.1802 | 7.94E-10 | 3.295033552 | 1.18022874 | 7.94E-10 |
| 139 | ENSG00000138674 | SEC31A | 1.0548 | -1.1751 | 9.38E-10 | 2.381868363 | 1.175143455 | 9.38E-10 |
| 140 | ENSG00000198888 | MT-ND1 | 5.7878 | -1.177 | 1.05E-09 | 13.08708593 | 1.177045421 | 1.05E-09 |
| 141 | ENSG00000114125 | RNF7 | 1.509 | -1.1706 | 1.07E-09 | 3.396815385 | 1.170552069 | 1.07E-09 |
| 142 | ENSG00000204628 | RACK1 | 9.6661 | -1.1672 | 1.17E-09 | 21.70752563 | 1.167184992 | 1.17E-09 |
| 143 | ENSG00000143256 | PFDN2 | 2.8633 | 1.1553 | 1.30E-09 | 1.285507839 | -1.15533822 | 1.30E-09 |
| 144 | ENSG00000134755 | DSC2 | 2.5972 | -1.1641 | 1.50E-09 | 5.820287783 | 1.164149677 | 1.50E-09 |
| 145 | ENSG00000108654 | DDX5 | 13.282 | 1.1499 | 1.51E-09 | 5.98565709 | -1.14990764 | 1.51E-09 |
| 146 | ENSG00000126524 | SBDS | 5.0384 | 1.1491 | 1.56E-09 | 2.271861434 | -1.14908575 | 1.56E-09 |
| 147 | ENSG00000172270 | BSG | 1.2878 | -1.1569 | 1.70E-09 | 2.871575665 | 1.156872901 | 1.70E-09 |
| 148 | ENSG00000121774 | KHDRBS1 | 1.3882 | -1.1558 | 1.74E-09 | 3.093084997 | 1.15576537 | 1.74E-09 |
| 149 | ENSG00000141543 | EIF4A3 | 3.529 | 1.1435 | 1.92E-09 | 1.597403505 | -1.1435095 | 1.92E-09 |
| 150 | ENSG00000122406 | RPL5 | 15.783 | -1.1436 | 2.51E-09 | 34.86836825 | 1.143574792 | 2.51E-09 |
| 151 | ENSG00000136156 | ITM2B | 13.113 | 1.1317 | 2.74E-09 | 5.984580351 | -1.1316987 | 2.74E-09 |
| 152 | ENSG00000120438 | TCP1 | 3.419 | 1.1283 | 3.17E-09 | 1.563964749 | -1.12834293 | 3.17E-09 |
| 153 | ENSG00000182774 | RPS17 | 6.8133 | -1.1332 | 3.52E-09 | 14.94482123 | 1.133223202 | 3.52E-09 |
| 154 | ENSG00000087460 | GNAS | 5.2028 | -1.1326 | 3.59E-09 | 11.40704264 | 1.132562861 | 3.59E-09 |
| 155 | ENSG00000154518 | ATP5MC3 | 2.4794 | -1.1292 | 4.05E-09 | 5.423359571 | 1.129206949 | 4.05E-09 |
| 156 | ENSG00000168653 | NDUFS5 | 1.5032 | -1.1275 | 4.31E-09 | 3.284236244 | 1.127497448 | 4.31E-09 |
| 157 | ENSG00000108107 | RPL28 | 11.826 | -1.1262 | 4.39E-09 | 25.81427155 | 1.126152952 | 4.39E-09 |
| 158 | ENSG00000074695 | LMAN1 | 1.7998 | -1.1227 | 5.04E-09 | 3.919303436 | 1.122726202 | 5.04E-09 |
| 159 | ENSG00000136603 | SKIL | 1.1596 | -1.1223 | 5.28E-09 | 2.524446676 | 1.122287012 | 5.28E-09 |
| 160 | ENSG00000140612 | SEC11A | 1.8525 | -1.1187 | 5.66E-09 | 4.022700379 | 1.118670706 | 5.66E-09 |
| 161 | ENSG00000131469 | RPL27 | 5.1052 | -1.1177 | 5.75E-09 | 11.07815841 | 1.117662459 | 5.75E-09 |
| 162 | ENSG00000100219 | XBP1 | 2.1713 | 1.1061 | 6.57E-09 | 1.00863613 | -1.10611392 | 6.57E-09 |
| 163 | ENSG00000260032 | NORAD | 1.6278 | -1.1121 | 7.07E-09 | 3.518546816 | 1.11207231 | 7.07E-09 |
| 164 | ENSG00000083845 | RPS5 | 11.455 | -1.1108 | 7.12E-09 | 24.74040292 | 1.110832512 | 7.12E-09 |
| 165 | ENSG00000164687 | FABP5 | 16.846 | 1.1043 | 7.85E-09 | 7.835197196 | -1.10432343 | 7.85E-09 |
| 166 | ENSG00000108256 | NUFIP2 | 2.746 | 1.0979 | 8.45E-09 | 1.282905717 | -1.09790877 | 8.45E-09 |
| 167 | ENSG00000198712 | MT-CO2 | 24.903 | -1.111 | 8.94E-09 | 53.78929095 | 1.110996756 | 8.94E-09 |
| 168 | ENSG00000100201 | DDX17 | 1.754 | -1.1017 | 9.71E-09 | 3.76407343 | 1.101652901 | 9.71E-09 |
| 169 | ENSG00000178053 | MLF1 | 2.4514 | 1.087 | 1.20E-08 | 1.153936201 | -1.08704293 | 1.20E-08 |
| 170 | ENSG00000197111 | PCBP2 | 2.3918 | -1.0939 | 1.22E-08 | 5.105182922 | 1.093860538 | 1.22E-08 |
| 171 | ENSG00000113811 | SELENOK | 3.6708 | 1.0815 | 1.39E-08 | 1.734538299 | -1.08154052 | 1.39E-08 |
| 172 | ENSG00000124766 | SOX4 | 1.9201 | -1.0917 | 1.43E-08 | 4.092180014 | 1.091656281 | 1.43E-08 |
| 173 | ENSG00000265681 | RPL17 | 16.658 | -1.0869 | 1.49E-08 | 35.38517351 | 1.086932324 | 1.49E-08 |
| 174 | ENSG00000254772 | EEF1G | 5.1424 | -1.087 | 1.50E-08 | 10.92448369 | 1.08704189 | 1.50E-08 |
| 175 | ENSG00000186468 | RPS23 | 14.713 | -1.0862 | 1.53E-08 | 31.23610574 | 1.086157612 | 1.53E-08 |
| 176 | ENSG00000149273 | RPS3 | 17.991 | -1.0807 | 1.81E-08 | 38.05342478 | 1.080721082 | 1.81E-08 |
| 177 | ENSG00000203875 | SNHG5 | 5.8065 | 1.0731 | 1.81E-08 | 2.75971435 | -1.07313701 | 1.81E-08 |
| 178 | ENSG00000100300 | TSPO | 3.556 | 1.0707 | 1.95E-08 | 1.692964174 | -1.07070062 | 1.95E-08 |
| 179 | ENSG00000249915 | PDCD6 | 1.1101 | -1.0789 | 1.97E-08 | 2.344990021 | 1.078926816 | 1.97E-08 |
| 180 | ENSG00000159335 | PTMS | 1.6832 | -1.0762 | 2.14E-08 | 3.549203995 | 1.07623704 | 2.14E-08 |
| 181 | ENSG00000164292 | RHOBTB3 | 1.2227 | -1.0739 | 2.43E-08 | 2.573827165 | 1.073864212 | 2.43E-08 |
| 182 | ENSG00000112773 | TENT5A | 2.2418 | 1.0641 | 2.51E-08 | 1.072193695 | -1.06408101 | 2.51E-08 |
| 183 | ENSG00000142168 | SOD1 | 5.2541 | 1.0617 | 2.54E-08 | 2.517059043 | -1.0616975 | 2.54E-08 |
| 184 | ENSG00000067167 | TRAM1 | 1.0753 | -1.0706 | 2.55E-08 | 2.258521822 | 1.070581565 | 2.55E-08 |

| | | | | | | | | |
|---|---|---|---|---|---|---|---|---|
| 185 | ENSG00000142173 | COL6A2 | 4.8734 | 1.0618 | 2.70E-08 | 2.334581535 | -1.06175851 | 2.70E-08 |
| 186 | ENSG00000089157 | RPLP0 | 11.977 | -1.066 | 2.84E-08 | 25.07583732 | 1.065975147 | 2.84E-08 |
| 187 | ENSG00000231500 | RPS18 | 25.833 | -1.066 | 2.84E-08 | 54.08315122 | 1.065965728 | 2.84E-08 |
| 188 | ENSG00000169230 | PRELID1 | 1.6882 | -1.0642 | 3.06E-08 | 3.530002133 | 1.06416848 | 3.06E-08 |
| 189 | ENSG00000148303 | RPL7A | 17.751 | -1.0632 | 3.08E-08 | 37.09019119 | 1.063166223 | 3.08E-08 |
| 190 | ENSG00000134419 | RPS15A | 10.956 | -1.0624 | 3.16E-08 | 22.88000569 | 1.062395695 | 3.16E-08 |
| 191 | ENSG00000076043 | REXO2 | 1.3719 | -1.0631 | 3.19E-08 | 2.866640607 | 1.063120745 | 3.19E-08 |
| 192 | ENSG00000205581 | HMGN1 | 1.7538 | -1.0619 | 3.27E-08 | 3.661424223 | 1.061920209 | 3.27E-08 |
| 193 | ENSG00000125266 | EFNB2 | 1.0062 | -1.0636 | 3.30E-08 | 2.103022631 | 1.063591321 | 3.30E-08 |
| 194 | ENSG00000101782 | RIOK3 | 2.4751 | 1.0537 | 3.32E-08 | 1.192280107 | -1.05374878 | 3.32E-08 |
| 195 | ENSG00000108424 | KPNB1 | 1.0452 | -1.0619 | 3.33E-08 | 2.182163016 | 1.061907552 | 3.33E-08 |
| 196 | ENSG00000145425 | RPS3A | 26.904 | -1.0596 | 3.42E-08 | 56.07820076 | 1.05963101 | 3.42E-08 |
| 197 | ENSG00000203930 | LINC00632 | 1.2055 | -1.0618 | 3.45E-08 | 2.516550583 | 1.061786132 | 3.45E-08 |
| 198 | ENSG00000074800 | ENO1 | 12.024 | 1.0465 | 4.07E-08 | 5.821364523 | -1.04654109 | 4.07E-08 |
| 199 | ENSG00000128272 | ATF4 | 3.5632 | 1.0449 | 4.29E-08 | 1.72700112 | -1.04488113 | 4.29E-08 |
| 200 | ENSG00000122786 | CALD1 | 3.3089 | -1.0532 | 4.32E-08 | 6.866310716 | 1.053199358 | 4.32E-08 |
| 201 | ENSG00000118418 | HMGN3 | 1.55 | -1.0535 | 4.35E-08 | 3.217298911 | 1.053525977 | 4.35E-08 |
| 202 | ENSG00000164587 | RPS14 | 18.993 | -1.0491 | 4.70E-08 | 39.30019989 | 1.049075209 | 4.70E-08 |
| 203 | ENSG00000162704 | ARPC5 | 1.5735 | -1.048 | 4.97E-08 | 3.253549156 | 1.0479991 | 4.97E-08 |
| 204 | ENSG00000187109 | NAP1L1 | 3.9155 | -1.0424 | 5.79E-08 | 8.064841896 | 1.042446178 | 5.79E-08 |
| 205 | ENSG00000188612 | SUMO2 | 4.593 | -1.0389 | 6.40E-08 | 9.436847842 | 1.038863959 | 6.40E-08 |
| 206 | ENSG00000140941 | MAP1LC3B | 5.0137 | 1.0299 | 6.71E-08 | 2.455385773 | -1.02992552 | 6.71E-08 |
| 207 | ENSG00000085063 | CD59 | 2.8995 | 1.0299 | 6.84E-08 | 1.419980692 | -1.0298976 | 6.84E-08 |
| 208 | ENSG00000106803 | SEC61B | 2.8771 | -1.035 | 7.19E-08 | 5.895659578 | 1.035018835 | 7.19E-08 |
| 209 | ENSG00000130522 | JUND | 8.3403 | 1.0257 | 7.61E-08 | 4.096516883 | -1.02569723 | 7.61E-08 |
| 210 | ENSG00000154582 | ELOC | 3.4288 | 1.0234 | 8.19E-08 | 1.686772919 | -1.02342425 | 8.19E-08 |
| 211 | ENSG00000175756 | AURKAIP1 | 1.1817 | -1.0293 | 8.71E-08 | 2.412046991 | 1.02932464 | 8.71E-08 |
| 212 | ENSG00000105404 | RABAC1 | 1.9496 | -1.0265 | 9.37E-08 | 3.971555232 | 1.026528054 | 9.37E-08 |
| 213 | ENSG00000166913 | YWHAB | 2.3646 | -1.0259 | 9.44E-08 | 4.814851964 | 1.025889853 | 9.44E-08 |
| 214 | ENSG00000125691 | RPL23 | 9.7718 | -1.0245 | 9.71E-08 | 19.87874268 | 1.024528521 | 9.71E-08 |
| 215 | ENSG00000198604 | BAZ1A | 2.1493 | 1.0184 | 9.76E-08 | 1.061037473 | -1.0183833 | 9.76E-08 |
| 216 | ENSG00000166012 | TAF1D | 2.0933 | 1.0139 | 1.10E-07 | 1.036571549 | -1.01393816 | 1.10E-07 |
| 217 | ENSG00000188229 | TUBB4B | 4.8894 | 1.012 | 1.15E-07 | 2.424369681 | -1.01203713 | 1.15E-07 |
| 218 | ENSG00000100316 | RPL3 | 19.298 | -1.0172 | 1.20E-07 | 39.05924943 | 1.017199634 | 1.20E-07 |
| 219 | ENSG00000185650 | ZFP36L1 | 4.5474 | 1.01 | 1.25E-07 | 2.257953543 | -1.01001692 | 1.25E-07 |
| 220 | ENSG00000135390 | ATP5MC2 | 1.8178 | -1.0151 | 1.30E-07 | 3.673986189 | 1.015131671 | 1.30E-07 |
| 221 | ENSG00000143742 | SRP9 | 1.3585 | -1.0153 | 1.30E-07 | 2.746105554 | 1.015316972 | 1.30E-07 |
| 222 | ENSG00000175768 | TOMM5 | 1.3814 | -1.011 | 1.48E-07 | 2.783970908 | 1.010981491 | 1.48E-07 |
| 223 | ENSG00000089737 | DDX24 | 2.8721 | 0.9948 | 1.90E-07 | 1.441216396 | -0.99478924 | 1.90E-07 |
| 224 | ENSG00000175390 | EIF3F | 1.2986 | -1.0007 | 1.98E-07 | 2.598472546 | 1.000679247 | 1.98E-07 |
| 225 | ENSG00000198900 | TOP1 | 3.2448 | 0.9794 | 2.95E-07 | 1.645737164 | -0.97939231 | 2.95E-07 |
| 226 | ENSG00000174695 | TMEM167A | 1.0679 | -0.9864 | 2.98E-07 | 2.115734144 | 0.9863722 | 2.98E-07 |
| 227 | ENSG00000241343 | RPL36A | 4.7609 | -0.9842 | 3.09E-07 | 9.417825437 | 0.98416576 | 3.09E-07 |
| 228 | ENSG00000171222 | SCAND1 | 1.2071 | -0.9843 | 3.17E-07 | 2.388029708 | 0.98426021 | 3.17E-07 |
| 229 | ENSG00000110321 | EIF4G2 | 3.9594 | -0.9781 | 3.68E-07 | 7.799365684 | 0.978067029 | 3.68E-07 |
| 230 | ENSG00000117592 | PRDX6 | 2.6817 | -0.9778 | 3.73E-07 | 5.281648633 | 0.97781166 | 3.73E-07 |
| 231 | ENSG00000173726 | TOMM20 | 2.9693 | -0.9747 | 4.04E-07 | 5.83545187 | 0.974713612 | 4.04E-07 |

| | | | | | | | | |
|---|---|---|---|---|---|---|---|---|
| 232 | ENSG00000245532 | NEAT1 | 14.87 | -0.9735 | 4.35E-07 | 29.19903343 | 0.973491621 | 4.35E-07 |
| 233 | ENSG00000138326 | RPS24 | 17.186 | -0.9634 | 5.48E-07 | 33.51140675 | 0.963442068 | 5.48E-07 |
| 234 | ENSG00000198763 | MT-ND2 | 7.6903 | -0.9687 | 5.73E-07 | 15.05082029 | 0.968735935 | 5.73E-07 |
| 235 | ENSG00000184840 | TMED9 | 1.0618 | -0.9589 | 6.41E-07 | 2.063841261 | 0.958854938 | 6.41E-07 |
| 236 | ENSG00000174748 | RPL15 | 21.075 | -0.9569 | 6.56E-07 | 40.90810161 | 0.956875941 | 6.56E-07 |
| 237 | ENSG00000142937 | RPS8 | 21.598 | -0.9554 | 6.85E-07 | 41.87976967 | 0.955367768 | 6.85E-07 |
| 238 | ENSG00000063177 | RPL18 | 19.841 | -0.9541 | 7.08E-07 | 38.43982487 | 0.954113344 | 7.08E-07 |
| 239 | ENSG00000134333 | LDHA | 16.956 | 0.9419 | 8.35E-07 | 8.826246573 | -0.94192457 | 8.35E-07 |
| 240 | ENSG00000142541 | RPL13A | 14.861 | -0.9484 | 8.36E-07 | 28.67612664 | 0.948358668 | 8.36E-07 |
| 241 | ENSG00000176340 | COX8A | 1.066 | -0.9475 | 8.78E-07 | 2.055735802 | 0.947490162 | 8.78E-07 |
| 242 | ENSG00000171863 | RPS7 | 17.662 | -0.9439 | 9.37E-07 | 33.97566112 | 0.943889876 | 9.37E-07 |
| 243 | ENSG00000265241 | RBM8A | 3.576 | 0.9373 | 9.50E-07 | 1.867366133 | -0.9373226 | 9.50E-07 |
| 244 | ENSG00000119801 | YPEL5 | 2.0515 | 0.9382 | 9.56E-07 | 1.070608494 | -0.93819524 | 9.56E-07 |
| 245 | ENSG00000114850 | SSR3 | 1.5881 | -0.9421 | 1.01E-06 | 3.051301505 | 0.942100185 | 1.01E-06 |
| 246 | ENSG00000119335 | SET | 3.4761 | -0.9391 | 1.08E-06 | 6.664631345 | 0.939058159 | 1.08E-06 |
| 247 | ENSG00000136938 | ANP32B | 1.3477 | -0.9369 | 1.16E-06 | 2.580167967 | 0.936914203 | 1.16E-06 |
| 248 | ENSG00000111786 | SRSF9 | 2.2504 | -0.9322 | 1.30E-06 | 4.294308026 | 0.932218097 | 1.30E-06 |
| 249 | ENSG00000117410 | ATP6V0B | 1.7392 | -0.9304 | 1.37E-06 | 3.314654147 | 0.930422423 | 1.37E-06 |
| 250 | ENSG00000213719 | CLIC1 | 2.8277 | -0.9301 | 1.38E-06 | 5.388215976 | 0.930148683 | 1.38E-06 |
| 251 | ENSG00000189334 | S100A14 | 1.3264 | -0.9309 | 1.52E-06 | 2.528783545 | 0.930932835 | 1.52E-06 |
| 252 | ENSG00000002586 | CD99 | 4.4381 | 0.9186 | 1.59E-06 | 2.347801508 | -0.91861137 | 1.59E-06 |
| 253 | ENSG00000123562 | MORF4L2 | 3.5213 | -0.9243 | 1.60E-06 | 6.682636829 | 0.92431863 | 1.60E-06 |
| 254 | ENSG00000117450 | PRDX1 | 11.715 | 0.9161 | 1.68E-06 | 6.208482437 | -0.91609074 | 1.68E-06 |
| 255 | ENSG00000161016 | RPL8 | 16.219 | -0.921 | 1.73E-06 | 30.70862281 | 0.920967509 | 1.73E-06 |
| 256 | ENSG00000164713 | BRI3 | 2.0685 | -0.9178 | 1.91E-06 | 3.908057486 | 0.917846255 | 1.91E-06 |
| 257 | ENSG00000151366 | NDUFC2 | 1.5242 | -0.9152 | 2.05E-06 | 2.874506791 | 0.915238079 | 2.05E-06 |
| 258 | ENSG00000138071 | ACTR2 | 1.326 | -0.912 | 2.25E-06 | 2.49522515 | 0.912039727 | 2.25E-06 |
| 259 | ENSG00000163682 | RPL9 | 19.285 | -0.9106 | 2.27E-06 | 36.25386336 | 0.910637359 | 2.27E-06 |
| 260 | ENSG00000167526 | RPL13 | 33.231 | -0.9094 | 2.34E-06 | 62.41613127 | 0.909391139 | 2.34E-06 |
| 261 | ENSG00000122705 | CLTA | 2.8545 | -0.9079 | 2.46E-06 | 5.355913779 | 0.907888168 | 2.46E-06 |
| 262 | ENSG00000105640 | RPL18A | 16.143 | -0.9073 | 2.48E-06 | 30.27565372 | 0.907292591 | 2.48E-06 |
| 263 | ENSG00000135535 | CD164 | 1.3673 | -0.9047 | 2.71E-06 | 2.559859456 | 0.904679108 | 2.71E-06 |
| 264 | ENSG00000167552 | TUBA1A | 1.4322 | -0.9067 | 2.71E-06 | 2.685090291 | 0.906732256 | 2.71E-06 |
| 265 | ENSG00000124466 | LYPD3 | 2.2194 | 0.9027 | 2.83E-06 | 1.187135682 | -0.90269786 | 2.83E-06 |
| 266 | ENSG00000130402 | ACTN4 | 1.276 | -0.9018 | 2.99E-06 | 2.384141481 | 0.901805673 | 2.99E-06 |
| 267 | ENSG00000105669 | COPE | 1.4694 | -0.8995 | 3.09E-06 | 2.741200405 | 0.899536028 | 3.09E-06 |
| 268 | ENSG00000147676 | MAL2 | 2.4831 | 0.8969 | 3.13E-06 | 1.333482585 | -0.89693654 | 3.13E-06 |
| 269 | ENSG00000170889 | RPS9 | 16.022 | -0.8957 | 3.35E-06 | 29.81068152 | 0.895735556 | 3.35E-06 |
| 270 | ENSG00000101182 | PSMA7 | 3.4008 | -0.8951 | 3.43E-06 | 6.32474044 | 0.895110836 | 3.43E-06 |
| 271 | ENSG00000171858 | RPS21 | 6.5982 | -0.8942 | 3.50E-06 | 12.2634098 | 0.894216147 | 3.50E-06 |
| 272 | ENSG00000147403 | RPL10 | 37.212 | -0.893 | 3.59E-06 | 69.10472007 | 0.893022492 | 3.59E-06 |
| 273 | ENSG00000163466 | ARPC2 | 3.3759 | -0.891 | 3.83E-06 | 6.260674414 | 0.891042993 | 3.83E-06 |
| 274 | ENSG00000122545 | SEPTIN7 | 2.3645 | -0.8898 | 3.96E-06 | 4.381224866 | 0.889772335 | 3.96E-06 |
| 275 | ENSG00000104067 | TJP1 | 1.3211 | -0.8892 | 4.11E-06 | 2.446921401 | 0.8892061 | 4.11E-06 |
| 276 | ENSG00000189159 | JPT1 | 1.4085 | -0.8869 | 4.43E-06 | 2.604514253 | 0.886886242 | 4.43E-06 |
| 277 | ENSG00000117632 | STMN1 | 1.2731 | -0.8869 | 4.47E-06 | 2.354261948 | 0.886883748 | 4.47E-06 |
| 278 | ENSG00000233927 | RPS28 | 11.2 | -0.883 | 4.66E-06 | 20.65632837 | 0.883024203 | 4.66E-06 |

| | | | | | | | | |
|---|---|---|---|---|---|---|---|---|
| 279 | ENSG00000198363 | ASPH | 2.9821 | 0.8771 | 4.79E-06 | 1.623663996 | -0.87706674 | 4.79E-06 |
| 280 | ENSG00000079246 | XRCC5 | 1.4944 | -0.8823 | 4.83E-06 | 2.754599836 | 0.882256801 | 4.83E-06 |
| 281 | ENSG00000170144 | HNRNPA3 | 2.0006 | -0.881 | 4.97E-06 | 3.684275037 | 0.88097509 | 4.97E-06 |
| 282 | ENSG00000148180 | GSN | 2.241 | 0.8748 | 5.45E-06 | 1.222099821 | -0.87475763 | 5.45E-06 |
| 283 | ENSG00000023734 | STRAP | 1.4852 | -0.8763 | 5.62E-06 | 2.726425141 | 0.876317108 | 5.62E-06 |
| 284 | ENSG00000117519 | CNN3 | 1.72 | -0.8762 | 5.68E-06 | 3.157091203 | 0.876168982 | 5.68E-06 |
| 285 | ENSG00000163597 | SNHG16 | 1.1629 | -0.8754 | 5.81E-06 | 2.133320896 | 0.875397477 | 5.81E-06 |
| 286 | ENSG00000143543 | JTB | 1.3178 | -0.8748 | 5.85E-06 | 2.416623135 | 0.874809449 | 5.85E-06 |
| 287 | ENSG00000100804 | PSMB5 | 1.1584 | -0.8749 | 5.86E-06 | 2.124377973 | 0.874866434 | 5.86E-06 |
| 288 | ENSG00000136810 | TXN | 12.571 | 0.8633 | 6.72E-06 | 6.909888773 | -0.86331542 | 6.72E-06 |
| 289 | ENSG00000146674 | IGFBP3 | 1.7613 | -0.8708 | 7.24E-06 | 3.220858135 | 0.870754495 | 7.24E-06 |
| 290 | ENSG00000101084 | RAB5IF | 1.0334 | -0.8658 | 7.41E-06 | 1.883307866 | 0.865828138 | 7.41E-06 |
| 291 | ENSG00000197756 | RPL37A | 11.704 | -0.8626 | 7.80E-06 | 21.28245264 | 0.862614606 | 7.80E-06 |
| 292 | ENSG00000182220 | ATP6AP2 | 1.5206 | -0.8614 | 8.17E-06 | 2.762765113 | 0.861439636 | 8.17E-06 |
| 293 | ENSG00000115944 | COX7A2L | 1.8122 | -0.8587 | 8.75E-06 | 3.286210267 | 0.85868207 | 8.75E-06 |
| 294 | ENSG00000167978 | SRRM2 | 2.0712 | -0.8578 | 8.94E-06 | 3.753575215 | 0.857822203 | 8.94E-06 |
| 295 | ENSG00000099341 | PSMD8 | 1.9787 | -0.8555 | 9.43E-06 | 3.580369634 | 0.855531707 | 9.43E-06 |
| 296 | ENSG00000178913 | TAF7 | 3.1613 | 0.8498 | 9.59E-06 | 1.754099075 | -0.84977012 | 9.59E-06 |
| 297 | ENSG00000111716 | LDHB | 1.726 | -0.8544 | 9.78E-06 | 3.120511955 | 0.854354155 | 9.78E-06 |
| 298 | ENSG00000197747 | S100A10 | 26.293 | 0.8481 | 9.84E-06 | 14.60627625 | -0.84809723 | 9.84E-06 |
| 299 | ENSG00000185651 | UBE2L3 | 1.0471 | -0.8497 | 1.10E-05 | 1.887016637 | 0.849717485 | 1.10E-05 |
| 300 | ENSG00000230989 | HSBP1 | 1.5236 | -0.8489 | 1.12E-05 | 2.74419135 | 0.848907059 | 1.12E-05 |
| 301 | ENSG00000161970 | RPL26 | 14.777 | -0.8473 | 1.14E-05 | 26.58596507 | 0.84729067 | 1.14E-05 |
| 302 | ENSG00000180817 | PPA1 | 2.6659 | 0.8423 | 1.15E-05 | 1.486858206 | -0.84233365 | 1.15E-05 |
| 303 | ENSG00000130255 | RPL36 | 8.1729 | -0.8466 | 1.16E-05 | 14.6970215 | 0.846609859 | 1.16E-05 |
| 304 | ENSG00000127922 | SEM1 | 1.1 | -0.8476 | 1.16E-05 | 1.979466724 | 0.84761063 | 1.16E-05 |
| 305 | ENSG00000181163 | NPM1 | 10.645 | -0.8442 | 1.23E-05 | 19.11117666 | 0.844195483 | 1.23E-05 |
| 306 | ENSG00000127603 | MACF1 | 1.4095 | -0.8369 | 1.52E-05 | 2.517627323 | 0.836896105 | 1.52E-05 |
| 307 | ENSG00000163041 | H3F3A | 7.7164 | -0.8351 | 1.54E-05 | 13.765462 | 0.83505551 | 1.54E-05 |
| 308 | ENSG00000029363 | BCLAF1 | 1.3808 | -0.8313 | 1.72E-05 | 2.456911154 | 0.831348934 | 1.72E-05 |
| 309 | ENSG00000135316 | SYNCRIP | 1.3896 | -0.8306 | 1.74E-05 | 2.471357415 | 0.830573907 | 1.74E-05 |
| 310 | ENSG00000213585 | VDAC1 | 1.1729 | -0.8288 | 1.83E-05 | 2.083342218 | 0.828793545 | 1.83E-05 |
| 311 | ENSG00000254999 | BRK1 | 1.5649 | -0.8255 | 1.96E-05 | 2.773083871 | 0.825453448 | 1.96E-05 |
| 312 | ENSG00000197061 | HIST1H4C | 1.0917 | -0.8252 | 2.22E-05 | 1.934303466 | 0.825231468 | 2.22E-05 |
| 313 | ENSG00000177600 | RPLP2 | 8.2714 | -0.8199 | 2.22E-05 | 14.60122155 | 0.819880191 | 2.22E-05 |
| 314 | ENSG00000233016 | SNHG7 | 1.4528 | -0.8184 | 2.35E-05 | 2.562042845 | 0.818400894 | 2.35E-05 |
| 315 | ENSG00000134762 | DSC3 | 1.1602 | -0.818 | 2.48E-05 | 2.045476863 | 0.818010151 | 2.48E-05 |
| 316 | ENSG00000113282 | CLINT1 | 1.0518 | -0.8157 | 2.51E-05 | 1.851364582 | 0.815735016 | 2.51E-05 |
| 317 | ENSG00000109332 | UBE2D3 | 6.1673 | 0.8064 | 2.72E-05 | 3.526562547 | -0.80637512 | 2.72E-05 |
| 318 | ENSG00000104388 | RAB2A | 1.8381 | -0.8088 | 2.91E-05 | 3.22005058 | 0.808820859 | 2.91E-05 |
| 319 | ENSG00000074842 | MYDGF | 1.5379 | -0.808 | 2.99E-05 | 2.692507833 | 0.808000588 | 2.99E-05 |
| 320 | ENSG00000144713 | RPL32 | 16.965 | -0.806 | 3.07E-05 | 29.66149323 | 0.805993947 | 3.07E-05 |
| 321 | ENSG00000198918 | RPL39 | 12.823 | -0.803 | 3.30E-05 | 22.37220317 | 0.802992174 | 3.30E-05 |
| 322 | ENSG00000162980 | ARL5A | 1.1055 | -0.8033 | 3.35E-05 | 1.929308589 | 0.803308807 | 3.35E-05 |
| 323 | ENSG00000099624 | ATP5F1D | 1.5882 | -0.8009 | 3.52E-05 | 2.767012254 | 0.800890474 | 3.52E-05 |
| 324 | ENSG00000109390 | NDUFC1 | 1.7519 | 0.7945 | 3.69E-05 | 1.010011964 | -0.79451227 | 3.69E-05 |
| 325 | ENSG00000041357 | PSMA4 | 1.1953 | -0.7983 | 3.76E-05 | 2.078676345 | 0.798303822 | 3.76E-05 |

| | | | | | | | | |
|---|---|---|---|---|---|---|---|---|
| 326 | ENSG00000167088 | SNRPD1 | 1.1546 | -0.7969 | 3.89E-05 | 2.005996399 | 0.796902577 | 3.89E-05 |
| 327 | ENSG00000128524 | ATP6V1F | 1.4796 | -0.7941 | 4.14E-05 | 2.565572159 | 0.794092511 | 4.14E-05 |
| 328 | ENSG00000186395 | KRT10 | 2.5484 | -0.7933 | 4.22E-05 | 4.416547918 | 0.793349777 | 4.22E-05 |
| 329 | ENSG00000151914 | DST | 3.4587 | 0.7883 | 4.63E-05 | 2.002646542 | -0.78831233 | 4.63E-05 |
| 330 | ENSG00000148834 | GSTO1 | 1.9777 | 0.7802 | 5.16E-05 | 1.151543446 | -0.78021786 | 5.16E-05 |
| 331 | ENSG00000075624 | ACTB | 17.753 | -0.7835 | 5.23E-05 | 30.55799885 | 0.783461921 | 5.23E-05 |
| 332 | ENSG00000136942 | RPL35 | 6.6841 | -0.7825 | 5.32E-05 | 11.49760843 | 0.782531939 | 5.32E-05 |
| 333 | ENSG00000123505 | AMD1 | 3.3101 | 0.7765 | 5.56E-05 | 1.932359352 | -0.77651039 | 5.56E-05 |
| 334 | ENSG00000115875 | SRSF7 | 4.5456 | 0.7752 | 5.73E-05 | 2.655988404 | -0.77521203 | 5.73E-05 |
| 335 | ENSG00000105193 | RPS16 | 11.238 | -0.7776 | 5.95E-05 | 19.26443264 | 0.777575078 | 5.95E-05 |
| 336 | ENSG00000163468 | CCT3 | 1.1732 | -0.7776 | 6.07E-05 | 2.011230552 | 0.777582143 | 6.07E-05 |
| 337 | ENSG00000213741 | RPS29 | 6.9278 | -0.7735 | 6.53E-05 | 11.84279331 | 0.773547308 | 6.53E-05 |
| 338 | ENSG00000166710 | B2M | 19.165 | 0.7662 | 6.98E-05 | 11.26796373 | -0.7662188 | 6.98E-05 |
| 339 | ENSG00000107223 | EDF1 | 2.2516 | -0.7674 | 7.54E-05 | 3.8327156 | 0.767407091 | 7.54E-05 |
| 340 | ENSG00000171530 | TBCA | 1.6017 | -0.7663 | 7.77E-05 | 2.724361389 | 0.766265043 | 7.77E-05 |
| 341 | ENSG00000145824 | CXCL14 | 6.3063 | 0.7657 | 7.84E-05 | 3.709040055 | -0.76573596 | 7.84E-05 |
| 342 | ENSG00000106245 | BUD31 | 1.2055 | -0.7647 | 8.08E-05 | 2.048198623 | 0.764698083 | 8.08E-05 |
| 343 | ENSG00000147604 | RPL7 | 13.798 | -0.7637 | 8.13E-05 | 23.42588292 | 0.763690254 | 8.13E-05 |
| 344 | ENSG00000147123 | NDUFB11 | 1.1875 | -0.7638 | 8.24E-05 | 2.016345066 | 0.763770045 | 8.24E-05 |
| 345 | ENSG00000197321 | SVIL | 1.2514 | -0.7631 | 8.51E-05 | 2.123839603 | 0.763126555 | 8.51E-05 |
| 346 | ENSG00000131508 | UBE2D2 | 1.3388 | -0.7622 | 8.54E-05 | 2.270724875 | 0.762159623 | 8.54E-05 |
| 347 | ENSG00000100650 | SRSF5 | 4.3921 | 0.7558 | 8.85E-05 | 2.601074667 | -0.7557812 | 8.85E-05 |
| 348 | ENSG00000169021 | UQCRFS1 | 1.1 | -0.7601 | 8.98E-05 | 1.862939536 | 0.760081029 | 8.98E-05 |
| 349 | ENSG00000136888 | ATP6V1G1 | 4.6785 | -0.7587 | 9.12E-05 | 7.915623687 | 0.758651797 | 9.12E-05 |
| 350 | ENSG00000146731 | CCT6A | 1.6586 | -0.7572 | 9.52E-05 | 2.803352227 | 0.757214953 | 9.52E-05 |
| 351 | ENSG00000269821 | KCNQ1OT1 | 3.8325 | -0.7578 | 9.55E-05 | 6.480359269 | 0.757772133 | 9.55E-05 |
| 352 | ENSG00000107438 | PDLIM1 | 2.0995 | 0.7522 | 9.72E-05 | 1.246446107 | -0.75223016 | 9.72E-05 |
| 353 | ENSG00000119537 | KDSR | 1.6993 | 0.7508 | 0.0001 | 1.009832507 | -0.75082662 | 0.0001004 |
| 354 | ENSG00000120686 | UFM1 | 1.5432 | -0.7509 | 0.00011 | 2.596977074 | 0.750914588 | 0.0001096 |
| 355 | ENSG00000162244 | RPL29 | 12.354 | -0.7498 | 0.000111 | 20.77348365 | 0.74981747 | 0.0001106 |
| 356 | ENSG00000089009 | RPL6 | 12.251 | -0.7492 | 0.000112 | 20.59250162 | 0.749191911 | 0.000112 |
| 357 | ENSG00000112308 | C6orf62 | 2.2165 | 0.7455 | 0.000113 | 1.322027268 | -0.74553304 | 0.0001128 |
| 358 | ENSG00000054267 | ARID4B | 2.1487 | 0.7442 | 0.000116 | 1.28275617 | -0.74421368 | 0.000116 |
| 359 | ENSG00000140990 | NDUFB10 | 1.2128 | -0.7471 | 0.000119 | 2.035636657 | 0.747140952 | 0.0001193 |
| 360 | ENSG00000105993 | DNAJB6 | 4.4451 | 0.7406 | 0.000125 | 2.660295363 | -0.74061846 | 0.0001246 |
| 361 | ENSG00000113387 | SUB1 | 3.4724 | -0.7443 | 0.000125 | 5.816758469 | 0.744279999 | 0.0001253 |
| 362 | ENSG00000156508 | EEF1A1 | 38.658 | -0.7439 | 0.000126 | 64.73940703 | 0.743881537 | 0.0001256 |
| 363 | ENSG00000145592 | RPL37 | 11.714 | -0.7434 | 0.000127 | 19.60994652 | 0.74339668 | 0.0001271 |
| 364 | ENSG00000143947 | RPS27A | 18.936 | -0.7424 | 0.00013 | 31.67795793 | 0.742354682 | 0.0001299 |
| 365 | ENSG00000170296 | GABARAP | 2.8876 | -0.7425 | 0.00013 | 4.831391885 | 0.742541917 | 0.0001303 |
| 366 | ENSG00000125844 | RRBP1 | 1.2382 | -0.7429 | 0.000132 | 2.072245814 | 0.742930129 | 0.0001321 |
| 367 | ENSG00000044574 | HSPA5 | 8.1385 | 0.735 | 0.000142 | 4.88959566 | -0.73504491 | 0.0001417 |
| 368 | ENSG00000115738 | ID2 | 4.7901 | 0.7352 | 0.000143 | 2.877527644 | -0.73522896 | 0.0001428 |
| 369 | ENSG00000215301 | DDX3X | 3.6526 | 0.7341 | 0.000144 | 2.195921359 | -0.73407256 | 0.0001438 |
| 370 | ENSG00000182899 | RPL35A | 10.435 | -0.7352 | 0.000152 | 17.3697891 | 0.735196021 | 0.0001518 |
| 371 | ENSG00000107581 | EIF3A | 3.1536 | -0.7353 | 0.000153 | 5.249884805 | 0.735267952 | 0.0001526 |
| 372 | ENSG00000142676 | RPL11 | 21.896 | -0.7334 | 0.000158 | 36.40242356 | 0.733367587 | 0.0001578 |

| | | | | | | | | |
|---|---|---|---|---|---|---|---|---|
| 373 | ENSG00000051620 | HEBP2 | 1.1244 | -0.7343 | 0.00016 | 1.870626262 | 0.734325254 | 0.0001596 |
| 374 | ENSG00000146425 | DYNLT1 | 1.3001 | -0.728 | 0.000181 | 2.153330313 | 0.727965635 | 0.0001813 |
| 375 | ENSG00000111639 | MRPL51 | 1.3341 | -0.7275 | 0.000182 | 2.209081514 | 0.727530185 | 0.0001819 |
| 376 | ENSG00000172115 | CYCS | 5.2045 | 0.7228 | 0.000183 | 3.15350207 | -0.72280978 | 0.0001831 |
| 377 | ENSG00000009413 | REV3L | 1.6808 | 0.7238 | 0.000185 | 1.017698691 | -0.7238465 | 0.0001849 |
| 378 | ENSG00000120690 | ELF1 | 1.7267 | 0.7235 | 0.000185 | 1.045783657 | -0.723454 | 0.000185 |
| 379 | ENSG00000137575 | SDCBP | 5.0571 | 0.7204 | 0.000194 | 3.069247171 | -0.72042062 | 0.0001936 |
| 380 | ENSG00000157916 | RER1 | 1.142 | -0.7231 | 0.000202 | 1.885132342 | 0.723059076 | 0.0002022 |
| 381 | ENSG00000171634 | BPTF | 1.2078 | -0.7223 | 0.000205 | 1.992746516 | 0.72232804 | 0.0002052 |
| 382 | ENSG00000165629 | ATP5F1C | 1.3658 | -0.7194 | 0.000217 | 2.248861072 | 0.719421488 | 0.0002167 |
| 383 | ENSG00000156411 | ATP5MPL | 1.4704 | -0.7186 | 0.00022 | 2.41961408 | 0.718571696 | 0.0002201 |
| 384 | ENSG00000137154 | RPS6 | 22.437 | -0.716 | 0.00023 | 36.85462442 | 0.715978601 | 0.0002296 |
| 385 | ENSG00000186081 | KRT5 | 12.65 | 0.7143 | 0.000232 | 7.710325274 | -0.71431842 | 0.0002318 |
| 386 | ENSG00000169567 | HINT1 | 4.9267 | -0.7151 | 0.000234 | 8.087602981 | 0.71509696 | 0.0002344 |
| 387 | ENSG00000188846 | RPL14 | 13.738 | -0.7144 | 0.000238 | 22.54092234 | 0.714353137 | 0.0002376 |
| 388 | ENSG00000099783 | HNRNPM | 1.5273 | -0.7142 | 0.000241 | 2.505783184 | 0.714243099 | 0.0002414 |
| 389 | ENSG00000133872 | SARAF | 1.6889 | -0.7127 | 0.000249 | 2.767999266 | 0.712749096 | 0.0002486 |
| 390 | ENSG00000164924 | YWHAZ | 6.1104 | -0.7058 | 0.000287 | 9.966155246 | 0.705765579 | 0.000287 |
| 391 | ENSG00000255302 | EID1 | 2.8462 | -0.7052 | 0.00029 | 4.640420095 | 0.705225752 | 0.00029 |
| 392 | ENSG00000163602 | RYBP | 1.7798 | 0.7013 | 0.000295 | 1.094595867 | -0.70130865 | 0.0002955 |
| 393 | ENSG00000132432 | SEC61G | 2.3908 | -0.7043 | 0.000296 | 3.895525429 | 0.704312706 | 0.0002958 |
| 394 | ENSG00000174021 | GNG5 | 2.6577 | -0.7031 | 0.000302 | 4.326909319 | 0.703143118 | 0.0003024 |
| 395 | ENSG00000069345 | DNAJA2 | 1.081 | -0.7036 | 0.000303 | 1.760499695 | 0.703582602 | 0.0003033 |
| 396 | ENSG00000117318 | ID3 | 22.441 | 0.6994 | 0.000304 | 13.81935881 | -0.69943184 | 0.0003043 |
| 397 | ENSG00000100852 | ARHGAP5 | 1.2677 | -0.7035 | 0.000306 | 2.064319812 | 0.703484358 | 0.0003057 |
| 398 | ENSG00000154640 | BTG3 | 1.656 | 0.6991 | 0.000309 | 1.020001718 | -0.69909909 | 0.0003087 |
| 399 | ENSG00000103035 | PSMD7 | 1.3432 | -0.6995 | 0.000329 | 2.181176004 | 0.699464668 | 0.0003289 |
| 400 | ENSG00000039068 | CDH1 | 1.2507 | -0.7005 | 0.000342 | 2.032376528 | 0.700463132 | 0.0003421 |
| 401 | ENSG00000122884 | P4HA1 | 1.1775 | -0.6971 | 0.000351 | 1.909059896 | 0.697141016 | 0.0003507 |
| 402 | ENSG00000102317 | RBM3 | 3.525 | -0.6945 | 0.000361 | 5.704418603 | 0.694472779 | 0.0003607 |
| 403 | ENSG00000111843 | TMEM14C | 1.5331 | -0.6892 | 0.000405 | 2.471925695 | 0.689194859 | 0.0004049 |
| 404 | ENSG00000085224 | ATRX | 1.1484 | -0.6896 | 0.000406 | 1.852112318 | 0.689559613 | 0.0004063 |
| 405 | ENSG00000116209 | TMEM59 | 3.9252 | 0.6848 | 0.000408 | 2.441776977 | -0.68483441 | 0.0004081 |
| 406 | ENSG00000100902 | PSMA6 | 1.1481 | -0.689 | 0.000408 | 1.850945849 | 0.689010027 | 0.0004082 |
| 407 | ENSG00000115758 | ODC1 | 1.5982 | -0.6899 | 0.000411 | 2.578164034 | 0.689872144 | 0.0004111 |
| 408 | ENSG00000101856 | PGRMC1 | 3.1768 | 0.6838 | 0.00042 | 1.977642248 | -0.68379517 | 0.0004197 |
| 409 | ENSG00000211450 | SELENOH | 1.4326 | -0.6869 | 0.000424 | 2.306257293 | 0.68693373 | 0.0004245 |
| 410 | ENSG00000184076 | UQCR10 | 1.1628 | -0.6832 | 0.00046 | 1.867126858 | 0.68315874 | 0.0004596 |
| 411 | ENSG00000145632 | PLK2 | 1.7858 | 0.6787 | 0.000483 | 1.115592296 | -0.67873185 | 0.0004826 |
| 412 | ENSG00000136930 | PSMB7 | 1.1436 | -0.6792 | 0.000497 | 1.831265436 | 0.679229279 | 0.0004975 |
| 413 | ENSG00000127022 | CANX | 2.0205 | -0.6771 | 0.000517 | 3.230578704 | 0.677069826 | 0.0005166 |
| 414 | ENSG00000129625 | REEP5 | 1.1334 | -0.6767 | 0.000525 | 1.811704661 | 0.67670146 | 0.0005249 |
| 415 | ENSG00000115484 | CCT4 | 2.7137 | 0.6674 | 0.000585 | 1.708666631 | -0.66738327 | 0.0005853 |
| 416 | ENSG00000163359 | COL6A3 | 1.7323 | -0.6693 | 0.000625 | 2.754809202 | 0.669250032 | 0.0006246 |
| 417 | ENSG00000138758 | SEPTIN11 | 1.8228 | 0.6638 | 0.000637 | 1.150556434 | -0.66381076 | 0.0006369 |
| 418 | ENSG00000221983 | UBA52 | 6.6832 | -0.666 | 0.000639 | 10.60385447 | 0.665972413 | 0.0006385 |
| 419 | ENSG00000214253 | FIS1 | 1.1755 | -0.6652 | 0.000656 | 1.864135913 | 0.665240967 | 0.0006563 |

| | | | | | | | | |
|---|---|---|---|---|---|---|---|---|
| 420 | ENSG00000106028 | SSBP1 | 1.4058 | -0.6651 | 0.000656 | 2.229210569 | 0.665144109 | 0.0006563 |
| 421 | ENSG00000134363 | FST | 1.7132 | 0.6718 | 0.000661 | 1.075423915 | -0.67178033 | 0.0006607 |
| 422 | ENSG00000173812 | EIF1 | 33.902 | 0.6598 | 0.000671 | 21.45826034 | -0.6598323 | 0.0006714 |
| 423 | ENSG00000167642 | SPINT2 | 1.8516 | -0.6631 | 0.0007 | 2.932022649 | 0.663124447 | 0.0006995 |
| 424 | ENSG00000242485 | MRPL20 | 1.0947 | -0.6616 | 0.000706 | 1.731637083 | 0.661614851 | 0.000706 |
| 425 | ENSG00000184007 | PTP4A2 | 2.1829 | -0.6592 | 0.000733 | 3.447422162 | 0.659240464 | 0.0007332 |
| 426 | ENSG00000122565 | CBX3 | 1.1857 | -0.6596 | 0.000734 | 1.873019018 | 0.659635305 | 0.0007337 |
| 427 | ENSG00000179820 | MYADM | 1.5935 | -0.6575 | 0.000768 | 2.513440001 | 0.657471785 | 0.0007682 |
| 428 | ENSG00000084234 | APLP2 | 3.2018 | -0.6556 | 0.00079 | 5.043778837 | 0.655603294 | 0.00079 |
| 429 | ENSG00000111678 | C12orf57 | 2.6224 | 0.6519 | 0.000796 | 1.66903662 | -0.65188855 | 0.0007962 |
| 430 | ENSG00000087302 | RTRAF | 1.5117 | -0.6546 | 0.000805 | 2.379625155 | 0.654589879 | 0.0008053 |
| 431 | ENSG00000173575 | CHD2 | 2.0596 | 0.6479 | 0.000868 | 1.31443027 | -0.647883 | 0.000868 |
| 432 | ENSG00000118257 | NRP2 | 1.1705 | -0.6468 | 0.000966 | 1.832611361 | 0.646770745 | 0.0009659 |
| 433 | ENSG00000198755 | RPL10A | 12.042 | -0.6447 | 0.000968 | 18.82661822 | 0.644699327 | 0.0009683 |
| 434 | ENSG00000272888 | LINC01578 | 3.1508 | 0.6415 | 0.000973 | 2.019784652 | -0.64149436 | 0.0009734 |
| 435 | ENSG00000161960 | EIF4A1 | 9.9458 | 0.6409 | 0.000978 | 6.378457799 | -0.64087676 | 0.000978 |
| 436 | ENSG00000156482 | RPL30 | 18.945 | -0.6424 | 0.001009 | 29.57296128 | 0.642432916 | 0.0010093 |
| 437 | ENSG00000173660 | UQCRH | 2.0907 | -0.6415 | 0.001035 | 3.261475158 | 0.641513279 | 0.0010345 |
| 438 | ENSG00000172809 | RPL38 | 5.804 | -0.6411 | 0.001036 | 9.051584313 | 0.641111537 | 0.0010364 |
| 439 | ENSG00000189171 | S100A13 | 1.3395 | -0.6409 | 0.001059 | 2.088636189 | 0.64085074 | 0.0010592 |
| 440 | ENSG00000102554 | KLF5 | 4.2268 | 0.6325 | 0.001192 | 2.72645505 | -0.63252054 | 0.0011922 |
| 441 | ENSG00000196531 | NACA | 15.61 | -0.6329 | 0.001212 | 24.20684838 | 0.632918475 | 0.0012116 |
| 442 | ENSG00000134186 | PRPF38B | 1.6983 | 0.6285 | 0.00126 | 1.098514004 | -0.62850235 | 0.0012596 |
| 443 | ENSG00000116251 | RPL22 | 6.6304 | -0.6303 | 0.001273 | 10.2634551 | 0.630345467 | 0.001273 |
| 444 | ENSG00000171346 | KRT15 | 3.1414 | 0.6343 | 0.001299 | 2.023882246 | -0.63428672 | 0.0012989 |
| 445 | ENSG00000062194 | GPBP1 | 2.6974 | 0.6234 | 0.001381 | 1.750928674 | -0.62342558 | 0.0013807 |
| 446 | ENSG00000108518 | PFN1 | 4.5514 | -0.6241 | 0.001436 | 7.014811098 | 0.624078883 | 0.0014359 |
| 447 | ENSG00000104979 | C19orf53 | 1.1671 | -0.6239 | 0.001456 | 1.798574416 | 0.623920737 | 0.0014555 |
| 448 | ENSG00000124942 | AHNAK | 3.5798 | -0.62 | 0.001562 | 5.501542854 | 0.619956093 | 0.0015619 |
| 449 | ENSG00000010404 | IDS | 1.2838 | -0.6183 | 0.001617 | 1.970673348 | 0.618253497 | 0.001617 |
| 450 | ENSG00000127184 | COX7C | 2.2209 | -0.6157 | 0.001685 | 3.403096368 | 0.615671217 | 0.001685 |
| 451 | ENSG00000117523 | PRRC2C | 2.6561 | -0.6132 | 0.001763 | 4.06289867 | 0.613178935 | 0.0017631 |
| 452 | ENSG00000140575 | IQGAP1 | 1.3364 | -0.6061 | 0.002023 | 2.034260822 | 0.606118095 | 0.0020228 |
| 453 | ENSG00000147649 | MTDH | 2.2624 | -0.6051 | 0.002047 | 3.441290726 | 0.605065358 | 0.002047 |
| 454 | ENSG00000067082 | KLF6 | 5.7735 | 0.6008 | 0.002137 | 3.807053299 | -0.60075935 | 0.0021368 |
| 455 | ENSG00000100097 | LGALS1 | 5.8502 | 0.5974 | 0.002251 | 3.866602999 | -0.59740856 | 0.0022506 |
| 456 | ENSG00000138757 | G3BP2 | 1.1701 | -0.5983 | 0.002327 | 1.771476461 | 0.598293019 | 0.0023274 |
| 457 | ENSG00000160932 | LY6E | 1.711 | 0.5949 | 0.002369 | 1.132820135 | -0.594868 | 0.0023689 |
| 458 | ENSG00000142156 | COL6A1 | 2.3246 | -0.5975 | 0.002387 | 3.517410257 | 0.597511876 | 0.0023872 |
| 459 | ENSG00000127314 | RAP1B | 1.7234 | -0.5957 | 0.00243 | 2.604544163 | 0.595731356 | 0.0024296 |
| 460 | ENSG00000115053 | NCL | 6.5294 | -0.5944 | 0.002475 | 9.85812234 | 0.594355026 | 0.0024749 |
| 461 | ENSG00000143761 | ARF1 | 2.1963 | -0.5937 | 0.002509 | 3.31447469 | 0.593710233 | 0.0025087 |
| 462 | ENSG00000134294 | SLC38A2 | 3.4258 | 0.5881 | 0.002649 | 2.278860243 | -0.58811355 | 0.0026486 |
| 463 | ENSG00000007168 | PAFAH1B1 | 1.0891 | -0.5867 | 0.002865 | 1.635687592 | 0.586681739 | 0.0028653 |
| 464 | ENSG00000071082 | RPL31 | 3.9471 | -0.5842 | 0.002964 | 5.91752338 | 0.584193065 | 0.0029638 |
| 465 | ENSG00000129562 | DAD1 | 2.1601 | -0.584 | 0.002985 | 3.238026155 | 0.583995005 | 0.0029851 |
| 466 | ENSG00000142227 | EMP3 | 6.1733 | 0.5811 | 0.002996 | 4.126545963 | -0.58110153 | 0.0029956 |

| | | | | | | | | |
|---|---|---|---|---|---|---|---|---|
| 467 | ENSG00000142192 | APP | 2.1822 | -0.583 | 0.003045 | 3.268952518 | 0.583034663 | 0.0030452 |
| 468 | ENSG00000196683 | TOMM7 | 2.6539 | 0.5799 | 0.00306 | 1.775454416 | -0.57990257 | 0.0030596 |
| 469 | ENSG00000167815 | PRDX2 | 2.4232 | -0.5812 | 0.003142 | 3.625233798 | 0.581181561 | 0.0031423 |
| 470 | ENSG00000116288 | PARK7 | 2.6071 | -0.5785 | 0.003279 | 3.893282221 | 0.578535746 | 0.0032787 |
| 471 | ENSG00000178741 | COX5A | 1.1631 | -0.5785 | 0.00331 | 1.736931055 | 0.578526263 | 0.0033104 |
| 472 | ENSG00000177469 | CAVIN1 | 1.9558 | -0.5784 | 0.00332 | 2.920328057 | 0.578357114 | 0.0033199 |
| 473 | ENSG00000174574 | AKIRIN1 | 1.5035 | -0.5705 | 0.003801 | 2.232739883 | 0.570455994 | 0.0038006 |
| 474 | ENSG00000112514 | CUTA | 1.3713 | -0.5688 | 0.003904 | 2.034171094 | 0.568841872 | 0.0039039 |
| 475 | ENSG00000111341 | MGP | 2.9431 | -0.5701 | 0.00412 | 4.369470455 | 0.570107753 | 0.0041199 |
| 476 | ENSG00000115419 | GLS | 2.1835 | 0.5629 | 0.004178 | 1.47806483 | -0.56292923 | 0.0041776 |
| 477 | ENSG00000101745 | ANKRD12 | 2.3352 | 0.56 | 0.004353 | 1.584004075 | -0.55996249 | 0.0043534 |
| 478 | ENSG00000100632 | ERH | 2.8269 | -0.5603 | 0.004492 | 4.168479002 | 0.560318398 | 0.0044917 |
| 479 | ENSG00000100387 | RBX1 | 1.5842 | -0.5603 | 0.004506 | 2.336077007 | 0.560322359 | 0.0045063 |
| 480 | ENSG00000143774 | GUK1 | 3.0486 | -0.5582 | 0.004658 | 4.488928769 | 0.558214259 | 0.0046578 |
| 481 | ENSG00000179271 | GADD45GI | 1.0601 | -0.5574 | 0.004765 | 1.56013634 | 0.557405333 | 0.004765 |
| 482 | ENSG00000122566 | HNRNPA2E | 8.691 | 0.5543 | 0.004766 | 5.91863003 | -0.55425751 | 0.0047661 |
| 483 | ENSG00000181029 | TRAPPC5 | 1.1993 | -0.556 | 0.004874 | 1.763191545 | 0.556007428 | 0.0048738 |
| 484 | ENSG00000086598 | TMED2 | 1.5231 | -0.5553 | 0.004916 | 2.238153492 | 0.555319991 | 0.0049157 |
| 485 | ENSG00000170606 | HSPA4 | 1.0288 | -0.5553 | 0.004947 | 1.511742862 | 0.555268505 | 0.0049474 |
| 486 | ENSG00000068697 | LAPTM4A | 5.7673 | 0.5519 | 0.00496 | 3.934048791 | -0.5518705 | 0.0049603 |
| 487 | ENSG00000075415 | SLC25A3 | 4.5679 | -0.5535 | 0.00503 | 6.704261356 | 0.553535487 | 0.0050298 |
| 488 | ENSG00000136238 | RAC1 | 4.2022 | -0.5521 | 0.005151 | 6.161554522 | 0.552135725 | 0.0051513 |
| 489 | ENSG00000088930 | XRN2 | 1.0888 | -0.548 | 0.005579 | 1.591870258 | 0.547971039 | 0.0055788 |
| 490 | ENSG00000165732 | DDX21 | 2.3446 | 0.5433 | 0.005769 | 1.608828912 | -0.54334651 | 0.0057692 |
| 491 | ENSG00000145907 | G3BP1 | 1.494 | -0.5446 | 0.005888 | 2.179291709 | 0.54464565 | 0.005888 |
| 492 | ENSG00000109046 | WSB1 | 4.5456 | 0.5409 | 0.005991 | 3.124310454 | -0.54093558 | 0.0059913 |
| 493 | ENSG00000135404 | CD63 | 9.1272 | -0.5411 | 0.006193 | 13.28107858 | 0.541129829 | 0.0061933 |
| 494 | ENSG00000136758 | YME1L1 | 2.4328 | 0.5374 | 0.006371 | 1.676155067 | -0.53742493 | 0.0063712 |
| 495 | ENSG00000144674 | GOLGA4 | 1.9771 | -0.5364 | 0.006743 | 2.867507981 | 0.536382478 | 0.0067426 |
| 496 | ENSG00000198886 | MT-ND4 | 12.054 | -0.5387 | 0.006743 | 17.51021393 | 0.538710704 | 0.0067433 |
| 497 | ENSG00000167283 | ATP5MG | 1.9641 | -0.5359 | 0.006775 | 2.847618202 | 0.535878324 | 0.0067749 |
| 498 | ENSG00000047410 | TPR | 1.3378 | -0.5332 | 0.007113 | 1.935978394 | 0.533168908 | 0.0071132 |
| 499 | ENSG00000198242 | RPL23A | 10.128 | -0.5321 | 0.007166 | 14.64485943 | 0.532089963 | 0.0071661 |
| 500 | ENSG00000126457 | PRMT1 | 1.1432 | -0.5324 | 0.007219 | 1.6534538 | 0.532354713 | 0.007219 |
| 501 | ENSG00000172757 | CFL1 | 6.3491 | -0.5299 | 0.007438 | 9.166855305 | 0.529872472 | 0.0074378 |
| 502 | ENSG00000198898 | CAPZA2 | 1.9588 | 0.5273 | 0.007517 | 1.359114977 | -0.52726696 | 0.007517 |
| 503 | ENSG00000091409 | ITGA6 | 1.5279 | -0.5305 | 0.00762 | 2.206957943 | 0.530472833 | 0.0076203 |
| 504 | ENSG00000174886 | NDUFA11 | 1.6271 | -0.5261 | 0.007952 | 2.343165545 | 0.526120981 | 0.0079519 |
| 505 | ENSG00000141867 | BRD4 | 1.0113 | -0.5248 | 0.008183 | 1.455004649 | 0.524797983 | 0.0081833 |
| 506 | ENSG00000182196 | ARL6IP4 | 1.7388 | -0.5242 | 0.008199 | 2.500638759 | 0.524181299 | 0.0081993 |
| 507 | ENSG00000164405 | UQCRQ | 1.1241 | -0.523 | 0.008386 | 1.615259442 | 0.523021979 | 0.0083863 |
| 508 | ENSG00000228474 | OST4 | 1.5854 | -0.5222 | 0.008462 | 2.27688622 | 0.522198054 | 0.0084625 |
| 509 | ENSG00000182768 | NGRN | 1.6434 | 0.5153 | 0.009118 | 1.149778789 | -0.51528736 | 0.009118 |
| 510 | ENSG00000144224 | UBXN4 | 1.954 | -0.5172 | 0.00915 | 2.796473055 | 0.517172354 | 0.0091495 |
| 511 | ENSG00000067560 | RHOA | 2.5247 | -0.5164 | 0.009237 | 3.611415635 | 0.516443161 | 0.0092369 |
| 512 | ENSG00000198804 | MT-CO1 | 25.42 | -0.5185 | 0.009456 | 36.41160576 | 0.518453242 | 0.0094557 |
| 513 | ENSG00000173674 | EIF1AX | 4.1007 | 0.512 | 0.009563 | 2.875523712 | -0.51202215 | 0.0095628 |

| | | | | | | | | |
|---|---|---|---|---|---|---|---|---|
| 514 | ENSG00000006327 | TNFRSF12A | 4.9381 | -0.5106 | 0.010214 | 7.034970062 | 0.510574769 | 0.010214 |
| 515 | ENSG00000189043 | NDUFA4 | 3.2101 | -0.5097 | 0.010277 | 4.57040209 | 0.509688316 | 0.010277 |
| 516 | ENSG00000155368 | DBI | 1.5563 | -0.5078 | 0.010685 | 2.212880013 | 0.507822476 | 0.0106851 |
| 517 | ENSG00000198727 | MT-CYB | 13.943 | -0.5084 | 0.010874 | 19.83366915 | 0.508408298 | 0.0108742 |
| 518 | ENSG00000091039 | OSBPL8 | 1.2555 | -0.5064 | 0.010934 | 1.783470147 | 0.506388432 | 0.0109336 |
| 519 | ENSG00000189403 | HMGB1 | 5.9218 | -0.5043 | 0.011178 | 8.399648195 | 0.504292849 | 0.0111778 |
| 520 | ENSG00000130770 | ATP5IF1 | 1.0405 | -0.5045 | 0.011268 | 1.476120716 | 0.504474208 | 0.011268 |
| 521 | ENSG00000158710 | TAGLN2 | 3.4392 | 0.5011 | 0.01136 | 2.429962747 | -0.50114216 | 0.0113604 |
| 522 | ENSG00000114416 | FXR1 | 1.4887 | 0.4997 | 0.011686 | 1.052872195 | -0.49972905 | 0.011686 |
| 523 | ENSG00000178982 | EIF3K | 2.3556 | -0.5008 | 0.011801 | 3.33325782 | 0.500832107 | 0.0118007 |
| 524 | ENSG00000104408 | EIF3E | 3.1467 | -0.499 | 0.012132 | 4.44702564 | 0.499011286 | 0.0121325 |
| 525 | ENSG00000135378 | PRRG4 | 1.732 | 0.4985 | 0.012212 | 1.226017958 | -0.49846455 | 0.0122125 |
| 526 | ENSG00000171159 | C9orf16 | 1.282 | -0.4986 | 0.012321 | 1.811256019 | 0.498629727 | 0.0123212 |
| 527 | ENSG00000065135 | GNAI3 | 1.2669 | -0.4981 | 0.012364 | 1.789392217 | 0.498122672 | 0.0123643 |
| 528 | ENSG00000109180 | OCIAD1 | 1.0902 | -0.4974 | 0.012538 | 1.538990364 | 0.49738213 | 0.0125379 |
| 529 | ENSG00000092199 | HNRNPC | 4.7438 | 0.4918 | 0.013098 | 3.373515929 | -0.49179678 | 0.0130977 |
| 530 | ENSG00000173230 | GOLGB1 | 2.0101 | -0.4937 | 0.013197 | 2.830360454 | 0.493748553 | 0.0131972 |
| 531 | ENSG00000165389 | SPTSSA | 1.2713 | -0.4917 | 0.013654 | 1.787567741 | 0.491666435 | 0.0136539 |
| 532 | ENSG00000070831 | CDC42 | 3.3424 | -0.4881 | 0.014294 | 4.688035927 | 0.488108244 | 0.0142941 |
| 533 | ENSG00000108298 | RPL19 | 18.928 | -0.4877 | 0.014333 | 26.54116073 | 0.487695466 | 0.0143332 |
| 534 | ENSG00000092841 | MYL6 | 8.6658 | -0.4843 | 0.0151 | 12.12241668 | 0.484280301 | 0.0150995 |
| 535 | ENSG00000166595 | CIAO2B | 1.2092 | -0.484 | 0.015262 | 1.691259336 | 0.484040549 | 0.0152623 |
| 536 | ENSG00000198832 | SELENOM | 4.0275 | 0.4791 | 0.015922 | 2.889491421 | -0.4790764 | 0.0159217 |
| 537 | ENSG00000198786 | MT-ND5 | 3.3239 | -0.4835 | 0.016129 | 4.647299267 | 0.483494039 | 0.0161286 |
| 538 | ENSG00000150753 | CCT5 | 1.4688 | -0.4791 | 0.016421 | 2.047391068 | 0.479144309 | 0.0164214 |
| 539 | ENSG00000023191 | RNH1 | 1.8985 | 0.4743 | 0.017099 | 1.366502609 | -0.47432956 | 0.0170993 |
| 540 | ENSG00000143878 | RHOB | 1.7099 | 0.4717 | 0.018182 | 1.233016767 | -0.47173987 | 0.0181819 |
| 541 | ENSG00000132507 | EIF5A | 1.7419 | -0.4709 | 0.018501 | 2.414320108 | 0.470947267 | 0.018501 |
| 542 | ENSG00000174444 | RPL4 | 5.8807 | -0.469 | 0.01894 | 8.13973514 | 0.468990097 | 0.0189396 |
| 543 | ENSG00000091527 | CDV3 | 2.1423 | -0.4685 | 0.019138 | 2.964354756 | 0.468532913 | 0.0191381 |
| 544 | ENSG00000111642 | CHD4 | 1.0054 | -0.4678 | 0.019498 | 1.390430163 | 0.467762594 | 0.0194984 |
| 545 | ENSG00000173457 | PPP1R14B | 2.6733 | -0.4671 | 0.019546 | 3.69540135 | 0.467094869 | 0.0195459 |
| 546 | ENSG00000125534 | PPDPF | 3.7457 | -0.4636 | 0.02055 | 5.165121445 | 0.463580155 | 0.0205501 |
| 547 | ENSG00000196504 | PRPF40A | 1.3757 | -0.4604 | 0.021584 | 1.892908797 | 0.460401699 | 0.021584 |
| 548 | ENSG00000153147 | SMARCA5 | 1.4232 | -0.4585 | 0.022189 | 1.955598989 | 0.458464821 | 0.0221889 |
| 549 | ENSG00000126777 | KTN1 | 2.0774 | -0.4582 | 0.022238 | 2.853988913 | 0.458230674 | 0.0222379 |
| 550 | ENSG00000113013 | HSPA9 | 2.3106 | 0.4562 | 0.022289 | 1.684230617 | -0.45615614 | 0.022289 |
| 551 | ENSG00000149100 | EIF3M | 1.4039 | -0.4568 | 0.022721 | 1.926885924 | 0.456777941 | 0.022721 |
| 552 | ENSG00000128245 | YWHAH | 1.2316 | -0.4572 | 0.022725 | 1.690870513 | 0.457189758 | 0.0227254 |
| 553 | ENSG00000152661 | GJA1 | 2.383 | 0.4544 | 0.022974 | 1.739054625 | -0.45444452 | 0.0229742 |
| 554 | ENSG00000131871 | SELENOS | 1.5331 | -0.4556 | 0.023093 | 2.10254408 | 0.455638795 | 0.0230928 |
| 555 | ENSG00000188243 | COMMD6 | 1.5445 | -0.4534 | 0.023816 | 2.114806951 | 0.453423351 | 0.0238157 |
| 556 | ENSG00000165637 | VDAC2 | 2.5405 | -0.4523 | 0.024167 | 3.475925861 | 0.452280581 | 0.0241666 |
| 557 | ENSG00000088986 | DYNLL1 | 11.212 | 0.4492 | 0.02454 | 8.212504813 | -0.44919216 | 0.0245398 |
| 558 | ENSG00000119396 | RAB14 | 1.063 | -0.4508 | 0.024778 | 1.452910988 | 0.450838367 | 0.0247779 |
| 559 | ENSG00000197958 | RPL12 | 17.841 | -0.448 | 0.025496 | 24.33728346 | 0.44796864 | 0.0254961 |
| 560 | ENSG00000139343 | SNRPF | 1.0834 | -0.4474 | 0.025944 | 1.477317094 | 0.447362676 | 0.0259438 |

| | | | | | | | | |
|---|---|---|---|---|---|---|---|---|
| 561 | ENSG00000089693 | MLF2 | 1.0761 | -0.445 | 0.026839 | 1.464814947 | 0.444951346 | 0.026839 |
| 562 | ENSG00000008988 | RPS20 | 10.353 | -0.4435 | 0.027191 | 14.07903259 | 0.443464041 | 0.0271909 |
| 563 | ENSG00000115524 | SF3B1 | 1.3215 | -0.4438 | 0.027242 | 1.797557494 | 0.443831615 | 0.027242 |
| 564 | ENSG00000243678 | NME2 | 7.2067 | -0.443 | 0.027374 | 9.796748164 | 0.442974248 | 0.0273741 |
| 565 | ENSG00000106615 | RHEB | 2.4077 | -0.4429 | 0.027479 | 3.272930474 | 0.442915261 | 0.0274792 |
| 566 | ENSG00000143183 | TMCO1 | 1.0121 | -0.4398 | 0.028835 | 1.37290323 | 0.43982789 | 0.028835 |
| 567 | ENSG00000185222 | TCEAL9 | 2.5918 | -0.4392 | 0.029108 | 3.514090309 | 0.439183649 | 0.0291079 |
| 568 | ENSG00000170776 | AKAP13 | 1.0938 | -0.4379 | 0.029687 | 1.481773601 | 0.43793857 | 0.0296865 |
| 569 | ENSG00000089280 | FUS | 2.6367 | -0.4371 | 0.029783 | 3.569721872 | 0.437066401 | 0.029783 |
| 570 | ENSG00000067064 | IDI1 | 2.0003 | 0.4345 | 0.030309 | 1.480128581 | -0.43445948 | 0.0303088 |
| 571 | ENSG00000186184 | POLR1D | 2.0556 | 0.4342 | 0.030382 | 1.521343793 | -0.43421628 | 0.0303823 |
| 572 | ENSG00000131143 | COX4I1 | 4.5527 | -0.4306 | 0.032403 | 6.136281043 | 0.430627185 | 0.032403 |
| 573 | ENSG00000130779 | CLIP1 | 1.1869 | -0.4303 | 0.032971 | 1.59931771 | 0.430261871 | 0.0329706 |
| 574 | ENSG00000094975 | SUCO | 1.682 | -0.4289 | 0.033485 | 2.264294345 | 0.428899706 | 0.0334852 |
| 575 | ENSG00000158417 | EIF5B | 1.9302 | 0.4256 | 0.034093 | 1.437058984 | -0.42564654 | 0.0340932 |
| 576 | ENSG00000127914 | AKAP9 | 1.1353 | -0.427 | 0.034412 | 1.526278851 | 0.42699237 | 0.0344116 |
| 577 | ENSG00000142864 | SERBP1 | 3.4365 | -0.4247 | 0.035152 | 4.612753861 | 0.424703634 | 0.0351522 |
| 578 | ENSG00000110700 | RPS13 | 14.871 | -0.4242 | 0.035296 | 19.95375556 | 0.42415185 | 0.0352959 |
| 579 | ENSG00000120705 | ETF1 | 1.6667 | 0.422 | 0.035751 | 1.243993533 | -0.42199805 | 0.0357508 |
| 580 | ENSG00000154723 | ATP5PF | 1.8905 | -0.423 | 0.035978 | 2.534765433 | 0.423044917 | 0.0359777 |
| 581 | ENSG00000142507 | PSMB6 | 1.6054 | -0.423 | 0.036035 | 2.152313392 | 0.422951859 | 0.0360354 |
| 582 | ENSG00000100567 | PSMA3 | 1.5601 | -0.4218 | 0.036629 | 2.089892385 | 0.421796729 | 0.0366286 |
| 583 | ENSG00000063046 | EIF4B | 1.442 | -0.4205 | 0.037299 | 1.929936687 | 0.420496755 | 0.0372992 |
| 584 | ENSG00000137210 | TMEM14B | 1.2805 | -0.4201 | 0.037571 | 1.713242776 | 0.420051818 | 0.0375712 |
| 585 | ENSG00000180530 | NRIP1 | 1.7729 | 0.4183 | 0.037925 | 1.326663232 | -0.41833762 | 0.0379248 |
| 586 | ENSG00000179010 | MRFAP1 | 2.0542 | -0.418 | 0.038437 | 2.744490444 | 0.417989368 | 0.038437 |
| 587 | ENSG00000168906 | MAT2A | 2.0325 | 0.4135 | 0.0401 | 1.526039575 | -0.4134661 | 0.0401002 |
| 588 | ENSG00000084733 | RAB10 | 1.212 | -0.4122 | 0.04173 | 1.612747049 | 0.412169682 | 0.0417302 |
| 589 | ENSG00000232112 | TMA7 | 2.5267 | -0.4091 | 0.043216 | 3.355061803 | 0.409057372 | 0.0432158 |
| 590 | ENSG00000166598 | HSP90B1 | 6.2006 | -0.4078 | 0.043915 | 8.2260837 | 0.40779965 | 0.0439152 |
| 591 | ENSG00000146223 | RPL7L1 | 1.062 | -0.4071 | 0.04454 | 1.408196372 | 0.407122172 | 0.0445402 |
| 592 | ENSG00000104529 | EEF1D | 4.6653 | -0.405 | 0.045533 | 6.177107432 | 0.404973395 | 0.0455328 |
| 593 | ENSG00000132963 | POMP | 2.8654 | -0.4047 | 0.04569 | 3.793294955 | 0.404728485 | 0.0456904 |
| 594 | ENSG00000277443 | MARCKS | 2.5022 | -0.4044 | 0.046064 | 3.31181275 | 0.404431919 | 0.0460642 |
| 595 | ENSG00000134248 | LAMTOR5 | 1.6526 | -0.4019 | 0.047486 | 2.183628578 | 0.401943798 | 0.0474862 |
| 596 | ENSG00000103342 | GSPT1 | 1.4361 | -0.4018 | 0.047655 | 1.897335395 | 0.401843444 | 0.0476554 |
| 597 | ENSG00000241837 | ATP5PO | 1.3323 | -0.4016 | 0.047702 | 1.759871597 | 0.401587335 | 0.047702 |
| 598 | ENSG00000080371 | RAB21 | 1.371 | 0.3989 | 0.048474 | 1.039831678 | -0.39886111 | 0.0484744 |
| 599 | ENSG00000101421 | CHMP4B | 1.0433 | -0.3988 | 0.049513 | 1.375535261 | 0.398831104 | 0.0495135 |
| 600 | ENSG00000197170 | PSMD12 | 1.1811 | -0.3976 | 0.050242 | 1.555919108 | 0.397619545 | 0.0502421 |
| 601 | ENSG00000120963 | ZNF706 | 2.8196 | -0.3958 | 0.051269 | 3.709668154 | 0.395807057 | 0.051269 |
| 602 | ENSG00000150459 | SAP18 | 4.4546 | 0.3938 | 0.051502 | 3.390534402 | -0.39378622 | 0.0515016 |
| 603 | ENSG00000164919 | COX6C | 1.7983 | -0.395 | 0.051851 | 2.364670434 | 0.394998351 | 0.0518507 |
| 604 | ENSG00000124172 | ATP5F1E | 3.2499 | -0.393 | 0.053056 | 4.267479256 | 0.392997265 | 0.0530564 |
| 605 | ENSG00000172354 | GNB2 | 1.0954 | -0.3928 | 0.053484 | 1.438135724 | 0.392762205 | 0.0534839 |
| 606 | ENSG00000009307 | CSDE1 | 2.5733 | -0.3917 | 0.053951 | 3.376058232 | 0.391721106 | 0.0539508 |
| 607 | ENSG00000219200 | RNASEK | 2.6631 | -0.3912 | 0.054318 | 3.492615329 | 0.391204358 | 0.0543182 |

| | | | | | | | | |
|---|---|---|---|---|---|---|---|---|
| 608 | ENSG00000242372 | EIF6 | 1.5675 | -0.3913 | 0.054492 | 2.05585544 | 0.391277102 | 0.0544917 |
| 609 | ENSG00000120742 | SERP1 | 4.0569 | 0.389 | 0.05478 | 3.098049964 | -0.38901135 | 0.0547798 |
| 610 | ENSG00000141552 | ANAPC11 | 1.1113 | -0.3902 | 0.055284 | 1.456410393 | 0.390163579 | 0.0552842 |
| 611 | ENSG00000138385 | SSB | 1.4854 | -0.3888 | 0.056151 | 1.944891408 | 0.388786495 | 0.0561506 |
| 612 | ENSG00000106588 | PSMA2 | 1.7908 | -0.3878 | 0.056621 | 2.343105726 | 0.387809925 | 0.0566205 |
| 613 | ENSG00000224032 | EPB41L4A- | 2.2457 | 0.3831 | 0.059053 | 1.721976333 | -0.38308895 | 0.0590525 |
| 614 | ENSG00000183696 | UPP1 | 1.3651 | 0.3826 | 0.060233 | 1.047039854 | -0.38262894 | 0.0602332 |
| 615 | ENSG00000134352 | IL6ST | 2.2342 | -0.383 | 0.060432 | 2.913418975 | 0.382957099 | 0.0604315 |
| 616 | ENSG00000132475 | H3F3B | 24.344 | 0.3796 | 0.061345 | 18.71260343 | -0.37955438 | 0.0613447 |
| 617 | ENSG00000104904 | OAZ1 | 7.2461 | 0.3788 | 0.061989 | 5.572936693 | -0.37876638 | 0.0619895 |
| 618 | ENSG00000163191 | S100A11 | 22.325 | 0.3781 | 0.06261 | 17.17836866 | -0.37808955 | 0.0626096 |
| 619 | ENSG00000167004 | PDIA3 | 2.5712 | -0.3786 | 0.063403 | 3.342828841 | 0.378609316 | 0.0634033 |
| 620 | ENSG00000062716 | VMP1 | 2.1041 | 0.3765 | 0.064949 | 1.62076278 | -0.37650458 | 0.0649491 |
| 621 | ENSG00000165119 | HNRNPK | 3.3589 | -0.376 | 0.065399 | 4.358882513 | 0.375969331 | 0.065399 |
| 622 | ENSG00000116754 | SRSF11 | 2.2664 | -0.3759 | 0.065564 | 2.941025391 | 0.375886231 | 0.0655637 |
| 623 | ENSG00000163463 | KRTCAP2 | 1.0324 | -0.3762 | 0.065617 | 1.340002843 | 0.37618038 | 0.0656171 |
| 624 | ENSG00000196262 | PPIA | 7.9065 | -0.3723 | 0.068317 | 10.23459249 | 0.372338511 | 0.068317 |
| 625 | ENSG00000026025 | VIM | 34.651 | 0.3699 | 0.069319 | 26.81429376 | -0.3698769 | 0.0693194 |
| 626 | ENSG00000092820 | EZR | 9.247 | 0.3699 | 0.069539 | 7.155415387 | -0.36994891 | 0.0695389 |
| 627 | ENSG00000205339 | IPO7 | 1.1646 | -0.3713 | 0.069654 | 1.506418981 | 0.371261509 | 0.0696544 |
| 628 | ENSG00000087365 | SF3B2 | 1.1717 | -0.3707 | 0.070026 | 1.515002991 | 0.370708019 | 0.0700258 |
| 629 | ENSG00000177954 | RPS27 | 17.819 | -0.3688 | 0.071239 | 23.00906493 | 0.36880011 | 0.0712387 |
| 630 | ENSG00000177700 | POLR2L | 2.2869 | 0.3653 | 0.073266 | 1.775394598 | -0.36525071 | 0.0732661 |
| 631 | ENSG00000118058 | KMT2A | 1.0053 | -0.367 | 0.07335 | 1.296544423 | 0.367041984 | 0.0733496 |
| 632 | ENSG00000166848 | TERF2IP | 1.3678 | 0.3651 | 0.073558 | 1.061964666 | -0.36512875 | 0.073558 |
| 633 | ENSG00000162191 | UBXN1 | 1.2977 | -0.366 | 0.07386 | 1.672476206 | 0.366034362 | 0.0738597 |
| 634 | ENSG00000108953 | YWHAE | 3.9523 | -0.3645 | 0.074916 | 5.088523363 | 0.364544399 | 0.0749163 |
| 635 | ENSG00000244754 | N4BP2L2 | 1.7262 | -0.3647 | 0.075033 | 2.222660401 | 0.364705192 | 0.0750331 |
| 636 | ENSG00000173915 | ATP5MD | 1.6285 | -0.3643 | 0.075399 | 2.096203278 | 0.364260126 | 0.0753991 |
| 637 | ENSG00000087086 | FTL | 108.78 | 0.3601 | 0.07772 | 84.75283193 | -0.36007757 | 0.0777201 |
| 638 | ENSG00000221914 | PPP2R2A | 1.4254 | 0.3577 | 0.080395 | 1.112391986 | -0.35765235 | 0.0803955 |
| 639 | ENSG00000147684 | NDUFB9 | 1.6341 | 0.356 | 0.081802 | 1.276744372 | -0.35598143 | 0.0818021 |
| 640 | ENSG00000129084 | PSMA1 | 1.1663 | -0.3572 | 0.081943 | 1.493976653 | 0.357211646 | 0.0819433 |
| 641 | ENSG00000116030 | SUMO1 | 2.0228 | -0.3568 | 0.082112 | 2.590396996 | 0.356843786 | 0.0821124 |
| 642 | ENSG00000105185 | PDCD5 | 1.3352 | -0.357 | 0.082118 | 1.710042466 | 0.356987958 | 0.0821178 |
| 643 | ENSG00000158195 | WASF2 | 1.4616 | -0.3555 | 0.083598 | 1.870028074 | 0.355458847 | 0.0835978 |
| 644 | ENSG00000170540 | ARL6IP1 | 4.2688 | 0.3536 | 0.083943 | 3.340884728 | -0.35360801 | 0.0839432 |
| 645 | ENSG00000156261 | CCT8 | 1.2172 | -0.3546 | 0.084489 | 1.55636775 | 0.354568417 | 0.0844889 |
| 646 | ENSG00000115268 | RPS15 | 13.068 | -0.354 | 0.084602 | 16.70197106 | 0.353997231 | 0.0846022 |
| 647 | ENSG00000161011 | SQSTM1 | 6.8197 | 0.3509 | 0.086704 | 5.34724004 | -0.35090265 | 0.086704 |
| 648 | ENSG00000229117 | RPL41 | 21.319 | -0.3456 | 0.092861 | 27.08892226 | 0.345594155 | 0.0928606 |
| 649 | ENSG00000119048 | UBE2B | 3.9302 | 0.3442 | 0.093173 | 3.095986212 | -0.34421457 | 0.0931729 |
| 650 | ENSG00000185883 | ATP6V0C | 2.4209 | -0.3449 | 0.093892 | 3.074750508 | 0.344922798 | 0.0938915 |
| 651 | ENSG00000136943 | CTSV | 1.7821 | 0.3449 | 0.094264 | 1.403141676 | -0.34488247 | 0.0942637 |
| 652 | ENSG00000143198 | MGST3 | 2.0962 | -0.3433 | 0.095573 | 2.659457899 | 0.343335649 | 0.0955728 |
| 653 | ENSG00000105974 | CAV1 | 1.9622 | -0.3443 | 0.095684 | 2.491097647 | 0.344330738 | 0.0956839 |
| 654 | ENSG00000125835 | SNRPB | 2.8934 | -0.3428 | 0.096093 | 3.669410044 | 0.342783498 | 0.0960931 |

| | | | | | | | | |
|---|---|---|---|---|---|---|---|---|
| 655 | ENSG00000166295 | ANAPC16 | 1.0528 | -0.3422 | 0.097156 | 1.334589234 | 0.342200334 | 0.0971556 |
| 656 | ENSG00000169020 | ATP5ME | 1.1022 | -0.3421 | 0.097327 | 1.397129878 | 0.342101432 | 0.0973272 |
| 657 | ENSG00000168003 | SLC3A2 | 3.7046 | 0.3396 | 0.09848 | 2.927476413 | -0.33964642 | 0.0984796 |
| 658 | ENSG00000008018 | PSMB1 | 3.3587 | -0.3386 | 0.10071 | 4.247230564 | 0.338629238 | 0.1007101 |
| 659 | ENSG00000153113 | CAST | 5.4043 | 0.3367 | 0.101654 | 4.279443033 | -0.33669285 | 0.1016539 |
| 660 | ENSG00000240972 | MIF | 9.7746 | -0.3359 | 0.10354 | 12.33749549 | 0.335944819 | 0.1035402 |
| 661 | ENSG00000123144 | TRIR | 1.9108 | -0.3321 | 0.107968 | 2.405437004 | 0.332081134 | 0.1079679 |
| 662 | ENSG00000120533 | ENY2 | 1.7022 | -0.3321 | 0.108005 | 2.142802189 | 0.332107653 | 0.1080053 |
| 663 | ENSG00000127540 | UQCR11 | 1.4899 | -0.3312 | 0.109222 | 1.874364943 | 0.331185686 | 0.1092219 |
| 664 | ENSG00000132780 | NASP | 1.3297 | -0.3312 | 0.1094 | 1.672835119 | 0.331203442 | 0.1093998 |
| 665 | ENSG00000183255 | PTTG1IP | 1.4915 | -0.3307 | 0.109826 | 1.875710868 | 0.330714426 | 0.1098261 |
| 666 | ENSG00000152795 | HNRNPDL | 4.8181 | 0.3261 | 0.113705 | 3.843303543 | -0.32611264 | 0.1137045 |
| 667 | ENSG00000129559 | NEDD8 | 1.6811 | -0.3263 | 0.11523 | 2.107778232 | 0.32629874 | 0.1152303 |
| 668 | ENSG00000169504 | CLIC4 | 1.0409 | -0.3194 | 0.124641 | 1.29887736 | 0.319360111 | 0.1246414 |
| 669 | ENSG00000166226 | CCT2 | 1.9393 | -0.319 | 0.124668 | 2.419165438 | 0.318978832 | 0.1246681 |
| 670 | ENSG00000138668 | HNRNPD | 1.6863 | -0.3176 | 0.126626 | 2.101616887 | 0.31761038 | 0.126626 |
| 671 | ENSG00000110955 | ATP5F1B | 1.7624 | -0.3158 | 0.129057 | 2.193678151 | 0.315808924 | 0.1290566 |
| 672 | ENSG00000125743 | SNRPD2 | 1.3595 | -0.3152 | 0.129958 | 1.69155843 | 0.31524706 | 0.1299581 |
| 673 | ENSG00000100664 | EIF5 | 7.4347 | 0.3136 | 0.130268 | 5.982157686 | -0.31360526 | 0.1302682 |
| 674 | ENSG00000105583 | WDR83OS | 1.3907 | -0.314 | 0.131642 | 1.728915324 | 0.314023815 | 0.131642 |
| 675 | ENSG00000095787 | WAC | 1.9551 | 0.3116 | 0.133411 | 1.575300427 | -0.31161799 | 0.1334111 |
| 676 | ENSG00000118181 | RPS25 | 9.6113 | -0.312 | 0.134096 | 11.93162435 | 0.311985853 | 0.1340958 |
| 677 | ENSG00000169714 | CNBP | 3.7765 | 0.3098 | 0.135764 | 3.04675527 | -0.30976543 | 0.1357641 |
| 678 | ENSG00000198431 | TXNRD1 | 3.7376 | 0.3092 | 0.137386 | 3.016576643 | -0.309192 | 0.137386 |
| 679 | ENSG00000225921 | NOL7 | 1.4939 | -0.3091 | 0.138664 | 1.85088603 | 0.30912864 | 0.138664 |
| 680 | ENSG00000164615 | CAMLG | 1.3617 | 0.3076 | 0.13945 | 1.100248752 | -0.30758718 | 0.1394503 |
| 681 | ENSG00000160789 | LMNA | 8.6547 | 0.3071 | 0.139797 | 6.995160594 | -0.30711797 | 0.1397965 |
| 682 | ENSG00000169756 | LIMS1 | 1.0229 | -0.3042 | 0.146283 | 1.262986029 | 0.304211702 | 0.1462833 |
| 683 | ENSG00000111328 | CDK2AP1 | 1.0067 | -0.3035 | 0.147247 | 1.242348513 | 0.303482335 | 0.147247 |
| 684 | ENSG00000123349 | PFDN5 | 4.4941 | -0.2983 | 0.15465 | 5.526487329 | 0.298344388 | 0.1546504 |
| 685 | ENSG00000169045 | HNRNPH1 | 3.1583 | -0.2984 | 0.154813 | 3.883920566 | 0.298363564 | 0.1548133 |
| 686 | ENSG00000164171 | ITGA2 | 1.4738 | -0.2987 | 0.156774 | 1.81284122 | 0.298690202 | 0.1567741 |
| 687 | ENSG00000110651 | CD81 | 3.9949 | 0.2945 | 0.15914 | 3.257257926 | -0.29448737 | 0.15914 |
| 688 | ENSG00000069275 | NUCKS1 | 2.7812 | -0.2947 | 0.160487 | 3.411500921 | 0.294715717 | 0.1604872 |
| 689 | ENSG00000198843 | SELENOT | 1.1122 | -0.2942 | 0.161653 | 1.363810759 | 0.294237602 | 0.161653 |
| 690 | ENSG00000186591 | UBE2H | 2.1357 | -0.2936 | 0.162761 | 2.617644498 | 0.293586432 | 0.1627613 |
| 691 | ENSG00000162434 | JAK1 | 2.0055 | -0.2933 | 0.162898 | 2.4576888 | 0.293309382 | 0.1628979 |
| 692 | ENSG00000188529 | SRSF10 | 1.2814 | -0.293 | 0.163725 | 1.569946637 | 0.292966442 | 0.1637254 |
| 693 | ENSG00000103769 | RAB11A | 1.0174 | -0.2926 | 0.164338 | 1.246206831 | 0.292645915 | 0.1643385 |
| 694 | ENSG00000084623 | EIF3I | 1.5013 | 0.2907 | 0.165592 | 1.227274154 | -0.29071947 | 0.1655916 |
| 695 | ENSG00000137776 | SLTM | 1.0206 | -0.2919 | 0.165649 | 1.249556689 | 0.291930232 | 0.1656492 |
| 696 | ENSG00000169976 | SF3B5 | 1.5351 | -0.2916 | 0.165683 | 1.879000906 | 0.291618355 | 0.1656833 |
| 697 | ENSG00000140319 | SRP14 | 7.6003 | -0.2906 | 0.166818 | 9.296602466 | 0.290639979 | 0.166818 |
| 698 | ENSG00000144746 | ARL6IP5 | 1.7667 | 0.2892 | 0.168151 | 1.445822451 | -0.28917547 | 0.1681515 |
| 699 | ENSG00000089220 | PEBP1 | 4.3327 | 0.2886 | 0.168671 | 3.547200062 | -0.28857235 | 0.1686713 |
| 700 | ENSG00000108561 | C1QBP | 1.4861 | -0.2892 | 0.169698 | 1.815951802 | 0.289238159 | 0.1696984 |
| 701 | ENSG00000134644 | PUM1 | 1.2832 | -0.2872 | 0.172454 | 1.565789225 | 0.287174831 | 0.1724543 |

| | | | | | | | | |
|---|---|---|---|---|---|---|---|---|
| 702 | ENSG00000100941 | PNN | 1.9341 | -0.2856 | 0.175011 | 2.357551986 | 0.285593888 | 0.1750108 |
| 703 | ENSG00000174953 | DHX36 | 1.5122 | 0.2824 | 0.178517 | 1.243305615 | -0.28243203 | 0.1785166 |
| 704 | ENSG00000128989 | ARPP19 | 1.0189 | -0.2832 | 0.179216 | 1.239925848 | 0.283195334 | 0.1792157 |
| 705 | ENSG00000170759 | KIF5B | 2.4206 | -0.2823 | 0.180192 | 2.94377706 | 0.282330792 | 0.180192 |
| 706 | ENSG00000110696 | C11orf58 | 2.1218 | -0.2821 | 0.180638 | 2.579958601 | 0.282054428 | 0.1806378 |
| 707 | ENSG00000104695 | PPP2CB | 1.3459 | 0.2808 | 0.181462 | 1.107815841 | -0.28083638 | 0.1814619 |
| 708 | ENSG00000196586 | MYO6 | 1.0369 | -0.2818 | 0.182488 | 1.260623183 | 0.281792356 | 0.1824883 |
| 709 | ENSG00000169564 | PCBP1 | 3.2486 | 0.2764 | 0.188666 | 2.682248894 | -0.27635934 | 0.1886661 |
| 710 | ENSG00000104765 | BNIP3L | 1.5065 | 0.2727 | 0.195272 | 1.247104115 | -0.27265553 | 0.1952723 |
| 711 | ENSG00000187735 | TCEA1 | 1.253 | -0.2719 | 0.198388 | 1.512849511 | 0.271898016 | 0.1983875 |
| 712 | ENSG00000125977 | EIF2S2 | 1.7776 | 0.2667 | 0.20649 | 1.477586279 | -0.26669655 | 0.2064899 |
| 713 | ENSG00000120616 | EPC1 | 1.3896 | 0.2664 | 0.207307 | 1.155341945 | -0.26638997 | 0.2073068 |
| 714 | ENSG00000182117 | NOP10 | 1.1364 | -0.2646 | 0.212381 | 1.365156684 | 0.264631117 | 0.2123813 |
| 715 | ENSG00000114902 | SPCS1 | 2.3003 | -0.2642 | 0.212735 | 2.762585657 | 0.264205753 | 0.2127349 |
| 716 | ENSG00000177410 | ZFAS1 | 6.879 | -0.262 | 0.217039 | 8.248994333 | 0.262016071 | 0.2170386 |
| 717 | ENSG00000185088 | RPS27L | 1.3127 | -0.262 | 0.217895 | 1.574133959 | 0.262030972 | 0.2178949 |
| 718 | ENSG00000065978 | YBX1 | 8.5801 | -0.2589 | 0.223345 | 10.26686478 | 0.258937291 | 0.2233454 |
| 719 | ENSG00000067225 | PKM | 4.1177 | 0.2569 | 0.226028 | 3.446046327 | -0.25689794 | 0.2260285 |
| 720 | ENSG00000083857 | FAT1 | 1.2954 | -0.258 | 0.227954 | 1.549039937 | 0.258006081 | 0.2279539 |
| 721 | ENSG00000171735 | CAMTA1 | 1.1891 | -0.257 | 0.228031 | 1.420997613 | 0.256973249 | 0.2280313 |
| 722 | ENSG00000198931 | APRT | 1.4349 | -0.2565 | 0.229064 | 1.71411015 | 0.256480576 | 0.2290642 |
| 723 | ENSG00000124193 | SRSF6 | 1.0095 | -0.2565 | 0.229341 | 1.205888903 | 0.25646335 | 0.2293413 |
| 724 | ENSG00000214736 | TOMM6 | 1.865 | -0.2547 | 0.232497 | 2.225172794 | 0.25474022 | 0.2324968 |
| 725 | ENSG00000171988 | JMJD1C | 1.3526 | -0.2536 | 0.2357 | 1.612537683 | 0.253563612 | 0.2357002 |
| 726 | ENSG00000025796 | SEC63 | 1.0369 | -0.2516 | 0.239835 | 1.23448233 | 0.251562168 | 0.2398349 |
| 727 | ENSG00000251562 | MALAT1 | 199.38 | -0.2437 | 0.256482 | 236.0732856 | 0.243744735 | 0.2564817 |
| 728 | ENSG00000196937 | FAM3C | 2.1687 | 0.2424 | 0.257939 | 1.833329188 | -0.24238314 | 0.2579394 |
| 729 | ENSG00000122026 | RPL21 | 17.591 | -0.2417 | 0.260833 | 20.79917586 | 0.24168908 | 0.2608334 |
| 730 | ENSG00000123200 | ZC3H13 | 1.0067 | -0.242 | 0.261322 | 1.190515449 | 0.241954707 | 0.2613222 |
| 731 | ENSG00000180879 | SSR4 | 4.0288 | 0.2405 | 0.262032 | 3.410244725 | -0.2404742 | 0.2620316 |
| 732 | ENSG00000023909 | GCLM | 1.2642 | 0.2385 | 0.267706 | 1.071535687 | -0.23849963 | 0.2677056 |
| 733 | ENSG00000112531 | QKI | 1.3939 | -0.2386 | 0.268724 | 1.644630515 | 0.238628987 | 0.2687242 |
| 734 | ENSG00000114023 | FAM162A | 1.2961 | -0.2382 | 0.270036 | 1.528761335 | 0.238181613 | 0.2700359 |
| 735 | ENSG00000107937 | GTPBP4 | 1.0177 | -0.2382 | 0.270038 | 1.200355656 | 0.238160188 | 0.2700379 |
| 736 | ENSG00000102144 | PGK1 | 2.3087 | -0.2378 | 0.270349 | 2.722387366 | 0.237787769 | 0.2703486 |
| 737 | ENSG00000188186 | LAMTOR4 | 1.4366 | 0.2357 | 0.273467 | 1.220065979 | -0.23567014 | 0.2734669 |
| 738 | ENSG00000134308 | YWHAQ | 2.9501 | -0.2361 | 0.274033 | 3.474639755 | 0.236092677 | 0.2740331 |
| 739 | ENSG00000178980 | SELENOW | 2.5573 | 0.2319 | 0.282542 | 2.17764669 | -0.23187122 | 0.2825423 |
| 740 | ENSG00000023287 | RB1CC1 | 1.2135 | -0.23 | 0.289665 | 1.423240822 | 0.230018485 | 0.2896646 |
| 741 | ENSG00000136950 | ARPC5L | 1.0634 | -0.23 | 0.290075 | 1.247163934 | 0.229987546 | 0.2900753 |
| 742 | ENSG00000173113 | TRMT112 | 1.7271 | -0.2296 | 0.290285 | 2.025018805 | 0.229611472 | 0.2902852 |
| 743 | ENSG00000050405 | LIMA1 | 1.3036 | -0.2286 | 0.292753 | 1.527475229 | 0.228609472 | 0.2927535 |
| 744 | ENSG00000150687 | PRSS23 | 1.1508 | -0.2275 | 0.296077 | 1.347360566 | 0.22746236 | 0.2960769 |
| 745 | ENSG00000072110 | ACTN1 | 1.0471 | -0.2249 | 0.302226 | 1.223714931 | 0.224890252 | 0.3022259 |
| 746 | ENSG00000065548 | ZC3H15 | 1.3824 | -0.2236 | 0.304985 | 1.614212612 | 0.223617711 | 0.3049848 |
| 747 | ENSG00000135940 | COX5B | 2.1373 | 0.2226 | 0.305295 | 1.831714078 | -0.22261852 | 0.3052954 |
| 748 | ENSG00000143621 | ILF2 | 1.4948 | -0.2211 | 0.310574 | 1.742374573 | 0.221140573 | 0.3105737 |

| | | | | | | | | |
|---|---|---|---|---|---|---|---|---|
| 749 | ENSG00000168374 | ARF4 | 4.9763 | 0.2154 | 0.321002 | 4.286112839 | -0.21540189 | 0.3210018 |
| 750 | ENSG00000116560 | SFPQ | 4.6662 | -0.2155 | 0.322203 | 5.418035691 | 0.215518096 | 0.3222029 |
| 751 | ENSG00000120727 | PAIP2 | 1.1524 | -0.2155 | 0.322793 | 1.33799891 | 0.215492848 | 0.3227926 |
| 752 | ENSG00000171490 | RSL1D1 | 1.6263 | -0.213 | 0.329022 | 1.885132342 | 0.213036869 | 0.3290216 |
| 753 | ENSG00000166136 | NDUFB8 | 1.6098 | -0.2121 | 0.331435 | 1.864764012 | 0.21209629 | 0.3314346 |
| 754 | ENSG00000169764 | UGP2 | 1.3753 | 0.2103 | 0.334659 | 1.188750792 | -0.21034349 | 0.334659 |
| 755 | ENSG00000048649 | RSF1 | 1.0639 | -0.2108 | 0.335548 | 1.23128202 | 0.210765472 | 0.3355478 |
| 756 | ENSG00000164096 | C4orf3 | 2.2485 | 0.2091 | 0.337598 | 1.945100774 | -0.20910745 | 0.337598 |
| 757 | ENSG00000170348 | TMED10 | 1.6324 | -0.2097 | 0.338014 | 1.887764373 | 0.209676138 | 0.3380142 |
| 758 | ENSG00000112378 | PERP | 10.833 | -0.2099 | 0.338196 | 12.52972347 | 0.209915069 | 0.3381961 |
| 759 | ENSG00000140307 | GTF2A2 | 1.8108 | -0.2087 | 0.339723 | 2.092584235 | 0.208663909 | 0.3397231 |
| 760 | ENSG00000137818 | RPLP1 | 47.122 | 0.206 | 0.34334 | 40.85270933 | -0.20595141 | 0.3433399 |
| 761 | ENSG00000083937 | CHMP2B | 1.7806 | 0.2061 | 0.343531 | 1.54350669 | -0.20612471 | 0.3435306 |
| 762 | ENSG00000244687 | UBE2V1 | 1.2235 | -0.2067 | 0.344314 | 1.411994871 | 0.206685745 | 0.344314 |
| 763 | ENSG00000155438 | NIFK | 1.2764 | 0.2051 | 0.346655 | 1.107247561 | -0.20513243 | 0.3466549 |
| 764 | ENSG00000115310 | RTN4 | 7.1018 | -0.2045 | 0.34958 | 8.183462745 | 0.204522573 | 0.3495798 |
| 765 | ENSG00000111237 | VPS29 | 1.1673 | 0.2035 | 0.351182 | 1.013720735 | -0.20345065 | 0.3511822 |
| 766 | ENSG00000077782 | FGFR1 | 1.4256 | 0.2032 | 0.352315 | 1.238340648 | -0.20320273 | 0.3523147 |
| 767 | ENSG00000164190 | NIPBL | 1.2507 | -0.2014 | 0.358814 | 1.438075905 | 0.201407373 | 0.3588137 |
| 768 | ENSG00000182718 | ANXA2 | 11.489 | 0.2003 | 0.359021 | 9.999534184 | -0.20027286 | 0.3590205 |
| 769 | ENSG00000112306 | RPS12 | 28.069 | -0.2001 | 0.361482 | 32.24435304 | 0.200079257 | 0.3614821 |
| 770 | ENSG00000090273 | NUDC | 1.4952 | -0.2002 | 0.361923 | 1.717759102 | 0.200153909 | 0.3619227 |
| 771 | ENSG00000213923 | CSNK1E | 1.0334 | -0.1988 | 0.366649 | 1.186088852 | 0.198829195 | 0.3666489 |
| 772 | ENSG00000111229 | ARPC3 | 2.5148 | -0.1983 | 0.367213 | 2.88527419 | 0.198266238 | 0.3672135 |
| 773 | ENSG00000101608 | MYL12A | 3.9767 | -0.1976 | 0.368959 | 4.560352517 | 0.197586438 | 0.3689585 |
| 774 | ENSG00000108946 | PRKAR1A | 1.1056 | -0.1967 | 0.371877 | 1.267083622 | 0.196665392 | 0.3718775 |
| 775 | ENSG00000113732 | ATP6V0E1 | 2.8841 | -0.192 | 0.384611 | 3.294584911 | 0.191967105 | 0.384611 |
| 776 | ENSG00000111142 | METAP2 | 1.2897 | 0.1882 | 0.393886 | 1.131952761 | -0.1882314 | 0.3938863 |
| 777 | ENSG00000177733 | HNRNPA0 | 1.9003 | -0.1887 | 0.394262 | 2.16583246 | 0.188668285 | 0.394262 |
| 778 | ENSG00000137815 | RTF1 | 1.3615 | 0.1854 | 0.402483 | 1.197274983 | -0.18542989 | 0.4024831 |
| 779 | ENSG00000186010 | NDUFA13 | 2.0921 | -0.186 | 0.402568 | 2.379954159 | 0.185962913 | 0.4025684 |
| 780 | ENSG00000185787 | MORF4L1 | 3.5104 | -0.1846 | 0.406367 | 3.989620536 | 0.184629274 | 0.4063673 |
| 781 | ENSG00000180370 | PAK2 | 1.4158 | -0.1836 | 0.410012 | 1.607961538 | 0.183616323 | 0.4100125 |
| 782 | ENSG00000159140 | SON | 2.6435 | -0.1829 | 0.411739 | 3.000874185 | 0.182906774 | 0.4117385 |
| 783 | ENSG00000115461 | IGFBP5 | 2.6894 | -0.1839 | 0.414703 | 3.054980367 | 0.183878673 | 0.4147034 |
| 784 | ENSG00000135241 | PNPLA8 | 1.2882 | 0.1781 | 0.425266 | 1.138592657 | -0.17812218 | 0.4252656 |
| 785 | ENSG00000173436 | MICOS10 | 1.1278 | -0.1785 | 0.426095 | 1.27632564 | 0.178496171 | 0.4260954 |
| 786 | ENSG00000114956 | DGUOK | 1.1716 | -0.1783 | 0.426618 | 1.325736039 | 0.178302372 | 0.4266176 |
| 787 | ENSG00000105968 | H2AFV | 1.1937 | 0.1777 | 0.426618 | 1.055354679 | -0.17769443 | 0.4266176 |
| 788 | ENSG00000164346 | NSA2 | 1.187 | -0.178 | 0.427763 | 1.342784421 | 0.177960924 | 0.4277632 |
| 789 | ENSG00000106052 | TAX1BP1 | 2.0629 | -0.1775 | 0.428672 | 2.332996334 | 0.177536159 | 0.4286722 |
| 790 | ENSG00000105373 | NOP53 | 1.4245 | -0.1773 | 0.429555 | 1.610862754 | 0.177340396 | 0.4295546 |
| 791 | ENSG00000143549 | TPM3 | 3.0739 | -0.1772 | 0.429638 | 3.475507128 | 0.177164449 | 0.4296382 |
| 792 | ENSG00000186432 | KPNA4 | 1.6107 | -0.177 | 0.43051 | 1.820916769 | 0.177000121 | 0.4305097 |
| 793 | ENSG00000187514 | PTMA | 16.414 | -0.1765 | 0.431457 | 18.5490586 | 0.176464602 | 0.431457 |
| 794 | ENSG00000130396 | AFDN | 1.0957 | -0.1757 | 0.435527 | 1.237652731 | 0.175713535 | 0.4355266 |
| 795 | ENSG00000136153 | LMO7 | 1.7283 | 0.1732 | 0.442977 | 1.5327692 | -0.17318478 | 0.4429774 |

| | | | | | | | | |
|---|---|---|---|---|---|---|---|---|
| 796 | ENSG00000176171 | BNIP3 | 1.3754 | 0.1709 | 0.448727 | 1.221770817 | -0.17091708 | 0.4487265 |
| 797 | ENSG00000163956 | LRPAP1 | 1.137 | 0.1689 | 0.455014 | 1.011357889 | -0.168897 | 0.4550145 |
| 798 | ENSG00000165527 | ARF6 | 1.3948 | 0.1662 | 0.463671 | 1.243006521 | -0.16624557 | 0.4636713 |
| 799 | ENSG00000090060 | PAPOLA | 1.659 | -0.1666 | 0.464069 | 1.862042252 | 0.16659626 | 0.4640688 |
| 800 | ENSG00000010244 | ZNF207 | 1.5035 | -0.165 | 0.468044 | 1.685606451 | 0.164975801 | 0.4680443 |
| 801 | ENSG00000034713 | GABARAPL | 2.3339 | 0.1638 | 0.469534 | 2.083402036 | -0.16379873 | 0.4695339 |
| 802 | ENSG00000132341 | RAN | 4.7348 | -0.1643 | 0.469856 | 5.305815462 | 0.16427677 | 0.4698559 |
| 803 | ENSG00000100380 | ST13 | 2.4844 | -0.1643 | 0.469963 | 2.784060637 | 0.164281543 | 0.4699629 |
| 804 | ENSG00000106153 | CHCHD2 | 7.8554 | -0.1637 | 0.471553 | 8.799208437 | 0.163696162 | 0.4715527 |
| 805 | ENSG00000077721 | UBE2A | 1.1148 | -0.1616 | 0.479416 | 1.246894749 | 0.161602123 | 0.4794156 |
| 806 | ENSG00000034510 | TMSB10 | 14.366 | -0.1613 | 0.479584 | 16.06492985 | 0.16129418 | 0.479584 |
| 807 | ENSG00000184990 | SIVA1 | 1.509 | -0.161 | 0.480961 | 1.687221561 | 0.161031566 | 0.4809615 |
| 808 | ENSG00000107262 | BAG1 | 2.3994 | 0.16 | 0.482424 | 2.147557791 | -0.15998602 | 0.4824237 |
| 809 | ENSG00000010278 | CD9 | 6.0541 | 0.1597 | 0.48349 | 5.419770438 | -0.15968821 | 0.4834898 |
| 810 | ENSG00000091164 | TXNL1 | 1.5912 | -0.1595 | 0.484618 | 1.777219074 | 0.159481123 | 0.4846181 |
| 811 | ENSG00000100138 | SNU13 | 2.0632 | 0.1585 | 0.485679 | 1.848583003 | -0.15848985 | 0.4856786 |
| 812 | ENSG00000170515 | PA2G4 | 1.5739 | 0.1572 | 0.49014 | 1.411426592 | -0.15722252 | 0.4901398 |
| 813 | ENSG00000183726 | TMEM50A | 1.5989 | -0.1576 | 0.490695 | 1.783559875 | 0.157635648 | 0.4906955 |
| 814 | ENSG00000197451 | HNRNPAB | 2.6841 | -0.1573 | 0.491634 | 2.993247277 | 0.157293742 | 0.4916337 |
| 815 | ENSG00000245910 | SNHG6 | 2.9488 | 0.155 | 0.497525 | 2.648451224 | -0.15499624 | 0.4975246 |
| 816 | ENSG00000160948 | VPS28 | 1.0089 | -0.1553 | 0.499094 | 1.123608027 | 0.155325403 | 0.499094 |
| 817 | ENSG00000226950 | DANCR | 1.34 | 0.154 | 0.501528 | 1.204303702 | -0.1540435 | 0.5015278 |
| 818 | ENSG00000087191 | PSMC5 | 1.2532 | 0.1525 | 0.506464 | 1.127526163 | -0.15245588 | 0.5064636 |
| 819 | ENSG00000163399 | ATP1A1 | 1.4103 | 0.1513 | 0.510768 | 1.2698652 | -0.15132179 | 0.5107678 |
| 820 | ENSG00000131051 | RBM39 | 3.2451 | 0.1506 | 0.512266 | 2.92337882 | -0.15061394 | 0.5122655 |
| 821 | ENSG00000163331 | DAPL1 | 1.3433 | 0.1512 | 0.512927 | 1.209597674 | -0.15123487 | 0.5129273 |
| 822 | ENSG00000090013 | BLVRB | 1.6221 | 0.1505 | 0.512995 | 1.46140527 | -0.15048405 | 0.5129946 |
| 823 | ENSG00000143321 | HDGF | 1.4482 | -0.1489 | 0.520792 | 1.605688421 | 0.148890638 | 0.5207916 |
| 824 | ENSG00000065518 | NDUFB4 | 1.3264 | 0.1473 | 0.524472 | 1.197693716 | -0.1472673 | 0.5244722 |
| 825 | ENSG00000198258 | UBL5 | 2.3127 | -0.1476 | 0.524955 | 2.561863388 | 0.147610296 | 0.5249549 |
| 826 | ENSG00000119655 | NPC2 | 1.844 | -0.1476 | 0.525149 | 2.042665376 | 0.147616707 | 0.5251489 |
| 827 | ENSG00000008952 | SEC62 | 1.8331 | 0.1455 | 0.530634 | 1.657282209 | -0.14548488 | 0.5306343 |
| 828 | ENSG00000133112 | TPT1 | 61.132 | -0.1448 | 0.53424 | 67.58687569 | 0.144826673 | 0.53424 |
| 829 | ENSG00000126432 | PRDX5 | 4.3647 | 0.1428 | 0.539768 | 3.95346002 | -0.14276423 | 0.5397675 |
| 830 | ENSG00000124795 | DEK | 1.6974 | -0.1398 | 0.552853 | 1.870117802 | 0.139840323 | 0.5528529 |
| 831 | ENSG00000070756 | PABPC1 | 5.5742 | 0.1371 | 0.559484 | 5.068753221 | -0.1371184 | 0.5594835 |
| 832 | ENSG00000196754 | S100A2 | 7.9873 | -0.1366 | 0.567296 | 8.780544945 | 0.136604409 | 0.5672956 |
| 833 | ENSG00000115128 | SF3B6 | 3.4849 | 0.133 | 0.573889 | 3.177908175 | -0.13304387 | 0.5738888 |
| 834 | ENSG00000170027 | YWHAG | 1.8685 | 0.1329 | 0.57481 | 1.704030668 | -0.1328966 | 0.5748098 |
| 835 | ENSG00000119707 | RBM25 | 2.5124 | 0.1316 | 0.579145 | 2.293306504 | -0.13164136 | 0.5791454 |
| 836 | ENSG00000126267 | COX6B1 | 1.6398 | -0.1311 | 0.583155 | 1.795852656 | 0.131111405 | 0.5831549 |
| 837 | ENSG00000153037 | SRP19 | 1.6002 | 0.1303 | 0.58402 | 1.462033368 | -0.1303174 | 0.5840197 |
| 838 | ENSG00000134001 | EIF2S1 | 1.0969 | 0.127 | 0.584377 | 1.004448808 | -0.12702798 | 0.5843769 |
| 839 | ENSG00000133398 | MED10 | 1.1553 | -0.1299 | 0.584377 | 1.264152497 | 0.129933189 | 0.5843769 |
| 840 | ENSG00000105887 | MTPN | 2.0756 | -0.1275 | 0.584377 | 2.267345108 | 0.127450962 | 0.5843769 |
| 841 | ENSG00000081154 | PCNP | 1.2816 | 0.1283 | 0.584377 | 1.172599693 | -0.12827957 | 0.5843769 |
| 842 | ENSG00000162545 | CAMK2N1 | 2.388 | -0.1268 | 0.586885 | 2.607445379 | 0.126844755 | 0.5868849 |

| | | | | | | | | |
|---|---|---|---|---|---|---|---|---|
| 843 | ENSG00000197728 | RPS26 | 6.4839 | -0.1259 | 0.58931 | 7.075377719 | 0.125946968 | 0.5893098 |
| 844 | ENSG00000114391 | RPL24 | 18.741 | -0.1253 | 0.591819 | 20.44142902 | 0.125266938 | 0.5918192 |
| 845 | ENSG00000167863 | ATP5PD | 1.1422 | 0.1241 | 0.595134 | 1.047996956 | -0.12411447 | 0.5951339 |
| 846 | ENSG00000111775 | COX6A1 | 2.9506 | 0.1234 | 0.596653 | 2.70874866 | -0.12335821 | 0.5966534 |
| 847 | ENSG00000184897 | H1FX | 1.3998 | 0.1222 | 0.596653 | 1.286135937 | -0.12218801 | 0.5966534 |
| 848 | ENSG00000163714 | U2SURP | 1.2115 | 0.121 | 0.599656 | 1.114007095 | -0.12101263 | 0.5996562 |
| 849 | ENSG00000213625 | LEPROT | 1.0178 | -0.1197 | 0.606593 | 1.105841818 | 0.119711242 | 0.6065929 |
| 850 | ENSG00000138398 | PPIG | 1.6412 | 0.1174 | 0.612262 | 1.512969149 | -0.11739999 | 0.6122615 |
| 851 | ENSG00000065613 | SLK | 1.2185 | 0.1169 | 0.614638 | 1.123667845 | -0.11685623 | 0.6146381 |
| 852 | ENSG00000118363 | SPCS2 | 2.1117 | -0.114 | 0.627078 | 2.285290773 | 0.113994814 | 0.6270776 |
| 853 | ENSG00000175550 | DRAP1 | 2.6342 | 0.1132 | 0.627818 | 2.435376356 | -0.11323819 | 0.6278176 |
| 854 | ENSG00000071462 | BUD23 | 1.1102 | 0.1132 | 0.628255 | 1.026372429 | -0.11324218 | 0.6282552 |
| 855 | ENSG00000165502 | RPL36AL | 4.6618 | 0.1113 | 0.634827 | 4.315663368 | -0.1112914 | 0.6348271 |
| 856 | ENSG00000150093 | ITGB1 | 3.5478 | -0.1095 | 0.643496 | 3.827720723 | 0.10954904 | 0.6434962 |
| 857 | ENSG00000115457 | IGFBP2 | 8.3323 | 0.1089 | 0.644132 | 7.726386644 | -0.10892031 | 0.6441322 |
| 858 | ENSG00000101558 | VAPA | 2.5506 | -0.1093 | 0.644521 | 2.751249978 | 0.10926511 | 0.644521 |
| 859 | ENSG00000122042 | UBL3 | 1.5759 | 0.1086 | 0.645313 | 1.461584727 | -0.10861972 | 0.6453132 |
| 860 | ENSG00000166483 | WEE1 | 1.3611 | -0.1088 | 0.646978 | 1.467775981 | 0.108844813 | 0.6469779 |
| 861 | ENSG00000163605 | PPP4R2 | 1.2753 | 0.1061 | 0.654793 | 1.184892474 | -0.10606321 | 0.6547927 |
| 862 | ENSG00000173933 | RBM4 | 1.4617 | 0.1044 | 0.660772 | 1.359713166 | -0.10438416 | 0.6607717 |
| 863 | ENSG00000069849 | ATP1B3 | 1.382 | -0.1047 | 0.661871 | 1.486050651 | 0.10470383 | 0.6618713 |
| 864 | ENSG00000130724 | CHMP2A | 1.4116 | -0.1036 | 0.665447 | 1.516707829 | 0.10364094 | 0.6654474 |
| 865 | ENSG00000126247 | CAPNS1 | 1.06 | -0.1029 | 0.668665 | 1.138353382 | 0.102864925 | 0.6686649 |
| 866 | ENSG00000138069 | RAB1A | 1.7487 | -0.1023 | 0.670528 | 1.87717643 | 0.1022789 | 0.6705285 |
| 867 | ENSG00000177889 | UBE2N | 1.1534 | 0.1011 | 0.67339 | 1.075304277 | -0.10110365 | 0.6733905 |
| 868 | ENSG00000109787 | KLF3 | 1.3114 | 0.1011 | 0.673758 | 1.22269801 | -0.10105383 | 0.6737582 |
| 869 | ENSG00000099622 | CIRBP | 4.5453 | 0.1003 | 0.675792 | 4.240201845 | -0.10025332 | 0.6757918 |
| 870 | ENSG00000125868 | DSTN | 4.7799 | 0.0992 | 0.679461 | 4.462159818 | -0.09923707 | 0.6794612 |
| 871 | ENSG00000103994 | ZNF106 | 1.1571 | -0.0994 | 0.682043 | 1.239596845 | 0.099374402 | 0.6820432 |
| 872 | ENSG00000008294 | SPAG9 | 2.6207 | 0.0985 | 0.682672 | 2.447758865 | -0.09847172 | 0.6826717 |
| 873 | ENSG00000159377 | PSMB4 | 1.0809 | -0.099 | 0.683004 | 1.157644972 | 0.098972891 | 0.6830038 |
| 874 | ENSG00000149806 | FAU | 9.6755 | -0.0981 | 0.685665 | 10.35644356 | 0.098117454 | 0.6856654 |
| 875 | ENSG00000134717 | BTF3L4 | 1.1068 | -0.0963 | 0.693366 | 1.183187636 | 0.096260271 | 0.6933661 |
| 876 | ENSG00000134825 | TMEM258 | 1.1958 | -0.0959 | 0.694597 | 1.278030478 | 0.095922101 | 0.6945975 |
| 877 | ENSG00000175416 | CLTB | 2.2146 | 0.0939 | 0.700498 | 2.075087211 | -0.09386273 | 0.7004982 |
| 878 | ENSG00000118680 | MYL12B | 5.7617 | 0.0916 | 0.706949 | 5.40735802 | -0.0915728 | 0.7069487 |
| 879 | ENSG00000108848 | LUC7L3 | 2.1215 | 0.0903 | 0.709989 | 1.992716607 | -0.09033594 | 0.7099886 |
| 880 | ENSG00000137876 | RSL24D1 | 1.8779 | -0.0908 | 0.709989 | 1.999954692 | 0.090847419 | 0.7099886 |
| 881 | ENSG00000109475 | RPL34 | 13.892 | -0.0907 | 0.710146 | 14.7936589 | 0.090716301 | 0.7101463 |
| 882 | ENSG00000156467 | UQCRB | 2.0468 | -0.0905 | 0.711302 | 2.1792618 | 0.090458915 | 0.7113022 |
| 883 | ENSG00000148154 | UGCG | 1.254 | -0.0899 | 0.714079 | 1.334589234 | 0.089858865 | 0.7140789 |
| 884 | ENSG00000113558 | SKP1 | 5.828 | 0.0889 | 0.714964 | 5.479858508 | -0.08886787 | 0.7149643 |
| 885 | ENSG00000143158 | MPC2 | 1.3585 | 0.0886 | 0.716402 | 1.277522018 | -0.0886364 | 0.7164025 |
| 886 | ENSG00000126698 | DNAJC8 | 1.1703 | -0.0866 | 0.724329 | 1.242707427 | 0.086563562 | 0.7243293 |
| 887 | ENSG00000163220 | S100A9 | 4.7844 | -0.0906 | 0.72879 | 5.094505251 | 0.090619556 | 0.7287905 |
| 888 | ENSG00000162493 | PDPN | 1.0625 | -0.0854 | 0.729744 | 1.127316797 | 0.085440637 | 0.7297443 |
| 889 | ENSG00000185043 | CIB1 | 1.2312 | 0.0842 | 0.731711 | 1.161413562 | -0.08419907 | 0.7317112 |

| | | | | | | | | |
|---|---|---|---|---|---|---|---|---|
| 890 | ENSG00000167468 | GPX4 | 5.8063 | -0.0842 | 0.73289 | 6.155393177 | 0.084233799 | 0.7328903 |
| 891 | ENSG00000138085 | ATRAID | 1.2866 | 0.0835 | 0.734263 | 1.214233637 | -0.08347862 | 0.7342629 |
| 892 | ENSG00000179262 | RAD23A | 1.1061 | -0.0837 | 0.735285 | 1.172180961 | 0.083688488 | 0.7352855 |
| 893 | ENSG00000100353 | EIF3D | 1.0596 | 0.0825 | 0.738143 | 1.000740037 | -0.08249206 | 0.7381435 |
| 894 | ENSG00000112695 | COX7A2 | 2.4368 | 0.0812 | 0.742756 | 2.303415896 | -0.08120375 | 0.7427558 |
| 895 | ENSG00000183291 | SELENOF | 1.2655 | -0.0786 | 0.755062 | 1.3363838 | 0.078588498 | 0.7550615 |
| 896 | ENSG00000178127 | NDUFV2 | 2.5498 | 0.0768 | 0.759523 | 2.417610147 | -0.0768249 | 0.759523 |
| 897 | ENSG00000112081 | SRSF3 | 5.9042 | -0.0768 | 0.761311 | 6.227175839 | 0.076830951 | 0.761311 |
| 898 | ENSG00000139218 | SCAF11 | 1.3576 | -0.0756 | 0.766325 | 1.430658363 | 0.075636172 | 0.766325 |
| 899 | ENSG00000164032 | H2AFZ | 10.379 | -0.0753 | 0.766635 | 10.935281 | 0.075289668 | 0.7666345 |
| 900 | ENSG00000277791 | PSMB3 | 1.5272 | 0.0718 | 0.776025 | 1.453060536 | -0.07182116 | 0.7760249 |
| 901 | ENSG00000167470 | MIDN | 1.604 | -0.0707 | 0.782547 | 1.68458953 | 0.070694719 | 0.7825467 |
| 902 | ENSG00000118816 | CCNI | 4.4836 | -0.0701 | 0.784635 | 4.706729329 | 0.070057472 | 0.7846348 |
| 903 | ENSG00000132424 | PNISR | 2.1477 | -0.0694 | 0.78711 | 2.253616674 | 0.069449854 | 0.7871099 |
| 904 | ENSG00000143933 | CALM2 | 7.5675 | 0.0689 | 0.787197 | 7.214695902 | -0.06887359 | 0.7871971 |
| 905 | ENSG00000234741 | GAS5 | 5.3174 | 0.0682 | 0.789929 | 5.071863803 | -0.0682052 | 0.7899288 |
| 906 | ENSG00000075785 | RAB7A | 1.9081 | 0.0669 | 0.795422 | 1.821724324 | -0.06686691 | 0.7954218 |
| 907 | ENSG00000116539 | ASH1L | 1.332 | 0.0649 | 0.803855 | 1.273454333 | -0.06485096 | 0.8038553 |
| 908 | ENSG00000100811 | YY1 | 2.1255 | -0.0636 | 0.809863 | 2.221284566 | 0.063577905 | 0.8098634 |
| 909 | ENSG00000168710 | AHCYL1 | 1.1635 | 0.0603 | 0.820123 | 1.11589139 | -0.06027664 | 0.8201226 |
| 910 | ENSG00000134108 | ARL8B | 1.0601 | 0.0592 | 0.824565 | 1.017489325 | -0.05915189 | 0.8245646 |
| 911 | ENSG00000117394 | SLC2A1 | 2.2323 | -0.0582 | 0.830185 | 2.324143139 | 0.058178189 | 0.8301846 |
| 912 | ENSG00000147677 | EIF3H | 2.2708 | 0.0576 | 0.830195 | 2.18195365 | -0.05759593 | 0.8301953 |
| 913 | ENSG00000133226 | SRRM1 | 1.9598 | -0.0574 | 0.832443 | 2.039285609 | 0.057370086 | 0.8324427 |
| 914 | ENSG00000018408 | WWTR1 | 1.1399 | -0.0569 | 0.834579 | 1.185819667 | 0.056943039 | 0.8345789 |
| 915 | ENSG00000278311 | GGNBP2 | 1.1845 | 0.056 | 0.836415 | 1.139400212 | -0.05597094 | 0.836415 |
| 916 | ENSG00000181222 | POLR2A | 1.1495 | -0.0559 | 0.838346 | 1.194912137 | 0.055869064 | 0.8383464 |
| 917 | ENSG00000124831 | LRRFIP1 | 2.1914 | -0.0553 | 0.840523 | 2.276975948 | 0.055272859 | 0.8405227 |
| 918 | ENSG00000084207 | GSTP1 | 14.279 | 0.05 | 0.859099 | 13.7923805 | -0.04999456 | 0.8590989 |
| 919 | ENSG00000111897 | SERINC1 | 1.5085 | -0.049 | 0.865294 | 1.5606448 | 0.048982161 | 0.8652942 |
| 920 | ENSG00000150787 | PTS | 1.066 | 0.0482 | 0.866266 | 1.031008393 | -0.04817377 | 0.8662658 |
| 921 | ENSG00000166200 | COPS2 | 1.1115 | 0.0468 | 0.871569 | 1.075962285 | -0.04682161 | 0.8715691 |
| 922 | ENSG00000142669 | SH3BGRL3 | 5.6771 | 0.046 | 0.874498 | 5.498940733 | -0.04599863 | 0.874498 |
| 923 | ENSG00000113575 | PPP2CA | 1.3432 | -0.046 | 0.876913 | 1.386661573 | 0.045951414 | 0.8769132 |
| 924 | ENSG00000213853 | EMP2 | 1.3205 | 0.0433 | 0.885274 | 1.281470064 | -0.04332114 | 0.8852742 |
| 925 | ENSG00000140264 | SERF2 | 6.5384 | 0.0422 | 0.889172 | 6.349714825 | -0.04224213 | 0.8891715 |
| 926 | ENSG00000131174 | COX7B | 1.6403 | 0.0407 | 0.895712 | 1.594651837 | -0.04067644 | 0.8957124 |
| 927 | ENSG00000101361 | NOP56 | 1.051 | 0.0405 | 0.896649 | 1.021856103 | -0.04050943 | 0.8966485 |
| 928 | ENSG00000125968 | ID1 | 15.301 | 0.0392 | 0.901701 | 14.89041595 | -0.03921264 | 0.9017012 |
| 929 | ENSG00000119326 | CTNNAL1 | 1.3499 | 0.038 | 0.90695 | 1.314759274 | -0.03801309 | 0.90695 |
| 930 | ENSG00000162664 | ZNF326 | 1.3317 | -0.0384 | 0.907099 | 1.367639168 | 0.03843132 | 0.9070995 |
| 931 | ENSG00000111832 | RWDD1 | 1.185 | -0.0384 | 0.907129 | 1.216955397 | 0.038409274 | 0.9071286 |
| 932 | ENSG00000136754 | ABI1 | 1.1482 | 0.0363 | 0.914164 | 1.11965998 | -0.03625661 | 0.9141639 |
| 933 | ENSG00000185896 | LAMP1 | 1.2053 | -0.0366 | 0.914647 | 1.236187168 | 0.036557727 | 0.9146472 |
| 934 | ENSG00000204435 | CSNK2B | 1.0748 | -0.0364 | 0.915118 | 1.102312503 | 0.03643826 | 0.9151181 |
| 935 | ENSG00000111669 | TPI1 | 3.9639 | -0.0358 | 0.917282 | 4.063586587 | 0.035825847 | 0.9172819 |
| 936 | ENSG00000086065 | CHMP5 | 1.4225 | 0.0344 | 0.921092 | 1.38899451 | -0.03438706 | 0.9210916 |

| | | | | | | | | |
|---|---|---|---|---|---|---|---|---|
| 937 | ENSG00000110958 | PTGES3 | 3.9113 | -0.0346 | 0.921998 | 4.006190367 | 0.034570838 | 0.9219979 |
| 938 | ENSG00000122482 | ZNF644 | 1.0494 | 0.0331 | 0.926231 | 1.025564874 | -0.03314403 | 0.9262308 |
| 939 | ENSG00000139644 | TMBIM6 | 3.6688 | -0.0314 | 0.934881 | 3.749507531 | 0.031382492 | 0.9348806 |
| 940 | ENSG00000143977 | SNRPG | 1.6697 | 0.0296 | 0.940438 | 1.635807229 | -0.02956567 | 0.9404378 |
| 941 | ENSG00000072364 | AFF4 | 1.3709 | -0.0296 | 0.942464 | 1.399313268 | 0.029597505 | 0.942464 |
| 942 | ENSG00000005022 | SLC25A5 | 2.6778 | 0.0287 | 0.944051 | 2.625002221 | -0.02871145 | 0.944051 |
| 943 | ENSG00000175602 | CCDC85B | 1.2735 | -0.0289 | 0.945179 | 1.299296092 | 0.028922008 | 0.9451791 |
| 944 | ENSG00000079332 | SAR1A | 1.3354 | -0.0272 | 0.95222 | 1.360759996 | 0.027168935 | 0.9522196 |
| 945 | ENSG00000169813 | HNRNPF | 2.2573 | -0.0264 | 0.955305 | 2.298929479 | 0.026390467 | 0.9553046 |
| 946 | ENSG00000137409 | MTCH1 | 1.5582 | -0.0257 | 0.958101 | 1.586277193 | 0.025724768 | 0.9581008 |
| 947 | ENSG00000134884 | ARGLU1 | 2.1583 | 0.0225 | 0.969578 | 2.124946252 | -0.02247618 | 0.9695783 |
| 948 | ENSG00000120306 | CYSTM1 | 1.1504 | 0.0208 | 0.976456 | 1.133926784 | -0.02082219 | 0.9764561 |
| 949 | ENSG00000065809 | FAM107B | 2.6456 | 0.0204 | 0.977672 | 2.60843239 | -0.02043489 | 0.9776719 |
| 950 | ENSG00000165678 | GHITM | 1.2084 | -0.0209 | 0.977672 | 1.226017958 | 0.020918434 | 0.9776719 |
| 951 | ENSG00000159199 | ATP5MC1 | 1.0319 | 0.0199 | 0.980088 | 1.017818328 | -0.01985483 | 0.9800876 |
| 952 | ENSG00000123416 | TUBA1B | 2.8012 | -0.0188 | 0.985978 | 2.837957452 | 0.018834103 | 0.9859781 |
| 953 | ENSG00000102580 | DNAJC3 | 1.0565 | 0.0167 | 0.993168 | 1.044288185 | -0.01670867 | 0.9931676 |
| 954 | ENSG00000164111 | ANXA5 | 5.8401 | 0.0011 | 1 | 5.835601417 | -0.00110365 | 1 |
| 955 | ENSG00000145741 | BTF3 | 9.5807 | -0.0054 | 1 | 9.616334407 | 0.005365181 | 1 |
| 956 | ENSG00000082153 | BZW1 | 5.1724 | 0.0096 | 1 | 5.137933762 | -0.00963152 | 1 |
| 957 | ENSG00000198668 | CALM1 | 6.7368 | -0.0125 | 1 | 6.795335608 | 0.012490984 | 1 |
| 958 | ENSG00000113712 | CSNK1A1 | 3.7392 | 0.0021 | 1 | 3.733834983 | -0.00208645 | 1 |
| 959 | ENSG00000160213 | CSTB | 2.9503 | -0.0102 | 1 | 2.971204018 | 0.010201352 | 1 |
| 960 | ENSG00000114784 | EIF1B | 3.4171 | 0.0001 | 1 | 3.416854712 | -0.0001057 | 1 |
| 961 | ENSG00000103363 | ELOB | 2.5631 | -0.005 | 1 | 2.57194287 | 0.004973608 | 1 |
| 962 | ENSG00000111640 | GAPDH | 18.936 | -0.0072 | 1 | 19.0308399 | 0.00723243 | 1 |
| 963 | ENSG00000112972 | HMGCS1 | 1.3969 | 0.01 | 1 | 1.387170034 | -0.01004021 | 1 |
| 964 | ENSG00000153187 | HNRNPU | 4.3755 | 0.0035 | 1 | 4.364984039 | -0.00347123 | 1 |
| 965 | ENSG00000196549 | MME | 2.1607 | 0.0095 | 1 | 2.14654087 | -0.00945994 | 1 |
| 966 | ENSG00000163110 | PDLIM5 | 1.186 | -0.0033 | 1 | 1.188661064 | 0.003272511 | 1 |
| 967 | ENSG00000147669 | POLR2K | 1.2681 | -0.0014 | 1 | 1.26935674 | 0.001386215 | 1 |
| 968 | ENSG00000125995 | ROMO1 | 1.1601 | 0.0016 | 1 | 1.15884135 | -0.00160933 | 1 |
| 969 | ENSG00000112335 | SNX3 | 2.1067 | 0.0041 | 1 | 2.100779423 | -0.00408785 | 1 |
| 970 | ENSG00000161547 | SRSF2 | 3.7764 | -0.0103 | 1 | 3.803464165 | 0.010308566 | 1 |
| 971 | ENSG00000198492 | YTHDF2 | 1.1374 | -0.0012 | 1 | 1.138353382 | 0.001188773 | 1 |
