## Supplementary material for "Single cell RNA-seq identifies developing corneal cell fates in the human cornea organoid": Table S4

| FeatureID | FeatureName | Cornea Average | Cornea Log2 Fold Change | Cornea P-Value | Organoid Average | Organoid Log2 Fold Change | Organoid P-Value |
|---|---|---|---|---|---|---|---|
| ENSG00000179163 | FUCA1 | 0.020724 | -6.46054 | 4.88E-40 | 2.0115537 | 6.460543651 | 4.88E-40 |
| ENSG00000205542 | TMSB4X | 10.181708 | -3.73602 | 2.34E-36 | 136.49599 | 3.736023383 | 2.34E-36 |
| ENSG00000204287 | HLA-DRA | 19.40804 | 5.4119 | 7.17E-28 | 0.450588 | -5.41189559 | 7.17E-28 |
| ENSG00000196735 | HLA-DQA1 | 5.0918904 | 6.72987 | 7.17E-28 | 0.0402311 | -6.72987498 | 7.17E-28 |
| ENSG00000068366 | ACSL4 | 0.3502359 | -3.61318 | 5.03E-25 | 4.3288636 | 3.613182242 | 5.03E-25 |
| ENSG00000126709 | IFI6 | 0.1450681 | -4.21654 | 2.39E-24 | 2.7437593 | 4.216537111 | 2.39E-24 |
| ENSG00000135111 | TBX3 | 0.0145068 | -6.11836 | 2.21E-23 | 1.1506087 | 6.118356717 | 2.21E-23 |
| ENSG00000168542 | COL3A1 | 0.0041448 | -8.31184 | 2.48E-23 | 1.9793689 | 8.311836446 | 2.48E-23 |
| ENSG00000196126 | HLA-DRB1 | 11.302878 | 4.87997 | 4.79E-23 | 0.3781721 | -4.87996927 | 4.79E-23 |
| ENSG00000204389 | HSPA1A | 17.511793 | 5.70416 | 3.96E-22 | 0.3298948 | -5.70415721 | 3.96E-22 |
| ENSG00000146070 | PLA2G7 | 0.0808237 | -4.08594 | 5.63E-22 | 1.4080876 | 4.085935239 | 5.63E-22 |
| ENSG00000198502 | HLA-DRB5 | 2.4661578 | 5.9476 | 5.82E-22 | 0.0321849 | -5.94759789 | 5.82E-22 |
| ENSG00000099860 | GADD45B | 8.0347005 | 4.8852 | 5.42E-20 | 0.2655251 | -4.88520253 | 5.42E-20 |
| ENSG00000129538 | RNASE1 | 0.8248158 | -3.15211 | 2.13E-19 | 7.3864253 | 3.152107831 | 2.13E-19 |
| ENSG00000271614 | ATP2B1-AS1 | 4.1530926 | 4.69903 | 4.30E-19 | 0.1528781 | -4.69902671 | 4.30E-19 |
| ENSG00000204388 | HSPA1B | 8.1175966 | 5.90001 | 4.46E-19 | 0.1287394 | -5.90000713 | 4.46E-19 |
| ENSG00000067048 | DDX3Y | 1.2185721 | 7.25369 | 7.38E-19 | 0 | -7.25369211 | 7.38E-19 |
| ENSG00000223865 | HLA-DPB1 | 4.6048761 | 4.84793 | 1.96E-18 | 0.1528781 | -4.84793242 | 1.96E-18 |
| ENSG00000102172 | SMS | 0.3647427 | -2.96867 | 2.70E-18 | 2.8805449 | 2.968666199 | 2.70E-18 |
| ENSG00000232810 | TNF | 15.225934 | 4.6093 | 4.45E-18 | 0.6195585 | -4.60930198 | 4.45E-18 |
| ENSG00000165168 | CYBB | 0.2072401 | -3.37237 | 7.21E-18 | 2.172478 | 3.372369274 | 7.21E-18 |
| ENSG00000118515 | SGK1 | 2.9718237 | 5.21648 | 7.21E-18 | 0.0724159 | -5.21647521 | 7.21E-18 |
| ENSG00000231389 | HLA-DPA1 | 5.1623521 | 4.47668 | 1.10E-17 | 0.225294 | -4.47667564 | 1.10E-17 |
| ENSG00000170345 | FOS | 9.0978425 | 4.94245 | 4.56E-17 | 0.2896637 | -4.94244874 | 4.56E-17 |
| ENSG00000160888 | IER2 | 6.0762811 | 4.2478 | 2.01E-16 | 0.3138024 | -4.24780155 | 2.01E-16 |
| ENSG00000198695 | MT-ND6 | 0.1119097 | -3.22235 | 2.41E-16 | 1.0621004 | 3.222354438 | 2.41E-16 |
| ENSG00000226380 | AC016831.1 | 2.7832352 | 4.12196 | 5.04E-16 | 0.1528781 | -4.12195761 | 5.04E-16 |
| ENSG00000163220 | S100A9 | 1.4568982 | -3.1083 | 5.19E-16 | 12.64865 | 3.108303052 | 5.19E-16 |
| ENSG00000176788 | BASP1 | 5.3447234 | 3.89287 | 5.78E-16 | 0.3540335 | -3.89287054 | 5.78E-16 |
| ENSG00000125740 | FOSB | 2.1656595 | 4.76031 | 1.59E-15 | 0.0724159 | -4.76030733 | 1.59E-15 |
| ENSG00000019582 | CD74 | 8.5921765 | 3.53026 | 7.25E-15 | 0.7402518 | -3.53026181 | 7.25E-15 |
| ENSG00000132002 | DNAJB1 | 6.6814224 | 4.61919 | 1.65E-14 | 0.2655251 | -4.61918889 | 1.65E-14 |
| ENSG00000175445 | LPL | 0.1119097 | -3.15576 | 1.67E-14 | 1.0138231 | 3.155756689 | 1.67E-14 |
| ENSG00000152558 | TMEM123 | 0.4580007 | -2.54203 | 1.76E-14 | 2.6874358 | 2.542033135 | 1.76E-14 |
| ENSG00000196743 | GM2A | 0.1305613 | -3.31475 | 2.34E-14 | 1.3195792 | 3.31475393 | 2.34E-14 |
| ENSG00000163735 | CXCL5 | 1.4154502 | -2.96937 | 3.11E-14 | 11.1601 | 2.969371053 | 3.11E-14 |
| ENSG00000138061 | CYP1B1 | 0.8641914 | -2.65258 | 3.79E-14 | 5.4714261 | 2.652583571 | 3.79E-14 |
| ENSG00000198938 | MT-CO3 | 10.432469 | -2.22997 | 4.99E-14 | 49.234789 | 2.22997159 | 4.99E-14 |
| ENSG00000198804 | MT-CO1 | 15.802061 | -2.21458 | 5.01E-14 | 73.78379 | 2.214584237 | 5.01E-14 |
| ENSG00000130429 | ARPC1B | 0.3937563 | -2.54593 | 6.07E-14 | 2.3173099 | 2.54592857 | 6.07E-14 |
| ENSG00000120738 | EGR1 | 5.1892933 | 4.88273 | 8.41E-14 | 0.1689705 | -4.88273155 | 8.41E-14 |
| ENSG00000109321 | AREG | 5.4110403 | 4.87895 | 9.25E-14 | 0.1770167 | -4.87894551 | 9.25E-14 |
| ENSG00000234883 | MIR155HG | 5.3177822 | 3.59608 | 1.16E-13 | 0.4344956 | -3.59607615 | 1.16E-13 |

| | | | | | | | |
|---|---|---|---|---|---|---|---|
| ENSG00000136826 | KLF4 | 1.2496581 | 4.70501 | 1.16E-13 | 0.0402311 | -4.70501052 | 1.16E-13 |
| ENSG00000042753 | AP2S1 | 0.3481634 | -2.72248 | 1.35E-13 | 2.3173099 | 2.722477962 | 1.35E-13 |
| ENSG00000151012 | SLC7A11 | 0.8849154 | -2.40909 | 2.02E-13 | 4.7311744 | 2.409088553 | 2.02E-13 |
| ENSG00000118785 | SPP1 | 1.7988445 | -3.18099 | 2.70E-13 | 16.422325 | 3.1809865 | 2.70E-13 |
| ENSG00000198886 | MT-ND4 | 7.1456403 | -2.18267 | 2.75E-13 | 32.635448 | 2.182666889 | 2.75E-13 |
| ENSG00000102575 | ACP5 | 0.0269412 | -6.2496 | 3.85E-13 | 2.2127091 | 6.24960125 | 3.85E-13 |
| ENSG00000198899 | MT-ATP6 | 9.761011 | -2.15202 | 5.43E-13 | 43.64267 | 2.152021521 | 5.43E-13 |
| ENSG00000198712 | MT-CO2 | 14.243615 | -2.10876 | 7.13E-13 | 61.802977 | 2.108763828 | 7.13E-13 |
| ENSG00000110852 | CLEC2B | 1.5501563 | 4.60039 | 1.14E-12 | 0.0563235 | -4.60039019 | 1.14E-12 |
| ENSG00000173110 | HSPA6 | 5.8089413 | 6.50484 | 1.29E-12 | 0.0563235 | -6.50483892 | 1.29E-12 |
| ENSG00000150938 | CRIM1 | 0.1865161 | -2.49592 | 3.92E-12 | 1.0621004 | 2.495919511 | 3.92E-12 |
| ENSG00000260708 | AL118516.1 | 1.3221921 | 4.37124 | 4.14E-12 | 0.0563235 | -4.37124041 | 4.14E-12 |
| ENSG00000090104 | RGS1 | 2.828828 | 3.82345 | 4.85E-12 | 0.1931092 | -3.82345386 | 4.85E-12 |
| ENSG00000178607 | ERN1 | 1.2496581 | 4.28997 | 4.96E-12 | 0.0563235 | -4.28997302 | 4.96E-12 |
| ENSG00000160307 | S100B | 0.0331584 | -4.90536 | 5.22E-12 | 1.0540542 | 4.905362994 | 5.22E-12 |
| ENSG00000120694 | HSPH1 | 5.9395026 | 4.11063 | 5.46E-12 | 0.337941 | -4.11062972 | 5.46E-12 |
| ENSG00000141682 | PMAIP1 | 5.0048496 | 3.76641 | 5.59E-12 | 0.3620797 | -3.76641107 | 5.59E-12 |
| ENSG00000198786 | MT-ND5 | 2.756294 | -2.10054 | 8.73E-12 | 11.892306 | 2.100543197 | 8.73E-12 |
| ENSG00000140403 | DNAJA4 | 1.1688344 | 4.60871 | 1.10E-11 | 0.0402311 | -4.60871284 | 1.10E-11 |
| ENSG00000105223 | PLD3 | 0.2569778 | -2.49044 | 1.27E-11 | 1.4563649 | 2.490442071 | 1.27E-11 |
| ENSG00000171988 | JMJD1C | 2.5946467 | 3.38854 | 1.27E-11 | 0.2413864 | -3.38854267 | 1.27E-11 |
| ENSG00000120129 | DUSP1 | 4.3105951 | 3.43077 | 2.01E-11 | 0.3942645 | -3.43077334 | 2.01E-11 |
| ENSG00000089351 | GRAMD1A | 0.1844437 | -2.5651 | 2.17E-11 | 1.1023314 | 2.565103076 | 2.17E-11 |
| ENSG00000184205 | TSPYL2 | 1.2413685 | 3.82096 | 3.11E-11 | 0.0804621 | -3.82095536 | 3.11E-11 |
| ENSG00000248323 | LUCAT1 | 6.762246 | 3.12408 | 3.57E-11 | 0.7724366 | -3.12408078 | 3.57E-11 |
| ENSG00000198840 | MT-ND3 | 5.0773836 | -1.97814 | 5.36E-11 | 20.123583 | 1.978143021 | 5.36E-11 |
| ENSG00000117984 | CTSD | 0.3647427 | -2.96867 | 5.48E-11 | 2.8805449 | 2.968666199 | 5.48E-11 |
| ENSG00000212907 | MT-ND4L | 0.8641914 | -2.15396 | 6.32E-11 | 3.8702294 | 2.15396192 | 6.32E-11 |
| ENSG00000112137 | PHACTR1 | 1.3387714 | 3.4823 | 6.33E-11 | 0.112647 | -3.48229959 | 6.33E-11 |
| ENSG00000128016 | ZFP36 | 3.0795886 | 3.34183 | 6.57E-11 | 0.29771 | -3.3418297 | 6.57E-11 |
| ENSG00000179344 | HLA-DQB1 | 1.9024646 | 3.80803 | 7.69E-11 | 0.1287394 | -3.80802649 | 7.69E-11 |
| ENSG00000030582 | GRN | 0.2238194 | -2.78758 | 1.10E-10 | 1.5609657 | 2.787577705 | 1.10E-10 |
| ENSG00000096384 | HSP90AB1 | 18.952112 | 2.75517 | 1.29E-10 | 2.8161752 | -2.75516627 | 1.29E-10 |
| ENSG00000198888 | MT-ND1 | 5.8980546 | -1.92339 | 2.28E-10 | 22.505263 | 1.923385185 | 2.28E-10 |
| ENSG00000015475 | BID | 0.495304 | -2.21147 | 2.33E-10 | 2.3092637 | 2.211466121 | 2.33E-10 |
| ENSG00000188229 | TUBB4B | 2.1179943 | 2.9627 | 4.24E-10 | 0.2655251 | -2.96269587 | 4.24E-10 |
| ENSG00000137801 | THBS1 | 2.7998144 | -2.1372 | 4.90E-10 | 12.391171 | 2.137203426 | 4.90E-10 |
| ENSG00000182287 | AP1S2 | 0.4352043 | -2.18153 | 6.35E-10 | 1.9874151 | 2.181528837 | 6.35E-10 |
| ENSG00000151929 | BAG3 | 1.4278846 | 4.02259 | 6.41E-10 | 0.0804621 | -4.02258922 | 6.41E-10 |
| ENSG00000116717 | GADD45A | 1.7594689 | 3.46096 | 7.18E-10 | 0.1528781 | -3.46095922 | 7.18E-10 |
| ENSG00000116741 | RGS2 | 1.935623 | 3.46096 | 8.70E-10 | 0.1689705 | -3.46095922 | 8.70E-10 |
| ENSG00000179630 | LACC1 | 0.1844437 | -2.45658 | 1.13E-09 | 1.0218693 | 2.456578619 | 1.13E-09 |
| ENSG00000143545 | RAB13 | 0.3378014 | -2.23474 | 1.40E-09 | 1.6011968 | 2.234735901 | 1.40E-09 |
| ENSG00000181649 | PHLDA2 | 4.2028302 | 2.77133 | 1.52E-09 | 0.6115123 | -2.77133489 | 1.52E-09 |
| ENSG00000104714 | ERICH1 | 2.5366194 | 4.60969 | 1.58E-09 | 0.0965546 | -4.6096946 | 1.58E-09 |
| ENSG00000106211 | HSPB1 | 4.4411564 | 3.98837 | 1.84E-09 | 0.2735713 | -3.98837446 | 1.84E-09 |
| ENSG00000081041 | CXCL2 | 41.814845 | 2.57399 | 2.34E-09 | 7.0565305 | -2.57398672 | 2.34E-09 |

| | | | | | | | |
|---|---|---|---|---|---|---|---|
| ENSG00000143387 | CTSK | 0.0393756 | -5.6265 | 2.36E-09 | 2.0517848 | 5.626503621 | 2.36E-09 |
| ENSG00000104870 | FCGRT | 0.325367 | -2.2957 | 2.40E-09 | 1.609243 | 2.295702659 | 2.40E-09 |
| ENSG00000087074 | PPP1R15A | 11.476959 | 2.38895 | 2.80E-09 | 2.1966167 | -2.38894603 | 2.80E-09 |
| ENSG00000158050 | DUSP2 | 4.4867492 | 3.67295 | 2.82E-09 | 0.3459872 | -3.67295419 | 2.82E-09 |
| ENSG00000080824 | HSP90AA1 | 30.605225 | 2.78291 | 2.86E-09 | 4.4656493 | -2.78291174 | 2.86E-09 |
| ENSG00000163739 | CXCL1 | 2.4267821 | -2.24553 | 3.01E-09 | 11.578503 | 2.245527957 | 3.01E-09 |
| ENSG00000066336 | SPI1 | 0.2984258 | -2.17791 | 3.35E-09 | 1.3598103 | 2.177913562 | 3.35E-09 |
| ENSG00000162711 | NLRP3 | 1.3449886 | 3.14794 | 3.40E-09 | 0.1448319 | -3.14793668 | 3.40E-09 |
| ENSG00000151247 | EIF4E | 2.7438596 | 2.61599 | 3.94E-09 | 0.4425418 | -2.61599001 | 3.94E-09 |
| ENSG00000182220 | ATP6AP2 | 0.6072136 | -1.96308 | 4.19E-09 | 2.3816796 | 1.963078492 | 4.19E-09 |
| ENSG00000101966 | XIAP | 0.2590502 | -2.10043 | 4.25E-09 | 1.1184239 | 2.100434809 | 4.25E-09 |
| ENSG00000258667 | HIF1A-AS3 | 1.3263369 | 3.91632 | 5.05E-09 | 0.0804621 | -3.91631721 | 5.05E-09 |
| ENSG00000162645 | GBP2 | 1.0672868 | 3.0628 | 5.23E-09 | 0.1206932 | -3.06279554 | 5.23E-09 |
| ENSG00000075624 | ACTB | 5.3260718 | -1.83646 | 5.74E-09 | 19.133899 | 1.836457377 | 5.74E-09 |
| ENSG00000067082 | KLF6 | 7.4668625 | 2.55407 | 6.51E-09 | 1.271302 | -2.55406863 | 6.51E-09 |
| ENSG00000197632 | SERPINB2 | 4.6691205 | 4.66628 | 8.55E-09 | 0.1770167 | -4.66627813 | 8.55E-09 |
| ENSG00000185650 | ZFP36L1 | 2.8992897 | 2.94819 | 1.05E-08 | 0.3701259 | -2.94819055 | 1.05E-08 |
| ENSG00000130222 | GADD45G | 1.8693061 | 3.87015 | 1.20E-08 | 0.1206932 | -3.87015046 | 1.20E-08 |
| ENSG00000124882 | EREG | 10.167202 | 2.72723 | 1.42E-08 | 1.536827 | -2.72723134 | 1.42E-08 |
| ENSG00000151726 | ACSL1 | 0.4807971 | -2.06724 | 1.44E-08 | 2.0276462 | 2.067239146 | 1.44E-08 |
| ENSG00000115541 | HSPE1 | 3.7987119 | 2.72242 | 1.51E-08 | 0.5712813 | -2.72242121 | 1.51E-08 |
| ENSG00000254999 | BRK1 | 0.3067154 | -2.05119 | 1.56E-08 | 1.2793482 | 2.05119129 | 1.56E-08 |
| ENSG00000166592 | RRAD | 1.1709068 | 3.8743 | 2.12E-08 | 0.0724159 | -3.87429843 | 2.12E-08 |
| ENSG00000253320 | AZIN1-AS1 | 2.3397413 | 2.83612 | 2.33E-08 | 0.3218486 | -2.83612334 | 2.33E-08 |
| ENSG00000143546 | S100A8 | 0.5595484 | -2.45125 | 2.37E-08 | 3.0817003 | 2.451245175 | 2.37E-08 |
| ENSG00000197461 | PDGFA | 1.398871 | 4.13052 | 2.91E-08 | 0.0724159 | -4.13051963 | 2.91E-08 |
| ENSG00000136244 | IL6 | 4.8763607 | 3.44511 | 3.02E-08 | 0.4425418 | -3.44511197 | 3.02E-08 |
| ENSG00000137710 | RDX | 0.4206975 | -1.97645 | 3.05E-08 | 1.6655665 | 1.976446092 | 3.05E-08 |
| ENSG00000157227 | MMP14 | 0.242471 | -2.12107 | 3.63E-08 | 1.0621004 | 2.121071102 | 3.63E-08 |
| ENSG00000163466 | ARPC2 | 1.5232151 | -1.74228 | 3.66E-08 | 5.1254389 | 1.742282373 | 3.66E-08 |
| ENSG00000175197 | DDIT3 | 1.3056129 | 2.70921 | 4.48E-08 | 0.1931092 | -2.70920829 | 4.48E-08 |
| ENSG00000138821 | SLC39A8 | 0.3564531 | -2.10626 | 4.95E-08 | 1.5448733 | 2.106260525 | 4.95E-08 |
| ENSG00000153234 | NR4A2 | 2.1739492 | 3.18085 | 5.21E-08 | 0.2333402 | -3.1808513 | 5.21E-08 |
| ENSG00000136634 | IL10 | 2.4371441 | 3.34558 | 5.49E-08 | 0.2333402 | -3.34557629 | 5.49E-08 |
| ENSG00000175352 | NRIP3 | 1.4942015 | 2.64053 | 6.40E-08 | 0.2333402 | -2.64053272 | 6.40E-08 |
| ENSG00000260231 | KDM7A-DT | 1.1812688 | 2.96099 | 7.67E-08 | 0.1448319 | -2.96098771 | 7.67E-08 |
| ENSG00000269893 | SNHG8 | 1.5563735 | 3.02119 | 8.78E-08 | 0.1850629 | -3.02119464 | 8.78E-08 |
| ENSG00000269821 | KCNQ1OT1 | 2.3770445 | 3.08719 | 1.07E-07 | 0.2735713 | -3.08719219 | 1.07E-07 |
| ENSG00000115875 | SRSF7 | 1.6289076 | 2.46233 | 1.39E-07 | 0.2896637 | -2.46233474 | 1.39E-07 |
| ENSG00000100097 | LGALS1 | 2.6236603 | -1.67328 | 1.68E-07 | 8.4163408 | 1.673276633 | 1.68E-07 |
| ENSG00000198431 | TXNRD1 | 1.1999205 | -1.80195 | 1.70E-07 | 4.2081704 | 1.801945627 | 1.70E-07 |
| ENSG00000102786 | INTS6 | 1.0051147 | 2.72845 | 2.73E-07 | 0.1448319 | -2.72845328 | 2.73E-07 |
| ENSG00000171223 | JUNB | 3.7158159 | 2.35274 | 3.36E-07 | 0.7241593 | -2.35273782 | 3.36E-07 |
| ENSG00000139278 | GLIPR1 | 2.5739226 | 2.49829 | 3.36E-07 | 0.450588 | -2.49828885 | 3.36E-07 |
| ENSG00000108602 | ALDH3A1 | 2.0972703 | 5.03599 | 3.56E-07 | 0.0563235 | -5.03598674 | 3.56E-07 |
| ENSG00000125347 | IRF1 | 1.1688344 | 2.67011 | 4.31E-07 | 0.1770167 | -2.67011339 | 4.31E-07 |
| ENSG00000115956 | PLEK | 4.9924152 | 2.13664 | 4.38E-07 | 1.1345163 | -2.1366386 | 4.38E-07 |

| | | | | | | | |
|---|---|---|---|---|---|---|---|
| ENSG00000092820 | EZR | 1.997795 | 2.41136 | 5.61E-07 | 0.3701259 | -2.41136457 | 5.61E-07 |
| ENSG00000111678 | C12orf57 | 1.3263369 | 3.12782 | 6.76E-07 | 0.1448319 | -3.12782132 | 6.76E-07 |
| ENSG00000152229 | PSTPIP2 | 0.3067154 | -1.93872 | 7.60E-07 | 1.1827936 | 1.938716561 | 7.60E-07 |
| ENSG00000169429 | CXCL8 | 270.52922 | 1.95791 | 7.60E-07 | 70.042301 | -1.9579064 | 7.60E-07 |
| ENSG00000112110 | MRPL18 | 1.2890337 | 2.63422 | 7.62E-07 | 0.2011554 | -2.63421692 | 7.62E-07 |
| ENSG00000104763 | ASAH1 | 0.3916839 | -1.94082 | 8.25E-07 | 1.5126884 | 1.940818531 | 8.25E-07 |
| ENSG00000198805 | PNP | 1.0589972 | 2.59214 | 8.48E-07 | 0.1689705 | -2.59213667 | 8.48E-07 |
| ENSG00000162772 | ATF3 | 1.5770975 | 2.53775 | 8.91E-07 | 0.2655251 | -2.53775263 | 8.91E-07 |
| ENSG00000164687 | FABP5 | 2.2464832 | -1.75522 | 1.03E-06 | 7.6278117 | 1.755216665 | 1.03E-06 |
| ENSG00000198763 | MT-ND2 | 7.8585464 | -1.54363 | 1.05E-06 | 23.044359 | 1.543627483 | 1.05E-06 |
| ENSG00000204569 | PPP1R10 | 1.5770975 | 2.26766 | 1.06E-06 | 0.3218486 | -2.26766347 | 1.06E-06 |
| ENSG00000140379 | BCL2A1 | 13.414655 | 2.03958 | 1.12E-06 | 3.2748095 | -2.03958459 | 1.12E-06 |
| ENSG00000100906 | NFKBIA | 24.162129 | 2.02169 | 1.13E-06 | 5.9783377 | -2.02168964 | 1.13E-06 |
| ENSG00000107317 | PTGDS | 11.043827 | 4.8769 | 1.22E-06 | 0.3701259 | -4.87689925 | 1.22E-06 |
| ENSG00000197405 | C5AR1 | 1.6392696 | 2.39552 | 1.24E-06 | 0.3057562 | -2.39552269 | 1.24E-06 |
| ENSG00000267519 | AC020916.1 | 1.0983728 | 2.51917 | 1.44E-06 | 0.1850629 | -2.51917384 | 1.44E-06 |
| ENSG00000105939 | ZC3HAV1 | 1.0092595 | 3.17495 | 1.55E-06 | 0.1046008 | -3.1749507 | 1.55E-06 |
| ENSG00000144802 | NFKBIZ | 3.680585 | 2.05238 | 2.02E-06 | 0.8850836 | -2.05238038 | 2.02E-06 |
| ENSG00000104765 | BNIP3L | 0.5947792 | -1.86066 | 2.24E-06 | 2.172478 | 1.860655755 | 2.24E-06 |
| ENSG00000078401 | EDN1 | 1.0589972 | 7.05157 | 2.49E-06 | 0 | -7.05156828 | 2.49E-06 |
| ENSG00000145901 | TNIP1 | 0.6382997 | -1.60178 | 3.08E-06 | 1.947184 | 1.601781191 | 3.08E-06 |
| ENSG00000044574 | HSPA5 | 5.1851485 | 2.21173 | 3.29E-06 | 1.1184239 | -2.21172784 | 3.29E-06 |
| ENSG00000141543 | EIF4A3 | 1.429957 | 2.27466 | 3.38E-06 | 0.2896637 | -2.27465682 | 3.38E-06 |
| ENSG00000177606 | JUN | 7.6575235 | 2.58139 | 3.87E-06 | 1.2793482 | -2.58138923 | 3.87E-06 |
| ENSG00000159335 | PTMS | 0.294281 | -1.85465 | 4.20E-06 | 1.0701466 | 1.854649569 | 4.20E-06 |
| ENSG00000128524 | ATP6V1F | 1.5750251 | -1.49042 | 4.24E-06 | 4.4495568 | 1.490421238 | 4.24E-06 |
| ENSG00000109971 | HSPA8 | 6.0845707 | 2.19666 | 4.34E-06 | 1.3276255 | -2.1966564 | 4.34E-06 |
| ENSG00000135046 | ANXA1 | 9.8273278 | 2.28587 | 4.78E-06 | 2.0195999 | -2.28587251 | 4.78E-06 |
| ENSG00000198727 | MT-CYB | 7.7694331 | -1.47824 | 4.93E-06 | 21.773057 | 1.478236008 | 4.93E-06 |
| ENSG00000150991 | UBC | 12.29763 | 1.90361 | 5.42E-06 | 3.2989481 | -1.90360602 | 5.42E-06 |
| ENSG00000138071 | ACTR2 | 0.9719563 | -1.50237 | 5.58E-06 | 2.7678979 | 1.502367321 | 5.58E-06 |
| ENSG00000043462 | LCP2 | 1.1750516 | 2.24712 | 5.66E-06 | 0.2413864 | -2.24711909 | 5.66E-06 |
| ENSG00000127540 | UQCR11 | 0.2880638 | -1.84152 | 5.90E-06 | 1.0379617 | 1.841516512 | 5.90E-06 |
| ENSG00000123975 | CKS2 | 1.7055864 | 2.41614 | 5.93E-06 | 0.3138024 | -2.41614072 | 5.93E-06 |
| ENSG00000086061 | DNAJA1 | 3.9976625 | 1.96595 | 6.43E-06 | 1.0218693 | -1.96595342 | 6.43E-06 |
| ENSG00000155307 | SAMSN1 | 1.0237663 | -1.52071 | 6.61E-06 | 2.9529609 | 1.520708957 | 6.61E-06 |
| ENSG00000155097 | ATP6V1C1 | 0.3979011 | -1.62276 | 7.24E-06 | 1.2310709 | 1.622761219 | 7.24E-06 |
| ENSG00000171530 | TBCA | 0.3502359 | -1.64757 | 7.42E-06 | 1.1023314 | 1.647565236 | 7.42E-06 |
| ENSG00000251230 | MIR3945HG | 1.3967986 | 3.86535 | 8.07E-06 | 0.0885084 | -3.86534948 | 8.07E-06 |
| ENSG00000011465 | DCN | 12.397106 | 4.81686 | 8.17E-06 | 0.4344956 | -4.81686192 | 8.17E-06 |
| ENSG00000275302 | CCL4 | 65.04232 | 2.01141 | 8.38E-06 | 16.221169 | -2.01140647 | 8.38E-06 |
| ENSG00000204472 | AIF1 | 0.6465893 | -1.68078 | 8.42E-06 | 2.0839697 | 1.680780682 | 8.42E-06 |
| ENSG00000123983 | ACSL3 | 1.0921556 | 2.18907 | 8.42E-06 | 0.2333402 | -2.18907181 | 8.42E-06 |
| ENSG00000108106 | UBE2S | 1.7781205 | 2.0432 | 1.01E-05 | 0.4264494 | -2.0431951 | 1.01E-05 |
| ENSG00000169508 | GPR183 | 3.5355169 | 1.9944 | 1.05E-05 | 0.8850836 | -1.99439976 | 1.05E-05 |
| ENSG00000196352 | CD55 | 1.4755499 | 2.06989 | 1.12E-05 | 0.3459872 | -2.06989493 | 1.12E-05 |
| ENSG00000172757 | CFL1 | 2.3894789 | -1.38333 | 1.15E-05 | 6.2680014 | 1.383334521 | 1.15E-05 |

| | | | | | | | |
|---|---|---|---|---|---|---|---|
| ENSG00000189058 | APOD | 6.0141091 | 4.00031 | 1.47E-05 | 0.3701259 | -4.00030829 | 1.47E-05 |
| ENSG00000134531 | EMP1 | 1.4133778 | 2.51311 | 1.89E-05 | 0.2413864 | -2.51311374 | 1.89E-05 |
| ENSG00000077549 | CAPZB | 0.3979011 | -1.62276 | 2.06E-05 | 1.2310709 | 1.622761219 | 2.06E-05 |
| ENSG00000197329 | PELI1 | 1.2310065 | 2.05886 | 2.13E-05 | 0.2896637 | -2.05886078 | 2.13E-05 |
| ENSG00000167996 | FTH1 | 104.30811 | -1.32952 | 2.29E-05 | 263.70665 | 1.329522907 | 2.29E-05 |
| ENSG00000171819 | ANGPTL7 | 1.1626172 | 6.18599 | 2.38E-05 | 0.0080462 | -6.18599461 | 2.38E-05 |
| ENSG00000277632 | CCL3 | 136.68317 | 1.76012 | 2.41E-05 | 40.585108 | -1.76012414 | 2.41E-05 |
| ENSG00000165997 | ARL5B | 1.3035405 | 2.02885 | 2.49E-05 | 0.3138024 | -2.02884821 | 2.49E-05 |
| ENSG00000147872 | PLIN2 | 3.1894259 | 2.23089 | 2.52E-05 | 0.6758821 | -2.23089198 | 2.52E-05 |
| ENSG00000009413 | REV3L | 1.367785 | 2.09815 | 2.53E-05 | 0.3138024 | -2.09814665 | 2.53E-05 |
| ENSG00000168685 | IL7R | 0.6776753 | -1.73554 | 2.82E-05 | 2.2690326 | 1.735537954 | 2.82E-05 |
| ENSG00000006652 | IFRD1 | 1.2372237 | 2.02764 | 3.46E-05 | 0.29771 | -2.02764245 | 3.46E-05 |
| ENSG00000167468 | GPX4 | 1.4879843 | -1.36547 | 3.51E-05 | 3.8541369 | 1.365474351 | 3.51E-05 |
| ENSG00000144381 | HSPD1 | 4.7561614 | 2.20525 | 3.51E-05 | 1.0299155 | -2.20524796 | 3.51E-05 |
| ENSG00000169100 | SLC25A6 | 0.7253405 | -1.41792 | 3.82E-05 | 1.947184 | 1.417916999 | 3.82E-05 |
| ENSG00000178104 | PDE4DIP | 2.2361212 | -1.35745 | 3.82E-05 | 5.7610899 | 1.357445427 | 3.82E-05 |
| ENSG00000111817 | DSE | 2.7417872 | 1.7929 | 3.84E-05 | 0.7885291 | -1.79289907 | 3.84E-05 |
| ENSG00000113070 | HBEGF | 1.4133778 | 2.00788 | 4.31E-05 | 0.3459872 | -2.00787843 | 4.31E-05 |
| ENSG00000240972 | MIF | 1.894175 | -1.35398 | 4.99E-05 | 4.86796 | 1.353977766 | 4.99E-05 |
| ENSG00000170315 | UBB | 6.9715586 | 1.86107 | 5.02E-05 | 1.9230454 | -1.86106848 | 5.02E-05 |
| ENSG00000051108 | HERPUD1 | 1.5066359 | 2.00477 | 6.64E-05 | 0.3701259 | -2.00477407 | 6.64E-05 |
| ENSG00000142669 | SH3BGRL3 | 2.3956961 | -1.36285 | 6.65E-05 | 6.1955855 | 1.362845616 | 6.65E-05 |
| ENSG00000178381 | ZFAND2A | 1.1128796 | 2.47917 | 6.98E-05 | 0.1931092 | -2.47917446 | 6.98E-05 |
| ENSG00000078369 | GNB1 | 0.4061907 | -1.42441 | 7.14E-05 | 1.0942852 | 1.424411979 | 7.14E-05 |
| ENSG00000164713 | BRI3 | 1.5397943 | -1.32216 | 7.29E-05 | 3.8702294 | 1.322162241 | 7.29E-05 |
| ENSG00000243678 | NME2 | 1.1501828 | -1.38791 | 7.54E-05 | 3.0253768 | 1.387911356 | 7.54E-05 |
| ENSG00000062716 | VMP1 | 2.8516244 | 1.80646 | 8.06E-05 | 0.8126677 | -1.80645579 | 8.06E-05 |
| ENSG00000099624 | ATP5F1D | 0.3191498 | -1.66099 | 8.72E-05 | 1.0138231 | 1.660991997 | 8.72E-05 |
| ENSG00000187608 | ISG15 | 0.6362273 | -1.8911 | 8.97E-05 | 2.3736334 | 1.89109854 | 8.97E-05 |
| ENSG00000172183 | ISG20 | 1.5252875 | 2.21954 | 9.18E-05 | 0.3218486 | -2.21953709 | 9.18E-05 |
| ENSG00000005483 | KMT2E | 1.2289341 | 1.83962 | 9.29E-05 | 0.337941 | -1.83962265 | 9.29E-05 |
| ENSG00000130066 | SAT1 | 14.973101 | -1.28617 | 9.94E-05 | 36.730971 | 1.28616532 | 9.94E-05 |
| ENSG00000122705 | CLTA | 0.6942545 | -1.39532 | 9.98E-05 | 1.834537 | 1.395318081 | 9.98E-05 |
| ENSG00000145779 | TNFAIP8 | 5.1996553 | 1.67261 | 0.0001 | 1.6333816 | -1.67261459 | 0.0001018 |
| ENSG00000115009 | CCL20 | 46.912952 | 1.89415 | 0.0001 | 12.688881 | -1.89414568 | 0.0001044 |
| ENSG00000156976 | EIF4A2 | 1.3967986 | 1.86535 | 0.00011 | 0.3781721 | -1.86534948 | 0.0001054 |
| ENSG00000092841 | MYL6 | 2.7417872 | -1.26074 | 0.00012 | 6.6059424 | 1.260738892 | 0.000116 |
| ENSG00000035681 | NSMAF | 1.2310065 | 1.98291 | 0.00014 | 0.3057562 | -1.98291192 | 0.0001431 |
| ENSG00000101608 | MYL12A | 1.2662373 | -1.3131 | 0.00014 | 3.1621624 | 1.313095692 | 0.000145 |
| ENSG00000108654 | DDX5 | 7.1104095 | 1.53901 | 0.00014 | 2.4540955 | -1.53901428 | 0.000145 |
| ENSG00000038427 | VCAN | 0.2880638 | -1.80783 | 0.00015 | 1.0138231 | 1.807833385 | 0.0001542 |
| ENSG00000198743 | SLC5A3 | 0.3875391 | -1.53339 | 0.00015 | 1.1264701 | 1.533394216 | 0.0001547 |
| ENSG00000122257 | RBBP6 | 1.2268617 | 1.9059 | 0.00015 | 0.3218486 | -1.90590457 | 0.0001548 |
| ENSG00000170296 | GABARAP | 0.9346531 | -1.35707 | 0.00016 | 2.4058183 | 1.357071444 | 0.0001578 |
| ENSG00000143226 | FCGR2A | 2.2651348 | 1.70402 | 0.00016 | 0.6919745 | -1.70402181 | 0.0001589 |
| ENSG00000162783 | IER5 | 1.5605183 | 2.02503 | 0.00016 | 0.3781721 | -2.0250265 | 0.0001616 |
| ENSG00000140941 | MAP1LC3B | 3.6308474 | 1.60801 | 0.00018 | 1.1908398 | -1.60801005 | 0.0001791 |

| | | | | | | | |
|---|---|---|---|---|---|---|---|
| ENSG00000184009 | ACTG1 | 3.0671542 | -1.31543 | 0.00018 | 7.676089 | 1.315432713 | 0.0001831 |
| ENSG00000145390 | USP53 | 1.347061 | 1.93865 | 0.00018 | 0.3459872 | -1.9386504 | 0.000184 |
| ENSG00000177189 | RPS6KA3 | 0.4642179 | -1.39204 | 0.00018 | 1.2230247 | 1.392038367 | 0.0001846 |
| ENSG00000119655 | NPC2 | 1.0444903 | -1.34767 | 0.00018 | 2.6713433 | 1.347670505 | 0.0001846 |
| ENSG00000164949 | GEM | 1.0279111 | 2.83875 | 0.00019 | 0.1367857 | -2.83874533 | 0.0001874 |
| ENSG00000095794 | CREM | 2.5801398 | 1.73474 | 0.00022 | 0.7724366 | -1.7347438 | 0.0002181 |
| ENSG00000164266 | SPINK1 | 1.3408438 | 4.80646 | 0.00027 | 0.0402311 | -4.80645579 | 0.0002717 |
| ENSG00000153823 | PID1 | 0.3419462 | -1.55067 | 0.00028 | 1.0057769 | 1.550672208 | 0.0002814 |
| ENSG00000139330 | KERA | 1.2538029 | 5.70978 | 0.00029 | 0.0160924 | -5.70977977 | 0.0002883 |
| ENSG00000176340 | COX8A | 0.52639 | -1.41351 | 0.0003 | 1.4080876 | 1.413509897 | 0.0002951 |
| ENSG00000132510 | KDM6B | 2.2900036 | 1.59285 | 0.00032 | 0.7563442 | -1.59284835 | 0.0003157 |
| ENSG00000168461 | RAB31 | 0.335729 | -1.61093 | 0.00032 | 1.0299155 | 1.610930817 | 0.0003158 |
| ENSG00000184292 | TACSTD2 | 1.0693592 | 3.89566 | 0.00033 | 0.0643697 | -3.89566375 | 0.0003316 |
| ENSG00000100985 | MMP9 | 0.7232681 | -2.63933 | 0.00033 | 4.5380652 | 2.639327661 | 0.0003328 |
| ENSG00000082397 | EPB41L3 | 0.7709334 | -1.25089 | 0.00033 | 1.8425832 | 1.250889946 | 0.0003328 |
| ENSG00000167553 | TUBA1C | 1.1750516 | 1.77505 | 0.00035 | 0.337941 | -1.77505065 | 0.0003452 |
| ENSG00000170776 | AKAP13 | 1.6786452 | 1.64904 | 0.00039 | 0.5310502 | -1.6490372 | 0.0003857 |
| ENSG00000276085 | CCL3L1 | 66.68988 | 1.58878 | 0.00039 | 22.296061 | -1.58877828 | 0.0003907 |
| ENSG00000122566 | HNRNPA2B1 | 5.9788783 | 1.48033 | 0.0004 | 2.1483394 | -1.48033468 | 0.0003995 |
| ENSG00000113732 | ATP6V0E1 | 0.9657391 | -1.32908 | 0.00041 | 2.4380031 | 1.329080489 | 0.0004136 |
| ENSG00000026751 | SLAMF7 | 0.7460645 | -1.31056 | 0.00041 | 1.8586756 | 1.310557684 | 0.0004136 |
| ENSG00000117410 | ATP6V0B | 1.0817936 | -1.32716 | 0.00044 | 2.7276668 | 1.327155515 | 0.0004419 |
| ENSG00000154518 | ATP5MC3 | 0.5740552 | -1.39935 | 0.00046 | 1.5207346 | 1.399346251 | 0.0004598 |
| ENSG00000103187 | COTL1 | 0.7087613 | -1.26108 | 0.00047 | 1.7057976 | 1.26107657 | 0.0004718 |
| ENSG00000186847 | KRT14 | 1.6434144 | 6.0996 | 0.00053 | 0.0160924 | -6.09960098 | 0.0005272 |
| ENSG00000161547 | SRSF2 | 1.8029893 | 1.55131 | 0.00058 | 0.6115123 | -1.55131065 | 0.0005776 |
| ENSG00000135821 | GLUL | 2.3459585 | 1.72177 | 0.00059 | 0.7080669 | -1.721767 | 0.00059 |
| ENSG00000100650 | SRSF5 | 1.8630889 | 1.56157 | 0.00068 | 0.6276048 | -1.56156873 | 0.0006766 |
| ENSG00000131143 | COX4I1 | 1.4527534 | -1.19007 | 0.0007 | 3.331133 | 1.190072021 | 0.0007039 |
| ENSG00000109846 | CRYAB | 2.2444108 | 4.96379 | 0.00071 | 0.0643697 | -4.96379232 | 0.0007051 |
| ENSG00000170017 | ALCAM | 0.6279376 | -1.2474 | 0.00074 | 1.496596 | 1.247398662 | 0.0007448 |
| ENSG00000197989 | SNHG12 | 1.1543276 | 1.92776 | 0.00089 | 0.29771 | -1.92776208 | 0.000891 |
| ENSG00000196154 | S100A4 | 2.4226373 | -1.20529 | 0.00095 | 5.616258 | 1.205287546 | 0.0009468 |
| ENSG00000143384 | MCL1 | 3.324132 | 1.46152 | 0.00096 | 1.2069322 | -1.46152113 | 0.0009555 |
| ENSG00000011600 | TYROBP | 1.6641384 | -1.16466 | 0.00096 | 3.7495361 | 1.164658764 | 0.0009572 |
| ENSG00000164032 | H2AFZ | 5.7281177 | 1.46226 | 0.00098 | 2.0839697 | -1.46226424 | 0.0009842 |
| ENSG00000168003 | SLC3A2 | 1.7946997 | 1.50768 | 0.00102 | 0.6276048 | -1.50767572 | 0.0010228 |
| ENSG00000006451 | RALA | 0.6009964 | -1.23116 | 0.00107 | 1.4161338 | 1.231161923 | 0.001074 |
| ENSG00000197982 | C1orf122 | 0.4434939 | -1.30876 | 0.00122 | 1.1023314 | 1.308763323 | 0.0012247 |
| ENSG00000164919 | COX6C | 0.4165527 | -1.33461 | 0.00126 | 1.0540542 | 1.334614352 | 0.0012588 |
| ENSG00000124172 | ATP5F1E | 1.6268352 | -1.12121 | 0.00128 | 3.556427 | 1.121209102 | 0.001281 |
| ENSG00000255112 | CHMP1B | 1.336699 | 1.96069 | 0.00138 | 0.337941 | -1.96069388 | 0.0013818 |
| ENSG00000168209 | DDIT4 | 1.1128796 | 1.83763 | 0.00153 | 0.3057562 | -1.83762843 | 0.0015325 |
| ENSG00000176597 | B3GNT5 | 1.1108072 | 1.66091 | 0.00157 | 0.3459872 | -1.66091494 | 0.0015747 |
| ENSG00000120438 | TCP1 | 1.0714316 | 1.60894 | 0.00161 | 0.3459872 | -1.60894495 | 0.001607 |
| ENSG00000165312 | OTUD1 | 2.3190173 | 1.75459 | 0.00165 | 0.6839283 | -1.75458655 | 0.0016534 |
| ENSG00000166913 | YWHAB | 0.862119 | -1.13327 | 0.00167 | 1.8989067 | 1.133271391 | 0.0016674 |

| | | | | | | | |
|---|---|---|---|---|---|---|---|
| ENSG00000137331 | IER3 | 18.577007 | 1.29596 | 0.00168 | 7.6036731 | -1.29595978 | 0.0016806 |
| ENSG00000116679 | IVNS1ABP | 1.1564 | 1.50585 | 0.00178 | 0.4023107 | -1.50584742 | 0.0017829 |
| ENSG00000197555 | SIPA1L1 | 0.4621455 | -1.21789 | 0.00184 | 1.0781928 | 1.217892391 | 0.0018406 |
| ENSG00000125968 | ID1 | 1.094228 | 2.1918 | 0.0019 | 0.2333402 | -2.1918016 | 0.0019018 |
| ENSG00000112715 | VEGFA | 1.5750251 | 1.53586 | 0.00193 | 0.5390964 | -1.53585809 | 0.001926 |
| ENSG00000102144 | PGK1 | 0.8455398 | -1.16122 | 0.00194 | 1.8989067 | 1.161217932 | 0.001944 |
| ENSG00000198034 | RPS4X | 5.0255736 | -1.0539 | 0.00194 | 10.492264 | 1.053901972 | 0.001944 |
| ENSG00000067225 | PKM | 1.9066094 | -1.05848 | 0.00195 | 3.9909226 | 1.058476411 | 0.001947 |
| ENSG00000167460 | TPM4 | 0.8994222 | -1.17224 | 0.00202 | 2.0356924 | 1.172244812 | 0.0020181 |
| ENSG00000108518 | PFN1 | 2.5055334 | -1.06739 | 0.00211 | 5.278317 | 1.067389944 | 0.0021114 |
| ENSG00000142634 | EFHD2 | 0.4870143 | -1.27524 | 0.00225 | 1.1827936 | 1.275242032 | 0.0022474 |
| ENSG00000185215 | TNFAIP2 | 1.0113319 | -1.14917 | 0.00265 | 2.2529402 | 1.149167381 | 0.0026504 |
| ENSG00000115738 | ID2 | 3.2826839 | 1.38721 | 0.00278 | 1.2552095 | -1.38721466 | 0.0027835 |
| ENSG00000130203 | APOE | 0.2694122 | -2.62237 | 0.0028 | 1.6736127 | 2.622367846 | 0.0027997 |
| ENSG00000123416 | TUBA1B | 1.4030158 | 1.41232 | 0.00286 | 0.523004 | -1.41231563 | 0.0028619 |
| ENSG00000245532 | NEAT1 | 13.692357 | -1.03815 | 0.00303 | 28.282445 | 1.038153099 | 0.0030324 |
| ENSG00000120885 | CLU | 1.3657126 | 3.71745 | 0.0031 | 0.0965546 | -3.71745078 | 0.0030969 |
| ENSG00000138166 | DUSP5 | 1.0589972 | 1.49698 | 0.00311 | 0.3701259 | -1.49697943 | 0.0031055 |
| ENSG00000183696 | UPP1 | 1.641342 | 1.47329 | 0.00312 | 0.5873737 | -1.47329198 | 0.0031232 |
| ENSG00000162704 | ARPC5 | 0.9408703 | -1.08449 | 0.00321 | 2.0035075 | 1.084493265 | 0.0032138 |
| ENSG00000056558 | TRAF1 | 1.4672602 | 1.41269 | 0.00328 | 0.5471426 | -1.41268565 | 0.0032821 |
| ENSG00000244754 | N4BP2L2 | 0.5927068 | -1.11444 | 0.00385 | 1.2873944 | 1.114441667 | 0.0038533 |
| ENSG00000178913 | TAF7 | 1.2745269 | 1.38762 | 0.00386 | 0.4827729 | -1.38761749 | 0.0038556 |
| ENSG00000115419 | GLS | 1.662066 | 1.36097 | 0.00386 | 0.6436972 | -1.36097446 | 0.0038628 |
| ENSG00000178695 | KCTD12 | 1.2206445 | 1.70155 | 0.0039 | 0.3701259 | -1.70155058 | 0.0039 |
| ENSG00000115310 | RTN4 | 2.5677054 | 1.27241 | 0.0044 | 1.0621004 | -1.27241025 | 0.0043991 |
| ENSG00000113811 | SELENOK | 1.1957757 | 1.36851 | 0.00457 | 0.4586342 | -1.36851297 | 0.0045678 |
| ENSG00000185022 | MAFF | 1.5543011 | 1.30046 | 0.00462 | 0.6276048 | -1.30045663 | 0.0046223 |
| ENSG00000101160 | CTSZ | 1.2662373 | -1.10043 | 0.00471 | 2.7276668 | 1.100434809 | 0.0047076 |
| ENSG00000099341 | PSMD8 | 0.5512588 | -1.11655 | 0.00481 | 1.198886 | 1.116554474 | 0.0048076 |
| ENSG00000118503 | TNFAIP3 | 5.2038001 | 1.30179 | 0.00496 | 2.1161545 | -1.30179491 | 0.00496 |
| ENSG00000136527 | TRA2B | 1.4382466 | 1.40497 | 0.00523 | 0.5390964 | -1.40497461 | 0.0052299 |
| ENSG00000177156 | TALDO1 | 0.6175756 | -1.12531 | 0.00529 | 1.3517641 | 1.125309478 | 0.0052856 |
| ENSG00000108256 | NUFIP2 | 1.1315312 | 1.39208 | 0.00542 | 0.4264494 | -1.39207781 | 0.0054166 |
| ENSG00000186594 | MIR22HG | 1.5543011 | 1.30046 | 0.00567 | 0.6276048 | -1.30045663 | 0.0056722 |
| ENSG00000188612 | SUMO2 | 0.8952774 | -1.02941 | 0.00614 | 1.834537 | 1.029412289 | 0.00614 |
| ENSG00000221869 | CEBPD | 1.744962 | 1.43111 | 0.00655 | 0.6436972 | -1.4311071 | 0.0065485 |
| ENSG00000120708 | TGFBI | 0.7709334 | -1.0919 | 0.00656 | 1.6494741 | 1.091900422 | 0.0065558 |
| ENSG00000276070 | CCL4L2 | 29.426029 | 1.49755 | 0.00656 | 10.476172 | -1.49755023 | 0.0065558 |
| ENSG00000184557 | SOCS3 | 1.7573965 | 1.35492 | 0.00659 | 0.6839283 | -1.35492427 | 0.0065929 |
| ENSG00000124762 | CDKN1A | 2.4329993 | 1.23879 | 0.00668 | 1.0299155 | -1.23878607 | 0.0066838 |
| ENSG00000135390 | ATP5MC2 | 0.495304 | -1.13957 | 0.00696 | 1.0942852 | 1.139573203 | 0.0069571 |
| ENSG00000105835 | NAMPT | 5.2100173 | -0.94556 | 0.00726 | 10.089953 | 0.94556068 | 0.0072648 |
| ENSG00000067334 | DNTTIP2 | 1.2061377 | 1.30818 | 0.00783 | 0.4827729 | -1.30818302 | 0.0078329 |
| ENSG00000130255 | RPL36 | 2.5324746 | -0.92344 | 0.00877 | 4.8277289 | 0.923444208 | 0.0087688 |
| ENSG00000100219 | XBP1 | 1.2496581 | 1.24558 | 0.00909 | 0.523004 | -1.2455789 | 0.0090915 |
| ENSG00000173432 | SAA1 | 3.0567922 | 3.49158 | 0.00914 | 0.2655251 | -3.49158245 | 0.009137 |

| | | | | | | | |
|---|---|---|---|---|---|---|---|
| ENSG00000133657 | ATP13A3 | 1.6910796 | -0.94666 | 0.00915 | 3.2748095 | 0.946664789 | 0.0091462 |
| ENSG00000131236 | CAP1 | 0.5077384 | -1.1352 | 0.00926 | 1.1184239 | 1.135200227 | 0.0092569 |
| ENSG00000148926 | ADM | 1.2869613 | 1.35506 | 0.00939 | 0.4988653 | -1.35505913 | 0.0093943 |
| ENSG00000164733 | CTSB | 2.1345735 | -1.06014 | 0.00948 | 4.4736955 | 1.060136616 | 0.009484 |
| ENSG00000189043 | NDUFA4 | 0.7108337 | -1.01402 | 0.00948 | 1.4402725 | 1.014020057 | 0.009484 |
| ENSG00000118257 | NRP2 | 2.9614617 | 1.20245 | 0.00948 | 1.2873944 | -1.20245084 | 0.009484 |
| ENSG00000104904 | OAZ1 | 3.4878517 | -0.89609 | 0.01028 | 6.5254803 | 0.89609121 | 0.0102836 |
| ENSG00000105669 | COPE | 0.5201728 | -1.05861 | 0.01077 | 1.086239 | 1.058614633 | 0.0107701 |
| ENSG00000126353 | CCR7 | 1.5833147 | 1.38296 | 0.01132 | 0.6034661 | -1.38295671 | 0.0113243 |
| ENSG00000186480 | INSIG1 | 1.7408172 | -0.93637 | 0.01147 | 3.3472254 | 0.936373299 | 0.0114652 |
| ENSG00000101182 | PSMA7 | 1.336699 | -0.93045 | 0.0117 | 2.5586963 | 0.930453975 | 0.0117033 |
| ENSG00000187193 | MT1X | 1.7387448 | 1.90783 | 0.0121 | 0.4586342 | -1.90783281 | 0.0121016 |
| ENSG00000092199 | HNRNPC | 4.1054273 | 1.09742 | 0.01263 | 1.9230454 | -1.09741894 | 0.0126298 |
| ENSG00000003402 | CFLAR | 3.985228 | 1.15411 | 0.01282 | 1.7943059 | -1.15410645 | 0.0128174 |
| ENSG00000163660 | CCNL1 | 2.0060846 | 1.21703 | 0.0136 | 0.860945 | -1.21703364 | 0.0135998 |
| ENSG00000142541 | RPL13A | 4.3416811 | -0.88223 | 0.01363 | 8.0462149 | 0.882234973 | 0.0136295 |
| ENSG00000125538 | IL1B | 106.42196 | 1.09171 | 0.014 | 50.224473 | -1.09170547 | 0.013999 |
| ENSG00000170144 | HNRNPA3 | 1.4859119 | 1.14709 | 0.01481 | 0.6678358 | -1.1470909 | 0.0148075 |
| ENSG00000233927 | RPS28 | 4.0018073 | -0.87176 | 0.01496 | 7.3622866 | 0.871756125 | 0.0149557 |
| ENSG00000060138 | YBX3 | 1.6496316 | 1.16644 | 0.01517 | 0.7322056 | -1.16644224 | 0.015167 |
| ENSG00000152795 | HNRNPDL | 1.3139025 | 1.21243 | 0.01537 | 0.563235 | -1.21243395 | 0.0153715 |
| ENSG00000122862 | SRGN | 20.367562 | 1.02323 | 0.01583 | 10.073861 | -1.02322651 | 0.0158287 |
| ENSG00000131669 | NINJ1 | 1.8112789 | -0.92349 | 0.01589 | 3.4518262 | 0.923485358 | 0.0158888 |
| ENSG00000277443 | MARCKS | 2.4682302 | -0.89919 | 0.01615 | 4.6265736 | 0.899188197 | 0.0161461 |
| ENSG00000175793 | SFN | 1.114952 | 3.42527 | 0.01654 | 0.0965546 | -3.42527003 | 0.0165402 |
| ENSG00000136888 | ATP6V1G1 | 1.367785 | -0.91532 | 0.01675 | 2.5908812 | 0.915315609 | 0.0167526 |
| ENSG00000163734 | CXCL3 | 46.164815 | 1.1052 | 0.01711 | 21.579948 | -1.10520429 | 0.0171089 |
| ENSG00000127184 | COX7C | 0.5782 | -0.99906 | 0.01738 | 1.1586549 | 0.999057789 | 0.0173788 |
| ENSG00000167283 | ATP5MG | 0.6776753 | -0.93073 | 0.01814 | 1.2954406 | 0.930729714 | 0.0181436 |
| ENSG00000112308 | C6orf62 | 0.7377749 | -0.92808 | 0.01834 | 1.4080876 | 0.92808307 | 0.0183428 |
| ENSG00000161960 | EIF4A1 | 5.2638998 | 1.03183 | 0.01898 | 2.582835 | -1.03183207 | 0.0189779 |
| ENSG00000136758 | YME1L1 | 1.2683097 | 1.20279 | 0.01906 | 0.5471426 | -1.20278709 | 0.019061 |
| ENSG00000102760 | RGCC | 2.2319764 | 1.23089 | 0.0195 | 0.9494534 | -1.23089198 | 0.0194992 |
| ENSG00000112149 | CD83 | 4.1593098 | 1.07474 | 0.01978 | 1.9793689 | -1.07474461 | 0.0197848 |
| ENSG00000139318 | DUSP6 | 2.1366459 | 1.25544 | 0.01979 | 0.8931299 | -1.25544062 | 0.0197868 |
| ENSG00000156467 | UQCRB | 0.505666 | -1.02302 | 0.0204 | 1.0299155 | 1.023021032 | 0.0203966 |
| ENSG00000162434 | JAK1 | 0.6610961 | -0.93939 | 0.02088 | 1.271302 | 0.939386576 | 0.0208797 |
| ENSG00000163191 | S100A11 | 6.499051 | -0.85486 | 0.02135 | 11.81989 | 0.854862341 | 0.021346 |
| ENSG00000065978 | YBX1 | 3.3469284 | -0.81574 | 0.02245 | 5.9220142 | 0.815741042 | 0.0224453 |
| ENSG00000219200 | RNASEK | 1.8506545 | -0.84474 | 0.02313 | 3.3391792 | 0.844740413 | 0.0231305 |
| ENSG00000197111 | PCBP2 | 0.8766258 | -0.85714 | 0.02431 | 1.5931505 | 0.857135882 | 0.024307 |
| ENSG00000139679 | LPAR6 | 1.1564 | 1.68642 | 0.02461 | 0.3540335 | -1.68641966 | 0.0246077 |
| ENSG00000173674 | EIF1AX | 1.2890337 | 1.08673 | 0.02542 | 0.6034661 | -1.08672912 | 0.0254239 |
| ENSG00000136238 | RAC1 | 1.5045635 | -0.8435 | 0.02568 | 2.7115744 | 0.843499598 | 0.0256762 |
| ENSG00000157557 | ETS2 | 3.659861 | 1.01211 | 0.02604 | 1.8184446 | -1.01210612 | 0.0260433 |
| ENSG00000153187 | HNRNPU | 2.9096517 | 1.05671 | 0.02613 | 1.4000414 | -1.05671159 | 0.0261262 |
| ENSG00000125835 | SNRPB | 1.357423 | 1.05157 | 0.02655 | 0.6517434 | -1.05156828 | 0.0265513 |

| | | | | | | | |
|---|---|---|---|---|---|---|---|
| ENSG00000110651 | CD81 | 0.7564265 | -0.87568 | 0.02675 | 1.3919952 | 0.875675373 | 0.0267469 |
| ENSG00000143774 | GUK1 | 0.6942545 | -0.89596 | 0.02682 | 1.2954406 | 0.895964296 | 0.0268192 |
| ENSG00000146278 | PNRC1 | 9.4356439 | 0.99017 | 0.02712 | 4.7714054 | -0.99016774 | 0.0271239 |
| ENSG00000169567 | HINT1 | 1.1895585 | -0.85783 | 0.02819 | 2.1644318 | 0.857829167 | 0.0281852 |
| ENSG00000148834 | GSTO1 | 1.3076853 | -0.84922 | 0.03144 | 2.3655872 | 0.849222112 | 0.0314359 |
| ENSG00000100902 | PSMA6 | 0.7812954 | -0.88604 | 0.0316 | 1.4483187 | 0.886035178 | 0.0316006 |
| ENSG00000107372 | ZFAND5 | 1.8796681 | 1.05794 | 0.0316 | 0.9011761 | -1.05793781 | 0.0316006 |
| ENSG00000120690 | ELF1 | 1.1398208 | 1.07001 | 0.0321 | 0.5390964 | -1.07001395 | 0.0321018 |
| ENSG00000143119 | CD53 | 1.1584724 | 1.07233 | 0.03237 | 0.5471426 | -1.07232685 | 0.0323726 |
| ENSG00000163131 | CTSS | 0.8745534 | -0.88925 | 0.03327 | 1.6253354 | 0.88925378 | 0.0332694 |
| ENSG00000011422 | PLAUR | 10.809646 | 0.98906 | 0.03366 | 5.4714261 | -0.98906201 | 0.0336646 |
| ENSG00000204592 | HLA-E | 2.2216144 | 0.98972 | 0.03419 | 1.1184239 | -0.98971963 | 0.0341853 |
| ENSG00000265972 | TXNIP | 2.4060581 | 1.18957 | 0.03645 | 1.0540542 | -1.18956852 | 0.0364468 |
| ENSG00000128272 | ATF4 | 1.9957226 | 0.99867 | 0.03672 | 0.9977306 | -0.99867334 | 0.036719 |
| ENSG00000113580 | NR3C1 | 1.0175491 | 1.11144 | 0.03718 | 0.4666805 | -1.11143974 | 0.0371823 |
| ENSG00000188042 | ARL4C | 0.8331054 | -0.94485 | 0.03912 | 1.609243 | 0.944847378 | 0.0391244 |
| ENSG00000135047 | CTSL | 8.1051622 | 1.02236 | 0.03916 | 4.007015 | -1.02236293 | 0.0391592 |
| ENSG00000188313 | PLSCR1 | 1.094228 | 1.09869 | 0.04138 | 0.5069115 | -1.0986922 | 0.0413797 |
| ENSG00000067560 | RHOA | 1.0880108 | -0.80433 | 0.04157 | 1.9069529 | 0.80433049 | 0.0415717 |
| ENSG00000101439 | CST3 | 5.5871944 | -1.20234 | 0.042 | 12.930267 | 1.202343601 | 0.0419998 |
| ENSG00000131981 | LGALS3 | 4.2152646 | -0.82335 | 0.04237 | 7.4990723 | 0.823351907 | 0.0423743 |
| ENSG00000127314 | RAP1B | 1.4216674 | -0.80039 | 0.04466 | 2.4862804 | 0.800389832 | 0.0446644 |
| ENSG00000104267 | CA2 | 0.5077384 | -1.03934 | 0.04513 | 1.0460079 | 1.039340212 | 0.0451269 |
| ENSG00000176845 | METRNL | 1.1398208 | 1.09139 | 0.04515 | 0.5310502 | -1.0913876 | 0.045152 |
| ENSG00000132475 | H3F3B | 9.8501242 | 0.91335 | 0.04614 | 5.2541783 | -0.91334765 | 0.0461436 |
| ENSG00000123610 | TNFAIP6 | 0.8579742 | -1.03937 | 0.0478 | 1.7701673 | 1.039366749 | 0.0477981 |
| ENSG00000140264 | SERF2 | 2.2050352 | -0.74977 | 0.04913 | 3.7253975 | 0.749774996 | 0.0491349 |
| ENSG00000118816 | CCNI | 1.0921556 | -0.78668 | 0.04947 | 1.8908605 | 0.786680646 | 0.0494659 |
